# Development of novel PCR primer sets for DNA metabarcoding of aquatic insects, and the discovery of some cryptic species

**DOI:** 10.1101/2021.11.05.467390

**Authors:** Masaki Takenaka, Koki Yano, Tomoya Suzuki, Koji Tojo

**Author notes:** **Co-Correspondence:** Masaki Takenaka and Koji Tojo.

## Abstract

DNA barcoding is a powerful tool that provides rapid, accurate, and automatable species identification by using standardized genetic region(s). It can be a powerful tool in various fields of biology such as for revealing the existence of cryptic species and/or rare species and in environmental science such as when monitoring river biota. Biodiversity reduction in recent times has become one of the most serious environmental issues on a worldwide scale. DNA barcoding techniques require the development of sets of universal PCR primers for DNA metabarcoding. We tried to develop universal primer sets for the DNA barcoding of all insect groups. In this study, we succeeded in designing not only universal primer sets for DNA barcoding regions of almost all insects, which were designed to include a hypervariable site between highly conserved sites, but also primer sets for longer fragment sequences for registration in a database. We confirmed successful amplification for 14 orders, 43 families, and 68 species with DNA barcoding in the mtDNA 16S rRNA region, and for 13 orders, 42 families, and 66 species with DNA barcoding in the mtDNA 12S rRNA region. A key feature is that the DNA fragments of the DNA barcoding regions amplified by these primer sets are both short at about 200-bp, and longer fragment sequences will increase the level of data registration in the DNA database. Such resulting database enhancements will serve as a powerful tool for increasingly accurate assessment of biodiversity and genetic diversity.

## Introduction

About 8.7 million eukaryotic species are estimated to inhabit the Earth (Mora et al. 2011). Insects are the largest and most diverse group of organisms on Earth, and about one million species have been described (Grimaldi and Engel 2005; Tojo et al. 2017; Stork 2018). It is known that there are still many undescribed insect species, and new species are still described on a daily basis. A more accurate understanding of species diversity and elucidation of the mechanisms of diversity are important issues. On the other hand, as many species have been evaluated to be threatened with extinction, environmental conservation and species conservation efforts are also urgent tasks (Ceballos et al. 2015; Dirzo et al. 2014). For effective conservation measures of a particular species, it is important to appropriately assess and understand the current state of its biodiversity.

DNA barcoding is a system which provides rapid, accurate, and automatable species identification by using a standardized genetic region(s) (Hebert and Gregory 2005). In general, numerous insects are identified based on their morphological characteristics, but this method requires specialist knowledge and it takes a lot of time to acquire enough skills. Under such circumstances, DNA barcoding can rapidly identify a species by sequencing a standardized short DNA fragment, even if the specimens are difficult to identify by morphology (Hebert and Gregory 2005; Miya et al. 2015). In addition, DNA barcoding even allows species identification of specimens that are not suitable for species identification by means of traditional morphological classification, such as larval specimens or parts of specimens (incomplete specimens). Since this method is easy and fast, and its results are highly reproducible, it is possible use it for a wide range of species to undertake long-term monitoring and gain an understanding of their biodiversity (Hänfling et al. 2016; Uchida et al. 2020; Chucholl et al. 2021).

DNA barcoding is also an effective tool for identifying the existence of cryptic species and/or rare species (Hebert et al. 2004). In recent years, it has been reported that many cryptic species or undescribed species have been being discovered by conducting DNA barcoding-based genetic analyses (Vuataz et al. 2013; Saitoh et al. 2015; Struck et al. 2018; Takenaka and Tojo 2019; Yano et al. 2019; Ohnishi et al. 2021; Tojo et al. 2021). Of course, DNA barcoding does not replace traditional taxonomy (Schindel 2005). There is no doubt that highly experienced taxonomists are still required to scrutinize the taxonomic descriptions of such assessed species. As an interesting example of the use of DNA barcoding in recent studies, it was possible to understand past biodiversity by detecting the DNA of a particular fish from hundreds of years ago collected from seafloor sediments (Kuwae et al. 2020). Kudoh et al. (2020) identified a particular herbivorous insect using leaves with external foliage feeding marks and environmental DNA (eDNA) techniques. Moreover, for endangered species, DNA based non-invasive assessment of biodiversity and corresponding genetic diversity is a breakthrough technique (Sekiya et al. 2017; Ahn et al. 2020; Yamazaki et al. 2020). Such methods are also expected to be applied to various other fields in addition to taxonomy.

For aquatic organisms, eDNA in aquatic environments has also facilitated the detection of an aquatic vertebrate species (Miya et al. 2015). It can be a powerful tool in the various fields of biology and environmental science such as in monitoring river biota. Biodiversity reduction in recent times has become one of the most serious environmental issues on a worldwide scale. In particular, freshwater organisms tend to account for a high proportion of Red List species. Under such circumstances, it is necessary to monitor biological fauna and flora to facilitate the conservation of biodiversity. For this purpose, eDNA analysis offers a powerful molecular tool capable of non-invasively surveying species richness within many ecosystems (Bohmann et al. 2014; Deiner et al. 2017; Uchida et al. 2020).

These techniques require the development of sets of universal PCR primers for DNA metabarcoding. For fish species, Miya et al. (2015) designed a set of universal PCR primers (i.e., “MiFish”) for the metabarcoding of eDNA. Also, these primers have been developed for various other animal taxa (e.g., “MiMammal” for mammals: Ushio et al. 2017; “MiBird” for birds: Ushio et al. 2018; “MiDeca” for crustaceans, especially Decapoda: Komai et al. 2019). As for insects, which have the highest species diversity on the Earth, the mitochondrial DNA (mtDNA) cytochrome c oxidase subunit I (COI) region is frequently targeted using Folmer’s universal primer set for DNA barcoding (Folmer et al. 1994). However, it is not a suitable primer set for DNA metabarcoding, as the protein-coding gene for amino acids of the mtDNA COI region is not highly conserved (Deagle et al. 2014); its third codon in particular is detected with a high number of polymorphisms. As such polymorphisms tend to be concentrated on the third base of each codon, we considered that it was not suitable for the development of a highly versatile primer set for amplification of short fragment sequences.

Many previous studies on non-insect groups using metabarcoding of eDNA have used primer sets developed from within their ribosomal RNA region (Miya et al. 2015; Ushio et al. 2017, 2018; Komai et al. 2019). Therefore, we also tried to design a universal primer set suitable for DNA metabarcoding of insects based on the ribosomal RNA region on the mtDNA.

The ideal characteristics of DNA fragments for a DNA barcoding region are listed below (Valentini et al. 2009; Miya et al. 2015). It must be 1) possible to reliably identify the specific insect species from it, so it needs to be completely the same or with only a minimal difference from other individuals of the same species, but with clear differences to the sequences of other species, 2) a homologous standardized region that is also able to be used for amplification in all insect groups, 3) in a target region which has sufficient phylogenetic information to easily assign undescribed species to a taxon (genus or family), 4) a highly preserved, reliable, robust fragment, 5) suitable for amplification of a short fragment and contain sufficient sequence variations in order to correctly assign the insect species.

Therefore, in this study, we tried to develop universal primer sets for DNA barcoding of all insect groups, by the methods set our below and as in Miya et al. (2015). Firstly, the primer sets developed in this study are applicable to all insect groups (especially aquatic insects), and the region contained between these versatile primers includes polymorphism-rich sites (hypervariable regions). Second, although the target region is a short-length sequence (about 200 bp), it is able to reliably distinguish species, even closely related species, and is also effective at capturing fragmented DNA such as samples recovered from eDNA present in an aquatic environment. However, it takes a lot of effort and is expensive to enrich a database that refers to the region amplified by using our newly designed primer sets. In order to enhance the database more efficiently, we designed versatile universal primer sets that amplify not only short fragments for DNA barcoding, but also longer fragments including a targeted barcoding region that can also be used for phylogenetic analyses. We consider that this primer set that can also amplify longer fragments is an ideal tool for phylogenetic studies; it will therefore be adopted as the optimal method and lead to the enhancement of databases that refer to eDNA.

We also examined whether the DNA region we selected for DNA metabarcoding contained sufficient polymorphisms to be effective in species differentiation even between closely related species, and also whether cryptic species could be readily detected from it. In aquatic insects, it is known that closely related species are niche-differentiated, each adapting to various river microhabitats (Ohgitani and Nakamura 2008; Ohgitani et al. 2021; Okamoto and Tojo 2021; Okamoto et al. 2021). Heptageniid mayflies are a typical group exhibiting niche differentiation between closely related species (Ohgitani and Nakamura 2008; Tojo 2010) and are therefore suitable for testing the newly developed primers in this study. Previous studies using molecular markers have reported cases of discovering undescribed species and/or cryptic species as a result of phylogenetic analysis of species inhabiting a wide range and/or a variety of environments (Ueda et al. 2012; Yano et al. 2019). Also, the study it based on the hypothesis that *Epeorus aesculus* (Heptageniidae) also contains a cryptic species because *Epeorus aesculus* Imanishi, 1934, inhabit in a relatively wide range of river flow (Ohgitani and Nakamura 2008). From these viewpoints, we assessed and verified the detection ability and sensitivity of this newly developed DNA metabarcoding primer set using heptageniid mayflies, including *E. aesculus*.

## Materials and Methods

### Development of primer sets

In order to perform DNA metabarcoding, we focused on the mtDNA 16S rRNA and 12S rRNA regions because these ribosomal RNA regions have been reported to provide almost the same potential to correctly identify different individual species as the mtDNA COI region, which is the standard DNA barcoding region (Collins et al. 2019). We considered that these rRNA regions also have the advantage of having fewer intraspecific polymorphisms than the COI region. Also, for previous studies of other vertebrates and invertebrates, primer sets have been developed for DNA metabarcoding in the 16S rRNA or 12S rRNA regions (Miya et al. 2015; Ushio et al. 2017, 2018; Komai et al. 2019). The general DNA barcoding region for insects is the mtDNA COI region, but this could not be as effective as the versatile primers sets in this study due to the presence of polymorphisms every three bases.

In order to select a few suitable regions, whole or partial mitogenome sequences of various insect groups, to increase versatility, were downloaded from GenBank: aquatic insects [Ephemeroptera, Odonata, Plecoptera, each family of Hemiptera (Hemiptera s. lat.: Belostomatidae, Nepidae, Gerridae, Corixidae), Corydalidae, Trichoptera, each family of Coleoptera (Dytiscidae, Gyrinidae, Lampyridae, Dryopoidae), and each family of Diptera (Simuliidae, Culicidae, Tipulidae)], and Apterygota (Diplura, Archaeognatha, Zygentoma). Initially, we referred only to the sequences of aquatic insects because we were focusing on the DNA barcoding of aquatic insects. However, since aquatic insects include a wide range of insect groups, the results were used to search for a genetic region that could be applied to almost all insect groups. All sequences were aligned using MAFFT v7.222 (Katoh & Standley 2013) with the default set of parameters. The highly versatile areas were graphically represented using MEGA 7.0.26 (Kumar et al. 2016) and highly versatile regions were identified by means of visual inspection.

For the mtDNA 16S rRNA region, we searched for a highly versatile region using all data sets of all insect groups downloaded. However, it was not possible to design a single set for all insects in the highly versatile mtDNA 12S rRNA region for amplification; therefore, we designed three specialized primer sets to amplify the mtDNA 12S rRNA region for each of the three groups: 1) Hemimetabola, 2) Holometabola excluding Trichoptera, and 3) Trichoptera. In designing these generic primers, we applied the following recommendations made by Miya et al. (2015), paying attention to both ends of each primer so that not only the complementarity on the 3’-end, but also the region with high complementarity on the 5ʼ-end were included.

### Testing the versatility of the newly developed primers

To evaluate the versatility of the primers designed in this study, PCR amplification was conducted using the total genomic DNA extracted and purified from a variety of insect groups stored in the Tojo laboratory of Shinshu University, Japan (Table 1). Each total genomic DNA sample was used to amplify DNA fragments [the mtDNA 16S rRNA and 12S rRNA regions] by polymerase chain reaction (PCR) with sets of primers designed in this study. Regarding each reaction, 1.0 μL of 10x Ex Taq buffer, 0.8μL dNTP Mixture (included 25 mM MgCl_2_), 0.05μL of 5U/μL Ex Taq polymerase (TAKARA, Shiga), 0.25μL of each primer, 1.0μL of extracted DNA in total 10μL. The PCR protocol for the DNA barcoding region in the mtDNA 16S rRNA and 12S rRNA regions was: 94 °C for 1 min; 30× (94 °C for 1 min, 50 °C for 30 sec or 1min, 72 °C for 30 sec); 72 °C for 3 min. The PCR protocol for the long fragment in the mtDNA 16S rRNA and 12S rRNA regions was: 94 °C for 1 min; 30× (94 °C for 1 min, 51 °C for 30 sec, 74 °C for 1 min); 74 °C for 3 min. The PCR products were purified using ExoSAP-IT Express (Thermo Fisher Scientific K.K., Tokyo, JP). Sequencing of purified DNA fragments was outsourced to Eurofins Genomics (Tokyo, Japan). A BigDye Terminator Cycle Sequence Kit v3.1 (ABI) was used prior to sequencing with the ABI sequencer. Sequence data have been submitted to the DNA data-bank of Japan (DDBJ database; Accession numbers are given in Table 1). All sequence data were aligned using MAFFT v7.222 (Katoh and Standley 2013). Phylogenetic analyses were performed by the Neighbor-Joining (NJ) method using MEGA 7.0.26 (Kumar et al. 2016).

**Table 1.**
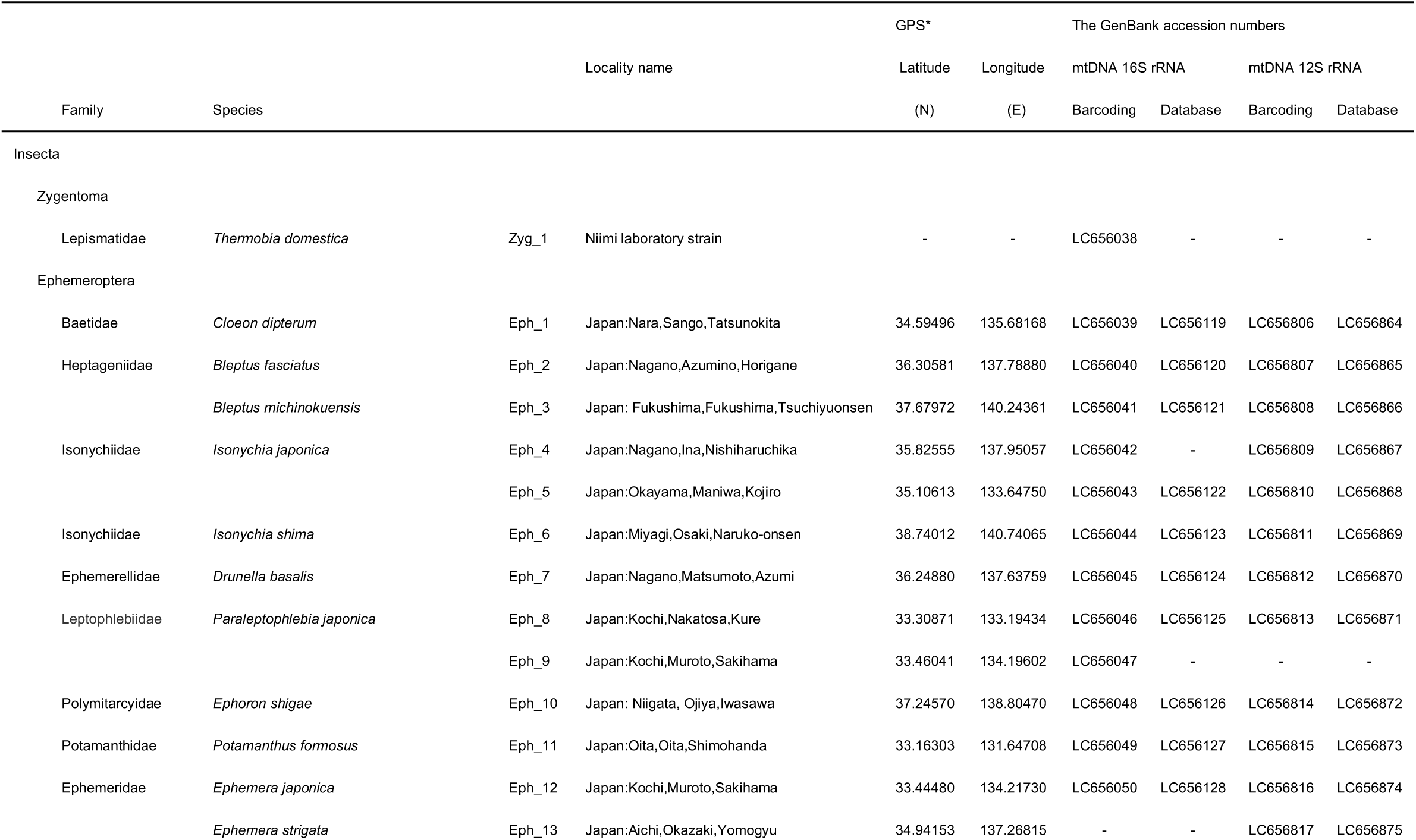
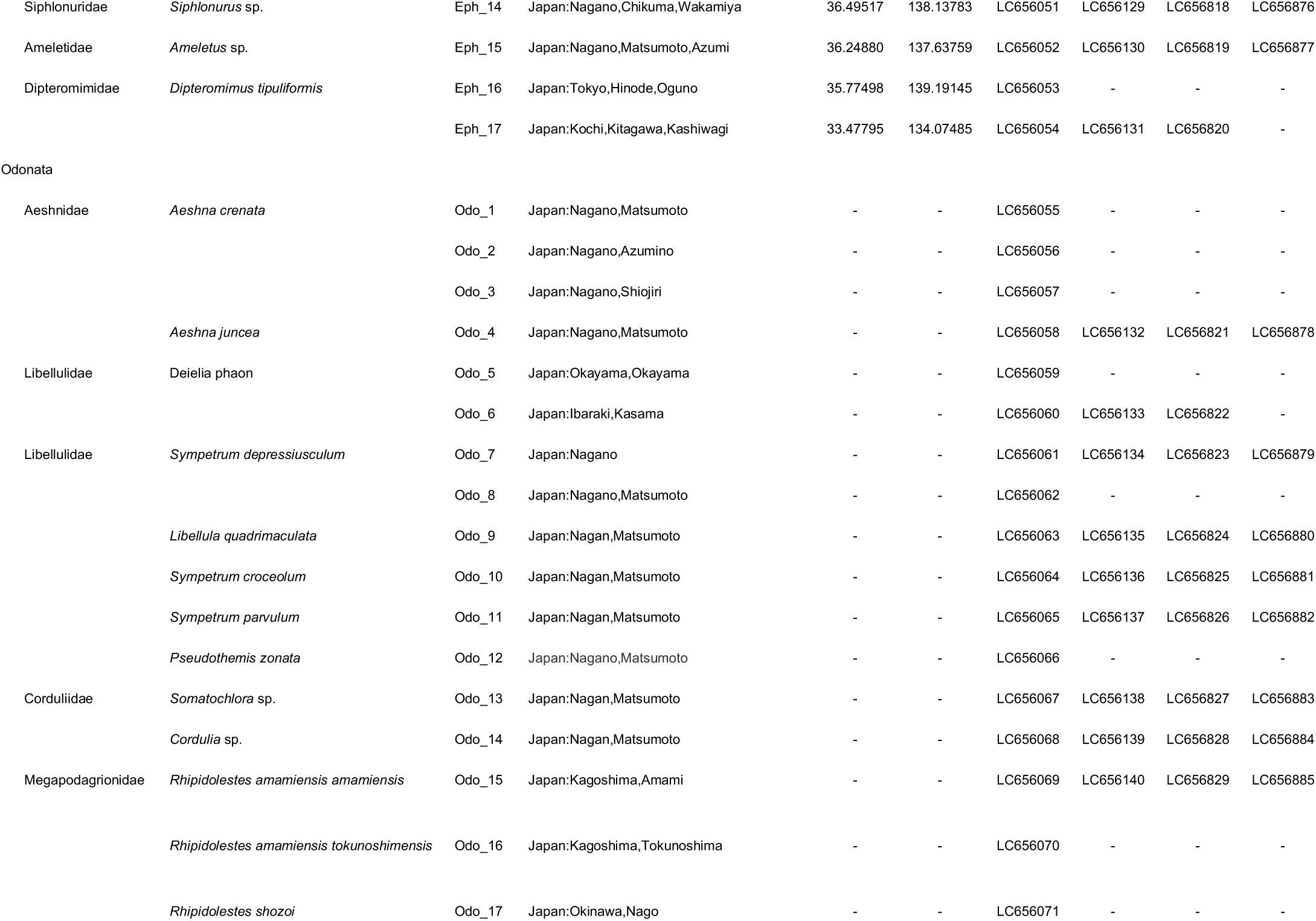
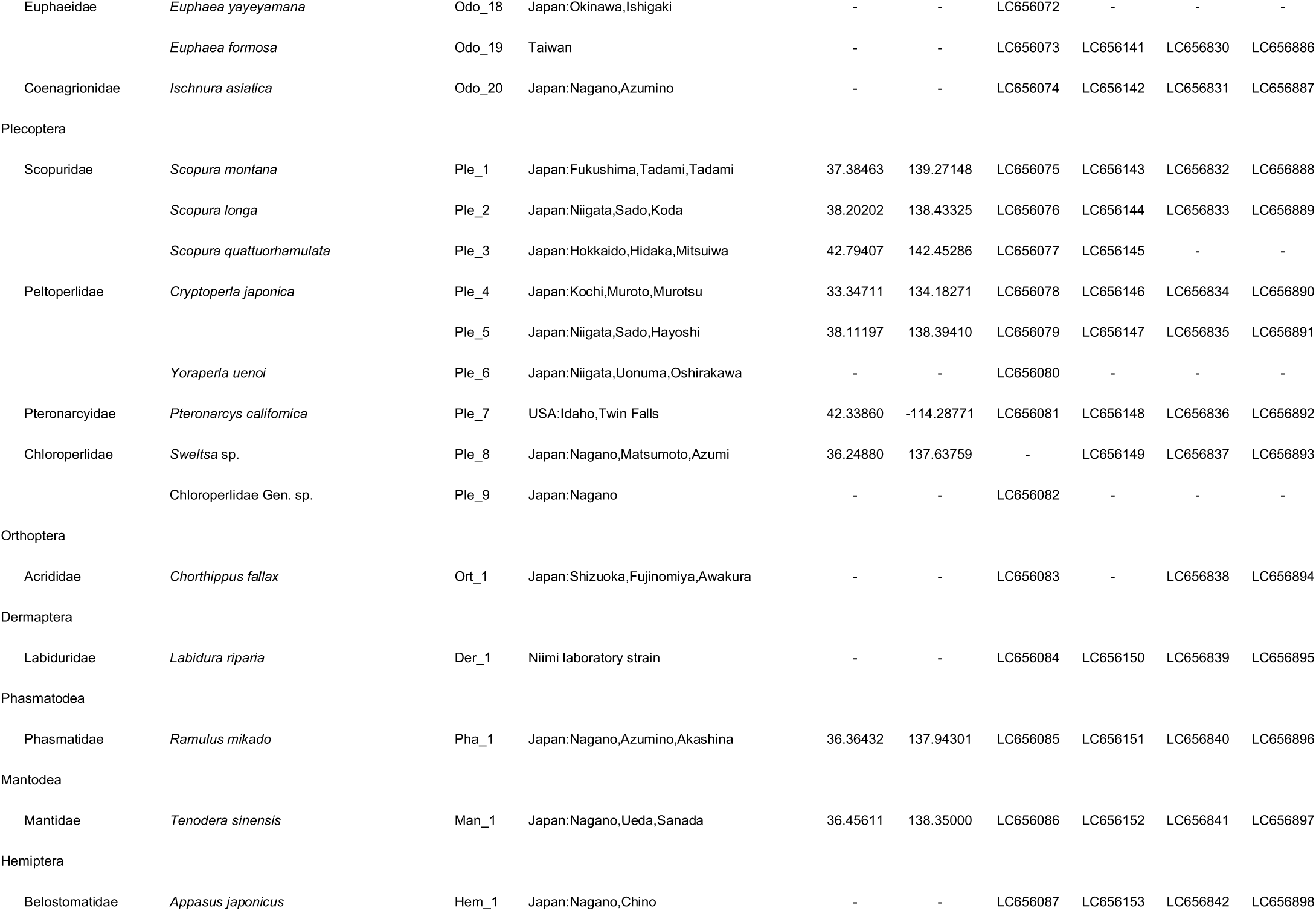
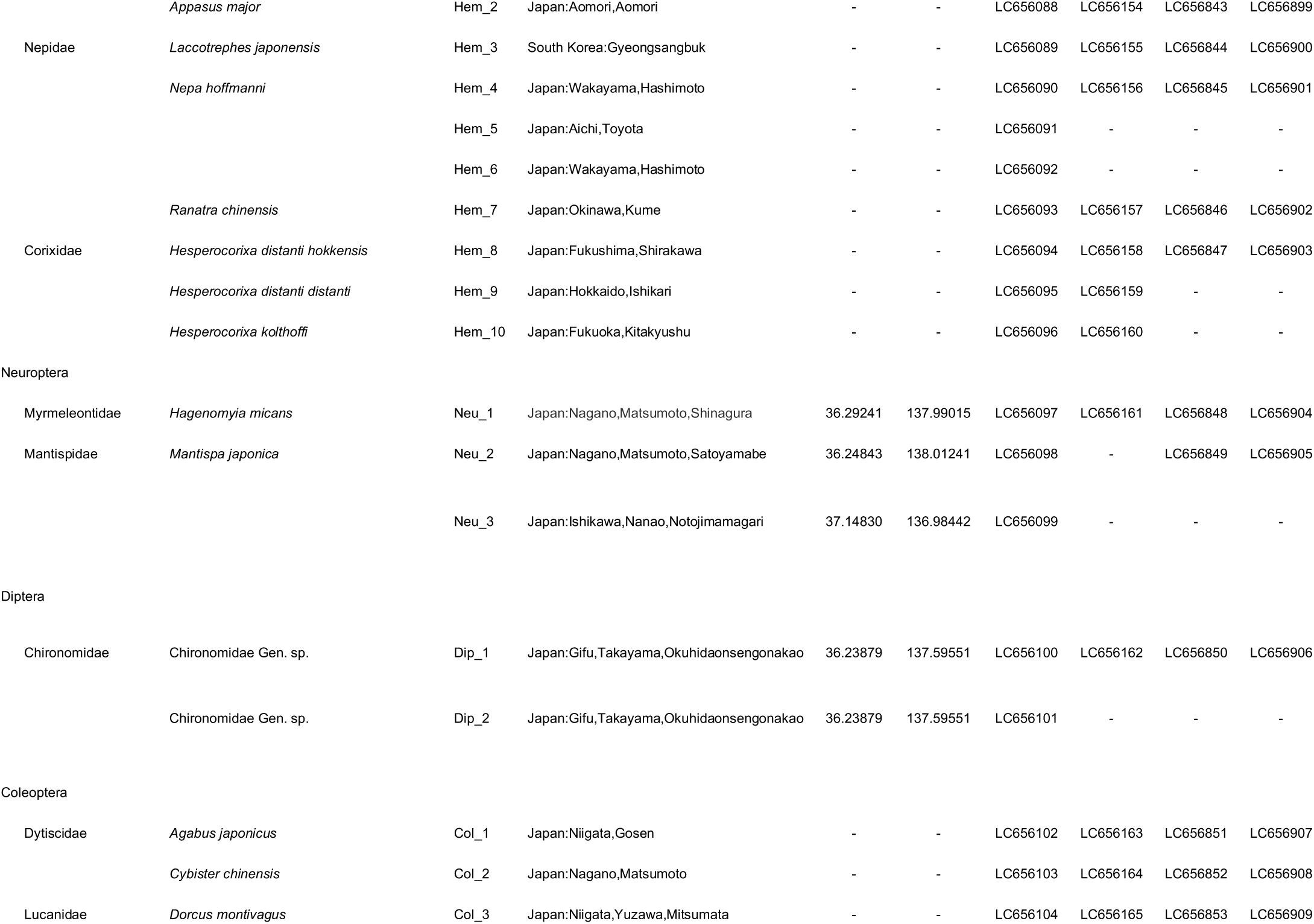
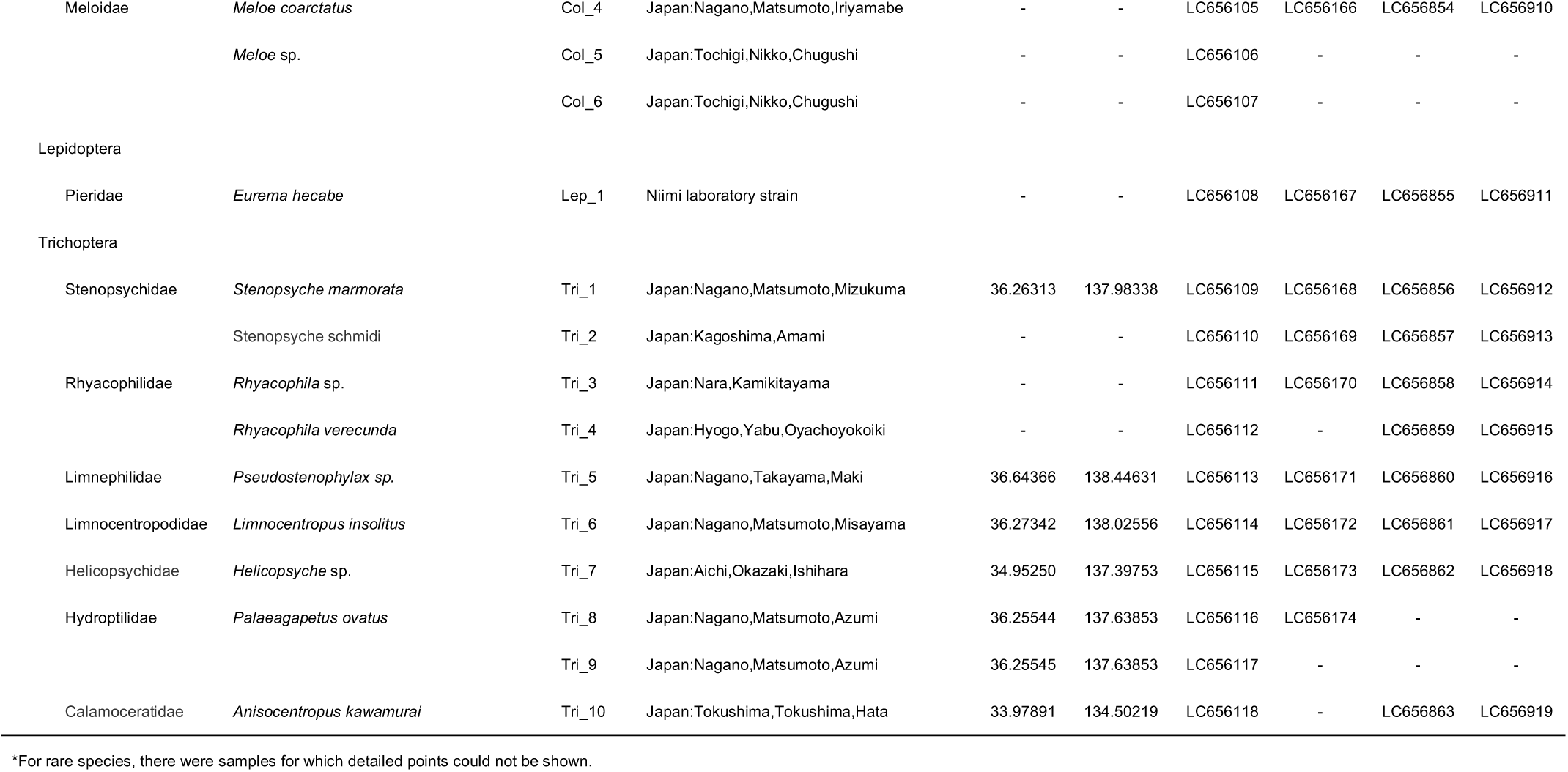
List of specimens examined in this study, sequence types, and the GenBank accession numbers

### Evaluation of interspecific variations and phylogenetic analysis

To check for genetic variation between closely related species and within a species (detection of cryptic species/lineages), this study focused on *Epeorus aesculus* (Heptageniidae, Ephemeroptera). We used the total genomic DNA of 14 specimens of *E. aesculus* Imanishi, 1934, from four localities including topotype specimens (i.e., specimens collected from the type locality: Kurobe-goro-zawa, Toyama, Toyama Prefecture) (Table S1).

With respect to the outgroups, we added appropriate DNA sequence data on *Epeorus dayongensis* (MK6422986, MT112895), *Epeorus herklotsi* (MG870104, NC_039612), *Epeorus carinatus* (MT112896), and *Afronurus yixingensis* (MK642297). Each total genomic DNA sample was used to amplify DNA fragments [the mtDNA 16S rRNA and 12S rRNA regions] by polymerase chain reaction (PCR) with sets of primers designed in this study. The PCR products were purified using ExoSAP-IT Express (Thermo Fisher Scientific K.K., Tokyo, JP). Sequencing of purified DNA fragments was outsourced to Eurofins Genomics (Tokyo, Japan). A BigDye Terminator Cycle Sequence Kit v3.1 (ABI) was used prior to sequencing with the ABI sequencer. Sequence data have been submitted to the DNA data-bank of Japan (DDBJ database; Accession numbers are given in Table S1).

Sequence alignment and editing were performed using the same methods for each gene separately using ATGC bundled with GENETYX ver. 15.2 (GENETYX Corporation). All sequence data were aligned using MAFFT v7.222 (Katoh and Standley, 2013). Phylogenetic analyses were performed by Bayesian analysis using MrBayes v3.2.6 (Ronquist et al. 2012). The program Kakusan4 (Tanabe 2007) was used to select appropriate models based on Schwarz’s Bayesian Information Criterion (BIC; Schwarz, 1978). Best-fit substitution models were chosen as follows: HKY + G for the mtDNA 16S rRNA; HKY + G for the mtDNA 12S rRNA regions. Bayesian MCMC simulations were run for 10 million generations, sampling every 1000 generations. The output files were checked for convergence after removing a 10% burn-in by examining Effective Sampling Size (ESS > 200) using Tracer v1.6 (Rambaut et al. 2014). Then, this was visualized in the resulting tree created by FigTree v1.3.1 (Rambaut 2009).

## Results and Discussion

### Design of versatile primer sets for DNA metabarcoding

We designed a primer set, “MtInsects-16S”, for amplification of the DNA barcoding regions in the mtDNA 16S rRNA region that is applicable to almost all insect groups. We also designed a primer “AQdb-16S” that can be used to PCR-amplify longer DNA fragments containing the DNA barcoding region and this is highly effective for phylogenetic analyses. This primer set will be very effective and useful when registering reference sequences. (Table 2; Fig. 1). The positional relationship of each primer is shown in Figure S1.

**Table 2.**
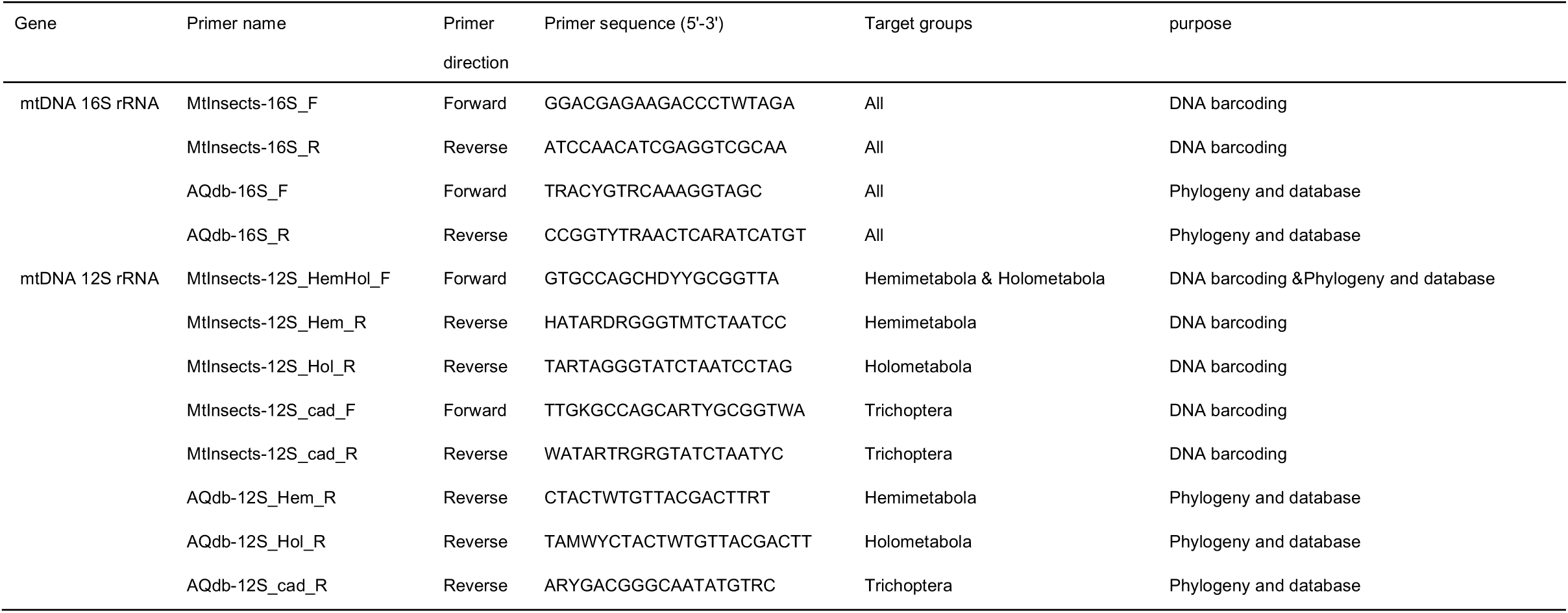
Information on the newly designed primer sets in this study

**Figure 1.**
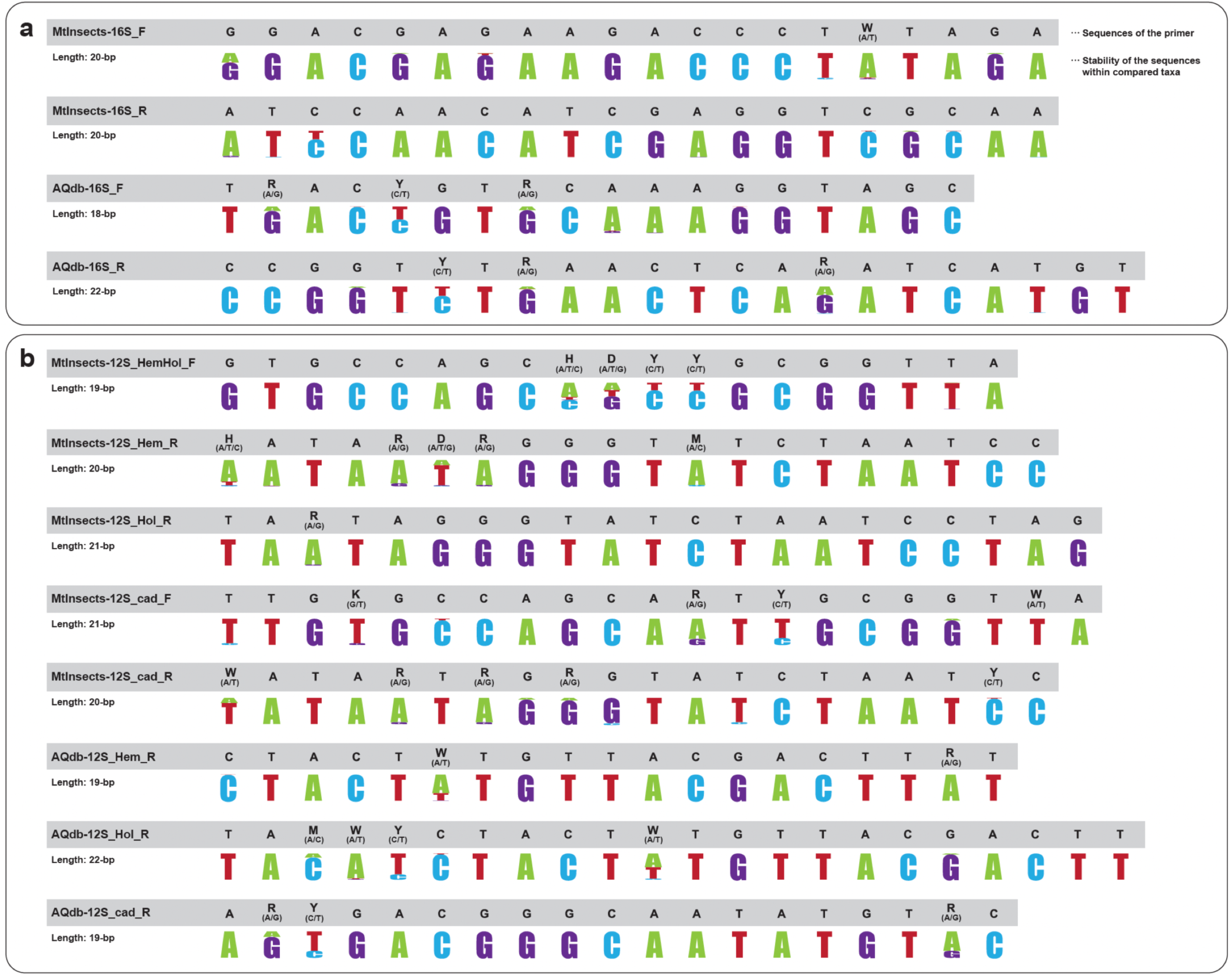
Concordance rate of primers site sequences. Gray bands show the sequences of each primer, while under gray bands show stability of the sequence within compared taxa.

On the other hand, in the mtDNA 12S rRNA region, we designed primer sets that amplified the DNA barcoding region for each of three groups: 1) Hemimetabola, 2) Holometabola excluding Trichoptera, and 3) Trichoptera. For Hemimetabola, “MtInsects-12S_HemHol_F” and “MtInsects-12S_Hem_R” were used as the primer set to amplify the DNA barcoding region. For Holometabola excluding Trichoptera, “MtInsects-12S_HemHol_F” (the same as Hemimetabola) and “MtInsects-12S_Hol_R” were used as the primer sets to amplify the DNA barcoding region. For Trichoptera, “MtInsects-12S_cad_F” and “MtInsects-12S_cad_R” were used as the primer sets to amplify the DNA barcoding region (Table 2; Fig. 1).

We also designed three primer sets to amplify longer fragments including each of the three DNA barcoding regions of each group: 1) Hemimetabola, 2) Holometabola excluding Trichoptera, and 3) Trichoptera. In Hemimetabola, “MtInsects-12S_HemHol_F” (the same as DNA barcoding) and “AQdb-12S_Hem_R” were used as a primer set; in Holometabola excluding Trichoptera, “MtInsects-12S_HemHol_F” (the same as DNA barcoding) and “AQdb-12S_Hol_R”; were used as a primer set, and in Trichoptera, “MtInsects-12S_cad_F” and “AQdb-12S_cad_R” were used as a primer set (Table 2; Fig. 1). The positional relationship of each primer is shown in Figure S2.

The above primer information is shown in Table 2. All primers were designed to put a hypervariable region between highly conserved regions (Fig. 2). It shows a comparison of the polymorphisms of primer sites in all insects used in the search for primer sites in this study (Table S2-13, Fig. 1). The concordance rate of each locus (the graph above) and rate of each nucleic acid sequence (include indel) in the full length mtDNA 16S rRNA, 12S rRNA regions, and the COI region, which is the standard DNA barcoding region for insects, are shown in Fig S3. These results show that the primer sets for DNA metabarcoding developed in this study have been optimized to select the best sites because the primer regions have a higher nucleic sequence concordance rate than other loci, except for the regions with a lot of indels. Also, we suggested that the mtDNA COI region, which is the standard DNA barcoding region for insects, is an unsuitable DNA barcoding region to use for all insects because there was no region with a continuous high concordance rate. For the mtDNA COI region, it was not possible to find regions with high concordance rates, even at the mayfly order level.

**Figure 2.**
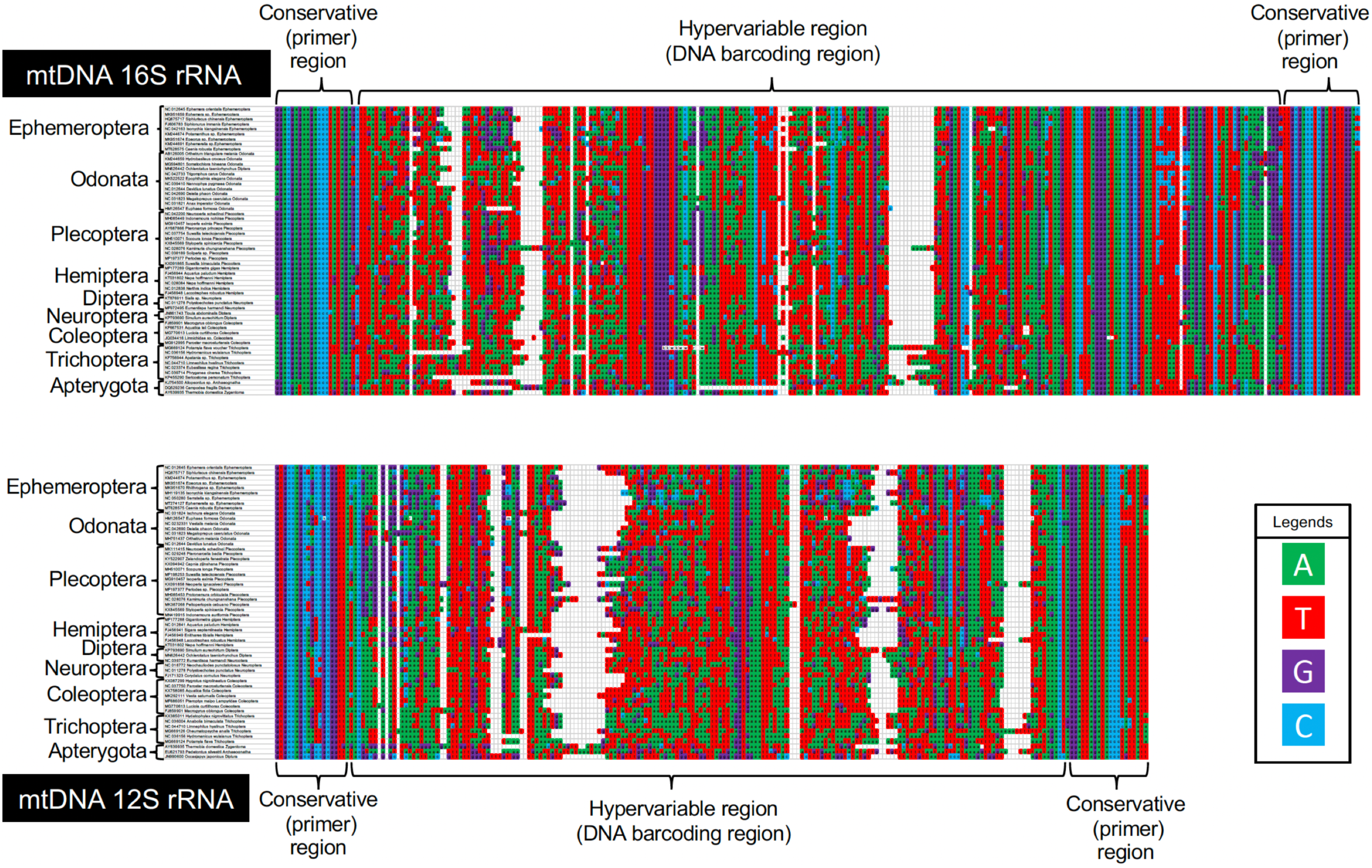
The barcoding region and the primer region were arranged using some of the sequences used in the development of each primer. The DNA barcoding regions suggested in this study in both the mtDNA 16S rRNA and 12S rRNA regions have high polymorphism and versatility by placing a hypervariable region between regions with highly conserved regions. Legends, green: “A” of nucleic acid sequence, red: “T” of nucleic acid sequence, “G” of nucleic acid sequence, “C” of nucleic acid sequence.

### Versatility and nucleotide polymorphisms of the DNA metabarcoding regions

In this study, using each of the applicable primer sets designed for PCR and Sanger sequencing, we succeeded in the PCR amplification and sequencing of DNA barcoding regions and longer fragments, including the respective DNA barcoding regions, for all species used. We confirmed successful amplification for 14 orders, 43 families, and 68 species using DNA barcoding in the mtDNA 16S rRNA region, and for 13 orders, 42 families, and 66 species for DNA barcoding in the mtDNA 12S rRNA (Table 1). It is necessary to confirm the methodology developed in this study in practice in future DNA barcoding and/or DNA metabarcoding research (e.g., eDNA), but there is no doubt that all the primers designed in this study have high versatility. Although PCR amplification was successful in all examined species of the various insect groups tested, care should be taken when using primer sets for the mtDNA 12S rRNA region in which Tm values differ by about 10°C between the forward and reverse primers. In the mtDNA 12S rRNA region, the GC nucleotides contained at the sites where the primers were designed was biased, so the Tm values of these primers could not be made uniform (Fig. S3).

A phylogenetic cladogram was constructed to investigate whether species could be identified using the DNA barcoding regions amplified using each primer set designed in this study (Fig. 3). As a result, the species analyzed could be identified and did not in any case share the same genotype among different species. In particular, as the DNA barcoding regions selected using our newly designed primer sets retain sufficient polymorphisms to identify each species, they can even differentiate between species of the same genus (Fig. 3).

**Figure 3.**
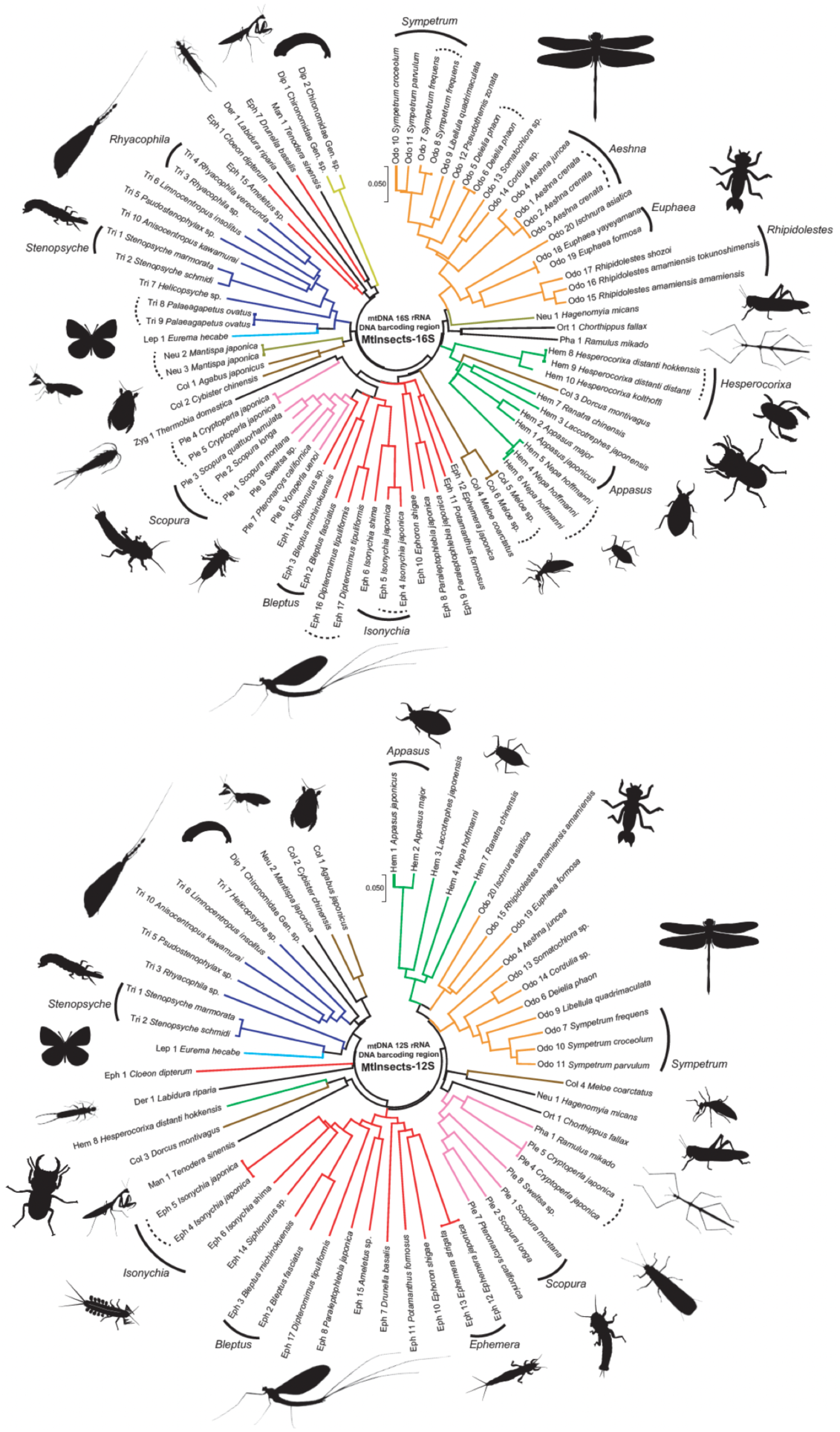
A phylogenetic cladogram based on concatenated data of the mtDNA 16S rRNA and the 12S rRNA regions to examine species identification sensitivity. The same species are shown by solid lines, and the same genus is shown by dashed lines.

Our study detected only two exceptional cases of inadequate DNA barcoding, both of which were due to species characteristics or for taxonomically problematic species. The marker (primer set) of the mtDNA 16S rRNA region we designed could not distinguish between two sub-species, *Hesperocorixa distanti hokkensis* and *Hesperocorixa distanti distanti*. However, these subspecies cannot be differentiated even using the sequence data of the mtDNA COI region (Yano et al. 2020). Therefore, these species cannot be differentiated by genetic markers or are possibly not in fact differentiated at the subspecies level. Regarding the second case, in the mtDNA 12S rRNA region, no genetic differentiation was observable between *Ephemera japonica* and *Ephemera strigata*. These species are sister species to each other, and are widely distributed sympatrically in the Japanese Islands (Okamoto and Tojo 2021). We confirmed interspecies introgression in areas where these two species had re-contacted after speciation (Takenaka et al. unpublished data). Therefore, we do not think such cases nullify the versatility of the primers we have developed.

### DNA barcoding method sufficiently sensitive to detect cryptic species

It is known that there are cryptic species within the mayfly species, *Epeorus aesculus* (Ogitani and Nakamura 2008; Tojo 2010). Therefore, this is a suitable species or species group to assess the potential of the genetic region we proposed for phylogenetic analysis. *Epeorus aesculus* were collected from multiple streams in the Japan Alps, including the type locality, and genetic analyses were conducted using the primer sets developed in this study to amplify longer fragments of their mtDNA 16S rRNA and 12S rRNA regions. As a result, a clade (i.e., Clade B) was detected that was largely genetically differentiated from another clade (i.e., Clade A: including the topotype specimens) collected at Kurobe-goro-zawa, which is the type locality of *E. aesculus* (Fig. 4). This means that *E. aesculus*, which has been treated as one species, has intraspecial large scale genetic differentiation, i.e., a cryptic species. Although it is necessary to investigate more samples and their morphology in detail, the results of this study adequately identified the existence of a known cryptic species or an undescribed species.

**Figure 4.**
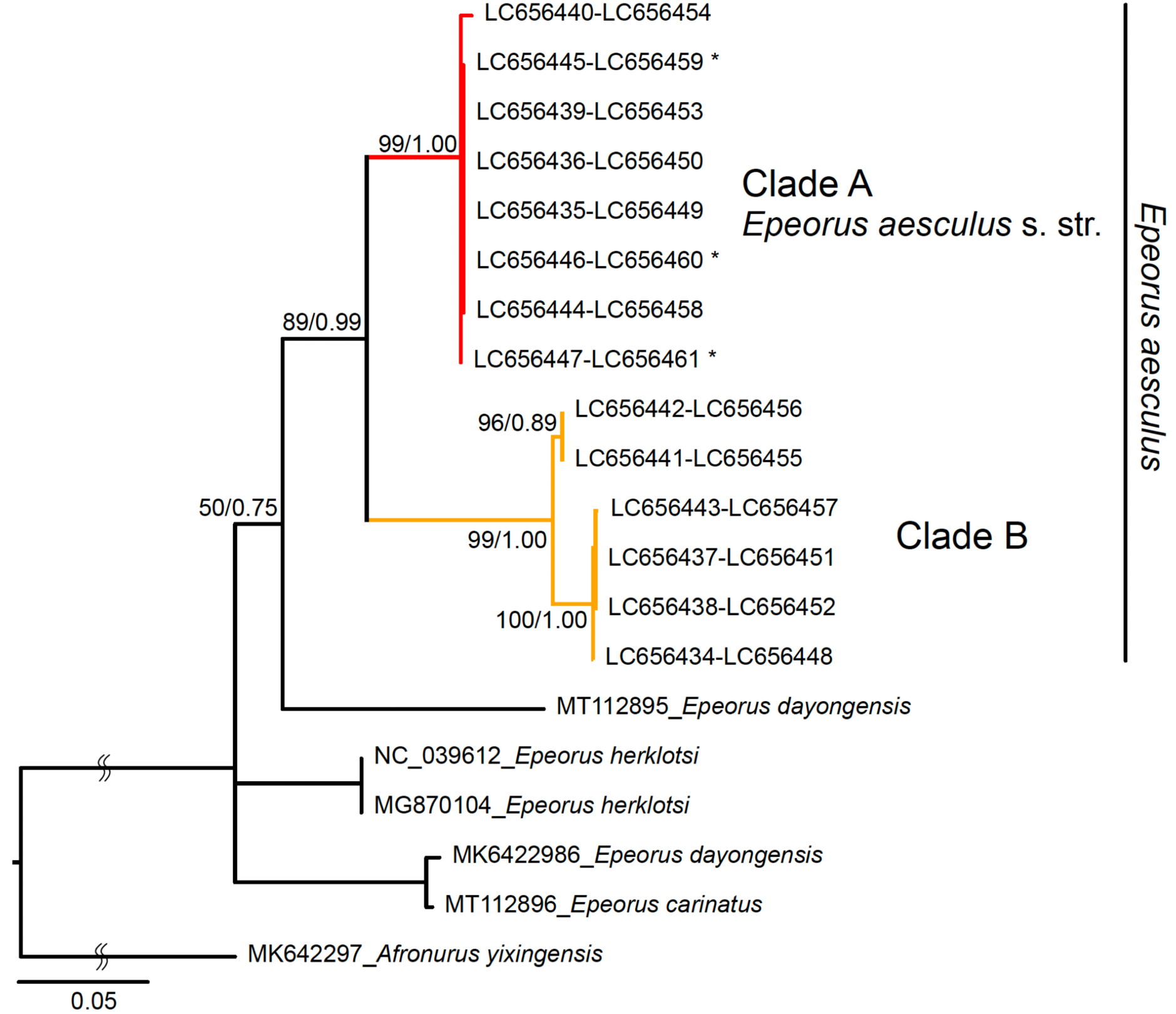
The estimated phylogenetic relationships (ML methods) based on the mtDNA 16S rRNA and 12S rRNA regions.

Today, DNA barcoding is a powerful tool for assessing biodiversity (Struck et al. 2018; Mosa et al. 2019). Most importantly, the genetic region of longer fragments amplified by the primer sets newly designed in this study contain a high number of polymorphisms by which detection of even cryptic species is possible. In addition, our development of the DNA barcoding methodology for short DNA fragments of about 200-bp will undoubtedly bring about further significant innovation in DNA metabarcoding methodology. Especially for aquatic insects, further development in the utilization of eDNA is anticipated. Under such circumstances, we believe that this study constitutes a methodological breakthrough that will underpin significant research advances.

## Conclusion

In this study, we succeeded in developing universal primer sets and newly selected optimal DNA barcoding regions. A key feature is that the DNA fragments amplified by these primer sets are both short at about 200-bp. By developing these primer sets for DNA barcoding or DNA metabarcoding that can be applied to almost all insect groups, it will enable relatively easy long-term monitoring of insect species composition and species diversity. In addition, it is also expected to facilitate the discovery of many undescribed species and/or cryptic species that have been overlooked or are difficult to identify morphologically.

We have established a barcoding region in the mtDNA ribosomal region; however, it is possible that closely related species that could not be identified using ribosomal regions will become identifiable in the future. For fish, it has been reported that the universal primer “MiFish” for the metabarcoding primers cannot differentiate species or amplify the barcoding region of some groups, so new primer sets focusing on specific groups have been designed (genus Anguilla: Takeuchi et al. 2019; Salmonidae: Morita et al. 2019; Cichlidae: Doble et al. 2019).

In the case of insects, if it is not possible to identify species using the ribosome DNA regions developed in this study, it may be able to identify them by using the mtDNA COI region, which is the traditional standard DNA barcoding region. In such cases, it would be best to make effective use of the existing database. Also, we recommend using multiple genetic regions to improve the accuracy and reliability of species identification. Finally, we were able to design not only DNA barcoding region markers, but also primer sets for longer fragment sequences for registration in the database. These primer sets that can amplify longer fragments can also be used for phylogenetic analyses, which will increase the level of data registration in the DNA database. In addition, such resulting database enhancements will serve as a powerful tool for increasingly accurate assessment of biodiversity and genetic diversity.

## Supplementary Information

**Table S1**. List of specimens of *Epeorus* mayflies examined in this study, sequence types, and the GenBank accession numbers

**Table S2**. Nucleotide sequences of universal primer (MtInsects-16S_F) and homologous region of 557 sequences of insects used for development in this study.

**Table S3**. Nucleotide sequences of universal primer (MtInsects-16S_R) and homologous region of 557 sequences of insects used for development in this study.

**Table S4**. Nucleotide sequences of universal primer (AQdb-16S_F) and homologous region of 557 sequences of insects used for development in this study.

**Table S5**. Nucleotide sequences of universal primer (AQdb-16S_R) and homologous region of 558 sequences of insects used for development in this study.

**Table S6**. Nucleotide sequences of universal primer (MtInsects-12S_HemHol_F) and homologous region of 491 sequences of insects used for development in this study.

**Table S7**. Nucleotide sequences of universal primer (MtInsects-12S-Hem_R) and homologous region of 265 sequences of insects used for development in this study.

**Table S8**. Nucleotide sequences of universal primer (MtInsects-12S_Hol_R) and homologous region of 209 sequences of insects used for development in this study.

**Table S9**. Nucleotide sequences of universal primer (MtInsects-12S_cad_F) and homologous region of 31 sequences of trichopteran insects used for development in this study.

**Table S10**. Nucleotide sequences of universal primer (MtInsects-12S_cad_R) and homologous region of 31 sequences of trichopteran insects used for development in this study.

**Table S11**. Nucleotide sequences of universal primer (AQdb-12S_Hem_R) and homologous region of 265 sequences of insects used for development in this study.

**Table S12**. Nucleotide sequences of universal primer (AQdb-12S_Hol_R) and homologous region of 214 sequences of insects used for development in this study.

**Table S13**. Nucleotide sequences of universal primer (AQdb-12S_cad_R) and homologous region of 31 sequences of trichopteran insects used for development in this study.

**Figure S1**. Position of primers sets in the mtDNA 16S rRNA region. Reference sequence data used the mtDNA 16S rRNA region in the complete mitochondrion genome of *Ephemera orientalis* (NC_012645).

**Figure S2**. Position of primers sets in the mtDNA 12S rRNA region. Reference sequence data used the mtDNA 12S rRNA region in the complete mitochondrion genome of *Ephemera orientalis* (NC_012645).

**Figure S3**. Graph of the nucleotide concordance rate (0.0-1.0; above) and the proportion of adenine (A: green), thymine (T: red), guanine (G: purple), cytosine (C: blue) and indel (white) (0.0-1.0; below) in each nucleotide position in the full length of each of the mtDNA 16S rRNA, 12S rRNA, and the COI regions. The nucleotide concordance rate of each locus is shown by gray bars, and the region of each newly designed primer is shown black bars.

## Acknowledgements

We express our thanks to Dr. Y. Iwasaki (National Institute of Advanced Industrial Science and Technology), Dr. N. Uchida (Tohoku University), Dr. N. Kondo (National Institute of Environmental studies), Dr. Y. Hasebe (Kanagawa Prefectural Government) for valuable advice and encouragement. We are indebted to Tojo Lab. members (Shinshu University) and Niimi Lab. members (NIBB) for their cooperation with the field research and collection of specimens. This study was supported by grants from the River Environment Fund (22-1215-021 to KT, 2017-5211-025 to KT, 2021-5311-005 to MT) of River and Watershed Environment Management.

## Contributions

M.T. designed, managed the study, and performed sample collection; M.T., K.Y. mainly performed laboratory work and phylogenetic analyses; M.T., K.Y., T.S. designed Figures and Tables; M.T., K.T. gathered funds; M.T., K.Y., and M.T., K.Y., T.S., K.T. wrote and reviewed the manuscript.

## Ethics declarations

The authors declare that they have no competing interests. The experiments comply with the current laws of the country in which they were performed.

## References

Ahn H, Kume M, Terashima Y, Ye F, Kameyama S, Miya M, Yamashita Y, Kasai A (2020) Evaluation of fish biodiversity in estuaries using environmental DNA metabarcoding. PloS ONE 15: e0231127. https://doi.org/10.1371/journal.pone.0231127

Bohmann K, Evans A, Gilbert MTP, Carvalho GR, Creer S, Knapp M, Douglas WY, De Bruyn M (2014) Environmental DNA for wildlife biology and biodiversity monitoring. Trends Ecol. Evol. 29: 358–367. https://doi.org/10.1016/j.tree.2014.04.003

Ceballos G, Ehrlich PR, Barnosky AD, García A, Pringle RM, Palmer TM (2015) Accelerated modern human–induced species losses: Entering the sixth mass extinction. Sci. Adv. 1: e1400253. https://doi.org/10.1126/sciadv.1400253

Chucholl F, Fiolka F, Segelbacher G, Epp LS (2021) eDNA detection of native and invasive crayfish species allows for year-round monitoring and large-scale screening of lotic systems. Front. Environ. Sci. 9: 23. https://doi.org/10.3389/fenvs.2021.639380

Collins RA, Bakker J, Wangensteen OS, Soto AZ, Corrigan L, Sims DW, Genner MJ, Mariani S (2019) Non-specific amplification compromises environmental DNA metabarcoding with COI. Methods Ecol. Evol. 10: 1985–2001. https://doi.org/10.1111/2041-210X.13276

Deagle BE, Jarman SN, Coissac E, Pompanon F, Taberlet P (2014) DNA metabarcoding and the cytochrome c oxidase subunit I marker: not a perfect match. Biol. Lett. 10: 20140562. https://doi.org/10.1098/rsbl.2014.0562

Deiner K, Bik HM, Mächler E, Seymour M, Lacoursière-Roussel A, Altermatt F, Creer S, Bista I, Lodge DM, de Vere N, Pfrender ME, Bernatchez L (2017) Environmental DNA metabarcoding: Transforming how we survey animal and plant communities. Mol. Ecol. 26: 5872–5895. https://doi.org/10.1111/mec.14350

Dirzo R, Young HS, Galetti M, Ceballos G, Isaac NJ, Collen B (2014) Defaunation in the Anthropocene. Science 345: 401–406. https://doi.org/10.1126/science.1251817

Doble CJ, Hipperson H, Salzburger W, Horsburgh GJ, Mwita C, Murrell DJ, Day JJ (2020) Testing the performance of environmental DNA metabarcoding for surveying highly diverse tropical fish communities: A case study from Lake Tanganyika. Environmental DNA 2: 24–41. https://doi.org/10.1002/edn3.43

Folmer O, Black M, Hoeh W, Lutz R, Vrijenhoek R (1994) DNA primers for amplification of mitochondrial cytochrome c oxidase subunit I from diverse metazoan invertebrates. Mol. Mar. Biol. Biotechnol. 3: 294–9.

Grimaldi D, Engel MS (2005) Evolution of the Insects. Cambridge University Press, Cambridge.

Hänfling B, Lawson Handley L, Read DS, Hahn C, Li J, Nichols P, Blackman RC, Oliver A, Winfield IJ (2016) Environmental DNA metabarcoding of lake fish communities reflects long-term data from established survey methods. Mol. Ecol. 25: 3101–3119. https://doi.org/10.1111/mec.13660

Hebert PD, Penton EH, Burns JM, Janzen DH, Hallwachs W (2004) Ten species in one: DNA barcoding reveals cryptic species in the neotropical skipper butterfly Astraptes fulgerator. Proc. Natl. Acad. Sci. 101: 14812–14817. https://doi.org/10.1073/pnas.0406166101

Hebert PD, Gregory TR (2005) The promise of DNA barcoding for taxonomy. Syst. Biol. 54: 852–859. https://doi.org/10.1080/10635150500354886

Katoh K, Standley DM (2013) MAFFT multiple sequence alignment software version 7: improvements in performance and usability. Mol. Biol. Evol. 30: 772–780. https://doi.org/10.1093/molbev/mst010

Komai T, Gotoh RO, Sado T, Miya M (2019) Development of a new set of PCR primers for eDNA metabarcoding decapod crustaceans. Metabarcoding Metagenom. 3: e33835. https://doi.org/10.3897/mbmg.3.33835

Kudoh A, Minamoto T, Yamamoto S (2020) Detection of herbivory: eDNA detection from feeding marks on leaves. Environmental DNA 2: 627–634.

Kumar S, Stecher G, Tamura K (2016) MEGA7: molecular evolutionary genetics analysis version 7.0 for bigger datasets. Mol. Biol. Evol. 33: 1870–1874. https://doi.org/10.1093/molbev/msw054

Kuwae M, Tamai H, Doi H, Sakata MK, Minamoto T, Suzuki Y (2020) Sedimentary DNA tracks decadal-centennial changes in fish abundance. Commun. Biol. 3: 1–12. https://doi.org/10.1038/s42003-020-01282-9

Miya M, Sato Y, Fukunaga T, Sado T, Poulsen JY, Sato K, Minamoto T, Yamamoto S, Yamanaka H, Araki H, Iwasaki W (2015) MiFish, a set of universal PCR primers for metabarcoding environmental DNA from fishes: detection of more than 230 subtropical marine species. Royal Soc. Open Sci. 2: 150088. https://doi.org/10.1098/rsos.150088

Mora C, Tittensor DP, Adl S, Simpson AG, Worm B (2011) How many species are there on Earth and in the ocean?. PLoS Biol. 9: e1001127. https://doi.org/10.1371/journal.pbio.1001127

Morita K, Sahashi G, Miya M, Kamada S, Kanbe T, Araki H (2019) Ongoing localized extinctions of stream-dwelling white-spotted charr populations in small dammed-off habitats of Hokkaido Island, Japan. Hydrobiologia 840: 207–213. https://doi.org/10.1007/s10750-019-3891-1

Mosa KA, Gairola S, Jamdade R, El-Keblawy A, Al Shaer KI, Al Harthi EK, Shabana HA, Mahmoud T (2019) The promise of molecular and genomic techniques for biodiversity research and DNA barcoding of the Arabian Peninsula flora. Front. Plant Sci. 9: 1929. https://doi.org/10.3389/fpls.2018.01929

Okamoto S, Saito T, Tojo K (2021) Geographical fine-scaled distributional differentiation caused by niche differentiation in three closely related mayflies. Limnol. https://doi.org/10.1007/s10201-021-00673-z.

Okamoto, S., & Tojo, K. (2021). Distribution patterns and niche segregation of three closely related Japanese ephemerid mayflies: a re-examination of each species’ habitat from “megadata” held in the “National Census on River Environments”. Limnol. 22: 227–287. https://doi.org/10.1007/s10201-021-00654-2

Ohgitani M, Nakamura H (2008) Distribution and Seasonal Population Change of Heptageniidae Nymph in the Oguro River (The Branch of Tenryu River). The Annals of environmental science, Shinshu University 30: 57-66. [Japanese]

Ohnishi O, Takenaka M, Okano R, Yoshitomi H, Tojo K (2021) Wide-scale gene flow, even in Insects that have lost their flight ability: presence of dispersion due to a unique parasitic ecological strategy of piggybacking hosts. Zool. Sci. 38: 122–139. https://doi.org/10.2108/zs200088

Rambaut A (2009) FigTree, Version 1.3.1. Available at: http://tree.bio.ed.ac.uk/software/figtree/ (cited 25 March 2019).

Rambaut A, Suchard MA, Xie D, Drummond AJ (2014) Tracer, version 1.6, MCMC trace analysis package. Available at: http://tree.bio.ed.ac.uk/software/tracer/ (xcited 25 March 2019).

Ronquist F, Teslenko M, van der Mark P, Ayres DL, Darling A, Höhna S, Larget B, Liu L, Suchard MA, Huelsenbeck JP (2012) MrBayes 3.2: efficient Bayesian phylogenetic inference and model choice across a large model space. Syst. Biol. 61: 539–542. https://doi.org/10.1093/sysbio/sys029

Saitoh T, Sugita N, Someya S, Iwami Y, Kobayashi S, Kamigaichi H, Higuchi A, Asai S, Yamamoto Y, Nishiumi I (2015) DNA barcoding reveals 24 distinct lineages as cryptic bird species candidates in and around the Japanese Archipelago. Mol. Ecol. Resour. 15: 177–186. https://doi.org/10.1111/1755-0998.12282

Schindel DE, Miller SE (2005) DNA barcoding a useful tool for taxonomists. Nature 435: 17.

Schwarz G (1978) Estimating the dimension of a model. Ann. Stat. 6: 461–464.

Sekiya T, Ichiyanagi H, Tojo K (2017) Establishing of genetic analyses methods of feces from the water shrew, Chimarrogale platycephalus (Erinaceidae, Eulipotyphla). JSM Biol. 2: 1010.

Stork NE (2018) How many species of insects and other terrestrial arthropods are there on Earth?. Annu. Rev. Entomol. 63: 31–45. https://doi.org/10.1146/annurev-ento-020117-043348

Struck TH, Feder JL, Bendiksby M, Birkeland S, Cerca J, Gusarov VI, Kistenich S, Larsson KH, Liow LH, Nowak MD, Stedje B, Bachmann L, Dimitrov D (2018) Finding evolutionary processes hidden in cryptic species. Trends Ecol. Evol. 33: 153–163. https://doi.org/10.1016/j.tree.2017.11.007

Takenaka M, Tojo K (2019) Ancient origin of a dipteromimid mayfly family endemic to the Japanese Islands and its genetic differentiation across tectonic faults. Biol. J. Linn. Soc. 126: 555–573. https://doi.org/10.1093/biolinnean/bly192

Takeuchi A, Iijima T, Kakuzen W, Watanabe S, Yamada Y, Okamura A, Horie N, Mikawa N, Miller MJ, Kojima T, Tsukamoto K (2019) Release of eDNA by different life history stages and during spawning activities of laboratory-reared Japanese eels for interpretation of oceanic survey data. Sci. Rep. 9: 1–9. https://doi.org/10.1038/s41598-019-42641-9

Tanabe AS (2007) Kakusan: a computer program to automate the selection of a nucleotide substitution model and the configuration of a mixed model on multilocus data. Mol. Ecol. Notes 7: 962–964. https://doi.org/10.1111/j.1471-8286.2007.01807.x

Tojo K (2010) The current distribution of aquatic insects in habiting river systems, with respect to their population and genetic structure. In Harris EL, Davies NE (eds). Insect habitats: characteristics, diversity and management. Nova Science, New York: 157–161.

Tojo K, Sekiné K, Takenaka M, Isaka Y, Komaki S, Suzuki T, Schoville SD (2017) Species diversity of insects in Japan: their origins and diversification processes. Entomol. Sci. 20: 357–381. https://doi.org/10.1111/ens.12261

Ueda S, Nozawa T, Matsuzuki T, Seki RI, Shimamoto S, Itino T (2012) Phylogeny and phylogeography of Myrmica rubra complex (Myrmicinae) in the Japanese Alps. Psyche 2012: 1–7. https://doi.org/10.1155/2012/319097

Uchida N, Kubota K, Aita S, Kazama S (2020) Aquatic insect community structure revealed by eDNA metabarcoding derives indices for environmental assessment. PeerJ 8: e9176. https://doi.org/10.7717/peerj.9176

Ushio M, Fukuda H, Inoue T, Makoto K, Kishida O, Sato K, Murata K, Nikaido M, Sado Y, Takeshita M, Iwasaki W, Yamanaka H, Kondoh M, Miya M. (2017) Environmental DNA enables detection of terrestrial mammals from forest pond water. Mol. Ecol. Resour. 17: e63–e75. https://doi.org/10.1111/1755-0998.12690

Ushio M, Murata K, Sado T, Nishiumi I, Takeshita M, Iwasaki W, Miya M (2018) Demonstration of the potential of environmental DNA as a tool for the detection of avian species. Sci. Rep. 8: 1–10. https://doi.org/10.1038/s41598-018-22817-5

Yamazaki H, Sekiya T, Nagayama S, Hirasawa K, Tokura K, Sasaki A, Ichiyanagi H, Tojo K (2020) Development of microsatellite markers for a soricid water shrew, Chimarrogale platycephalus, and their successful use for individual identification. Genes Genet. Syst. 95: 201–210. https://doi.org/10.1266/ggs.20-00017

Yano K, Takenaka M, Tojo K (2019) Genealogical position of Japanese populations of the globally distributed mayfly Cloeon dipterum and related species (Ephemeroptera, Baetidae): A molecular phylogeographic analysis. Zool. Sci. 36: 479–489. https://doi.org/10.2108/zs190049

Yano K, Takenaka M, Mitamura T, Tojo K (2020) Identifying a “pseudogene” for the mitochondrial DNA COI region of the corixid aquatic insect, Hesperocorixa distanti (Heteroptera, Corixidae). Limnology 21: 319–325. https://doi.org/10.1007/s10201-020-00618-y

Valentini A, Pompanon F, Taberlet P (2009. DNA barcoding for ecologists. Trends Ecol. Evol. 24: 110–117. https://doi.org/10.1016/j.tree.2008.09.011

Vuataz L, Sartori M, Gattolliat JL, Monaghan MT (2013) Endemism and diversification in freshwater insects of Madagascar revealed by coalescent and phylogenetic analysis of museum and field collections. Mol. Phylogenet. Evol. 66: 979–991. https://doi.org/10.1016/j.ympev.2012.12.003

