## Supplementary figures and images for "Development of novel PCR primer sets for DNA metabarcoding of aquatic insects, and the discovery of some cryptic species"

### sap_Figures.pdf

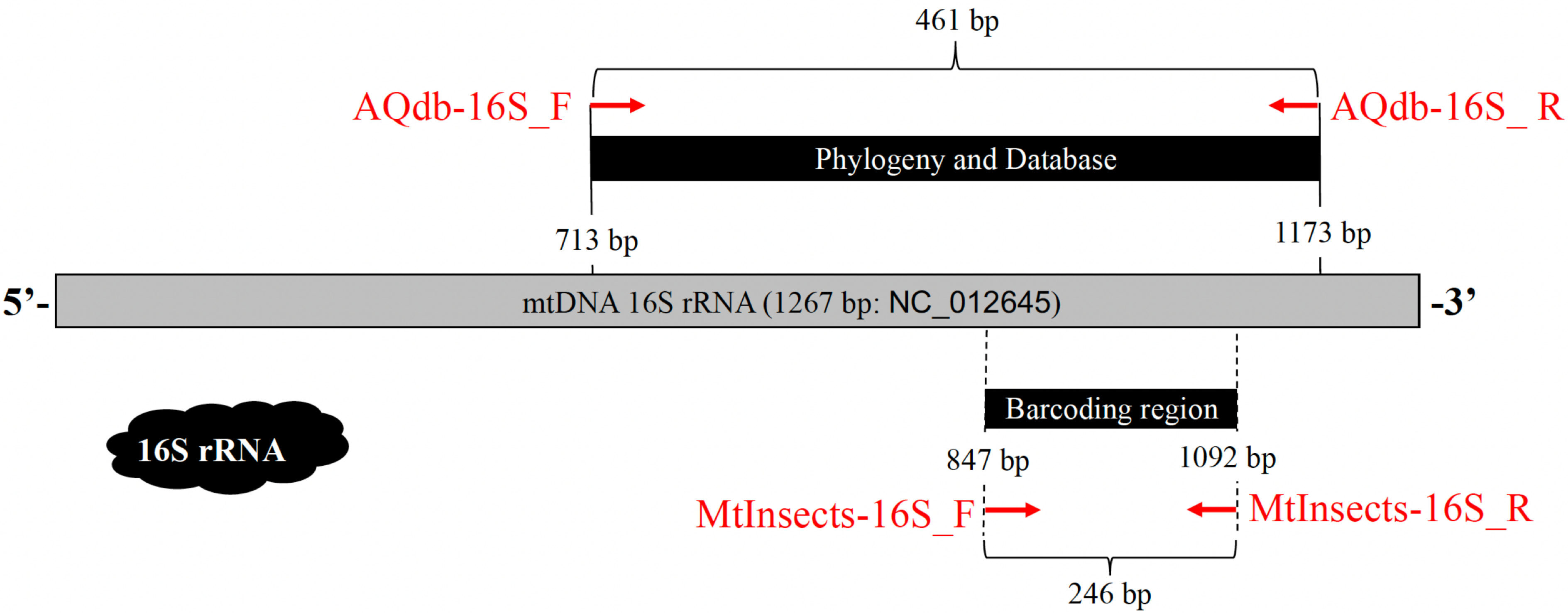

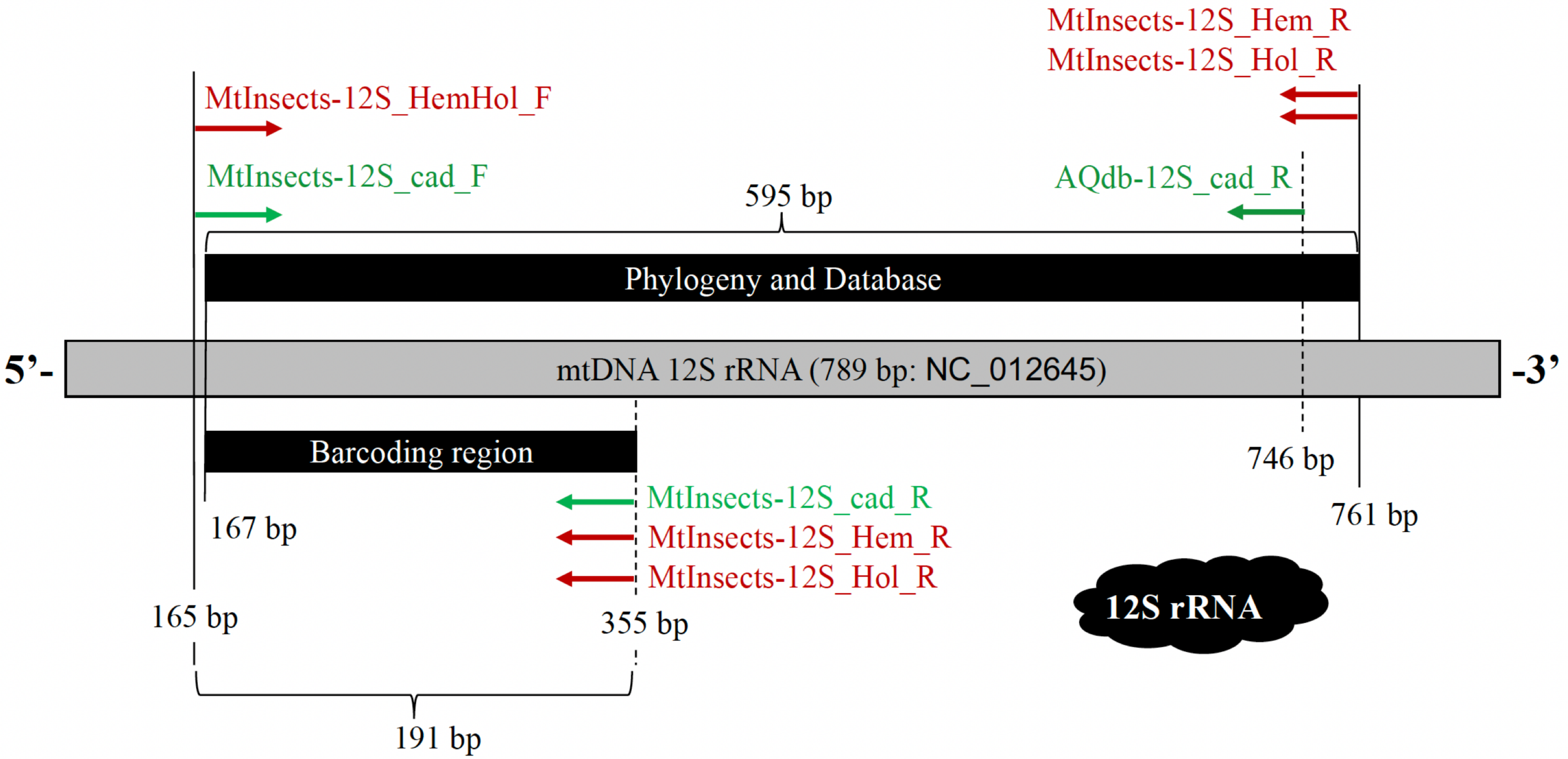

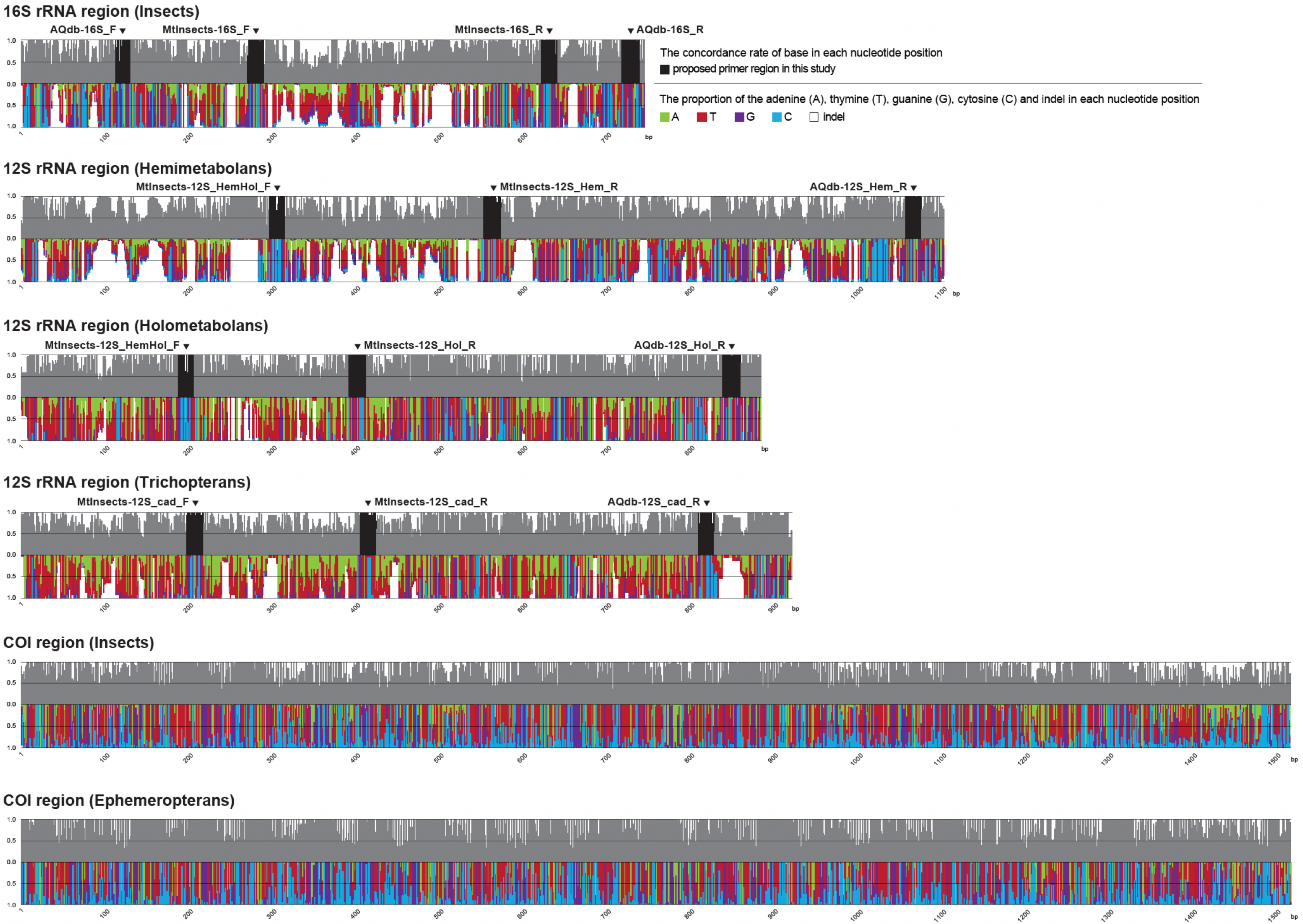
