## Supplemental Tables for "Development of novel PCR primer sets for DNA metabarcoding of aquatic insects, and the discovery of some cryptic species": supplement Table.pdf

Table S1. List of specimens of *Epeorus* mayflies examined in this study, sequence types, and the GenBank accession numbers

| | locality name | Latitude (N) | Longitude (E) | mtDNA 16S rRNA | mtDNA 12S rRNA | clade |
|---|---|---|---|---|---|---|
| 1 | Yokokawa, Tatsuno, Nagano | 36.01359 | 137.93795 | LC656434 | LC656448 | clade B |
| 2 | Nagawa, Matsumoto, Nagano | 36.09151 | 137.68030 | LC656435 | LC656449 | clade A |
| | | | | LC656436 | LC656450 | clade A |
| | | | | LC656437 | LC656451 | clade B |
| | | | | LC656438 | LC656452 | clade B |
| 3 | Narai, Shiojiri, Nagano | 35.88703 | 137.82736 | LC656439 | LC656453 | clade A |
| | | | | LC656440 | LC656454 | clade A |
| | | | | LC656441 | LC656455 | clade B |
| | | | | LC656442 | LC656456 | clade B |
| 4 | Iriyamabe, Matsumoto, Nagano | 36.18853 | 138.08131 | LC656443 | LC656457 | clade B |
| | | | | LC656444 | LC656458 | clade A |
| 5 | Kurobegoro, Toyama, Toyama | - | - | LC656445 | LC656459 | clade A |
| | | | | LC656446 | LC656460 | clade A |
| | | | | LC656447 | LC656461 | clade A |

Table S2. Nucleotide sequences of universal primer (MtInsects-16S_F) and homologous region of 557 sequences of insects used for development in this study.

| | | | | | | | | | | | | | | | | | | | | | | | |
|---|---|---|---|---|---|---|---|---|---|---|---|---|---|---|---|---|---|---|---|---|---|---|---|
| MtInsects-16S_F | | | | G | G | A | C | G | A | G | A | A | G | A | C | C | C | T | W | T | A | G | A |
| | | | A | 200 | 0 | 557 | 0 | 11 | 557 | 3 | 557 | 557 | 0 | 557 | 0 | 0 | 0 | 0 | 520 | 0 | 557 | 18 | 557 |
| | | | T | 2 | 0 | 0 | 0 | 0 | 0 | 42 | 0 | 0 | 0 | 0 | 0 | 0 | 0 | 528 | 22 | 555 | 0 | 0 | 0 |
| | | | G | 355 | 557 | 0 | 0 | 546 | 0 | 512 | 0 | 0 | 557 | 0 | 0 | 0 | 0 | 0 | 15 | 0 | 0 | 539 | 0 |
| | | | C | 0 | 0 | 0 | 557 | 0 | 0 | 0 | 0 | 0 | 0 | 0 | 557 | 557 | 557 | 29 | 0 | 0 | 0 | 0 | 0 |
| GenBank Acc. No. | Order | Species | | | | | | | | | | | | | | | | | | | | | |
| MN481418.1 | Diplura | *Campodea augens* | | G | G | A | C | G | A | T | A | A | G | A | C | C | C | T | A | T | A | G | A |
| DQ529236.1 | Diplura | *Campodea fragilis* | | G | G | A | C | G | A | T | A | A | G | A | C | C | C | T | A | T | A | G | A |
| NC008233.1 | Diplura | *Campodea fragilis* | | G | G | A | C | G | A | T | A | A | G | A | C | C | C | T | A | T | A | G | A |
| DQ529237.1 | Diplura | *Campodea lubbocki* | | G | G | A | C | G | A | T | A | A | G | A | C | C | C | T | A | T | A | G | A |
| NC008234.1 | Diplura | *Campodea lubbocki* | | G | G | A | C | G | A | T | A | A | G | A | C | C | C | T | A | T | A | G | A |
| AY771989.1 | Diplura | *Japyx solifugus* | | G | G | A | C | G | A | T | A | A | G | A | C | C | C | T | A | T | A | G | A |
| NC007214.1 | Diplura | *Japyx solifugus* | | G | G | A | C | G | A | T | A | A | G | A | C | C | C | T | A | T | A | G | A |
| JN990601.1 | Diplura | *Lepidocampa weberi* | | G | G | A | C | G | A | T | A | A | G | A | C | C | C | T | A | ? | A | A | A |
| NC022675.1 | Diplura | *Lepidocampa weberi* | | G | G | A | C | G | A | T | A | A | G | A | C | C | C | T | A | ? | A | A | A |

| | | | | | | | | | | | | | | | | | | | | | | |
|---|---|---|---|---|---|---|---|---|---|---|---|---|---|---|---|---|---|---|---|---|---|---|
| JN990600.1 | Diplura | *Occasjapyx japonicus* | G | G | A | C | G | A | T | A | A | G | A | C | C | C | T | A | T | A | G | A |
| NC022674.1 | Diplura | *Occasjapyx japonicus* | G | G | A | C | G | A | T | A | A | G | A | C | C | C | T | A | T | A | G | A |
| JN990598.1 | Diplura | *Octostigma sinensis* | G | G | A | C | G | A | T | A | A | G | A | C | C | C | T | T | T | A | G | A |
| NC022672.1 | Diplura | *Octostigma sinensis* | G | G | A | C | G | A | T | A | A | G | A | C | C | C | T | T | T | A | G | A |
| JN990599.1 | Diplura | *Parajapyx emeryanus* | G | G | A | C | G | A | T | A | A | G | A | C | C | C | T | A | T | A | G | A |
| NC022673.1 | Diplura | *Parajapyx emeryanus* | G | G | A | C | G | A | T | A | A | G | A | C | C | C | T | A | T | A | G | A |
| KJ754500.1 | Archaeognatha | *Allopsontus* sp. | G | G | A | C | A | A | A | A | A | G | A | C | C | C | T | A | T | A | G | A |
| KJ754501.1 | Archaeognatha | *Allopsontus* sp. | G | G | A | C | A | A | G | A | A | G | A | C | C | C | T | A | T | A | G | A |
| AY793551.1 | Archaeognatha | *Nesomachilis australica* | G | G | A | C | G | A | T | A | A | G | A | C | C | C | T | A | T | A | G | A |
| NC006895.1 | Archaeognatha | *Nesomachilis australica* | G | G | A | C | G | A | T | A | A | G | A | C | C | C | T | A | T | A | G | A |
| KJ754502.1 | Archaeognatha | *Pedetontinus luanchuanensis* | G | G | A | C | A | A | T | A | A | G | A | C | C | C | T | A | T | A | G | A |
| EU621793.1 | Archaeognatha | *Pedetontus silvestrii* | G | G | A | C | A | A | T | A | A | G | A | C | C | C | T | A | T | A | G | A |

| | | | | | | | | | | | | | | | | | | | | | | |
|---|---|---|---|---|---|---|---|---|---|---|---|---|---|---|---|---|---|---|---|---|---|---|
| NC011717.1 | Archaeognatha | *Pedetontus silvestrii* | G | G | A | C | A | A | T | A | A | G | A | C | C | C | T | A | T | A | G | A |
| KJ754503.1 | Archaeognatha | *Petrobiellus* sp. | G | G | A | C | G | A | T | A | A | G | A | C | C | C | T | A | T | A | G | A |
| KJ754504.1 | Archaeognatha | *Petrobiellus* sp. | G | G | A | C | G | A | T | A | A | G | A | C | C | C | T | A | T | A | G | A |
| AY956355.1 | Archaeognatha | *Petrobius brevistylis* | T | G | A | C | A | A | T | A | A | G | A | C | C | C | T | A | T | A | G | A |
| NC007688.1 | Archaeognatha | *Petrobius brevistylis* | T | G | A | C | A | A | T | A | A | G | A | C | C | C | T | A | T | A | G | A |
| JX308221.1 | Archaeognatha | *Songmachilis xinxiangensis* | G | G | A | C | A | A | A | A | A | G | A | C | C | C | T | A | T | A | G | A |
| NC021384.1 | Archaeognatha | *Songmachilis xinxiangensis* | G | G | A | C | A | A | A | A | A | G | A | C | C | C | T | A | T | A | G | A |
| EU016193.1 | Archaeognatha | *Trigoniophthalmus alternatus* | A | G | A | C | A | A | T | A | A | G | A | C | C | C | T | A | T | A | G | A |
| NC010532.1 | Archaeognatha | *Trigoniophthalmus alternatus* | A | G | A | C | A | A | T | A | A | G | A | C | C | C | T | A | T | A | G | A |
| EU084035.1 | Zygentoma | *Atelura formicaria* | G | G | A | C | G | A | T | A | A | G | A | C | C | C | T | A | T | A | G | A |
| NC011197.1 | Zygentoma | *Atelura formicaria* | G | G | A | C | G | A | T | A | A | G | A | C | C | C | T | A | T | A | G | A |
| MK301436.1 | Zygentoma | *Ctenolepisma villosa* | G | G | A | C | G | A | T | A | A | G | A | C | C | C | T | G | T | A | G | A |
| NC046478.1 | Zygentoma | *Ctenolepisma villosa* | G | G | A | C | G | A | T | A | A | G | A | C | C | C | T | G | T | A | G | A |

| | | | | | | | | | | | | | | | | | | | | | | |
|---|---|---|---|---|---|---|---|---|---|---|---|---|---|---|---|---|---|---|---|---|---|---|
| MT108230.1 | Zygentoma | *Lepisma saccharina* | G | G | A | C | G | A | T | A | A | G | A | C | C | C | T | A | T | A | G | A |
| NC047445.1 | Zygentoma | *Lepisma saccharina* | G | G | A | C | G | A | T | A | A | G | A | C | C | C | T | A | T | A | G | A |
| AY639935.1 | Zygentoma | *Thermobia domestica* | G | G | A | C | G | A | T | A | A | G | A | C | C | C | T | G | T | A | G | A |
| NC006080.1 | Zygentoma | *Thermobia domestica* | G | G | A | C | G | A | T | A | A | G | A | C | C | C | T | G | T | A | G | A |
| MK951657.1 | Zygentoma | *Thermobia* sp. | G | G | A | C | G | A | T | A | A | G | A | C | C | C | T | A | T | A | G | A |
| AY191994.1 | Zygentoma | *Tricholepidion gertschi* | G | G | A | C | G | A | T | A | A | G | A | C | C | C | T | A | T | A | G | A |
| NC005437.1 | Zygentoma | *Tricholepidion gertschi* | G | G | A | C | G | A | T | A | A | G | A | C | C | C | T | A | T | A | G | A |
| HM067837.1 | Zygentoma | *Vietnamella dabieshanensis* | G | G | A | C | G | A | G | A | A | G | A | C | C | C | T | A | T | A | G | A |
| KM244655.1 | Zygentoma | *Vietnamella* sp. | G | G | A | C | G | A | G | A | A | G | A | C | C | C | T | A | T | A | G | A |
| GU479735.1 | Ephemeroptera | *Alainites yixiani* | G | G | A | C | G | A | G | A | A | G | A | C | C | C | C | A | T | A | G | A |
| GU936204.1 | Ephemeroptera | *Baetis* sp. | G | G | A | C | G | A | G | A | A | G | A | C | C | C | C | A | T | A | G | A |
| MT628575.1 | Ephemeroptera | *Caenis robusta* | G | G | A | C | G | A | G | A | A | G | A | C | C | C | T | A | T | A | G | A |
| GQ502451.1 | Ephemeroptera | *Caenis* sp. | G | G | A | C | G | A | G | A | A | G | A | C | C | C | T | A | T | A | G | A |
| MG910499.1 | Ephemeroptera | *Caenis* sp. | G | G | A | C | G | A | G | A | A | G | A | C | C | C | T | A | T | A | G | A |
| MN356096.1 | Ephemeroptera | *Caenis* sp. | G | G | A | C | G | A | G | A | A | G | A | C | C | C | T | A | T | A | G | A |

| | | | | | | | | | | | | | | | | | | | | | | |
|---|---|---|---|---|---|---|---|---|---|---|---|---|---|---|---|---|---|---|---|---|---|---|
| MN807287.1 | Ephemeroptera | *Choroterpides apiculata* | A | G | A | C | G | A | G | A | A | G | A | C | C | C | T | A | T | A | G | A |
| MG870104.1 | Ephemeroptera | *Epeorus herklotsi* | G | G | A | C | G | A | G | A | A | G | A | C | C | C | T | A | T | A | G | A |
| MH752075.1 | Ephemeroptera | *Epeorus herklotsi* | G | G | A | C | G | A | G | A | A | G | A | C | C | C | T | A | T | A | G | A |
| NC039612.1 | Ephemeroptera | *Epeorus herklotsi* | G | G | A | C | G | A | G | A | A | G | A | C | C | C | T | A | T | A | G | A |
| KJ493406.1 | Ephemeroptera | *Epeorus* sp. | G | G | A | C | G | A | G | A | A | G | A | C | C | C | T | A | T | A | G | A |
| KM244708.1 | Ephemeroptera | *Epeorus* sp. | G | G | A | C | G | A | G | A | A | G | A | C | C | C | T | A | T | A | G | A |
| MK951674.1 | Ephemeroptera | *Epeorus* sp. | G | G | A | C | G | A | G | A | A | G | A | C | C | C | T | A | T | A | G | A |
| EU591678.1 | Ephemeroptera | *Ephemera orientalis* | G | G | A | C | G | A | G | A | A | G | A | C | C | C | T | A | T | A | G | A |
| NC012645.1 | Ephemeroptera | *Ephemera orientalis* | G | G | A | C | G | A | G | A | A | G | A | C | C | C | T | A | T | A | G | A |
| MK951659.1 | Ephemeroptera | *Ephemera* sp. | G | G | A | C | G | A | G | A | A | G | A | C | C | C | T | A | T | A | G | A |
| KM244691.1 | Ephemeroptera | *Ephemerella* sp. | G | G | A | C | G | A | G | A | A | G | A | C | C | C | T | A | T | A | G | A |
| MT274127.1 | Ephemeroptera | *Ephemerella* sp. | G | G | A | C | G | A | G | A | A | G | A | C | C | C | T | A | T | A | G | A |
| NC050279.1 | Ephemeroptera | *Ephemerella* sp. | G | G | A | C | G | A | G | A | A | G | A | C | C | C | T | A | T | A | G | A |
| GU936203.1 | Ephemeroptera | *Habrophlebiodes zijinensis* | A | G | A | C | G | A | G | A | A | G | A | C | C | C | T | A | T | A | G | A |
| HM143892.1 | Ephemeroptera | *Isonychia ignota* | G | G | A | C | G | A | G | A | A | G | A | C | C | C | T | A | T | A | G | A |
| MH119135.1 | Ephemeroptera | *Isonychia kiangsinensis* | G | G | A | C | G | A | G | A | A | G | A | C | C | C | T | A | T | A | G | A |
| NC042163.1 | Ephemeroptera | *Isonychia* | G | G | A | C | G | A | G | A | A | G | A | C | C | C | T | A | T | A | G | A |

| | | | | | | | | | | | | | | | | | | | | | | |
|---|---|---|---|---|---|---|---|---|---|---|---|---|---|---|---|---|---|---|---|---|---|---|
| | | *kiangsinensis* | | | | | | | | | | | | | | | | | | | | |
| MK951658.1 | Ephemeroptera | *Isonychia* sp. | G | G | A | C | G | A | G | A | A | G | A | C | C | C | T | A | T | A | G | A |
| HM004123.1 | Ephemeroptera | *Paegniodes cupulatus* | G | G | A | C | G | A | G | A | A | G | A | C | C | C | T | A | T | A | G | A |
| MK951673.1 | Ephemeroptera | *Parafronurus sp.* | G | G | A | C | G | A | G | A | A | G | A | C | C | C | T | A | T | A | G | A |
| EU349015.1 | Ephemeroptera | *Parafronurus youi* | G | G | A | C | G | A | G | A | A | G | A | C | C | C | T | A | T | A | G | A |
| NC011359.1 | Ephemeroptera | *Parafronurus youi* | G | G | A | C | G | A | G | A | A | G | A | C | C | C | T | A | T | A | G | A |
| KM244674.1 | Ephemeroptera | *Potamanthus* sp. | G | G | A | C | G | A | G | A | A | G | A | C | C | C | T | A | T | A | G | A |
| MK951670.1 | Ephemeroptera | *Rhithrogena* sp. | G | G | A | C | G | A | G | A | A | G | A | C | C | C | T | A | T | A | G | A |
| MT274128.1 | Ephemeroptera | *Serratella* sp. | G | G | A | C | G | A | G | A | A | G | A | C | C | C | T | A | T | A | G | A |
| MT274129.1 | Ephemeroptera | *Serratella* sp. | G | G | A | C | G | A | G | A | A | G | A | C | C | C | T | A | T | A | G | A |
| NC050280.1 | Ephemeroptera | *Serratella* sp. | G | G | A | C | G | A | G | A | A | G | A | C | C | C | T | A | T | A | G | A |
| NC050281.1 | Ephemeroptera | *Serratella* sp. | G | G | A | C | G | A | G | A | A | G | A | C | C | C | T | A | T | A | G | A |
| MT274130.1 | Ephemeroptera | *Serratella zapekinae* | G | G | A | C | G | A | G | A | A | G | A | C | C | C | T | A | T | A | G | A |
| NC050282.1 | Ephemeroptera | *Serratella zapekinae* | G | G | A | C | G | A | G | A | A | G | A | C | C | C | T | A | T | A | G | A |
| FJ606783.1 | Ephemeroptera | *Siphlonurus immanis* | G | G | A | C | G | A | G | A | A | G | A | C | C | C | T | A | T | A | G | A |
| NC013822.1 | Ephemeroptera | *Siphlonurus immanis* | G | G | A | C | G | A | G | A | A | G | A | C | C | C | T | A | T | A | G | A |
| KM244684.1 | Ephemeroptera | *Siphlonurus* sp. | G | G | A | C | G | A | G | A | A | G | A | C | C | C | T | A | T | A | G | A |

| | | | | | | | | | | | | | | | | | | | | | |
|---|---|---|---|---|---|---|---|---|---|---|---|---|---|---|---|---|---|---|---|---|---|---|
| HQ875717.1 | Ephemeroptera | *Siphluriscus chinensis* | G | G | A | C | G | A | G | A | A | G | A | C | C | C | T | A | T | A | G | A |
| MT274131.1 | Ephemeroptera | *Torleya grandiforceps* | G | G | A | C | G | A | G | A | A | G | A | C | C | C | T | A | T | A | G | A |
| NC050283.1 | Ephemeroptera | *Torleya grandiforceps* | G | G | A | C | G | A | G | A | A | G | A | C | C | C | T | A | T | A | G | A |
| MT274132.1 | Ephemeroptera | *Torleya nepalica* | G | G | A | C | G | A | G | A | A | G | A | C | C | C | T | A | T | A | G | A |
| NC050284.1 | Ephemeroptera | *Torleya nepalica* | G | G | A | C | G | A | G | A | A | G | A | C | C | C | T | A | T | A | G | A |
| MH751435.1 | Odonata | *Acisoma panorpoides* | A | G | A | C | G | A | G | A | A | G | A | C | C | C | T | A | T | A | G | A |
| NC042689.1 | Odonata | *Acisoma panorpoides* | A | G | A | C | G | A | G | A | A | G | A | C | C | C | T | A | T | A | G | A |
| KX161841.1 | Odonata | *Anax imperator* | A | G | A | C | G | A | G | A | A | G | A | C | C | C | T | A | T | A | G | A |
| NC031821.1 | Odonata | *Anax imperator* | A | G | A | C | G | A | G | A | A | G | A | C | C | C | T | A | T | A | G | A |
| KP233805.1 | Odonata | *Atrocalopteryx atrata* | G | G | A | C | G | A | G | A | A | G | A | C | C | C | T | A | T | A | G | A |
| NC027181.1 | Odonata | *Atrocalopteryx atrata* | G | G | A | C | G | A | G | A | A | G | A | C | C | C | T | A | T | A | G | A |
| MG011692.1 | Odonata | *Atrocalopteryx melli* | G | G | A | C | G | A | G | A | A | G | A | C | C | C | T | A | T | A | G | A |
| NC036749.1 | Odonata | *Atrocalopteryx melli* | G | G | A | C | G | A | G | A | A | G | A | C | C | C | T | A | T | A | G | A |

| | | | | | | | | | | | | | | | | | | | | | | |
|---|---|---|---|---|---|---|---|---|---|---|---|---|---|---|---|---|---|---|---|---|---|---|
| KM658172.1 | Odonata | *Brachythemis contaminata* | A | G | A | C | G | A | G | A | A | G | A | C | C | C | T | A | T | A | G | A |
| NC026305.1 | Odonata | *Brachythemis contaminata* | A | G | A | C | G | A | G | A | A | G | A | C | C | C | T | A | T | A | G | A |
| EU591677.1 | Odonata | *Davidius lunatus* | A | G | A | C | G | A | G | A | A | G | A | C | C | C | T | A | T | A | G | A |
| NC012644.1 | Odonata | *Davidius lunatus* | A | G | A | C | G | A | G | A | A | G | A | C | C | C | T | A | T | A | G | A |
| MH751436.1 | Odonata | *Deielia phaon* | A | G | A | C | G | A | G | A | A | G | A | C | C | C | T | A | T | A | G | A |
| NC042690.1 | Odonata | *Deielia phaon* | A | G | A | C | G | A | G | A | A | G | A | C | C | C | T | A | T | A | G | A |
| MF716899.1 | Odonata | *Enallagma cyathigerum* | G | G | A | C | G | A | G | A | A | G | A | C | C | C | T | A | T | A | G | A |
| JX050223.1 | Odonata | *Epiophlebia superstes* | A | G | A | C | G | A | G | A | A | G | A | C | C | C | T | A | T | A | G | A |
| NC023232.1 | Odonata | *Epiophlebia superstes* | A | G | A | C | G | A | G | A | A | G | A | C | C | C | T | A | T | A | G | A |
| MK522522.1 | Odonata | *Epophthalmia elegans* | A | G | A | C | G | A | G | A | A | G | A | C | C | C | T | A | T | A | G | A |
| NC046756.1 | Odonata | *Epophthalmia elegans* | A | G | A | C | G | A | G | A | A | G | A | C | C | C | T | A | T | A | G | A |
| HM126547.1 | Odonata | *Euphaea formosa* | A | G | A | C | G | A | G | A | A | G | A | C | C | C | T | A | T | A | G | A |
| NC014493.1 | Odonata | *Euphaea formosa* | A | G | A | C | G | A | G | A | A | G | A | C | C | C | T | A | T | A | G | A |
| KM244659.1 | Odonata | *Hydrobasileus croceus* | A | G | A | C | G | A | G | A | A | G | A | C | C | C | T | A | T | A | G | A |

| | | | | | | | | | | | | | | | | | | | | | | |
|---|---|---|---|---|---|---|---|---|---|---|---|---|---|---|---|---|---|---|---|---|---|---|
| NC025758.1 | Odonata | *Hydrobasileus croceus* | A | G | A | C | G | A | G | A | A | G | A | C | C | C | T | A | T | A | G | A |
| KM244673.1 | Odonata | *Ictinogomphus* sp. | A | G | A | C | G | A | G | A | A | G | A | C | C | C | T | A | T | A | G | A |
| KU958378.1 | Odonata | *Ischnura elegans* | G | G | A | C | G | A | G | A | A | G | A | C | C | C | T | A | T | A | G | A |
| MK951668.1 | Odonata | *Ischnura elegans* | G | G | A | C | G | A | G | A | A | G | A | C | C | C | T | A | T | A | G | A |
| NC031824.1 | Odonata | *Ischnura elegans* | G | G | A | C | G | A | G | A | A | G | A | C | C | C | T | A | T | A | G | A |
| KC878732.1 | Odonata | *Ischnura pumilio* | G | G | A | C | G | A | G | A | A | G | A | C | C | C | T | A | T | A | G | A |
| NC021617.1 | Odonata | *Ischnura pumilio* | G | G | A | C | G | A | G | A | A | G | A | C | C | C | T | A | T | A | G | A |
| MF990748.1 | Odonata | *Macromia daimoji* | A | G | A | C | G | A | G | A | A | G | A | C | C | C | T | A | T | A | G | A |
| NC041425.1 | Odonata | *Macromia daimoji* | A | G | A | C | G | A | G | A | A | G | A | C | C | C | T | A | T | A | G | A |
| MK722304.1 | Odonata | *Matrona basilaris* | G | G | A | C | G | A | G | A | A | G | A | C | C | C | T | A | T | A | G | A |
| MK722305.1 | Odonata | *Matrona basilaris* | G | G | A | C | G | A | G | A | A | G | A | C | C | C | T | A | T | A | G | A |
| KU958377.1 | Odonata | *Megaloprepus caerulatus* | G | G | A | C | G | A | G | A | A | G | A | C | C | C | T | A | T | A | G | A |
| NC031823.1 | Odonata | *Megaloprepus caerulatus* | G | G | A | C | G | A | G | A | A | G | A | C | C | C | T | A | T | A | G | A |
| KU871065.1 | Odonata | *Mnais costalis* | A | G | A | C | G | A | G | A | A | G | A | C | C | C | T | A | T | A | G | A |
| KY402222.1 | Odonata | *Nannophya pygmaea* | A | G | A | C | G | A | G | A | A | G | A | C | C | C | T | A | T | A | G | A |
| NC039410.1 | Odonata | *Nannophya pygmaea* | A | G | A | C | G | A | G | A | A | G | A | C | C | C | T | A | T | A | G | A |
| KU361233.1 | Odonata | *Orthetrum chrysis* | A | G | A | C | G | A | G | A | A | G | A | C | C | C | T | A | T | A | G | A |

| | | | | | | | | | | | | | | | | | | | | | | |
|---|---|---|---|---|---|---|---|---|---|---|---|---|---|---|---|---|---|---|---|---|---|---|
| NC032048.1 | Odonata | *Orthetrum chrysis* | A | G | A | C | G | A | G | A | A | G | A | C | C | C | T | A | T | A | G | A |
| KU361232.1 | Odonata | *Orthetrum glaucum* | A | G | A | C | G | A | G | A | A | G | A | C | C | C | T | A | T | A | G | A |
| NC032047.1 | Odonata | *Orthetrum glaucum* | A | G | A | C | G | A | G | A | A | G | A | C | C | C | T | A | T | A | G | A |
| MH751437.1 | Odonata | *Orthetrum melania* | A | G | A | C | G | A | G | A | A | G | A | C | C | C | T | A | T | A | G | A |
| NC042732.1 | Odonata | *Orthetrum melania* | A | G | A | C | G | A | G | A | A | G | A | C | C | C | T | A | T | A | G | A |
| KU361234.1 | Odonata | *Orthetrum sabina* | A | G | A | C | G | A | G | A | A | G | A | C | C | C | T | A | T | A | G | A |
| MH751438.1 | Odonata | *Orthetrum sabina* | A | G | A | C | G | A | G | A | A | G | A | C | C | C | T | A | T | A | G | A |
| NC032049.1 | Odonata | *Orthetrum sabina* | A | G | A | C | G | A | G | A | A | G | A | C | C | C | T | A | T | A | G | A |
| KU361235.1 | Odonata | *Orthetrum testaceum* | A | G | A | C | G | A | G | A | A | G | A | C | C | C | T | A | T | A | G | A |
| MH751439.1 | Odonata | *Orthetrum testaceum* | A | G | A | C | G | A | G | A | A | G | A | C | C | C | T | A | T | A | G | A |
| NC032050.1 | Odonata | *Orthetrum testaceum* | A | G | A | C | G | A | G | A | A | G | A | C | C | C | T | A | T | A | G | A |
| AB126005.1 | Odonata | *Orthetrum triangulare melania* | A | G | A | C | G | A | G | A | A | G | A | C | C | C | T | A | T | A | G | A |
| MK951669.1 | Odonata | *Paracercion v-nigrum* | G | G | A | C | G | A | G | A | A | G | A | C | C | C | T | A | T | A | G | A |
| KP233804.1 | Odonata | *Platycnemis* | G | G | A | C | G | A | G | A | A | G | A | C | C | C | T | A | T | A | G | A |

| | | | | | | | | | | | | | | | | | | | | | | |
|---|---|---|---|---|---|---|---|---|---|---|---|---|---|---|---|---|---|---|---|---|---|---|
| | | *foliacea* | | | | | | | | | | | | | | | | | | | | |
| NC027180.1 | Odonata | *Platycnemis foliacea* | G | G | A | C | G | A | G | A | A | G | A | C | C | C | T | A | T | A | G | A |
| FJ606784.1 | Odonata | *Pseudolestes mirabilis* | A | G | A | C | G | A | G | A | A | G | A | C | C | C | T | A | T | A | G | A |
| NC020636.1 | Odonata | *Pseudolestes mirabilis* | A | G | A | C | G | A | G | A | A | G | A | C | C | C | T | A | T | A | G | A |
| MF150044.1 | Odonata | *Psolodesmus mandarinus* | A | G | A | C | G | A | G | A | A | G | A | C | C | C | T | A | T | A | G | A |
| NC042442.1 | Odonata | *Psolodesmus mandarinus* | A | G | A | C | G | A | G | A | A | G | A | C | C | C | T | A | T | A | G | A |
| MG594801.1 | Odonata | *Somatochlora hineana* | A | G | A | C | G | A | G | A | A | G | A | C | C | C | T | A | T | A | G | A |
| MH751440.1 | Odonata | *Tramea virginia* | A | G | A | C | G | A | G | A | A | G | A | C | C | C | T | A | T | A | G | A |
| NC042691.1 | Odonata | *Tramea virginia* | A | G | A | C | G | A | G | A | A | G | A | C | C | C | T | A | T | A | G | A |
| MH751441.1 | Odonata | *Trigomphus carus* | A | G | A | C | G | A | G | A | A | G | A | C | C | C | T | A | T | A | G | A |
| NC042733.1 | Odonata | *Trigomphus carus* | A | G | A | C | G | A | G | A | A | G | A | C | C | C | T | A | T | A | G | A |
| JX050224.1 | Odonata | *Vestalis melania* | A | G | A | C | G | A | G | A | A | G | A | C | C | C | T | A | T | A | G | A |
| NC023233.1 | Odonata | *Vestalis melania* | A | G | A | C | G | A | G | A | A | G | A | C | C | C | T | A | T | A | G | A |
| KM199685.1 | Plecoptera | *Acroneuria hainana* | G | G | A | C | G | A | G | A | A | G | A | C | C | C | T | A | T | A | G | A |
| NC026104.1 | Plecoptera | *Acroneuria* | G | G | A | C | G | A | G | A | A | G | A | C | C | C | T | A | T | A | G | A |

| | | | | | | | | | | | | | | | | | | | | | | |
|---|---|---|---|---|---|---|---|---|---|---|---|---|---|---|---|---|---|---|---|---|---|---|
| | | *hainana* | | | | | | | | | | | | | | | | | | | | |
| KX091846.1 | Plecoptera | *Agnetina aequalis* | G | G | A | C | G | A | G | A | A | G | A | C | C | C | T | A | T | A | G | A |
| MH085446.1 | Plecoptera | *Amphinemura longispina* | G | G | A | C | G | A | G | A | A | G | A | C | C | C | T | A | T | A | G | A |
| NC044748.1 | Plecoptera | *Amphinemura longispina* | G | G | A | C | G | A | G | A | A | G | A | C | C | C | T | A | T | A | G | A |
| KX091847.1 | Plecoptera | *Amphinemura* sp. | G | G | A | C | G | A | G | A | A | G | A | C | C | C | T | A | T | A | G | A |
| MH085447.1 | Plecoptera | *Amphinemura yao* | G | G | A | C | G | A | G | A | A | G | A | C | C | C | T | A | T | A | G | A |
| NC044749.1 | Plecoptera | *Amphinemura yao* | G | G | A | C | G | A | G | A | A | G | A | C | C | C | T | A | T | A | G | A |
| MK111413.1 | Plecoptera | *Antarctoperla michaelseni* | G | G | A | C | G | A | G | A | A | G | A | C | C | C | C | A | T | A | G | A |
| NC042199.1 | Plecoptera | *Antarctoperla michaelseni* | G | G | A | C | G | A | G | A | A | G | A | C | C | C | C | A | T | A | G | A |
| KR604721.1 | Plecoptera | *Apteroperla tikumana* | G | G | A | C | G | A | G | A | A | G | A | C | C | C | T | A | T | A | G | A |
| NC027698.1 | Plecoptera | *Apteroperla tikumana* | G | G | A | C | G | A | G | A | A | G | A | C | C | C | T | A | T | A | G | A |
| MT410838.1 | Plecoptera | *Brachyptera risi* | G | G | A | C | G | A | G | A | A | G | A | C | C | C | T | A | T | A | G | A |
| MG677941.1 | Plecoptera | *Calineuria stigmatica* | G | G | A | C | G | A | G | A | A | G | A | C | C | C | T | A | T | A | G | A |
| KX094942.1 | Plecoptera | *Capnia zijinshana* | G | G | A | C | G | A | G | A | A | G | A | C | C | C | T | A | T | A | G | A |
| NC034661.1 | Plecoptera | *Capnia zijinshana* | G | G | A | C | G | A | G | A | A | G | A | C | C | C | T | A | T | A | G | A |

| | | | | | | | | | | | | | | | | | | | | | | |
|---|---|---|---|---|---|---|---|---|---|---|---|---|---|---|---|---|---|---|---|---|---|---|
| MG677942.1 | Plecoptera | *Caroperla siveci* | G | G | A | C | G | A | G | A | A | G | A | C | C | C | T | A | T | A | G | A |
| MF100783.1 | Plecoptera | *Cerconychia flectospina* | G | G | A | C | G | A | G | A | A | G | A | C | C | C | T | A | T | A | G | A |
| MN419914.1 | Plecoptera | *Claassenia* sp. | G | G | A | C | G | A | G | A | A | G | A | C | C | C | T | A | T | A | G | A |
| KC952026.1 | Plecoptera | *Cryptoperla* sp. | G | G | A | C | G | A | G | A | A | G | A | C | C | C | T | A | T | A | G | A |
| MK111416.1 | Plecoptera | *Diamphipnoa annulata* | G | G | A | C | G | A | G | A | A | G | A | C | C | C | T | A | T | A | G | A |
| NC042205.1 | Plecoptera | *Diamphipnoa annulata* | G | G | A | C | G | A | G | A | A | G | A | C | C | C | T | A | T | A | G | A |
| MK111417.1 | Plecoptera | *Diamphipnopsis sp.* | G | G | A | C | G | A | G | A | A | G | A | C | C | C | T | A | T | A | G | A |
| KF484757.1 | Plecoptera | *Dinocras cephalotes* | G | G | A | C | G | A | G | A | A | G | A | C | C | C | T | A | T | A | G | A |
| NC022843.1 | Plecoptera | *Dinocras cephalotes* | G | G | A | C | G | A | G | A | A | G | A | C | C | C | T | A | T | A | G | A |
| MG589787.1 | Plecoptera | *Doddsia occidentalis* | G | G | A | C | G | A | G | A | A | G | A | C | C | C | T | A | T | A | G | A |
| MK905888.1 | Plecoptera | *Etrocorema hochii* | G | G | A | C | G | A | G | A | A | G | A | C | C | C | T | A | T | A | G | A |
| MK905206.1 | Plecoptera | *Flavoperla* sp. | G | G | A | C | G | A | G | A | A | G | A | C | C | C | T | A | T | A | G | A |
| MN419916.1 | Plecoptera | *Flavoperla* sp. | G | G | A | C | G | A | G | A | A | G | A | C | C | C | T | A | T | A | G | A |
| MN419915.1 | Plecoptera | *Indonemoura auriformis* | G | G | A | C | G | A | G | A | A | G | A | C | C | C | T | A | T | A | G | A |

| | | | | | | | | | | | | | | | | | | | | | | |
|---|---|---|---|---|---|---|---|---|---|---|---|---|---|---|---|---|---|---|---|---|---|---|
| MH085448.1 | Plecoptera | *Indonemoura jacobsoni* | G | G | A | C | G | A | G | A | A | G | A | C | C | C | T | A | T | A | G | A |
| NC044750.1 | Plecoptera | *Indonemoura jacobsoni* | G | G | A | C | G | A | G | A | A | G | A | C | C | C | T | A | T | A | G | A |
| MH085449.1 | Plecoptera | *Indonemoura nohirae* | G | G | A | C | G | A | G | A | A | G | A | C | C | C | T | A | T | A | G | A |
| NC044751.1 | Plecoptera | *Indonemoura nohirae* | G | G | A | C | G | A | G | A | A | G | A | C | C | C | T | A | T | A | G | A |
| MF716959.1 | Plecoptera | *Isoperla bilineata* | G | G | A | C | G | A | G | A | A | G | A | C | C | C | T | A | T | A | G | A |
| NC038190.1 | Plecoptera | *Isoperla bilineata* | G | G | A | C | G | A | G | A | A | G | A | C | C | C | T | A | T | A | G | A |
| MG910457.1 | Plecoptera | *Isoperla eximia* | G | G | A | C | G | A | G | A | A | G | A | C | C | C | T | A | T | A | G | A |
| NC038167.1 | Plecoptera | *Isoperla eximia* | G | G | A | C | G | A | G | A | A | G | A | C | C | C | T | A | T | A | G | A |
| KT186102.1 | Plecoptera | *Kamimuria chungnanshana* | G | G | A | C | G | A | G | A | A | G | A | C | C | C | T | A | T | A | G | A |
| NC028076.1 | Plecoptera | *Kamimuria chungnanshana* | G | G | A | C | G | A | G | A | A | G | A | C | C | C | T | A | T | A | G | A |
| MN400755.1 | Plecoptera | *Kamimuria klapaleki* | G | G | A | C | G | A | G | A | A | G | A | C | C | C | T | A | T | A | G | A |
| MK111414.1 | Plecoptera | *Klapopteryx armillata* | G | G | A | C | G | A | G | A | A | G | A | C | C | C | T | A | T | A | G | A |
| MH374046.1 | Plecoptera | *Lednia tumana* | G | G | A | C | G | A | G | A | A | G | A | C | C | C | T | A | T | A | G | A |
| MK568475.1 | Plecoptera | *Leuctra fusca* | G | G | A | C | G | A | G | A | A | G | A | C | C | C | T | A | T | A | G | A |

| | | | | | | | | | | | | | | | | | | | | | | |
|---|---|---|---|---|---|---|---|---|---|---|---|---|---|---|---|---|---|---|---|---|---|---|
| KP642637.1 | Plecoptera | *Mesocapnia arizonensis* | G | G | A | C | G | A | G | A | A | G | A | C | C | C | T | A | T | A | G | A |
| KY568983.1 | Plecoptera | *Mesocapnia daxingana* | G | G | A | C | G | A | G | A | A | G | A | C | C | C | T | A | T | A | G | A |
| MH085450.1 | Plecoptera | *Mesonemoura metafiligera* | G | G | A | C | G | A | G | A | A | G | A | C | C | C | T | A | T | A | G | A |
| NC044719.1 | Plecoptera | *Mesonemoura metafiligera* | G | G | A | C | G | A | G | A | A | G | A | C | C | C | T | A | T | A | G | A |
| MH085451.1 | Plecoptera | *Mesonemoura tritaenia* | G | G | A | C | G | A | G | A | A | G | A | C | C | C | T | A | T | A | G | A |
| NC044720.1 | Plecoptera | *Mesonemoura tritaenia* | G | G | A | C | G | A | G | A | A | G | A | C | C | C | T | A | T | A | G | A |
| MN096323.1 | Plecoptera | *Microperla geei* | G | G | A | C | G | A | G | A | A | G | A | C | C | C | T | A | T | A | G | A |
| MT410776.1 | Plecoptera | *Nemoura avicularis* | G | G | A | C | G | A | G | A | A | G | A | C | C | C | T | A | T | A | G | A |
| MT410849.1 | Plecoptera | *Nemoura cinerea* | G | G | A | C | G | A | G | A | A | G | A | C | C | C | T | A | T | A | G | A |
| KY940360.1 | Plecoptera | *Nemoura nankinensis* | G | G | A | C | G | A | G | A | A | G | A | C | C | C | T | A | T | A | G | A |
| NC034939.1 | Plecoptera | *Nemoura nankinensis* | G | G | A | C | G | A | G | A | A | G | A | C | C | C | T | A | T | A | G | A |
| MK290826.1 | Plecoptera | *Nemoura papilla* | G | G | A | C | G | A | G | A | A | G | A | C | C | C | T | A | T | A | G | A |
| MK111418.1 | Plecoptera | *Neonemura* | G | G | A | C | G | A | G | A | A | G | A | C | C | C | T | A | T | A | G | A |

| | | | | | | | | | | | | | | | | | | | | | | |
|---|---|---|---|---|---|---|---|---|---|---|---|---|---|---|---|---|---|---|---|---|---|---|
| | | *barrosi* | | | | | | | | | | | | | | | | | | | | |
| NC042206.1 | Plecoptera | *Neonemura barrosi* | G | G | A | C | G | A | G | A | A | G | A | C | C | C | T | A | T | A | G | A |
| KX091858.1 | Plecoptera | *Neoperla ignacsiveci* | G | G | A | C | G | A | G | A | A | G | A | C | C | C | T | A | T | A | G | A |
| KX091859.1 | Plecoptera | *Neoperla* sp. | G | G | A | C | G | A | G | A | A | G | A | C | C | C | T | A | T | A | G | A |
| MN400756.1 | Plecoptera | *Neoperlops gressitti* | G | G | A | C | G | A | G | A | A | G | A | C | C | C | T | A | T | A | G | A |
| MK111415.1 | Plecoptera | *Neuroperla schedingi* | G | G | A | C | G | A | G | A | A | G | A | C | C | C | T | A | T | A | G | A |
| NC042200.1 | Plecoptera | *Neuroperla schedingi* | G | G | A | C | G | A | G | A | A | G | A | C | C | C | T | A | T | A | G | A |
| MK686067.1 | Plecoptera | *Niponiella limbatella* | G | G | A | C | G | A | G | A | A | G | A | C | C | C | T | A | T | A | G | A |
| MK387068.1 | Plecoptera | *Peltoperlopsis cebuano* | G | G | A | C | G | A | G | A | A | G | A | C | C | C | T | A | T | A | G | A |
| MF197377.1 | Plecoptera | *Perlodes* sp. | G | G | A | C | G | A | G | A | A | G | A | C | C | C | T | A | T | A | G | A |
| MH085452.1 | Plecoptera | *Protonemura kohnoae* | G | G | A | C | G | A | G | A | A | G | A | C | C | C | T | A | T | A | G | A |
| NC044752.1 | Plecoptera | *Protonemura kohnoae* | G | G | A | C | G | A | G | A | A | G | A | C | C | C | T | A | T | A | G | A |
| MT410850.1 | Plecoptera | *Protonemura* | G | G | A | C | G | A | G | A | A | G | A | C | C | C | T | A | T | A | G | A |

| | | | | | | | | | | | | | | | | | | | | | | |
|---|---|---|---|---|---|---|---|---|---|---|---|---|---|---|---|---|---|---|---|---|---|---|
| | | *meyeri* | | | | | | | | | | | | | | | | | | | | |
| NC050322.1 | Plecoptera | *Protonemura meyeri* | G | G | A | C | G | A | G | A | A | G | A | C | C | C | T | A | T | A | G | A |
| MH085453.1 | Plecoptera | *Protonemura orbiculata* | G | G | A | C | G | A | G | A | A | G | A | C | C | C | T | A | T | A | G | A |
| NC044753.1 | Plecoptera | *Protonemura orbiculata* | G | G | A | C | G | A | G | A | A | G | A | C | C | C | T | A | T | A | G | A |
| MG910458.1 | Plecoptera | *Pseudomegarcys japonica* | G | G | A | C | G | A | G | A | A | G | A | C | C | C | T | A | T | A | G | A |
| NC038168.1 | Plecoptera | *Pseudomegarcys japonica* | G | G | A | C | G | A | G | A | A | G | A | C | C | C | T | A | T | A | G | A |
| KU182360.1 | Plecoptera | *Pteronarcella badia* | G | G | A | C | G | A | G | A | A | G | A | C | C | C | T | A | T | A | G | A |
| NC029248.1 | Plecoptera | *Pteronarcella badia* | G | G | A | C | G | A | G | A | A | G | A | C | C | C | T | A | T | A | G | A |
| AY687866.1 | Plecoptera | *Pteronarcys princeps* | G | G | A | C | G | A | G | A | A | G | A | C | C | C | T | A | T | A | G | A |
| NC006133.1 | Plecoptera | *Pteronarcys princeps* | G | G | A | C | G | A | G | A | A | G | A | C | C | C | T | A | T | A | G | A |
| MK111419.1 | Plecoptera | *Rhopalopsole bulbifera* | G | G | A | C | G | A | G | A | A | G | A | C | C | C | T | A | T | A | G | A |
| NC042207.1 | Plecoptera | *Rhopalopsole* | G | G | A | C | G | A | G | A | A | G | A | C | C | C | T | A | T | A | G | A |

| | | | | | | | | | | | | | | | | | | | | | | |
|---|---|---|---|---|---|---|---|---|---|---|---|---|---|---|---|---|---|---|---|---|---|---|
| | | *bulbifera* | | | | | | | | | | | | | | | | | | | | |
| MH510071.1 | Plecoptera | *Scopura longa* | G | G | A | C | G | A | G | A | A | G | A | C | C | C | T | A | T | A | G | A |
| NC041105.1 | Plecoptera | *Scopura longa* | G | G | A | C | G | A | G | A | A | G | A | C | C | C | T | A | T | A | G | A |
| MK492253.1 | Plecoptera | *Sinacroneuria dabieshana* | G | G | A | C | G | A | G | A | A | G | A | C | C | C | T | A | T | A | G | A |
| MF716958.1 | Plecoptera | *Soliperla* sp. | G | G | A | C | G | A | G | A | A | G | A | C | C | C | T | A | T | A | G | A |
| NC038189.1 | Plecoptera | *Soliperla* sp. | G | G | A | C | G | A | G | A | A | G | A | C | C | C | T | A | T | A | G | A |
| MH085455.1 | Plecoptera | *Sphaeronemoura acutispina* | G | G | A | C | G | A | G | A | A | G | A | C | C | C | T | A | T | A | G | A |
| MH085454.1 | Plecoptera | *Sphaeronemoura grandicauda* | G | G | A | C | G | A | G | A | A | G | A | C | C | C | T | A | T | A | G | A |
| NC044754.1 | Plecoptera | *Sphaeronemoura grandicauda* | G | G | A | C | G | A | G | A | A | G | A | C | C | C | T | A | T | A | G | A |
| MK111420.1 | Plecoptera | *Sphaeronemoura hainana* | G | G | A | C | G | A | G | A | A | G | A | C | C | C | T | A | T | A | G | A |
| NC044755.1 | Plecoptera | *Sphaeronemoura hamistyla* | G | G | A | C | G | A | G | A | A | G | A | C | C | C | T | A | T | A | G | A |
| KR088971.1 | Plecoptera | *Styloperla* sp. | G | G | A | C | G | A | G | A | A | G | A | C | C | C | T | A | T | A | G | A |
| KX845569.1 | Plecoptera | *Styloperla spinicercia* | G | G | A | C | G | A | G | A | A | G | A | C | C | C | T | A | T | A | G | A |
| NC034809.1 | Plecoptera | *Styloperla spinicercia* | G | G | A | C | G | A | G | A | A | G | A | C | C | C | T | A | T | A | G | A |

| | | | | | | | | | | | | | | | | | | | | | | |
|---|---|---|---|---|---|---|---|---|---|---|---|---|---|---|---|---|---|---|---|---|---|---|
| KX091865.1 | Plecoptera | *Suwallia bimaculata* | G | G | A | C | G | A | G | A | A | G | A | C | C | C | T | A | T | A | G | A |
| MN121757.1 | Plecoptera | *Suwallia bimaculata* | G | G | A | C | G | A | G | A | A | G | A | C | C | C | T | A | T | A | G | A |
| MF198253.1 | Plecoptera | *Suwallia teleckojensis* | G | G | A | C | G | A | G | A | A | G | A | C | C | C | T | A | T | A | G | A |
| NC037754.1 | Plecoptera | *Suwallia teleckojensis* | G | G | A | C | G | A | G | A | A | G | A | C | C | C | T | A | T | A | G | A |
| MG589786.1 | Plecoptera | *Taeniopteryx ugola* | G | G | A | C | G | A | G | A | A | G | A | C | C | C | T | A | T | A | G | A |
| NC037897.1 | Plecoptera | *Taeniopteryx ugola* | G | G | A | C | G | A | G | A | A | G | A | C | C | C | T | A | T | A | G | A |
| KM409708.1 | Plecoptera | *Togoperla* sp. | G | G | A | C | G | A | G | A | A | G | A | C | C | C | T | A | T | A | G | A |
| KY522907.1 | Plecoptera | *Zelandoperla fenestrata* | G | G | A | C | G | A | G | A | A | G | A | C | C | C | C | A | T | A | G | A |
| NC034997.1 | Plecoptera | *Zelandoperla fenestrata* | G | G | A | C | G | A | G | A | A | G | A | C | C | C | C | A | T | A | G | A |
| FJ456939.1 | Heteroptera | *Aphelocheirus ellipsoideus* | G | G | A | C | G | A | G | A | A | G | A | C | C | C | T | A | T | A | G | A |
| KX505849.1 | Heteroptera | *Aphelocheirus jendeki* | G | G | A | C | G | A | G | A | A | G | A | C | C | C | T | A | T | A | G | A |
| NC037370.1 | Heteroptera | *Aphelocheirus* | G | G | A | C | G | A | G | A | A | G | A | C | C | C | T | A | T | A | G | A |

| | | | | | | | | | | | | | | | | | | | | | | |
|---|---|---|---|---|---|---|---|---|---|---|---|---|---|---|---|---|---|---|---|---|---|---|
| | | *jendeki* | | | | | | | | | | | | | | | | | | | | |
| FJ456944.1 | Heteroptera | *Aquarius paludum* | G | G | A | C | G | A | G | A | A | G | A | C | C | C | T | A | T | A | G | A |
| NC012841.1 | Heteroptera | *Aquarius paludum* | G | G | A | C | G | A | G | A | A | G | A | C | C | C | T | A | T | A | G | A |
| MK251109.1 | Heteroptera | *Diplonychus esakii* | G | G | A | C | G | A | G | A | A | G | A | C | C | C | T | A | T | A | G | A |
| FJ456940.1 | Heteroptera | *Diplonychus rusticus* | G | G | A | C | G | A | G | A | A | G | A | C | C | C | T | A | T | A | G | A |
| FJ456949.1 | Heteroptera | *Enithares tibialis* | G | G | A | C | G | A | G | A | A | G | A | C | C | C | T | A | T | A | G | A |
| NC012819.1 | Heteroptera | *Enithares tibialis* | G | G | A | C | G | A | G | A | A | G | A | C | C | C | T | A | T | A | G | A |
| KP400582.1 | Heteroptera | *Entomovelia* sp. | G | G | A | C | G | A | G | A | A | G | A | C | C | C | T | A | T | A | G | A |
| MF177288.1 | Heteroptera | *Gigantometra gigas* | G | G | A | C | G | A | G | A | A | G | A | C | C | C | T | A | T | A | G | A |
| MF177289.1 | Heteroptera | *Gigantometra gigas* | G | G | A | C | G | A | G | A | A | G | A | C | C | C | T | A | T | A | G | A |
| MF177290.1 | Heteroptera | *Gigantometra gigas* | G | G | A | C | G | A | G | A | A | G | A | C | C | C | T | A | T | A | G | A |
| MF177291.1 | Heteroptera | *Gigantometra gigas* | G | G | A | C | G | A | G | A | A | G | A | C | C | C | T | A | T | A | G | A |
| NC041084.1 | Heteroptera | *Gigantometra gigas* | G | G | A | C | G | A | G | A | A | G | A | C | C | C | T | A | T | A | G | A |
| KJ027513.1 | Heteroptera | *Helotrephes semiglobosus semiglobosus* | G | G | A | C | G | A | T | A | A | G | A | C | C | C | T | A | T | A | G | A |

| | | | | | | | | | | | | | | | | | | | | | | |
|---|---|---|---|---|---|---|---|---|---|---|---|---|---|---|---|---|---|---|---|---|---|---|
| FJ456951.1 | Heteroptera | *Helotrephes* sp. | G | G | A | C | G | A | T | A | A | G | A | C | C | C | T | A | T | A | G | A |
| NC012822.1 | Heteroptera | *Helotrephes* sp. | G | G | A | C | G | A | T | A | A | G | A | C | C | C | T | A | T | A | G | A |
| FJ456945.1 | Heteroptera | *Hydrometra greeni* | G | G | A | C | G | A | G | A | A | G | A | C | C | C | T | A | T | A | G | A |
| NC012842.1 | Heteroptera | *Hydrometra greeni* | G | G | A | C | G | A | G | A | A | G | A | C | C | C | T | A | T | A | G | A |
| FJ456947.1 | Heteroptera | *Ilyocoris cimicoides* | G | G | A | C | G | A | G | A | A | G | A | C | C | C | T | A | T | A | G | A |
| NC012845.1 | Heteroptera | *Ilyocoris cimicoides* | G | G | A | C | G | A | G | A | A | G | A | C | C | C | T | A | T | A | G | A |
| FJ456948.1 | Heteroptera | *Laccotrephes robustus* | G | G | A | C | G | A | G | A | A | G | A | C | C | C | T | A | T | A | G | A |
| NC012817.1 | Heteroptera | *Laccotrephes robustus* | G | G | A | C | G | A | G | A | A | G | A | C | C | C | T | A | T | A | G | A |
| FJ456946.1 | Heteroptera | *Leptopus* sp. | G | G | A | C | G | A | G | A | A | G | A | C | C | C | T | A | T | A | G | A |
| JQ910985.1 | Heteroptera | *Lethocerus deyrollei* | G | G | A | C | G | A | G | A | A | G | A | C | C | C | T | G | T | A | G | A |
| KU237288.1 | Heteroptera | *Lethocerus deyrollei* | G | G | A | C | G | A | G | A | A | G | A | C | C | C | T | G | T | A | G | A |
| KM588201.1 | Heteroptera | *Lethocerus indicus* | G | G | A | C | G | A | G | A | A | G | A | C | C | C | T | G | T | A | G | A |
| KR072671.1 | Heteroptera | *Lethocerus indicus* | G | G | A | C | G | A | G | A | A | G | A | C | C | C | T | G | T | A | G | A |
| NC027194.1 | Heteroptera | *Lethocerus indicus* | G | G | A | C | G | A | G | A | A | G | A | C | C | C | T | G | T | A | G | A |
| FJ456942.1 | Heteroptera | *Lycorma delicatula* | G | G | A | C | G | A | T | A | A | G | A | C | C | C | T | A | T | A | G | A |
| NC012835.1 | Heteroptera | *Lycorma delicatula* | G | G | A | C | G | A | T | A | A | G | A | C | C | C | T | A | T | A | G | A |

| | | | | | | | | | | | | | | | | | | | | | | |
|---|---|---|---|---|---|---|---|---|---|---|---|---|---|---|---|---|---|---|---|---|---|---|
| MN027278.1 | Heteroptera | *Metrocoris* sp. | A | G | A | C | G | A | G | A | A | G | A | C | C | C | T | T | T | A | G | A |
| MH382835.1 | Heteroptera | *Micronecta sahlbergii* | G | G | A | C | G | A | G | A | A | G | A | C | C | C | T | A | T | A | G | A |
| NC039588.1 | Heteroptera | *Micronecta sahlbergii* | G | G | A | C | G | A | G | A | A | G | A | C | C | C | T | A | T | A | G | A |
| KT031802.1 | Heteroptera | *Nepa hoffmanni* | G | G | A | C | G | A | G | A | A | G | A | C | C | C | T | G | T | A | G | A |
| NC028084.1 | Heteroptera | *Nepa hoffmanni* | G | G | A | C | G | A | G | A | A | G | A | C | C | C | T | G | T | A | G | A |
| FJ456943.1 | Heteroptera | *Nerthra indica* | G | G | A | C | G | A | G | A | A | G | A | C | C | C | T | A | T | A | G | A |
| NC012838.1 | Heteroptera | *Nerthra indica* | G | G | A | C | G | A | G | A | A | G | A | C | C | C | T | A | T | A | G | A |
| KX034086.1 | Heteroptera | *Notonecta chinensis* | G | G | A | C | G | A | G | A | A | G | A | C | C | C | T | A | T | A | G | A |
| NC036671.1 | Heteroptera | *Notonecta chinensis* | G | G | A | C | G | A | G | A | A | G | A | C | C | C | T | A | T | A | G | A |
| MH780861.1 | Heteroptera | *Notonecta montandoni* | G | G | A | C | G | A | G | A | A | G | A | C | C | C | T | A | T | A | G | A |
| NC045886.1 | Heteroptera | *Notonecta montandoni* | G | G | A | C | G | A | G | A | A | G | A | C | C | C | T | A | T | A | G | A |
| KM244707.1 | Heteroptera | *Notonectidae* sp. | G | G | A | C | G | A | G | A | A | G | A | C | C | C | T | A | T | A | G | A |
| FJ456950.1 | Heteroptera | *Ochterus marginatus* | G | G | A | C | G | A | G | A | A | G | A | C | C | C | T | A | T | A | G | A |
| KJ027516.1 | Heteroptera | *Paraplea frontalis* | G | G | A | C | G | A | G | A | A | G | A | C | C | C | T | A | T | A | G | A |
| NC028629.1 | Heteroptera | *Paraplea frontalis* | G | G | A | C | G | A | G | A | A | G | A | C | C | C | T | A | T | A | G | A |

| | | | | | | | | | | | | | | | | | | | | | | |
|---|---|---|---|---|---|---|---|---|---|---|---|---|---|---|---|---|---|---|---|---|---|---|
| JQ910988.1 | Heteroptera | *Perittopus* sp. | G | G | A | C | G | A | G | A | A | G | A | C | C | C | T | A | T | A | G | A |
| KP400583.1 | Heteroptera | *Ptilomera tigrina* | G | G | A | C | G | A | G | A | A | G | A | C | C | C | T | A | T | A | G | A |
| MN027271.1 | Heteroptera | *Rhagovelia reitteri* | G | G | A | C | G | A | G | A | A | G | A | C | C | C | T | A | T | A | G | A |
| NC044721.1 | Heteroptera | *Rhagovelia reitteri* | G | G | A | C | G | A | G | A | A | G | A | C | C | C | T | A | T | A | G | A |
| MN027279.1 | Heteroptera | *Rhagovelia* sp. | G | G | A | C | G | A | G | A | A | G | A | C | C | C | T | A | T | A | G | A |
| FJ456941.1 | Heteroptera | *Sigara septemlineata* | G | G | A | C | G | A | G | A | A | G | A | C | C | C | T | A | T | A | G | A |
| FJ456952.1 | Heteroptera | *Valentia hoffmanni* | G | G | A | C | G | A | G | A | A | G | A | C | C | C | T | A | T | A | G | A |
| NC012823.1 | Heteroptera | *Valentia hoffmanni* | G | G | A | C | G | A | G | A | A | G | A | C | C | C | T | A | T | A | G | A |
| KF840564.1 | Megaloptera | *Acanthacorydalis orientalis* | G | G | A | C | G | A | G | A | A | G | A | C | C | C | T | A | T | A | G | A |
| NC023462.1 | Megaloptera | *Acanthacorydalis orientalis* | G | G | A | C | G | A | G | A | A | G | A | C | C | C | T | A | T | A | G | A |
| KU925864.1 | Megaloptera | *Archichauliodes deceptor* | G | G | A | C | G | A | G | A | A | G | A | C | C | C | T | A | T | A | G | A |
| NC035504.1 | Megaloptera | *Archichauliodes deceptor* | G | G | A | C | G | A | G | A | A | G | A | C | C | C | T | A | T | A | G | A |
| KU302713.1 | Megaloptera | *Chloronia mirifica* | G | G | A | C | G | A | G | A | A | G | A | C | C | C | T | A | T | A | G | A |
| NC033349.1 | Megaloptera | *Chloronia mirifica* | G | G | A | C | G | A | G | A | A | G | A | C | C | C | T | A | T | A | G | A |
| FJ171323.1 | Megaloptera | *Corydalus cornutus* | G | G | A | C | G | A | G | A | A | G | A | C | C | C | T | A | T | A | G | A |
| NC011276.1 | Megaloptera | *Corydalus* | G | G | A | C | G | A | G | A | A | G | A | C | C | C | T | A | T | A | G | A |

| | | | | | | | | | | | | | | | | | | | | | | |
|---|---|---|---|---|---|---|---|---|---|---|---|---|---|---|---|---|---|---|---|---|---|---|
| | | *cornutus* | | | | | | | | | | | | | | | | | | | | |
| KJ806318.1 | Megaloptera | *Dysmicohermes ingens* | G | G | A | C | G | A | G | A | A | G | A | C | C | C | T | A | T | A | G | A |
| NC024657.1 | Megaloptera | *Dysmicohermes ingens* | G | G | A | C | G | A | G | A | A | G | A | C | C | C | T | A | T | A | G | A |
| JQ351950.1 | Megaloptera | *Neochauliodes bowringi* | A | G | A | C | G | A | G | A | A | G | A | C | C | C | T | A | T | A | G | A |
| NC023444.1 | Megaloptera | *Neochauliodes bowringi* | A | G | A | C | G | A | G | A | A | G | A | C | C | C | T | A | T | A | G | A |
| KF785799.1 | Megaloptera | *Neochauliodes fraternus* | G | G | A | C | G | A | G | A | A | G | A | C | C | C | T | A | T | A | G | A |
| NC025282.1 | Megaloptera | *Neochauliodes fraternus* | G | G | A | C | G | A | G | A | A | G | A | C | C | C | T | A | T | A | G | A |
| KX821680.1 | Megaloptera | *Neochauliodes parasparsus* | A | G | A | C | G | A | G | A | A | G | A | C | C | C | T | A | T | A | G | A |
| JX110703.1 | Megaloptera | *Neochauliodes punctatolosus* | A | G | A | C | G | A | G | A | A | G | A | C | C | C | T | A | T | A | G | A |
| MK301249.1 | Megaloptera | *Neochauliodes punctatolosus* | A | G | A | C | G | A | G | A | A | G | A | C | C | C | T | A | T | A | G | A |
| NC018772.1 | Megaloptera | *Neochauliodes punctatolosus* | A | G | A | C | G | A | G | A | A | G | A | C | C | C | T | A | T | A | G | A |
| KF771894.1 | Megaloptera | *Neochauliodes* | G | G | A | C | G | A | G | A | A | G | A | C | C | C | T | A | T | A | G | A |

| | | | | | | | | | | | | | | | | | | | | | | |
|---|---|---|---|---|---|---|---|---|---|---|---|---|---|---|---|---|---|---|---|---|---|---|
| | | *rotundatus* | | | | | | | | | | | | | | | | | | | | |
| NC025281.1 | Megaloptera | *Neochauliodes rotundatus* | G | G | A | C | G | A | G | A | A | G | A | C | C | C | T | A | T | A | G | A |
| KP126231.1 | Megaloptera | *Neoneuromus tonkinensis* | G | G | A | C | G | A | G | A | A | G | A | C | C | C | T | A | T | A | G | A |
| NC027852.1 | Megaloptera | *Neoneuromus tonkinensis* | G | G | A | C | G | A | G | A | A | G | A | C | C | C | T | A | T | A | G | A |
| KP126232.1 | Megaloptera | *Nevromus exterior* | G | G | A | C | G | A | G | A | A | G | A | C | C | C | T | A | T | A | G | A |
| NC027851.1 | Megaloptera | *Nevromus exterior* | G | G | A | C | G | A | G | A | A | G | A | C | C | C | T | A | T | A | G | A |
| KY230493.1 | Megaloptera | *Protochauliodes biconicus* | A | G | A | C | G | A | G | A | A | G | A | C | C | C | T | A | T | A | G | A |
| EU526394.1 | Megaloptera | *Protohermes concolorus* | G | G | A | C | G | A | G | A | A | G | A | C | C | C | T | A | T | A | G | A |
| NC011524.1 | Megaloptera | *Protohermes concolorus* | G | G | A | C | G | A | G | A | A | G | A | C | C | C | T | A | T | A | G | A |
| FJ859905.1 | Megaloptera | *Sialis hamata* | A | G | A | C | G | A | G | A | A | G | A | C | C | C | T | A | T | A | G | A |
| NC013256.1 | Megaloptera | *Sialis hamata* | A | G | A | C | G | A | G | A | A | G | A | C | C | C | T | A | T | A | G | A |
| KT876911.1 | Megaloptera | *Sialis* sp. | A | G | A | C | G | A | G | A | A | G | A | C | C | C | T | A | T | A | G | A |
| KT425084.1 | Megaloptera | *Stenosialis australiensis* | A | G | A | C | G | A | G | A | A | G | A | C | C | C | T | A | T | A | G | A |
| KY587199.1 | Neuroptera | *Abachrysa eureka* | G | G | A | C | G | A | G | A | A | G | A | C | C | C | T | A | T | A | G | A |
| NC015095.1 | Neuroptera | *Apochrysa* | G | G | A | C | G | A | G | A | A | G | A | C | C | C | T | A | T | A | G | A |

| | | | | | | | | | | | | | | | | | | | | | | |
|---|---|---|---|---|---|---|---|---|---|---|---|---|---|---|---|---|---|---|---|---|---|---|
| | | *matsumurae* | | | | | | | | | | | | | | | | | | | | |
| KC758703.1 | Neuroptera | *Ascalohybris subjacens* | G | G | A | C | G | A | G | A | A | G | A | C | C | C | T | A | T | A | G | A |
| NC021428.1 | Neuroptera | *Ascalohybris subjacens* | G | G | A | C | G | A | G | A | A | G | A | C | C | C | T | A | T | A | G | A |
| FJ171324.1 | Neuroptera | *Ascaloptynx appendiculatus* | G | G | A | C | G | A | G | A | A | G | A | C | C | C | T | A | T | A | G | A |
| NC011277.1 | Neuroptera | *Ascaloptynx appendiculatus* | G | G | A | C | G | A | G | A | A | G | A | C | C | C | T | A | T | A | G | A |
| KT425083.1 | Neuroptera | *Balmes birmanus* | A | G | A | C | G | A | G | A | A | G | A | C | C | C | T | A | T | A | G | A |
| KX369241.1 | Neuroptera | *Bullanga florida* | G | G | A | C | G | A | G | A | A | G | A | C | C | C | T | A | T | A | G | A |
| NC032298.1 | Neuroptera | *Bullanga florida* | G | G | A | C | G | A | G | A | A | G | A | C | C | C | T | A | T | A | G | A |
| KT425069.1 | Neuroptera | *Chasmoptera huttii* | G | G | A | C | G | A | G | A | A | G | A | C | C | C | T | A | T | A | G | A |
| JX033119.1 | Neuroptera | *Chrysopa pallens* | G | G | A | C | G | A | G | A | A | G | A | C | C | C | T | A | T | A | G | A |
| NC019618.1 | Neuroptera | *Chrysopa pallens* | G | G | A | C | G | A | G | A | A | G | A | C | C | C | T | A | T | A | G | A |
| KJ778879.1 | Neuroptera | *Chrysoperla carnea* | G | G | A | C | G | A | G | A | A | G | A | C | C | C | T | A | T | A | G | A |
| KU877169.1 | Neuroptera | *Chrysoperla externa* | G | G | A | C | G | A | G | A | A | G | A | C | C | C | T | A | T | A | G | A |
| NC030341.1 | Neuroptera | *Chrysoperla externa* | G | G | A | C | G | A | G | A | A | G | A | C | C | C | T | A | T | A | G | A |
| KP027407.1 | Neuroptera | *Chrysoperla* | G | G | A | C | G | A | G | A | A | G | A | C | C | C | T | A | T | A | G | A |

| | | | | | | | | | | | | | | | | | | | | | | |
|---|---|---|---|---|---|---|---|---|---|---|---|---|---|---|---|---|---|---|---|---|---|---|
| | | *nipponensis* | | | | | | | | | | | | | | | | | | | | |
| NC015093.1 | Neuroptera | *Chrysoperla nipponensis* | G | G | A | C | G | A | G | A | A | G | A | C | C | C | T | A | T | A | G | A |
| KT425088.1 | Neuroptera | *Climacia areolaris* | A | G | A | C | G | A | G | A | A | G | A | C | C | C | T | A | T | A | G | A |
| KT425078.1 | Neuroptera | *Coniopteryx* sp. | A | G | A | C | G | A | G | A | A | G | A | C | C | C | T | A | T | A | G | A |
| KT425068.1 | Neuroptera | *Dendroleon pantherinus* | G | G | A | C | G | A | G | A | A | G | A | C | C | C | T | A | T | A | G | A |
| MK301246.1 | Neuroptera | *Dendroleon pantherinus* | G | G | A | C | G | A | G | A | A | G | A | C | C | C | T | A | T | A | G | A |
| KT425073.1 | Neuroptera | *Dilar* sp. | A | G | A | C | G | A | G | A | A | G | A | C | C | C | T | A | T | A | G | A |
| FJ859906.1 | Neuroptera | *Ditaxis biseriata* | G | G | A | C | G | A | G | A | A | G | A | C | C | C | T | A | T | A | G | A |
| NC013257.1 | Neuroptera | *Ditaxis biseriata* | G | G | A | C | G | A | G | A | A | G | A | C | C | C | T | A | T | A | G | A |
| KT425087.1 | Neuroptera | *Drepanepteryx phalaenoides* | A | G | A | C | G | A | G | A | A | G | A | C | C | C | T | A | T | A | G | A |
| KF701327.1 | Neuroptera | *Epacanthaclisis banksi* | G | G | A | C | G | A | G | A | A | G | A | C | C | C | T | A | T | A | G | A |
| NC025905.1 | Neuroptera | *Epacanthaclisis banksi* | G | G | A | C | G | A | G | A | A | G | A | C | C | C | T | A | T | A | G | A |
| MF972496.1 | Neuroptera | *Euclimacia badia* | G | G | A | C | G | A | G | A | A | G | A | C | C | C | T | A | T | A | G | A |
| NC039773.1 | Neuroptera | *Euclimacia badia* | G | G | A | C | G | A | G | A | A | G | A | C | C | C | T | A | T | A | G | A |
| KT425080.1 | Neuroptera | *Eumantispa harmandi* | G | G | A | C | G | A | G | A | A | G | A | C | C | C | T | A | T | A | G | A |

| | | | | | | | | | | | | | | | | | | | | | | |
|---|---|---|---|---|---|---|---|---|---|---|---|---|---|---|---|---|---|---|---|---|---|---|
| MF972495.1 | Neuroptera | *Eumantispa harmandi* | G | G | A | C | G | A | G | A | A | G | A | C | C | C | T | A | T | A | G | A |
| NC039772.1 | Neuroptera | *Eumantispa harmandi* | G | G | A | C | G | A | G | A | A | G | A | C | C | C | T | A | T | A | G | A |
| KT425072.1 | Neuroptera | *Fontecilla graphicus* | G | G | A | C | G | A | G | A | A | G | A | C | C | C | T | A | T | A | G | A |
| KY364372.1 | Neuroptera | *Gatzara jezoensis* | G | G | A | C | G | A | G | A | A | G | A | C | C | C | T | A | T | A | G | A |
| MK341582.1 | Neuroptera | *Gryposmylus pennyi* | G | G | A | C | G | A | G | A | A | G | A | C | C | C | T | A | T | A | G | A |
| MN852445.1 | Neuroptera | *Hemerobius japonicus* | A | G | A | C | G | A | G | A | A | G | A | C | C | C | T | A | T | A | G | A |
| NC048517.1 | Neuroptera | *Hemerobius japonicus* | A | G | A | C | G | A | G | A | A | G | A | C | C | C | T | A | T | A | G | A |
| KT425077.1 | Neuroptera | *Heterosmylus* sp. | G | G | A | C | G | A | G | A | A | G | A | C | C | C | T | A | T | A | G | A |
| KT425085.1 | Neuroptera | *Inocellia fujiana* | G | G | A | C | G | A | G | A | A | G | A | C | C | C | T | A | T | A | G | A |
| KY587200.1 | Neuroptera | *Italochrysa insignis* | G | G | A | C | G | A | G | A | A | G | A | C | C | C | T | A | T | A | G | A |
| KY587201.1 | Neuroptera | *Leucochrysa pretiosa* | G | G | A | C | G | A | G | A | A | G | A | C | C | C | T | A | T | A | G | A |
| NC015609.1 | Neuroptera | *Libelloides macaronius* | G | G | A | C | G | A | G | A | A | G | A | C | C | C | T | A | T | A | G | A |
| MF972494.1 | Neuroptera | *Mantispa japonica* | G | G | A | C | G | A | G | A | A | G | A | C | C | C | T | A | T | A | G | A |

| | | | | | | | | | | | | | | | | | | | | | | |
|---|---|---|---|---|---|---|---|---|---|---|---|---|---|---|---|---|---|---|---|---|---|---|
| NC039771.1 | Neuroptera | *Mantispa japonica* | G | G | A | C | G | A | G | A | A | G | A | C | C | C | T | A | T | A | G | A |
| MK472073.1 | Neuroptera | *Micromus angulatus* | A | G | A | C | G | A | G | A | A | G | A | C | C | C | T | A | T | A | G | A |
| NC042677.1 | Neuroptera | *Micromus angulatus* | A | G | A | C | G | A | G | A | A | G | A | C | C | C | T | A | T | A | G | A |
| KT425075.1 | Neuroptera | *Micromus* sp. | A | G | A | C | G | A | G | A | A | G | A | C | C | C | T | A | T | A | G | A |
| KT425092.1 | Neuroptera | *Mucroberotha vesicaria* | A | G | A | C | G | A | G | A | A | G | A | C | C | C | T | A | T | A | G | A |
| KT425089.1 | Neuroptera | *Myiodactylus osmyloides* | A | G | A | C | G | A | G | A | A | G | A | C | C | C | T | A | T | A | G | A |
| MT118664.1 | Neuroptera | *Myrmeleon formicarius* | G | G | A | C | G | A | G | A | A | G | A | C | C | C | T | A | T | A | G | A |
| NC047285.1 | Neuroptera | *Myrmeleon formicarius* | G | G | A | C | G | A | G | A | A | G | A | C | C | C | T | A | T | A | G | A |
| KJ461323.1 | Neuroptera | *Myrmeleon immanis* | G | G | A | C | G | A | G | A | A | G | A | C | C | C | T | A | T | A | G | A |
| NC024826.1 | Neuroptera | *Myrmeleon immanis* | G | G | A | C | G | A | G | A | A | G | A | C | C | C | T | A | T | A | G | A |
| KT425071.1 | Neuroptera | *Nallachius americanus* | A | G | A | C | G | A | G | A | A | G | A | C | C | C | T | A | T | A | G | A |
| KT425086.1 | Neuroptera | *Negha inflata* | G | G | A | C | G | A | G | A | A | G | A | C | C | C | T | A | T | A | G | A |
| KT425079.1 | Neuroptera | *Nemoptera coa* | G | G | A | C | G | A | G | A | A | G | A | C | C | C | T | A | T | A | G | A |

| | | | | | | | | | | | | | | | | | | | | | | |
|---|---|---|---|---|---|---|---|---|---|---|---|---|---|---|---|---|---|---|---|---|---|---|
| KR078257.1 | Neuroptera | *Neuronema laminatum* | A | G | A | C | G | A | G | A | A | G | A | C | C | C | T | A | T | A | G | A |
| NC028153.1 | Neuroptera | *Neuronema laminatum* | A | G | A | C | G | A | G | A | A | G | A | C | C | C | T | A | T | A | G | A |
| KT425074.1 | Neuroptera | *Nevrorthus apatelios* | A | G | A | C | G | A | G | A | A | G | A | C | C | C | T | A | T | A | G | A |
| KT425076.1 | Neuroptera | *Nipponeurorthus fuscinervis* | A | G | A | C | G | A | G | A | A | G | A | C | C | C | T | A | T | A | G | A |
| KP264629.1 | Neuroptera | *Nothancyla verreauxi* | G | G | A | C | G | A | G | A | A | G | A | C | C | C | T | A | T | A | G | A |
| KP264630.1 | Neuroptera | *Nothochrysa* sp. | A | G | A | C | G | A | G | A | A | G | A | C | C | C | T | A | T | A | G | A |
| KJ461322.1 | Neuroptera | *Nymphes myrmeleonoides* | G | G | A | C | G | A | G | A | A | G | A | C | C | C | T | A | T | A | G | A |
| NC024825.1 | Neuroptera | *Nymphes myrmeleonoides* | G | G | A | C | G | A | G | A | A | G | A | C | C | C | T | A | T | A | G | A |
| KT425090.1 | Neuroptera | *Oliarces clara* | G | G | A | C | G | A | G | A | A | G | A | C | C | C | T | A | T | A | G | A |
| MN818868.1 | Neuroptera | *Osmylus atomatus* | G | G | A | C | G | A | G | A | A | G | A | C | C | C | T | A | T | A | G | A |
| MN818867.1 | Neuroptera | *Osmylus fulvicephalus* | G | G | A | C | G | A | G | A | A | G | A | C | C | C | T | A | T | A | G | A |
| MN617021.1 | Neuroptera | *Osmylus lucalatus* | G | G | A | C | G | A | G | A | A | G | A | C | C | C | T | A | T | A | G | A |
| MN818870.1 | Neuroptera | *Osmylus* sp. | G | G | A | C | G | A | G | A | A | G | A | C | C | C | T | A | T | A | G | A |
| MN818869.1 | Neuroptera | *Osmylus zheanus* | G | G | A | C | G | A | G | A | A | G | A | C | C | C | T | A | T | A | G | A |

| | | | | | | | | | | | | | | | | | | | | | | |
|---|---|---|---|---|---|---|---|---|---|---|---|---|---|---|---|---|---|---|---|---|---|---|
| KY587202.1 | Neuroptera | *Parankylopteryx* sp. | A | G | A | C | G | A | G | A | A | G | A | C | C | C | T | A | T | A | G | A |
| KT425091.1 | Neuroptera | *Podallea* sp. | G | G | A | C | G | A | G | A | A | G | A | C | C | C | T | A | T | A | A | A |
| FJ171325.1 | Neuroptera | *Polystoechotes punctatus* | G | G | A | C | G | A | G | A | A | G | A | C | C | C | T | A | T | A | G | A |
| NC011278.1 | Neuroptera | *Polystoechotes punctatus* | G | G | A | C | G | A | G | A | A | G | A | C | C | C | T | A | T | A | G | A |
| KT425082.1 | Neuroptera | *Psychopsis coelivaga* | A | G | A | C | G | A | G | A | A | G | A | C | C | C | T | A | T | A | G | A |
| KF626446.1 | Neuroptera | *Rapisma xizangense* | G | G | A | C | G | A | G | A | A | G | A | C | C | C | T | A | T | A | G | A |
| NC023362.1 | Neuroptera | *Rapisma xizangense* | G | G | A | C | G | A | G | A | A | G | A | C | C | C | T | A | T | A | G | A |
| KF626447.1 | Neuroptera | *Rapisma zayuanum* | G | G | A | C | G | A | G | A | A | G | A | C | C | C | T | A | T | A | G | A |
| NC023363.1 | Neuroptera | *Rapisma zayuanum* | G | G | A | C | G | A | G | A | A | G | A | C | C | C | T | A | T | A | G | A |
| KT425067.1 | Neuroptera | *Semidalis aleyrodiformis* | A | G | A | C | G | A | G | A | A | G | A | C | C | C | T | A | T | A | G | A |
| KT425070.1 | Neuroptera | *Sisyra nigra* | A | G | A | C | G | A | G | A | A | G | A | C | C | C | T | A | T | A | G | A |
| KT425081.1 | Neuroptera | *Stenobiella* sp. | G | G | A | C | G | A | G | A | A | G | A | C | C | C | T | A | T | A | A | A |
| MH361300.1 | Neuroptera | *Suhpalacsa* | G | G | A | C | G | A | G | A | A | G | A | C | C | C | T | A | T | A | G | A |

| | | | | | | | | | | | | | | | | | | | | | | |
|---|---|---|---|---|---|---|---|---|---|---|---|---|---|---|---|---|---|---|---|---|---|---|
| | | *longialata* | | | | | | | | | | | | | | | | | | | | |
| NC039948.1 | Neuroptera | *Suhpalacsa longialata* | G | G | A | C | G | A | G | A | A | G | A | C | C | C | T | A | T | A | G | A |
| MK301247.1 | Neuroptera | *Suhpalacsa* sp. | G | G | A | C | G | A | G | A | A | G | A | C | C | C | T | A | T | A | G | A |
| MK408756.1 | Neuroptera | *Thaumatosmylus hainanus* | G | G | A | C | G | A | G | A | A | G | A | C | C | C | T | A | T | A | G | A |
| NC046578.1 | Neuroptera | *Thaumatosmylus hainanus* | G | G | A | C | G | A | G | A | A | G | A | C | C | C | T | A | T | A | G | A |
| KC515397.1 | Neuroptera | *Thyridosmylus langii* | G | G | A | C | G | A | G | A | A | G | A | C | C | C | T | A | T | A | G | A |
| NC021415.1 | Neuroptera | *Thyridosmylus langii* | G | G | A | C | G | A | G | A | A | G | A | C | C | C | T | A | T | A | G | A |
| MH020192.1 | Coleoptera | *Abscondita anceyi* | A | G | A | C | G | A | G | A | A | G | A | C | C | C | T | A | T | A | G | A |
| NC039706.1 | Coleoptera | *Abscondita anceyi* | A | G | A | C | G | A | G | A | A | G | A | C | C | C | T | A | T | A | G | A |
| MK122952.1 | Coleoptera | *Abscondita chinensis* | A | G | A | C | G | A | G | A | A | G | A | C | C | C | T | A | T | A | G | A |
| MK292092.1 | Coleoptera | *Abscondita terminalis* | A | G | A | C | G | A | G | A | A | G | A | C | C | C | T | A | T | A | G | A |
| NC044776.1 | Coleoptera | *Abscondita terminalis* | A | G | A | C | G | A | G | A | A | G | A | C | C | C | T | A | T | A | G | A |
| KT876878.1 | Coleoptera | *Acilius sulcatus* | A | G | A | C | G | A | G | A | A | G | A | C | C | C | T | A | T | A | G | A |
| KX758085.1 | Coleoptera | *Aquatica ficta* | A | G | A | C | G | A | G | A | A | G | A | C | C | C | T | A | T | A | G | A |

| | | | | | | | | | | | | | | | | | | | | | | |
|---|---|---|---|---|---|---|---|---|---|---|---|---|---|---|---|---|---|---|---|---|---|---|
| NC035060.1 | Coleoptera | *Aquatica ficta* | A | G | A | C | G | A | G | A | A | G | A | C | C | C | T | A | T | A | G | A |
| NC035755.1 | Coleoptera | *Aquatica lateralis* | A | G | A | C | G | A | G | A | A | G | A | C | C | C | T | A | T | A | G | A |
| KF667531.1 | Coleoptera | *Aquatica leii* | A | G | A | C | G | A | G | A | A | G | A | C | C | C | T | A | T | A | G | A |
| NC025276.1 | Coleoptera | *Aquatica leii* | A | G | A | C | G | A | G | A | A | G | A | C | C | C | T | A | T | A | G | A |
| KX758086.1 | Coleoptera | *Aquatica wuhana* | A | G | A | C | G | A | G | A | A | G | A | C | C | C | T | A | T | A | G | A |
| NC035061.1 | Coleoptera | *Aquatica wuhana* | A | G | A | C | G | A | G | A | A | G | A | C | C | C | T | A | T | A | G | A |
| KX229747.1 | Coleoptera | *Asymmetricata circumdata* | A | G | A | C | G | A | G | A | A | G | A | C | C | C | T | A | T | A | G | A |
| MK292113.1 | Coleoptera | *Asymmetricata circumdata* | A | G | A | C | G | A | G | A | A | G | A | C | C | C | C | A | T | A | G | A |
| NC032062.1 | Coleoptera | *Asymmetricata circumdata* | A | G | A | C | G | A | G | A | A | G | A | C | C | C | T | A | T | A | G | A |
| KJ922151.1 | Coleoptera | *Bicellonychia lividipennis* | A | G | A | C | G | A | G | A | A | G | A | C | C | C | T | A | T | A | G | A |
| NC030060.1 | Coleoptera | *Bicellonychia lividipennis* | A | G | A | C | G | A | G | A | A | G | A | C | C | C | T | A | T | A | G | A |
| KT876885.1 | Coleoptera | *Colymbetes fuscus* | A | G | A | C | G | A | G | A | A | G | A | C | C | C | T | A | T | A | G | A |
| MK292114.1 | Coleoptera | *Curtos bilineatus* | A | G | A | C | G | A | G | A | A | G | A | C | C | C | T | A | T | A | G | A |
| NC044789.1 | Coleoptera | *Curtos bilineatus* | A | G | A | C | G | A | G | A | A | G | A | C | C | C | T | A | T | A | G | A |
| MK609965.1 | Coleoptera | *Curtos costipennis* | A | G | A | C | G | A | G | A | A | G | A | C | C | C | T | A | T | A | G | A |
| MH651351.1 | Coleoptera | *Diaphanes citrinus* | A | G | A | C | G | A | G | A | A | G | A | C | C | C | C | A | T | A | G | A |

| | | | | | | | | | | | | | | | | | | | | | | |
|---|---|---|---|---|---|---|---|---|---|---|---|---|---|---|---|---|---|---|---|---|---|---|
| MK292116.1 | Coleoptera | *Diaphanes mendax* | A | G | A | C | G | A | G | A | A | G | A | C | C | C | C | A | T | A | G | A |
| NC044791.1 | Coleoptera | *Diaphanes mendax* | A | G | A | C | G | A | G | A | A | G | A | C | C | C | C | A | T | A | G | A |
| MK292094.1 | Coleoptera | *Diaphanes nubilus* | A | G | A | C | G | A | G | A | A | G | A | C | C | C | C | A | T | A | G | A |
| NC044787.1 | Coleoptera | *Diaphanes nubilus* | A | G | A | C | G | A | G | A | A | G | A | C | C | C | C | A | T | A | G | A |
| MK292118.1 | Coleoptera | *Diaphanes pectinealis* | A | G | A | C | G | A | G | A | A | G | A | C | C | C | C | G | T | A | G | A |
| NC044793.1 | Coleoptera | *Diaphanes pectinealis* | A | G | A | C | G | A | G | A | A | G | A | C | C | C | C | G | T | A | G | A |
| MK292095.1 | Coleoptera | *Diaphanes* sp. | A | G | A | C | G | A | G | A | A | G | A | C | C | C | C | A | T | A | G | A |
| MG242622.1 | Coleoptera | *Ellychnia corrusca* | A | G | A | C | G | A | G | A | A | G | A | C | C | C | C | A | T | A | G | A |
| MK292112.1 | Coleoptera | *Emeia pseudosauteri* | A | G | A | C | G | A | G | A | A | G | A | C | C | C | T | A | T | A | G | A |
| NC044778.1 | Coleoptera | *Emeia pseudosauteri* | A | G | A | C | G | A | G | A | A | G | A | C | C | C | T | A | T | A | G | A |
| KT876897.1 | Coleoptera | *Hydroporus erythrocephalus* | A | G | A | C | G | A | G | A | A | G | A | C | C | C | T | A | T | A | G | A |
| KT876896.1 | Coleoptera | *Hydroporus obscurus* | A | G | A | C | G | A | G | A | A | G | A | C | C | C | T | A | T | A | G | A |
| KT876901.1 | Coleoptera | *Hygrotus confluens* | A | G | A | C | G | A | G | A | A | G | A | C | C | C | T | A | T | A | G | A |

| | | | | | | | | | | | | | | | | | | | | | | |
|---|---|---|---|---|---|---|---|---|---|---|---|---|---|---|---|---|---|---|---|---|---|---|
| KT876900.1 | Coleoptera | *Hygrotus impressopunctatus* | A | G | A | C | G | A | G | A | A | G | A | C | C | C | T | A | T | A | G | A |
| KT876899.1 | Coleoptera | *Hygrotus inaequalis* | A | G | A | C | G | A | G | A | A | G | A | C | C | C | T | A | T | A | G | A |
| KX087299.1 | Coleoptera | *Hygrotus nigrolineatus* | A | G | A | C | G | A | G | A | A | G | A | C | C | C | T | A | T | A | G | A |
| MH427718.1 | Coleoptera | *Inflata indica* | G | G | A | C | G | A | G | A | A | G | A | C | C | C | T | A | T | A | G | A |
| NC039700.1 | Coleoptera | *Inflata indica* | G | G | A | C | G | A | G | A | A | G | A | C | C | C | T | A | T | A | G | A |
| MK292091.1 | Coleoptera | *Lamprigera yunnana* | A | G | A | C | G | A | G | A | A | G | A | C | C | C | T | A | T | A | G | A |
| NC044786.1 | Coleoptera | *Lamprigera yunnana* | A | G | A | C | G | A | G | A | A | G | A | C | C | C | T | A | T | A | G | A |
| KX087302.1 | Coleoptera | *Lampyris noctiluca* | A | G | A | C | G | A | G | A | A | G | A | C | C | C | C | A | T | A | G | A |
| MG912994.1 | Coleoptera | *Limbodessus palmulaoides* | A | G | A | C | G | A | G | A | A | G | A | C | C | C | T | A | T | A | G | A |
| NC037749.1 | Coleoptera | *Limbodessus palmulaoides* | A | G | A | C | G | A | G | A | A | G | A | C | C | C | T | A | T | A | G | A |
| JQ034416.1 | Coleoptera | *Limnichidae* sp. | A | G | A | C | G | A | G | A | A | G | A | C | C | C | T | A | T | A | G | A |
| KT876902.1 | Coleoptera | *Liopterus haemorrhoidalis* | A | G | A | C | G | A | G | A | A | G | A | C | C | C | T | A | T | A | G | A |
| NC022472.1 | Coleoptera | *Luciola cruciata* | A | G | A | C | G | A | G | A | A | G | A | C | C | C | T | A | T | A | G | A |
| MG770613.1 | Coleoptera | *Luciola curtithorax* | A | G | A | C | G | A | G | A | A | G | A | C | C | C | T | A | T | A | G | A |

| | | | | | | | | | | | | | | | | | | | | | | |
|---|---|---|---|---|---|---|---|---|---|---|---|---|---|---|---|---|---|---|---|---|---|---|
| NC038225.1 | Coleoptera | *Luciola curtithorax* | A | G | A | C | G | A | G | A | A | G | A | C | C | C | T | A | T | A | G | A |
| KP313820.1 | Coleoptera | *Luciola substriata* | A | G | A | C | G | A | G | A | A | G | A | C | C | C | T | A | T | A | G | A |
| NC027176.1 | Coleoptera | *Luciola substriata* | A | G | A | C | G | A | G | A | A | G | A | C | C | C | T | A | T | A | G | A |
| FJ859901.1 | Coleoptera | *Macrogyrus oblongus* | A | G | A | C | G | A | G | A | A | G | A | C | C | C | T | A | T | A | G | A |
| NC013249.1 | Coleoptera | *Macrogyrus oblongus* | A | G | A | C | G | A | G | A | A | G | A | C | C | C | T | A | T | A | G | A |
| MG912995.1 | Coleoptera | *Paroster macrosturtensis* | A | G | A | C | G | A | G | A | A | G | A | C | C | C | T | A | T | A | G | A |
| NC037750.1 | Coleoptera | *Paroster macrosturtensis* | A | G | A | C | G | A | G | A | A | G | A | C | C | C | T | A | T | A | G | A |
| MG912996.1 | Coleoptera | *Paroster mesosturtensis* | A | G | A | C | G | A | G | A | A | G | A | C | C | C | T | A | T | A | G | A |
| NC037751.1 | Coleoptera | *Paroster mesosturtensis* | A | G | A | C | G | A | G | A | A | G | A | C | C | C | T | A | T | A | G | A |
| MG912997.1 | Coleoptera | *Paroster microsturtensis* | A | G | A | C | G | A | G | A | A | G | A | C | C | C | T | A | T | A | G | A |
| NC037752.1 | Coleoptera | *Paroster microsturtensis* | A | G | A | C | G | A | G | A | A | G | A | C | C | C | T | A | T | A | G | A |
| KY778696.1 | Coleoptera | *Photinus pyralis* | A | G | A | C | G | A | G | A | A | G | A | C | C | C | C | A | T | A | G | A |
| MF686051.1 | Coleoptera | *Pteroptyx maipo* | G | G | A | C | G | A | G | A | A | G | A | C | C | C | T | A | T | A | G | A |
| NC036353.1 | Coleoptera | *Pteroptyx maipo* | G | G | A | C | G | A | G | A | A | G | A | C | C | C | T | A | T | A | G | A |

| | | | | | | | | | | | | | | | | | | | | | | |
|---|---|---|---|---|---|---|---|---|---|---|---|---|---|---|---|---|---|---|---|---|---|---|
| MK292115.1 | Coleoptera | *Pyrocoelia praetexta* | A | G | A | C | G | A | G | A | A | G | A | C | C | C | C | A | T | A | G | A |
| NC044790.1 | Coleoptera | *Pyrocoelia praetexta* | A | G | A | C | G | A | G | A | A | G | A | C | C | C | C | A | T | A | G | A |
| AF452048.1 | Coleoptera | *Pyrocoelia rufa* | A | G | A | C | G | A | G | A | A | G | A | C | C | C | C | A | T | A | G | A |
| NC003970.1 | Coleoptera | *Pyrocoelia rufa* | A | G | A | C | G | A | G | A | A | G | A | C | C | C | C | A | T | A | G | A |
| MK292117.1 | Coleoptera | *Pyrocoelia thibetana* | A | G | A | C | G | A | G | A | A | G | A | C | C | C | C | A | T | A | G | A |
| NC044792.1 | Coleoptera | *Pyrocoelia thibetana* | A | G | A | C | G | A | G | A | A | G | A | C | C | C | C | A | T | A | G | A |
| MK292111.1 | Coleoptera | *Vesta saturnalis* | A | G | A | C | G | A | G | A | A | G | A | C | C | C | T | A | T | A | G | A |
| NC044788.1 | Coleoptera | *Vesta saturnalis* | A | G | A | C | G | A | G | A | A | G | A | C | C | C | T | A | T | A | G | A |
| MF680449.1 | Trichoptera | *Anabolia bimaculata* | A | G | A | C | G | A | G | A | A | G | A | C | C | C | T | A | T | A | A | A |
| NC036004.1 | Trichoptera | *Anabolia bimaculata* | A | G | A | C | G | A | G | A | A | G | A | C | C | C | T | A | T | A | A | A |
| KF756944.1 | Trichoptera | *Apatania* sp. | A | G | A | C | G | A | G | A | A | G | A | C | C | C | T | A | T | A | A | A |
| MT410856.1 | Trichoptera | *Beraea pullata* | A | G | A | C | G | A | G | A | A | G | A | C | C | C | T | A | T | A | G | A |
| MG669126.1 | Trichoptera | *Cheumatopsyche analis* | A | G | A | C | G | A | G | A | A | G | A | C | C | C | T | T | T | A | G | A |
| NC036955.1 | Trichoptera | *Cheumatopsyche analis* | A | G | A | C | G | A | G | A | A | G | A | C | C | C | T | T | T | A | G | A |

| | | | | | | | | | | | | | | | | | | | | | | |
|---|---|---|---|---|---|---|---|---|---|---|---|---|---|---|---|---|---|---|---|---|---|---|
| KX385010.1 | Trichoptera | *Cheumatopsyche brevilineata* | A | G | A | C | G | A | G | A | A | G | A | C | C | C | T | T | T | A | G | A |
| NC043769.1 | Trichoptera | *Cheumatopsyche brevilineata* | A | G | A | C | G | A | G | A | A | G | A | C | C | C | T | T | T | A | G | A |
| MG669125.1 | Trichoptera | *Cheumatopsyche campyla* | A | G | A | C | G | A | G | A | A | G | A | C | C | C | T | T | T | A | G | A |
| NC036954.1 | Trichoptera | *Cheumatopsyche campyla* | A | G | A | C | G | A | G | A | A | G | A | C | C | C | T | T | T | A | G | A |
| MG669123.1 | Trichoptera | *Cheumatopsyche speciosa* | A | G | A | C | G | A | G | A | A | G | A | C | C | C | T | T | T | A | G | A |
| NC036952.1 | Trichoptera | *Cheumatopsyche speciosa* | A | G | A | C | G | A | G | A | A | G | A | C | C | C | T | T | T | A | G | A |
| KF717094.1 | Trichoptera | *Eubasilissa regina* | A | G | A | C | G | A | G | A | A | G | A | C | C | C | T | A | T | A | A | A |
| KF756943.1 | Trichoptera | *Eubasilissa regina* | A | G | A | C | G | A | G | A | A | G | A | C | C | C | T | A | T | A | A | A |
| NC023374.1 | Trichoptera | *Eubasilissa regina* | A | G | A | C | G | A | G | A | A | G | A | C | C | C | T | A | T | A | A | A |
| KX385011.1 | Trichoptera | *Hydatophylax nigrovittatus* | A | G | A | C | G | A | G | A | A | G | A | C | C | C | T | A | T | A | A | A |
| NC043770.1 | Trichoptera | *Hydatophylax nigrovittatus* | A | G | A | C | G | A | G | A | A | G | A | C | C | C | T | A | T | A | A | A |
| KF717095.1 | Trichoptera | *Hydromanicus wulaianus* | A | G | A | C | G | A | G | A | A | G | A | C | C | C | T | T | T | A | G | A |
| NC036156.1 | Trichoptera | *Hydromanicus* | A | G | A | C | G | A | G | A | A | G | A | C | C | C | T | T | T | A | G | A |

| | | | | | | | | | | | | | | | | | | | | | | |
|---|---|---|---|---|---|---|---|---|---|---|---|---|---|---|---|---|---|---|---|---|---|---|
| | | *wulaianus* | | | | | | | | | | | | | | | | | | | | |
| MT628580.1 | Trichoptera | *Hydropsyche angustipennis* | A | G | A | C | G | A | G | A | A | G | A | C | C | C | T | T | T | A | G | A |
| MG669122.1 | Trichoptera | *Hydropsyche orris* | A | G | A | C | G | A | G | A | A | G | A | C | C | C | T | T | T | A | G | A |
| NC036951.1 | Trichoptera | *Hydropsyche orris* | A | G | A | C | G | A | G | A | A | G | A | C | C | C | T | T | T | A | G | A |
| KT876876.1 | Trichoptera | *Hydropsyche pellucidula* | A | G | A | C | G | A | G | A | A | G | A | C | C | C | T | T | T | A | G | A |
| NC029246.1 | Trichoptera | *Hydropsyche pellucidula* | A | G | A | C | G | A | G | A | A | G | A | C | C | C | T | T | T | A | G | A |
| MG669121.1 | Trichoptera | *Hydropsyche simulans* | A | G | A | C | G | A | G | A | A | G | A | C | C | C | T | T | T | A | G | A |
| NC036950.1 | Trichoptera | *Hydropsyche simulans* | A | G | A | C | G | A | G | A | A | G | A | C | C | C | T | T | T | A | G | A |
| MK077681.1 | Trichoptera | *Limnephilus hyalinus* | A | G | A | C | G | A | G | A | A | G | A | C | C | C | T | A | T | A | A | A |
| NC044710.1 | Trichoptera | *Limnephilus hyalinus* | A | G | A | C | G | A | G | A | A | G | A | C | C | C | T | A | T | A | A | A |
| MG980616.1 | Trichoptera | *Phryganea cinerea* | A | G | A | C | G | A | G | A | A | G | A | C | C | C | T | A | T | A | A | A |
| NC039714.1 | Trichoptera | *Phryganea cinerea* | A | G | A | C | G | A | G | A | A | G | A | C | C | C | T | A | T | A | A | A |
| KX385012.1 | Trichoptera | *Phryganopsyche* | A | G | A | C | G | A | G | A | A | G | A | C | C | C | C | G | T | A | G | A |

| | | | | | | | | | | | | | | | | | | | | | | |
|---|---|---|---|---|---|---|---|---|---|---|---|---|---|---|---|---|---|---|---|---|---|---|
| | | *latipennis* | | | | | | | | | | | | | | | | | | | | |
| NC043771.1 | Trichoptera | *Phryganopsyche latipennis* | A | G | A | C | G | A | G | A | A | G | A | C | C | C | C | G | T | A | G | A |
| MG669124.1 | Trichoptera | *Potamyia flava* | A | G | A | C | G | A | G | A | A | G | A | C | C | C | T | T | T | A | G | A |
| NC036953.1 | Trichoptera | *Potamyia flava* | A | G | A | C | G | A | G | A | A | G | A | C | C | C | T | T | T | A | G | A |
| KP455290.1 | Trichoptera | *Sericostoma personatum* | A | G | A | C | G | A | G | A | A | G | A | C | C | C | T | A | T | A | A | A |
| KP455291.1 | Trichoptera | *Thremma gallicum* | A | G | A | C | G | A | G | A | A | G | A | C | C | C | T | A | T | A | A | A |
| AY072044.1 | Diptera | *Aedes albopictus* | A | G | A | C | G | A | G | A | A | G | A | C | C | C | T | A | T | A | G | A |
| NC030249.1 | Diptera | *Anopheles bellator* | A | G | A | C | G | A | G | A | A | G | A | C | C | C | T | A | T | A | G | A |
| NC030717.1 | Diptera | *Anopheles janconnae* | G | G | A | C | G | A | G | A | A | G | A | C | C | C | T | A | T | A | G | A |
| HQ335345.1 | Diptera | *Anopheles oryzalimnetes* | G | G | A | C | G | A | G | A | A | G | A | C | C | C | T | A | T | A | G | A |
| MF381722.1 | Diptera | *Culex bilineatus* | G | G | A | C | G | A | G | A | A | G | A | C | C | C | T | A | T | A | G | A |
| MF381718.1 | Diptera | *Culex lygrus* | G | G | A | C | G | A | G | A | A | G | A | C | C | C | T | A | T | A | G | A |
| MH316119.1 | Diptera | *Lutzia halifaxii* | G | G | A | C | G | A | G | A | A | G | A | C | C | C | T | A | T | A | G | A |
| MN626442.1 | Diptera | *Ochlerotatus taeniorhynchus* | G | G | A | C | G | A | G | A | A | G | A | C | C | C | T | A | T | A | G | A |
| KP721463.1 | Diptera | *Ochlerotatus vigilax* | G | G | A | C | G | A | G | A | A | G | A | C | C | C | T | A | T | A | G | A |
| MF957171.1 | Diptera | *Sabethes* | A | G | A | C | G | A | G | A | A | G | A | C | C | C | T | A | T | A | G | A |

| | | | | | | | | | | | | | | | | | | | | | | |
|---|---|---|---|---|---|---|---|---|---|---|---|---|---|---|---|---|---|---|---|---|---|---|
| | | *belisarioi* | | | | | | | | | | | | | | | | | | | | |
| KP793690.1 | Diptera | *Simulium aureohirtum* | A | G | A | C | G | A | G | A | A | G | A | C | C | C | T | A | T | A | G | A |
| NC029753.1 | Diptera | *Simulium aureohirtum* | A | G | A | C | G | A | G | A | A | G | A | C | C | C | T | A | T | A | G | A |
| MH729190.1 | Diptera | *Simulium maculatum* | A | G | A | C | G | A | G | A | A | G | A | C | C | C | T | A | T | A | G | A |
| NC040120.1 | Diptera | *Simulium maculatum* | A | G | A | C | G | A | G | A | A | G | A | C | C | C | T | A | T | A | G | A |
| KU252587.1 | Diptera | *Simulium variegatum* | A | G | A | C | G | A | G | A | A | G | A | C | C | C | T | A | T | A | G | A |
| NC033348.1 | Diptera | *Simulium variegatum* | A | G | A | C | G | A | G | A | A | G | A | C | C | C | T | A | T | A | G | A |
| JN861743.1 | Diptera | *Tipula abdominalis* | G | G | A | C | G | A | G | A | A | G | A | C | C | C | C | A | T | A | G | A |
| KT970065.1 | Diptera | *Tipula cockerelliana* | G | G | A | C | G | A | G | A | A | G | A | C | C | C | C | A | T | A | G | A |
| NC030520.1 | Diptera | *Tipula cockerelliana* | G | G | A | C | G | A | G | A | A | G | A | C | C | C | C | A | T | A | G | A |
| MK575491.1 | Diptera | *Uranotaenia geometrica* | A | G | A | C | G | A | G | A | A | G | A | C | C | C | T | A | T | A | G | A |
| | | A | 35.9 | 0.0 | 100.0 | 0.0 | 2.0 | 100.0 | 0.5 | 100.0 | 100.0 | 0.0 | 100.0 | 0.0 | 0.0 | 0.0 | 0.0 | 93.4 | 0.0 | 100.0 | 3.2 | 100.0 |

| | | | | | | | | | | | | | | | | | | | | |
|---|---|---|---|---|---|---|---|---|---|---|---|---|---|---|---|---|---|---|---|---|
| T | 0.4 | 0.0 | 0.0 | 0.0 | 0.0 | 0.0 | 7.5 | 0.0 | 0.0 | 0.0 | 0.0 | 0.0 | 0.0 | 0.0 | 94.8 | 3.9 | 100.0 | 0.0 | 0.0 | 0.0 |
| G | 63.7 | 100.0 | 0.0 | 0.0 | 98.0 | 0.0 | 91.9 | 0.0 | 0.0 | 100.0 | 0.0 | 0.0 | 0.0 | 0.0 | 0.0 | 2.7 | 0.0 | 0.0 | 96.8 | 0.0 |
| C | 0.0 | 0.0 | 0.0 | 100.0 | 0.0 | 0.0 | 0.0 | 0.0 | 0.0 | 0.0 | 0.0 | 100.0 | 100.0 | 100.0 | 5.2 | 0.0 | 0.0 | 0.0 | 0.0 | 0.0 |

Table S3. Nucleotide sequences of universal primer (MtInsects-16S_R) and homologous region of 557 sequences of insects used for development in this study.

| | | | | | | | | | | | | | | | | | | | | | | |
|---|---|---|---|---|---|---|---|---|---|---|---|---|---|---|---|---|---|---|---|---|---|---|
| MtInsects-16S_R | | | A | T | C | C | A | A | C | A | T | C | G | A | G | G | T | C | G | C | A | A |
| | | A | 536 | 0 | 0 | 0 | 557 | 557 | 0 | 557 | 0 | 0 | 0 | 555 | 0 | 0 | 0 | 0 | 13 | 0 | 557 | 555 |
| | | T | 0 | 553 | 181 | 0 | 0 | 0 | 0 | 0 | 557 | 0 | 0 | 0 | 0 | 0 | 557 | 11 | 0 | 6 | 0 | 0 |
| | | G | 21 | 0 | 0 | 0 | 0 | 0 | 0 | 0 | 0 | 0 | 557 | 2 | 557 | 557 | 0 | 0 | 544 | 0 | 0 | 0 |
| | | C | 0 | 4 | 376 | 557 | 0 | 0 | 557 | 0 | 0 | 557 | 0 | 0 | 0 | 0 | 0 | 546 | 0 | 551 | 0 | 2 |
| GenBank Acc. No. | Order | Species | | | | | | | | | | | | | | | | | | | | |
| MN481418.1 | Diplura | *Campodea augens* | A | T | T | C | A | A | C | A | T | C | G | A | G | G | T | T | A | T | A | A |
| DQ529236.1 | Diplura | *Campodea fragilis* | A | T | T | C | A | A | C | A | T | C | G | A | G | G | T | T | A | T | A | A |
| NC008233.1 | Diplura | *Campodea fragilis* | A | T | T | C | A | A | C | A | T | C | G | A | G | G | T | T | A | T | A | A |
| DQ529237.1 | Diplura | *Campodea lubbocki* | A | T | T | C | A | A | C | A | T | C | G | A | G | G | T | T | A | T | A | A |
| NC008234.1 | Diplura | *Campodea lubbocki* | A | T | T | C | A | A | C | A | T | C | G | A | G | G | T | T | A | T | A | A |
| AY771989.1 | Diplura | *Japyx solifugus* | A | C | C | C | A | A | C | A | T | C | G | A | G | G | T | C | G | C | A | A |
| NC007214.1 | Diplura | *Japyx solifugus* | A | C | C | C | A | A | C | A | T | C | G | A | G | G | T | C | G | C | A | A |
| JN990601.1 | Diplura | *Lepidocampa weberi* | A | T | T | C | A | A | C | A | T | C | G | A | G | G | T | T | G | C | A | A |
| NC022675.1 | Diplura | *Lepidocampa weberi* | A | T | T | C | A | A | C | A | T | C | G | A | G | G | T | T | G | C | A | A |
| JN990600.1 | Diplura | *Occasjapyx japonicus* | A | C | C | C | A | A | C | A | T | C | G | A | G | G | T | C | G | C | A | A |
| NC022674.1 | Diplura | *Occasjapyx* | A | C | C | C | A | A | C | A | T | C | G | A | G | G | T | C | G | C | A | A |

| | | | | | | | | | | | | | | | | | | | | | | |
|---|---|---|---|---|---|---|---|---|---|---|---|---|---|---|---|---|---|---|---|---|---|---|
| | | *japonicus* | | | | | | | | | | | | | | | | | | | | |
| JN990598.1 | Diplura | *Octostigma sinensis* | A | T | T | C | A | A | C | A | T | C | G | A | G | G | T | C | G | C | A | A |
| NC022672.1 | Diplura | *Octostigma sinensis* | A | T | T | C | A | A | C | A | T | C | G | A | G | G | T | C | G | C | A | A |
| JN990599.1 | Diplura | *Parajapyx emeryanus* | A | T | C | C | A | A | C | A | T | C | G | A | G | G | T | C | G | C | A | A |
| NC022673.1 | Diplura | *Parajapyx emeryanus* | A | T | C | C | A | A | C | A | T | C | G | A | G | G | T | C | G | C | A | A |
| KJ754500.1 | Archaeognatha | *Allopsontus* sp. | A | T | C | C | A | A | C | A | T | C | G | A | G | G | T | C | G | C | A | A |
| KJ754501.1 | Archaeognatha | *Allopsontus* sp. | A | T | C | C | A | A | C | A | T | C | G | A | G | G | T | C | G | C | A | A |
| AY793551.1 | Archaeognatha | *Nesomachilis australica* | A | T | C | C | A | A | C | A | T | C | G | A | G | G | T | C | G | C | A | A |
| NC006895.1 | Archaeognatha | *Nesomachilis australica* | A | T | C | C | A | A | C | A | T | C | G | A | G | G | T | C | G | C | A | A |
| KJ754502.1 | Archaeognatha | *Pedetontinus luanchuanensis* | A | T | C | C | A | A | C | A | T | C | G | A | G | G | T | C | G | C | A | A |
| EU621793.1 | Archaeognatha | *Pedetontus silvestrii* | A | T | C | C | A | A | C | A | T | C | G | A | G | G | T | C | G | C | A | A |
| NC011717.1 | Archaeognatha | *Pedetontus silvestrii* | A | T | C | C | A | A | C | A | T | C | G | A | G | G | T | C | G | C | A | A |
| KJ754503.1 | Archaeognatha | *Petrobiellus* sp. | A | T | C | C | A | A | C | A | T | C | G | A | G | G | T | C | G | C | A | A |
| KJ754504.1 | Archaeognatha | *Petrobiellus* sp. | A | T | C | C | A | A | C | A | T | C | G | A | G | G | T | C | G | C | A | A |
| AY956355.1 | Archaeognatha | *Petrobius brevistylis* | A | T | C | C | A | A | C | A | T | C | G | A | G | G | T | T | G | C | A | A |
| NC007688.1 | Archaeognatha | *Petrobius brevistylis* | A | T | C | C | A | A | C | A | T | C | G | A | G | G | T | T | G | C | A | A |
| JX308221.1 | Archaeognatha | *Songmachilis* | A | T | C | C | A | A | C | A | T | C | G | A | G | G | T | C | G | C | A | A |

| | | | | | | | | | | | | | | | | | | | | | | |
|---|---|---|---|---|---|---|---|---|---|---|---|---|---|---|---|---|---|---|---|---|---|---|
| | | *xinxiangensis* | | | | | | | | | | | | | | | | | | | | |
| NC021384.1 | Archaeognatha | *Songmachilis xinxiangensis* | A | T | C | C | A | A | C | A | T | C | G | A | G | G | T | C | G | C | A | A |
| EU016193.1 | Archaeognatha | *Trigoniophthalmus alternatus* | A | T | C | C | A | A | C | A | T | C | G | A | G | G | T | C | G | C | A | A |
| NC010532.1 | Archaeognatha | *Trigoniophthalmus alternatus* | A | T | C | C | A | A | C | A | T | C | G | A | G | G | T | C | G | C | A | A |
| EU084035.1 | Zygentoma | *Atelura formicaria* | A | T | T | C | A | A | C | A | T | C | G | A | G | G | T | C | G | C | A | A |
| NC011197.1 | Zygentoma | *Atelura formicaria* | A | T | T | C | A | A | C | A | T | C | G | A | G | G | T | C | G | C | A | A |
| MK301436.1 | Zygentoma | *Ctenolepisma villosa* | A | T | C | C | A | A | C | A | T | C | G | A | G | G | T | C | G | C | A | A |
| NC046478.1 | Zygentoma | *Ctenolepisma villosa* | A | T | C | C | A | A | C | A | T | C | G | A | G | G | T | C | G | C | A | A |
| MT108230.1 | Zygentoma | *Lepisma saccharina* | A | T | C | C | A | A | C | A | T | C | G | A | G | G | T | C | G | C | A | A |
| NC047445.1 | Zygentoma | *Lepisma saccharina* | A | T | C | C | A | A | C | A | T | C | G | A | G | G | T | C | G | C | A | A |
| AY639935.1 | Zygentoma | *Thermobia domestica* | A | T | C | C | A | A | C | A | T | C | G | A | G | G | T | C | G | C | A | A |
| NC006080.1 | Zygentoma | *Thermobia domestica* | A | T | C | C | A | A | C | A | T | C | G | A | G | G | T | C | G | C | A | A |
| MK951657.1 | Zygentoma | *Thermobia* sp. | A | T | C | C | A | A | C | A | T | C | G | A | G | G | T | C | G | C | A | A |
| AY191994.1 | Zygentoma | *Tricholepidion gertschi* | A | T | C | C | A | A | C | A | T | C | G | A | G | G | T | C | G | C | A | A |

| | | | | | | | | | | | | | | | | | | | | | | |
|---|---|---|---|---|---|---|---|---|---|---|---|---|---|---|---|---|---|---|---|---|---|---|
| NC005437.1 | Zygentoma | *Tricholepidion gertschi* | A | T | C | C | A | A | C | A | T | C | G | A | G | G | T | C | G | C | A | A |
| GU479735.1 | Ephemeroptera | *Alainites yixiani* | A | T | C | C | A | A | C | A | T | C | G | A | G | G | T | C | G | C | A | A |
| GU936204.1 | Ephemeroptera | *Baetis sp.* | A | T | C | C | A | A | C | A | T | C | G | A | G | G | T | C | G | C | A | A |
| MT628575.1 | Ephemeroptera | *Caenis robusta* | G | T | C | C | A | A | C | A | T | C | G | A | G | G | T | C | G | C | A | A |
| GQ502451.1 | Ephemeroptera | *Caenis* sp. | G | T | C | C | A | A | C | A | T | C | G | A | G | G | T | C | G | C | A | A |
| MG910499.1 | Ephemeroptera | *Caenis* sp. | G | T | C | C | A | A | C | A | T | C | G | A | G | G | T | C | G | C | A | A |
| MN356096.1 | Ephemeroptera | *Caenis* sp. | G | T | C | C | A | A | C | A | T | C | G | A | G | G | T | C | G | C | A | A |
| MN807287.1 | Ephemeroptera | *Choroterpides apiculata* | A | T | C | C | A | A | C | A | T | C | G | A | G | G | T | C | A | C | A | A |
| MG870104.1 | Ephemeroptera | *Epeorus herklotsi* | G | T | C | C | A | A | C | A | T | C | G | A | G | G | T | C | G | C | A | A |
| MH752075.1 | Ephemeroptera | *Epeorus herklotsi* | G | T | C | C | A | A | C | A | T | C | G | A | G | G | T | C | G | C | A | A |
| NC039612.1 | Ephemeroptera | *Epeorus herklotsi* | G | T | C | C | A | A | C | A | T | C | G | A | G | G | T | C | G | C | A | A |
| KJ493406.1 | Ephemeroptera | *Epeorus* sp. | G | T | C | C | A | A | C | A | T | C | G | A | G | G | T | C | G | C | A | A |
| KM244708.1 | Ephemeroptera | *Epeorus* sp. | G | T | C | C | A | A | C | A | T | C | G | A | G | G | T | C | G | C | A | A |
| MK951674.1 | Ephemeroptera | *Epeorus* sp. | G | T | C | C | A | A | C | A | T | C | G | A | G | G | T | C | G | C | A | A |
| EU591678.1 | Ephemeroptera | *Ephemera orientalis* | G | T | C | C | A | A | C | A | T | C | G | A | G | G | T | C | G | C | A | A |
| NC012645.1 | Ephemeroptera | *Ephemera orientalis* | G | T | C | C | A | A | C | A | T | C | G | A | G | G | T | C | G | C | A | A |
| MK951659.1 | Ephemeroptera | *Ephemera* sp. | G | T | C | C | A | A | C | A | T | C | G | A | G | G | T | C | G | C | A | A |
| KM244691.1 | Ephemeroptera | *Ephemerella* sp. | A | T | C | C | A | A | C | A | T | C | G | A | G | G | T | C | G | C | A | A |
| MT274127.1 | Ephemeroptera | *Ephemerella* sp. | A | T | C | C | A | A | C | A | T | C | G | A | G | G | T | C | G | C | A | A |
| NC050279.1 | Ephemeroptera | *Ephemerella* sp. | A | T | C | C | A | A | C | A | T | C | G | A | G | G | T | C | G | C | A | A |

| | | | | | | | | | | | | | | | | | | | | | | |
|---|---|---|---|---|---|---|---|---|---|---|---|---|---|---|---|---|---|---|---|---|---|---|
| GU936203.1 | Ephemeroptera | *Habrophlebiodes zijinensis* | A | T | C | C | A | A | C | A | T | C | G | A | G | G | T | C | A | C | A | A |
| HM143892.1 | Ephemeroptera | *Isonychia ignota* | A | T | C | C | A | A | C | A | T | C | G | A | G | G | T | C | G | C | A | A |
| MH119135.1 | Ephemeroptera | *Isonychia kiangsinensis* | A | T | C | C | A | A | C | A | T | C | G | A | G | G | T | C | G | C | A | A |
| NC042163.1 | Ephemeroptera | *Isonychia kiangsinensis* | A | T | C | C | A | A | C | A | T | C | G | A | G | G | T | C | G | C | A | A |
| MK951658.1 | Ephemeroptera | *Isonychia* sp. | A | T | C | C | A | A | C | A | T | C | G | A | G | G | T | C | G | C | A | A |
| HM004123.1 | Ephemeroptera | *Paegniodes cupulatus* | G | T | C | C | A | A | C | A | T | C | G | A | G | G | T | C | G | C | A | A |
| MK951673.1 | Ephemeroptera | *Parafronurus* sp. | G | T | C | C | A | A | C | A | T | C | G | A | G | G | T | C | G | C | A | A |
| EU349015.1 | Ephemeroptera | *Parafronurus youi* | G | T | C | C | A | A | C | A | T | C | G | A | G | G | T | C | G | C | A | A |
| NC011359.1 | Ephemeroptera | *Parafronurus youi* | G | T | C | C | A | A | C | A | T | C | G | A | G | G | T | C | G | C | A | A |
| KM244674.1 | Ephemeroptera | *Potamanthus* sp. | G | T | C | C | A | A | C | A | T | C | G | A | G | G | T | C | G | C | A | A |
| MK951670.1 | Ephemeroptera | *Rhithrogena* sp. | G | T | C | C | A | A | C | A | T | C | G | A | G | G | T | C | G | C | A | A |
| MT274128.1 | Ephemeroptera | *Serratella* sp. | A | T | C | C | A | A | C | A | T | C | G | A | G | G | T | C | G | C | A | A |
| MT274129.1 | Ephemeroptera | *Serratella* sp. | A | T | C | C | A | A | C | A | T | C | G | A | G | G | T | C | G | C | A | A |
| NC050280.1 | Ephemeroptera | *Serratella* sp. | A | T | C | C | A | A | C | A | T | C | G | A | G | G | T | C | G | C | A | A |
| NC050281.1 | Ephemeroptera | *Serratella* sp. | A | T | C | C | A | A | C | A | T | C | G | A | G | G | T | C | G | C | A | A |
| MT274130.1 | Ephemeroptera | *Serratella zapekinae* | A | T | C | C | A | A | C | A | T | C | G | A | G | G | T | C | G | C | A | A |
| NC050282.1 | Ephemeroptera | *Serratella zapekinae* | A | T | C | C | A | A | C | A | T | C | G | A | G | G | T | C | G | C | A | A |
| FJ606783.1 | Ephemeroptera | *Siphlonurus* | G | T | C | C | A | A | C | A | T | C | G | A | G | G | T | C | G | C | A | A |

| | | | | | | | | | | | | | | | | | | | | | | |
|---|---|---|---|---|---|---|---|---|---|---|---|---|---|---|---|---|---|---|---|---|---|---|
| | | *immanis* | | | | | | | | | | | | | | | | | | | | |
| NC013822.1 | Ephemeroptera | *Siphlonurus immanis* | G | T | C | C | A | A | C | A | T | C | G | A | G | G | T | C | G | C | A | A |
| KM244684.1 | Ephemeroptera | *Siphlonurus* sp. | A | T | C | C | A | A | C | A | T | C | G | A | G | G | T | C | G | C | A | A |
| HQ875717.1 | Ephemeroptera | *Siphluriscus chinensis* | A | T | C | C | A | A | C | A | T | C | G | A | G | G | T | C | G | C | A | A |
| MT274131.1 | Ephemeroptera | *Torleya grandiforceps* | A | T | C | C | A | A | C | A | T | C | G | A | G | G | T | C | A | C | A | A |
| NC050283.1 | Ephemeroptera | *Torleya grandiforceps* | A | T | C | C | A | A | C | A | T | C | G | A | G | G | T | C | A | C | A | A |
| MT274132.1 | Ephemeroptera | *Torleya nepalica* | A | T | C | C | A | A | C | A | T | C | G | A | G | G | T | C | A | C | A | A |
| NC050284.1 | Ephemeroptera | *Torleya nepalica* | A | T | C | C | A | A | C | A | T | C | G | A | G | G | T | C | A | C | A | A |
| HM067837.1 | Ephemeroptera | *Vietnamella dabieshanensis* | A | T | C | C | A | A | C | A | T | C | G | A | G | G | T | C | G | C | A | A |
| KM244655.1 | Ephemeroptera | *Vietnamella* sp. | A | T | C | C | A | A | C | A | T | C | G | A | G | G | T | C | G | C | A | A |
| MH751435.1 | Odonata | *Acisoma panorpoides* | A | T | C | C | A | A | C | A | T | C | G | A | G | G | T | C | G | C | A | A |
| NC042689.1 | Odonata | *Acisoma panorpoides* | A | T | C | C | A | A | C | A | T | C | G | A | G | G | T | C | G | C | A | A |
| KX161841.1 | Odonata | *Anax imperator* | A | T | C | C | A | A | C | A | T | C | G | A | G | G | T | C | G | C | A | A |
| NC031821.1 | Odonata | *Anax imperator* | A | T | C | C | A | A | C | A | T | C | G | A | G | G | T | C | G | C | A | A |
| KP233805.1 | Odonata | *Atrocalopteryx* | A | T | C | C | A | A | C | A | T | C | G | A | G | G | T | C | G | C | A | A |

| | | | | | | | | | | | | | | | | | | | | | | |
|---|---|---|---|---|---|---|---|---|---|---|---|---|---|---|---|---|---|---|---|---|---|---|
| | | *atrata* | | | | | | | | | | | | | | | | | | | | |
| NC027181.1 | Odonata | *Atrocalopteryx atrata* | A | T | C | C | A | A | C | A | T | C | G | A | G | G | T | C | G | C | A | A |
| MG011692.1 | Odonata | *Atrocalopteryx melli* | A | T | C | C | A | A | C | A | T | C | G | A | G | G | T | C | G | C | A | A |
| NC036749.1 | Odonata | *Atrocalopteryx melli* | A | T | C | C | A | A | C | A | T | C | G | A | G | G | T | C | G | C | A | A |
| KM658172.1 | Odonata | *Brachythemis contaminata* | A | T | C | C | A | A | C | A | T | C | G | A | G | G | T | C | G | C | A | A |
| NC026305.1 | Odonata | *Brachythemis contaminata* | A | T | C | C | A | A | C | A | T | C | G | A | G | G | T | C | G | C | A | A |
| EU591677.1 | Odonata | *Davidius lunatus* | A | T | C | C | A | A | C | A | T | C | G | A | G | G | T | C | G | C | A | A |
| NC012644.1 | Odonata | *Davidius lunatus* | A | T | C | C | A | A | C | A | T | C | G | A | G | G | T | C | G | C | A | A |
| MH751436.1 | Odonata | *Deielia phaon* | A | T | C | C | A | A | C | A | T | C | G | A | G | G | T | C | G | C | A | A |
| NC042690.1 | Odonata | *Deielia phaon* | A | T | C | C | A | A | C | A | T | C | G | A | G | G | T | C | G | C | A | A |
| MF716899.1 | Odonata | *Enallagma cyathigerum* | A | T | C | C | A | A | C | A | T | C | G | A | G | G | T | C | G | C | A | A |
| JX050223.1 | Odonata | *Epiophlebia superstes* | A | T | C | C | A | A | C | A | T | C | G | A | G | G | T | C | G | C | A | A |
| NC023232.1 | Odonata | *Epiophlebia superstes* | A | T | C | C | A | A | C | A | T | C | G | A | G | G | T | C | G | C | A | A |
| MK522522.1 | Odonata | *Epophthalmia elegans* | A | T | C | C | A | A | C | A | T | C | G | A | G | G | T | C | G | C | A | A |
| NC046756.1 | Odonata | *Epophthalmia* | A | T | C | C | A | A | C | A | T | C | G | A | G | G | T | C | G | C | A | A |

| | | | | | | | | | | | | | | | | | | | | | | |
|---|---|---|---|---|---|---|---|---|---|---|---|---|---|---|---|---|---|---|---|---|---|---|
| | | *elegans* | | | | | | | | | | | | | | | | | | | | |
| HM126547.1 | Odonata | *Euphaea formosa* | A | T | C | C | A | A | C | A | T | C | G | A | G | G | T | C | G | C | A | A |
| NC014493.1 | Odonata | *Euphaea formosa* | A | T | C | C | A | A | C | A | T | C | G | A | G | G | T | C | G | C | A | A |
| KM244659.1 | Odonata | *Hydrobasileus croceus* | A | T | C | C | A | A | C | A | T | C | G | A | G | G | T | C | G | C | A | A |
| NC025758.1 | Odonata | *Hydrobasileus croceus* | A | T | C | C | A | A | C | A | T | C | G | A | G | G | T | C | G | C | A | A |
| KM244673.1 | Odonata | *Ictinogomphus* sp. | A | T | C | C | A | A | C | A | T | C | G | A | G | G | T | C | G | C | A | A |
| KU958378.1 | Odonata | *Ischnura elegans* | A | T | C | C | A | A | C | A | T | C | G | A | G | G | T | C | G | C | A | A |
| MK951668.1 | Odonata | *Ischnura elegans* | A | T | C | C | A | A | C | A | T | C | G | A | G | G | T | C | G | C | A | A |
| NC031824.1 | Odonata | *Ischnura elegans* | A | T | C | C | A | A | C | A | T | C | G | A | G | G | T | C | G | C | A | A |
| KC878732.1 | Odonata | *Ischnura pumilio* | A | T | C | C | A | A | C | A | T | C | G | A | G | G | T | C | G | C | A | A |
| NC021617.1 | Odonata | *Ischnura pumilio* | A | T | C | C | A | A | C | A | T | C | G | A | G | G | T | C | G | C | A | A |
| MF990748.1 | Odonata | *Macromia daimoji* | A | T | C | C | A | A | C | A | T | C | G | A | G | G | T | C | G | C | A | A |
| NC041425.1 | Odonata | *Macromia daimoji* | A | T | C | C | A | A | C | A | T | C | G | A | G | G | T | C | G | C | A | A |
| MK722304.1 | Odonata | *Matrona basilaris* | A | T | C | C | A | A | C | A | T | C | G | A | G | G | T | C | G | C | A | A |
| MK722305.1 | Odonata | *Matrona basilaris* | A | T | C | C | A | A | C | A | T | C | G | A | G | G | T | C | G | C | A | A |
| KU958377.1 | Odonata | *Megaloprepus caerulatus* | A | T | C | C | A | A | C | A | T | C | G | A | G | G | T | C | G | C | A | A |
| NC031823.1 | Odonata | *Megaloprepus caerulatus* | A | T | C | C | A | A | C | A | T | C | G | A | G | G | T | C | G | C | A | A |
| KU871065.1 | Odonata | *Mnais costalis* | A | T | C | C | A | A | C | A | T | C | G | A | G | G | T | C | G | C | A | A |

| | | | | | | | | | | | | | | | | | | | | | | |
|---|---|---|---|---|---|---|---|---|---|---|---|---|---|---|---|---|---|---|---|---|---|---|
| KY402222.1 | Odonata | *Nannophya pygmaea* | A | T | C | C | A | A | C | A | T | C | G | A | G | G | T | C | G | C | A | A |
| NC039410.1 | Odonata | *Nannophya pygmaea* | A | T | C | C | A | A | C | A | T | C | G | A | G | G | T | C | G | C | A | A |
| KU361233.1 | Odonata | *Orthetrum chrysis* | A | T | C | C | A | A | C | A | T | C | G | A | G | G | T | C | G | C | A | A |
| NC032048.1 | Odonata | *Orthetrum chrysis* | A | T | C | C | A | A | C | A | T | C | G | A | G | G | T | C | G | C | A | A |
| KU361232.1 | Odonata | *Orthetrum glaucum* | A | T | C | C | A | A | C | A | T | C | G | A | G | G | T | C | G | C | A | A |
| NC032047.1 | Odonata | *Orthetrum glaucum* | A | T | C | C | A | A | C | A | T | C | G | A | G | G | T | C | G | C | A | A |
| MH751437.1 | Odonata | *Orthetrum melania* | A | T | C | C | A | A | C | A | T | C | G | A | G | G | T | C | G | C | A | A |
| NC042732.1 | Odonata | *Orthetrum melania* | A | T | C | C | A | A | C | A | T | C | G | A | G | G | T | C | G | C | A | A |
| KU361234.1 | Odonata | *Orthetrum sabina* | A | T | C | C | A | A | C | A | T | C | G | A | G | G | T | C | G | C | A | A |
| MH751438.1 | Odonata | *Orthetrum sabina* | A | T | C | C | A | A | C | A | T | C | G | A | G | G | T | C | G | C | A | A |
| NC032049.1 | Odonata | *Orthetrum sabina* | A | T | C | C | A | A | C | A | T | C | G | A | G | G | T | C | G | C | A | A |
| KU361235.1 | Odonata | *Orthetrum testaceum* | A | T | C | C | A | A | C | A | T | C | G | A | G | G | T | C | G | C | A | A |
| MH751439.1 | Odonata | *Orthetrum testaceum* | A | T | C | C | A | A | C | A | T | C | G | A | G | G | T | C | G | C | A | A |
| NC032050.1 | Odonata | *Orthetrum testaceum* | A | T | C | C | A | A | C | A | T | C | G | A | G | G | T | C | G | C | A | A |
| AB126005.1 | Odonata | *Orthetrum triangulare melania* | A | T | C | C | A | A | C | A | T | C | G | A | G | G | T | C | G | C | A | A |
| MK951669.1 | Odonata | *Paracercion v-* | A | T | C | C | A | A | C | A | T | C | G | A | G | G | T | C | G | C | A | A |

| | | | | | | | | | | | | | | | | | | | | | | |
|---|---|---|---|---|---|---|---|---|---|---|---|---|---|---|---|---|---|---|---|---|---|---|
| | | *nigrum* | | | | | | | | | | | | | | | | | | | | |
| KP233804.1 | Odonata | *Platycnemis foliacea* | A | T | C | C | A | A | C | A | T | C | G | A | G | G | T | C | G | C | A | A |
| NC027180.1 | Odonata | *Platycnemis foliacea* | A | T | C | C | A | A | C | A | T | C | G | A | G | G | T | C | G | C | A | A |
| FJ606784.1 | Odonata | *Pseudolestes mirabilis* | A | T | C | C | A | A | C | A | T | C | G | A | G | G | T | C | G | C | A | A |
| NC020636.1 | Odonata | *Pseudolestes mirabilis* | A | T | C | C | A | A | C | A | T | C | G | A | G | G | T | C | G | C | A | A |
| MF150044.1 | Odonata | *Psolodesmus mandarinus* | A | T | C | C | A | A | C | A | T | C | G | A | G | G | T | C | G | C | A | A |
| NC042442.1 | Odonata | *Psolodesmus mandarinus* | A | T | C | C | A | A | C | A | T | C | G | A | G | G | T | C | G | C | A | A |
| MG594801.1 | Odonata | *Somatochlora hineana* | A | T | C | C | A | A | C | A | T | C | G | A | G | G | T | C | G | C | A | A |
| MH751440.1 | Odonata | *Tramea virginia* | A | T | C | C | A | A | C | A | T | C | G | A | G | G | T | C | G | C | A | A |
| NC042691.1 | Odonata | *Tramea virginia* | A | T | C | C | A | A | C | A | T | C | G | A | G | G | T | C | G | C | A | A |
| MH751441.1 | Odonata | *Trigomphus carus* | A | T | C | C | A | A | C | A | T | C | G | A | G | G | T | C | G | C | A | A |
| NC042733.1 | Odonata | *Trigomphus carus* | A | T | C | C | A | A | C | A | T | C | G | A | G | G | T | C | G | C | A | A |
| JX050224.1 | Odonata | *Vestalis melania* | A | T | C | C | A | A | C | A | T | C | G | A | G | G | T | C | G | C | A | A |
| NC023233.1 | Odonata | *Vestalis melania* | A | T | C | C | A | A | C | A | T | C | G | A | G | G | T | C | G | C | A | A |
| KM199685.1 | Plecoptera | *Acroneuria hainana* | A | T | C | C | A | A | C | A | T | C | G | A | G | G | T | C | G | C | A | A |
| NC026104.1 | Plecoptera | *Acroneuria hainana* | A | T | C | C | A | A | C | A | T | C | G | A | G | G | T | C | G | C | A | A |
| KX091846.1 | Plecoptera | *Agnetina aequalis* | A | T | C | C | A | A | C | A | T | C | G | A | G | G | T | C | G | C | A | A |

| | | | | | | | | | | | | | | | | | | | | | | |
|---|---|---|---|---|---|---|---|---|---|---|---|---|---|---|---|---|---|---|---|---|---|---|
| MH085446.1 | Plecoptera | *Amphinemura longispina* | A | T | C | C | A | A | C | A | T | C | G | A | G | G | T | C | G | C | A | A |
| NC044748.1 | Plecoptera | *Amphinemura longispina* | A | T | C | C | A | A | C | A | T | C | G | A | G | G | T | C | G | C | A | A |
| KX091847.1 | Plecoptera | *Amphinemura sp.* | A | T | C | C | A | A | C | A | T | C | G | A | G | G | T | C | G | C | A | A |
| MH085447.1 | Plecoptera | *Amphinemura yao* | A | T | C | C | A | A | C | A | T | C | G | A | G | G | T | C | G | C | A | A |
| NC044749.1 | Plecoptera | *Amphinemura yao* | A | T | C | C | A | A | C | A | T | C | G | A | G | G | T | C | G | C | A | A |
| MK111413.1 | Plecoptera | *Antarctoperla michaelseni* | A | T | C | C | A | A | C | A | T | C | G | A | G | G | T | C | G | C | A | A |
| NC042199.1 | Plecoptera | *Antarctoperla michaelseni* | A | T | C | C | A | A | C | A | T | C | G | A | G | G | T | C | G | C | A | A |
| KR604721.1 | Plecoptera | *Apteroperla tikumana* | A | T | C | C | A | A | C | A | T | C | G | A | G | G | T | C | G | C | A | A |
| NC027698.1 | Plecoptera | *Apteroperla tikumana* | A | T | C | C | A | A | C | A | T | C | G | A | G | G | T | C | G | C | A | A |
| MT410838.1 | Plecoptera | *Brachyptera risi* | A | T | C | C | A | A | C | A | T | C | G | A | G | G | T | C | G | C | A | A |
| MG677941.1 | Plecoptera | *Calineuria stigmatica* | A | T | C | C | A | A | C | A | T | C | G | A | G | G | T | C | G | C | A | A |
| KX094942.1 | Plecoptera | *Capnia zijinshana* | A | T | C | C | A | A | C | A | T | C | G | A | G | G | T | C | G | C | A | A |
| NC034661.1 | Plecoptera | *Capnia zijinshana* | A | T | C | C | A | A | C | A | T | C | G | A | G | G | T | C | G | C | A | A |
| MG677942.1 | Plecoptera | *Caroperla siveci* | A | T | C | C | A | A | C | A | T | C | G | A | G | G | T | C | G | C | A | A |
| MF100783.1 | Plecoptera | *Cerconychia* | A | T | C | C | A | A | C | A | T | C | G | A | G | G | T | C | G | C | A | A |

| | | | | | | | | | | | | | | | | | | | | | | |
|---|---|---|---|---|---|---|---|---|---|---|---|---|---|---|---|---|---|---|---|---|---|---|
| | | *flectospina* | | | | | | | | | | | | | | | | | | | | |
| MN419914.1 | Plecoptera | *Claassenia* sp. | A | T | C | C | A | A | C | A | T | C | G | A | G | G | T | C | G | C | A | A |
| KC952026.1 | Plecoptera | *Cryptoperla* sp. | A | T | C | C | A | A | C | A | T | C | G | A | G | G | T | C | G | C | A | A |
| MK111416.1 | Plecoptera | *Diamphipnoa annulata* | A | T | C | C | A | A | C | A | T | C | G | A | G | G | T | C | G | C | A | A |
| NC042205.1 | Plecoptera | *Diamphipnoa annulata* | A | T | C | C | A | A | C | A | T | C | G | A | G | G | T | C | G | C | A | A |
| MK111417.1 | Plecoptera | *Diamphipnopsis* sp. | A | T | C | C | A | A | C | A | T | C | G | A | G | G | T | C | G | C | A | A |
| KF484757.1 | Plecoptera | *Dinocras cephalotes* | A | T | C | C | A | A | C | A | T | C | G | A | G | G | T | C | G | C | A | A |
| NC022843.1 | Plecoptera | *Dinocras cephalotes* | A | T | C | C | A | A | C | A | T | C | G | A | G | G | T | C | G | C | A | A |
| MG589787.1 | Plecoptera | *Doddsia occidentalis* | A | T | C | C | A | A | C | A | T | C | G | A | G | G | T | C | G | C | A | A |
| MK905888.1 | Plecoptera | *Etrocorema hochii* | A | T | C | C | A | A | C | A | T | C | G | A | G | G | T | C | G | C | A | A |
| MK905206.1 | Plecoptera | *Flavoperla* sp. | A | T | C | C | A | A | C | A | T | C | G | A | G | G | T | C | G | C | A | A |
| MN419916.1 | Plecoptera | *Flavoperla* sp. | A | T | C | C | A | A | C | A | T | C | G | A | G | G | T | C | G | C | A | A |
| MN419915.1 | Plecoptera | *Indonemoura auriformis* | A | T | C | C | A | A | C | A | T | C | G | A | G | G | T | C | G | C | A | A |
| MH085448.1 | Plecoptera | *Indonemoura jacobsoni* | A | T | C | C | A | A | C | A | T | C | G | A | G | G | T | C | G | C | A | A |
| NC044750.1 | Plecoptera | *Indonemoura jacobsoni* | A | T | C | C | A | A | C | A | T | C | G | A | G | G | T | C | G | C | A | A |
| MH085449.1 | Plecoptera | *Indonemoura* | A | T | C | C | A | A | C | A | T | C | G | A | G | G | T | C | G | C | A | A |

| | | | | | | | | | | | | | | | | | | | | | | |
|---|---|---|---|---|---|---|---|---|---|---|---|---|---|---|---|---|---|---|---|---|---|---|
| | | *nohirae* | | | | | | | | | | | | | | | | | | | | |
| NC044751.1 | Plecoptera | *Indonemoura nohirae* | A | T | C | C | A | A | C | A | T | C | G | A | G | G | T | C | G | C | A | A |
| MF716959.1 | Plecoptera | *Isoperla bilineata* | A | T | C | C | A | A | C | A | T | C | G | A | G | G | T | C | G | C | A | A |
| NC038190.1 | Plecoptera | *Isoperla bilineata* | A | T | C | C | A | A | C | A | T | C | G | A | G | G | T | C | G | C | A | A |
| MG910457.1 | Plecoptera | *Isoperla eximia* | A | T | C | C | A | A | C | A | T | C | G | A | G | G | T | C | G | C | A | A |
| NC038167.1 | Plecoptera | *Isoperla eximia* | A | T | C | C | A | A | C | A | T | C | G | A | G | G | T | C | G | C | A | A |
| KT186102.1 | Plecoptera | *Kamimuria chungnanshana* | A | T | C | C | A | A | C | A | T | C | G | A | G | G | T | C | G | C | A | A |
| NC028076.1 | Plecoptera | *Kamimuria chungnanshana* | A | T | C | C | A | A | C | A | T | C | G | A | G | G | T | C | G | C | A | A |
| MN400755.1 | Plecoptera | *Kamimuria klapaleki* | A | T | C | C | A | A | C | A | T | C | G | A | G | G | T | C | G | C | A | A |
| MK111414.1 | Plecoptera | *Klapopteryx armillata* | A | T | C | C | A | A | C | A | T | C | G | A | G | G | T | C | G | C | A | A |
| MH374046.1 | Plecoptera | *Lednia tumana* | A | T | C | C | A | A | C | A | T | C | G | A | G | G | T | C | G | C | A | A |
| MK568475.1 | Plecoptera | *Leuctra fusca* | A | T | C | C | A | A | C | A | T | C | G | A | G | G | T | C | G | C | A | A |
| KP642637.1 | Plecoptera | *Mesocapnia arizonensis* | A | T | C | C | A | A | C | A | T | C | G | A | G | G | T | C | G | C | A | A |
| KY568983.1 | Plecoptera | *Mesocapnia daxingana* | A | T | C | C | A | A | C | A | T | C | G | A | G | G | T | C | G | C | A | A |
| MH085450.1 | Plecoptera | *Mesonemoura metafiligera* | A | T | C | C | A | A | C | A | T | C | G | A | G | G | T | C | G | C | A | A |

| | | | | | | | | | | | | | | | | | | | | | | |
|---|---|---|---|---|---|---|---|---|---|---|---|---|---|---|---|---|---|---|---|---|---|---|
| NC044719.1 | Plecoptera | *Mesonemoura metafiligera* | A | T | C | C | A | A | C | A | T | C | G | A | G | G | T | C | G | C | A | A |
| MH085451.1 | Plecoptera | *Mesonemoura tritaenia* | A | T | C | C | A | A | C | A | T | C | G | A | G | G | T | C | G | C | A | A |
| NC044720.1 | Plecoptera | *Mesonemoura tritaenia* | A | T | C | C | A | A | C | A | T | C | G | A | G | G | T | C | G | C | A | A |
| MN096323.1 | Plecoptera | *Microperla geei* | A | T | C | C | A | A | C | A | T | C | G | A | G | G | T | C | G | C | A | A |
| MT410776.1 | Plecoptera | *Nemoura avicularis* | A | T | C | C | A | A | C | A | T | C | G | A | G | G | T | C | G | C | A | A |
| MT410849.1 | Plecoptera | *Nemoura cinerea* | A | T | C | C | A | A | C | A | T | C | G | A | G | G | T | C | G | C | A | A |
| KY940360.1 | Plecoptera | *Nemoura nankinensis* | A | T | C | C | A | A | C | A | T | C | G | A | G | G | T | C | G | C | A | A |
| NC034939.1 | Plecoptera | *Nemoura nankinensis* | A | T | C | C | A | A | C | A | T | C | G | A | G | G | T | C | G | C | A | A |
| MK290826.1 | Plecoptera | *Nemoura papilla* | A | T | C | C | A | A | C | A | T | C | G | A | G | G | T | C | G | C | A | A |
| MK111418.1 | Plecoptera | *Neonemura barrosi* | A | T | C | C | A | A | C | A | T | C | G | A | G | G | T | C | G | C | A | A |
| NC042206.1 | Plecoptera | *Neonemura barrosi* | A | T | C | C | A | A | C | A | T | C | G | A | G | G | T | C | G | C | A | A |
| KX091858.1 | Plecoptera | *Neoperla ignacsiveci* | A | T | C | C | A | A | C | A | T | C | G | A | G | G | T | C | G | C | A | A |
| KX091859.1 | Plecoptera | *Neoperla* sp. | A | T | C | C | A | A | C | A | T | C | G | A | G | G | T | C | G | C | A | A |
| MN400756.1 | Plecoptera | *Neoperlops gressitti* | A | T | C | C | A | A | C | A | T | C | G | A | G | G | T | C | G | C | A | A |
| MK111415.1 | Plecoptera | *Neuroperla schedingi* | A | T | C | C | A | A | C | A | T | C | G | A | G | G | T | C | G | C | A | A |

| | | | | | | | | | | | | | | | | | | | | | | |
|---|---|---|---|---|---|---|---|---|---|---|---|---|---|---|---|---|---|---|---|---|---|---|
| NC042200.1 | Plecoptera | *Neuroperla schedingi* | A | T | C | C | A | A | C | A | T | C | G | A | G | G | T | C | G | C | A | A |
| MK686067.1 | Plecoptera | *Niponiella limbatella* | A | T | C | C | A | A | C | A | T | C | G | A | G | G | T | C | G | C | A | A |
| MK387068.1 | Plecoptera | *Peltoperlopsis cebuano* | A | T | C | C | A | A | C | A | T | C | G | A | G | G | T | C | G | C | A | A |
| MF197377.1 | Plecoptera | *Perlodes* sp. | A | T | C | C | A | A | C | A | T | C | G | A | G | G | T | C | G | C | A | A |
| MH085452.1 | Plecoptera | *Protonemura kohnoae* | A | T | C | C | A | A | C | A | T | C | G | A | G | G | T | C | G | C | A | A |
| NC044752.1 | Plecoptera | *Protonemura kohnoae* | A | T | C | C | A | A | C | A | T | C | G | A | G | G | T | C | G | C | A | A |
| MT410850.1 | Plecoptera | *Protonemura meyeri* | A | T | C | C | A | A | C | A | T | C | G | A | G | G | T | C | G | C | A | A |
| NC050322.1 | Plecoptera | *Protonemura meyeri* | A | T | C | C | A | A | C | A | T | C | G | A | G | G | T | C | G | C | A | A |
| MH085453.1 | Plecoptera | *Protonemura orbiculata* | A | T | C | C | A | A | C | A | T | C | G | A | G | G | T | C | G | C | A | A |
| NC044753.1 | Plecoptera | *Protonemura orbiculata* | A | T | C | C | A | A | C | A | T | C | G | A | G | G | T | C | G | C | A | A |
| MG910458.1 | Plecoptera | *Pseudomegarcys japonica* | A | T | C | C | A | A | C | A | T | C | G | A | G | G | T | C | G | C | A | A |
| NC038168.1 | Plecoptera | *Pseudomegarcys japonica* | A | T | C | C | A | A | C | A | T | C | G | A | G | G | T | C | G | C | A | A |
| KU182360.1 | Plecoptera | *Pteronarcella badia* | A | T | C | C | A | A | C | A | T | C | G | A | G | G | T | C | G | C | A | A |
| NC029248.1 | Plecoptera | *Pteronarcella badia* | A | T | C | C | A | A | C | A | T | C | G | A | G | G | T | C | G | C | A | A |

| | | | | | | | | | | | | | | | | | | | | | | |
|---|---|---|---|---|---|---|---|---|---|---|---|---|---|---|---|---|---|---|---|---|---|---|
| AY687866.1 | Plecoptera | *Pteronarcys princeps* | A | T | C | C | A | A | C | A | T | C | G | A | G | G | T | C | G | C | A | A |
| NC006133.1 | Plecoptera | *Pteronarcys princeps* | A | T | C | C | A | A | C | A | T | C | G | A | G | G | T | C | G | C | A | A |
| MK111419.1 | Plecoptera | *Rhopalopsole bulbifera* | A | T | C | C | A | A | C | A | T | C | G | A | G | G | T | C | G | C | A | A |
| NC042207.1 | Plecoptera | *Rhopalopsole bulbifera* | A | T | C | C | A | A | C | A | T | C | G | A | G | G | T | C | G | C | A | A |
| MH510071.1 | Plecoptera | *Scopura longa* | A | T | C | C | A | A | C | A | T | C | G | A | G | G | T | C | G | C | A | A |
| NC041105.1 | Plecoptera | *Scopura longa* | A | T | C | C | A | A | C | A | T | C | G | A | G | G | T | C | G | C | A | A |
| MK492253.1 | Plecoptera | *Sinacroneuria dabieshana* | A | T | C | C | A | A | C | A | T | C | G | A | G | G | T | C | G | C | A | A |
| MF716958.1 | Plecoptera | *Soliperla* sp. | A | T | C | C | A | A | C | A | T | C | G | A | G | G | T | C | G | C | A | A |
| NC038189.1 | Plecoptera | *Soliperla* sp. | A | T | C | C | A | A | C | A | T | C | G | A | G | G | T | C | G | C | A | A |
| MH085455.1 | Plecoptera | *Sphaeronemoura acutispina* | A | T | C | C | A | A | C | A | T | C | G | A | G | G | T | C | G | C | A | A |
| MH085454.1 | Plecoptera | *Sphaeronemoura grandicauda* | A | T | C | C | A | A | C | A | T | C | G | A | G | G | T | C | G | C | A | A |
| NC044754.1 | Plecoptera | *Sphaeronemoura grandicauda* | A | T | C | C | A | A | C | A | T | C | G | A | G | G | T | C | G | C | A | A |
| MK111420.1 | Plecoptera | *Sphaeronemoura hainana* | A | T | C | C | A | A | C | A | T | C | G | A | G | G | T | C | G | C | A | A |

| | | | | | | | | | | | | | | | | | | | | | | |
|---|---|---|---|---|---|---|---|---|---|---|---|---|---|---|---|---|---|---|---|---|---|---|
| NC044755.1 | Plecoptera | *Sphaeronemoura hamistyla* | A | T | C | C | A | A | C | A | T | C | G | A | G | G | T | C | G | C | A | A |
| KR088971.1 | Plecoptera | *Styloperla* sp. | A | T | C | C | A | A | C | A | T | C | G | A | G | G | T | C | G | C | A | A |
| KX845569.1 | Plecoptera | *Styloperla spinicercia* | A | T | C | C | A | A | C | A | T | C | G | A | G | G | T | C | G | C | A | A |
| NC034809.1 | Plecoptera | *Styloperla spinicercia* | A | T | C | C | A | A | C | A | T | C | G | A | G | G | T | C | G | C | A | A |
| KX091865.1 | Plecoptera | *Suwallia bimaculata* | A | T | C | C | A | A | C | A | T | C | G | A | G | G | T | C | G | C | A | A |
| MN121757.1 | Plecoptera | *Suwallia bimaculata* | A | T | C | C | A | A | C | A | T | C | G | A | G | G | T | C | G | C | A | A |
| MF198253.1 | Plecoptera | *Suwallia teleckojensis* | A | T | C | C | A | A | C | A | T | C | G | A | G | G | T | C | G | C | A | A |
| NC037754.1 | Plecoptera | *Suwallia teleckojensis* | A | T | C | C | A | A | C | A | T | C | G | A | G | G | T | C | G | C | A | A |
| MG589786.1 | Plecoptera | *Taeniopteryx ugola* | A | T | C | C | A | A | C | A | T | C | G | A | G | G | T | C | G | C | A | A |
| NC037897.1 | Plecoptera | *Taeniopteryx ugola* | A | T | C | C | A | A | C | A | T | C | G | A | G | G | T | C | G | C | A | A |
| KM409708.1 | Plecoptera | *Togoperla* sp. | A | T | C | C | A | A | C | A | T | C | G | A | G | G | T | C | G | C | A | A |
| KY522907.1 | Plecoptera | *Zelandoperla fenestrata* | A | T | C | C | A | A | C | A | T | C | G | A | G | G | T | C | G | C | A | A |
| NC034997.1 | Plecoptera | *Zelandoperla fenestrata* | A | T | C | C | A | A | C | A | T | C | G | A | G | G | T | C | G | C | A | A |
| FJ456939.1 | Heteroptera | *Aphelocheirus ellipsoideus* | A | T | T | C | A | A | C | A | T | C | G | A | G | G | T | C | G | C | A | A |

| | | | | | | | | | | | | | | | | | | | | | |
|---|---|---|---|---|---|---|---|---|---|---|---|---|---|---|---|---|---|---|---|---|---|---|
| KX505849.1 | Heteroptera | *Aphelocheirus jendeki* | A | T | T | C | A | A | C | A | T | C | G | A | G | G | T | C | G | C | A | A |
| NC037370.1 | Heteroptera | *Aphelocheirus jendeki* | A | T | T | C | A | A | C | A | T | C | G | A | G | G | T | C | G | C | A | A |
| FJ456944.1 | Heteroptera | *Aquarius paludum* | A | T | T | C | A | A | C | A | T | C | G | A | G | G | T | C | G | C | A | A |
| NC012841.1 | Heteroptera | *Aquarius paludum* | A | T | T | C | A | A | C | A | T | C | G | A | G | G | T | C | G | C | A | A |
| MK251109.1 | Heteroptera | *Diplonychus esakii* | A | T | T | C | A | A | C | A | T | C | G | A | G | G | T | C | G | C | A | A |
| FJ456940.1 | Heteroptera | *Diplonychus rusticus* | A | T | T | C | A | A | C | A | T | C | G | A | G | G | T | C | G | C | A | A |
| FJ456949.1 | Heteroptera | *Enithares tibialis* | A | T | T | C | A | A | C | A | T | C | G | A | G | G | T | C | G | C | A | A |
| NC012819.1 | Heteroptera | *Enithares tibialis* | A | T | T | C | A | A | C | A | T | C | G | A | G | G | T | C | G | C | A | A |
| KP400582.1 | Heteroptera | *Entomovelia* sp. | A | T | T | C | A | A | C | A | T | C | G | A | G | G | T | C | G | C | A | A |
| MF177288.1 | Heteroptera | *Gigantometra gigas* | A | T | T | C | A | A | C | A | T | C | G | A | G | G | T | C | G | C | A | A |
| MF177289.1 | Heteroptera | *Gigantometra gigas* | A | T | T | C | A | A | C | A | T | C | G | A | G | G | T | C | G | C | A | A |
| MF177290.1 | Heteroptera | *Gigantometra gigas* | A | T | T | C | A | A | C | A | T | C | G | A | G | G | T | C | G | C | A | A |
| MF177291.1 | Heteroptera | *Gigantometra gigas* | A | T | T | C | A | A | C | A | T | C | G | A | G | G | T | C | G | C | A | A |
| NC041084.1 | Heteroptera | *Gigantometra gigas* | A | T | T | C | A | A | C | A | T | C | G | A | G | G | T | C | G | C | A | A |
| KJ027513.1 | Heteroptera | *Helotrephes semiglobosus semiglobosus* | A | T | T | C | A | A | C | A | T | C | G | A | G | G | T | C | G | C | A | A |
| FJ456951.1 | Heteroptera | *Helotrephes* sp. | A | T | T | C | A | A | C | A | T | C | G | A | G | G | T | C | G | C | A | A |
| NC012822.1 | Heteroptera | *Helotrephes* sp. | A | T | T | C | A | A | C | A | T | C | G | A | G | G | T | C | G | C | A | A |

| | | | | | | | | | | | | | | | | | | | | | | |
|---|---|---|---|---|---|---|---|---|---|---|---|---|---|---|---|---|---|---|---|---|---|---|
| FJ456945.1 | Heteroptera | *Hydrometra greeni* | A | T | T | C | A | A | C | A | T | C | G | G | G | G | T | C | G | C | A | A |
| NC012842.1 | Heteroptera | *Hydrometra greeni* | A | T | T | C | A | A | C | A | T | C | G | G | G | G | T | C | G | C | A | A |
| FJ456947.1 | Heteroptera | *Ilyocoris cimicoides* | A | T | T | C | A | A | C | A | T | C | G | A | G | G | T | C | G | C | A | A |
| NC012845.1 | Heteroptera | *Ilyocoris cimicoides* | A | T | T | C | A | A | C | A | T | C | G | A | G | G | T | C | G | C | A | A |
| FJ456948.1 | Heteroptera | *Laccotrephes robustus* | A | T | T | C | A | A | C | A | T | C | G | A | G | G | T | C | G | C | A | A |
| NC012817.1 | Heteroptera | *Laccotrephes robustus* | A | T | T | C | A | A | C | A | T | C | G | A | G | G | T | C | G | C | A | A |
| FJ456946.1 | Heteroptera | *Leptopus* sp. | A | T | C | C | A | A | C | A | T | C | G | A | G | G | T | C | G | C | A | A |
| JQ910985.1 | Heteroptera | *Lethocerus deyrollei* | A | T | T | C | A | A | C | A | T | C | G | A | G | G | T | C | G | C | A | A |
| KU237288.1 | Heteroptera | *Lethocerus deyrollei* | A | T | T | C | A | A | C | A | T | C | G | A | G | G | T | C | G | C | A | A |
| KM588201.1 | Heteroptera | *Lethocerus indicus* | A | T | T | C | A | A | C | A | T | C | G | A | G | G | T | C | G | C | A | A |
| KR072671.1 | Heteroptera | *Lethocerus indicus* | A | T | T | C | A | A | C | A | T | C | G | A | G | G | T | C | G | C | A | A |
| NC027194.1 | Heteroptera | *Lethocerus indicus* | A | T | T | C | A | A | C | A | T | C | G | A | G | G | T | C | G | C | A | A |
| FJ456942.1 | Heteroptera | *Lycorma delicatula* | A | T | C | C | A | A | C | A | T | C | G | A | G | G | T | C | G | C | A | A |
| NC012835.1 | Heteroptera | *Lycorma delicatula* | A | T | C | C | A | A | C | A | T | C | G | A | G | G | T | C | G | C | A | A |
| MN027278.1 | Heteroptera | *Metrocoris* sp. | A | T | T | C | A | A | C | A | T | C | G | A | G | G | T | C | G | C | A | A |
| MH382835.1 | Heteroptera | *Micronecta sahlbergii* | A | T | T | C | A | A | C | A | T | C | G | A | G | G | T | C | G | C | A | A |
| NC039588.1 | Heteroptera | *Micronecta sahlbergii* | A | T | T | C | A | A | C | A | T | C | G | A | G | G | T | C | G | C | A | A |
| KT031802.1 | Heteroptera | *Nepa hoffmanni* | A | T | T | C | A | A | C | A | T | C | G | A | G | G | T | C | G | C | A | A |

| | | | | | | | | | | | | | | | | | | | | | | |
|---|---|---|---|---|---|---|---|---|---|---|---|---|---|---|---|---|---|---|---|---|---|---|
| NC028084.1 | Heteroptera | *Nepa hoffmanni* | A | T | T | C | A | A | C | A | T | C | G | A | G | G | T | C | G | C | A | A |
| FJ456943.1 | Heteroptera | *Nerthra indica* | A | T | T | C | A | A | C | A | T | C | G | A | G | G | T | C | G | C | A | A |
| NC012838.1 | Heteroptera | *Nerthra indica* | A | T | T | C | A | A | C | A | T | C | G | A | G | G | T | C | G | C | A | A |
| KX034086.1 | Heteroptera | *Notonecta chinensis* | A | T | T | C | A | A | C | A | T | C | G | A | G | G | T | C | G | C | A | A |
| NC036671.1 | Heteroptera | *Notonecta chinensis* | A | T | T | C | A | A | C | A | T | C | G | A | G | G | T | C | G | C | A | A |
| MH780861.1 | Heteroptera | *Notonecta montandoni* | A | T | T | C | A | A | C | A | T | C | G | A | G | G | T | C | G | C | A | A |
| NC045886.1 | Heteroptera | *Notonecta montandoni* | A | T | T | C | A | A | C | A | T | C | G | A | G | G | T | C | G | C | A | A |
| KM244707.1 | Heteroptera | *Notonectidae* sp. | A | T | T | C | A | A | C | A | T | C | G | A | G | G | T | C | G | C | A | A |
| FJ456950.1 | Heteroptera | *Ochterus marginatus* | A | T | T | C | A | A | C | A | T | C | G | A | G | G | T | C | G | C | A | A |
| KJ027516.1 | Heteroptera | *Paraplea frontalis* | A | T | T | C | A | A | C | A | T | C | G | A | G | G | T | C | G | C | A | A |
| NC028629.1 | Heteroptera | *Paraplea frontalis* | A | T | T | C | A | A | C | A | T | C | G | A | G | G | T | C | G | C | A | A |
| JQ910988.1 | Heteroptera | *Perittopus* sp. | A | T | T | C | A | A | C | A | T | C | G | A | G | G | T | C | G | C | A | A |
| KP400583.1 | Heteroptera | *Ptilomera tigrina* | A | T | T | C | A | A | C | A | T | C | G | A | G | G | T | C | G | C | A | A |
| MN027271.1 | Heteroptera | *Rhagovelia reitteri* | A | T | T | C | A | A | C | A | T | C | G | A | G | G | T | C | G | C | A | A |
| NC044721.1 | Heteroptera | *Rhagovelia reitteri* | A | T | T | C | A | A | C | A | T | C | G | A | G | G | T | C | G | C | A | A |
| MN027279.1 | Heteroptera | *Rhagovelia* sp. | A | T | T | C | A | A | C | A | T | C | G | A | G | G | T | C | G | C | A | A |
| FJ456941.1 | Heteroptera | *Sigara septemlineata* | A | T | T | C | A | A | C | A | T | C | G | A | G | G | T | C | G | C | A | A |
| FJ456952.1 | Heteroptera | *Valentia hoffmanni* | A | T | C | C | A | A | C | A | T | C | G | A | G | G | T | C | G | C | A | A |

| | | | | | | | | | | | | | | | | | | | | | | |
|---|---|---|---|---|---|---|---|---|---|---|---|---|---|---|---|---|---|---|---|---|---|---|
| NC012823.1 | Heteroptera | *Valentia hoffmanni* | A | T | C | C | A | A | C | A | T | C | G | A | G | G | T | C | G | C | A | A |
| KF840564.1 | Megaloptera | *Acanthacorydalis orientalis* | A | T | T | C | A | A | C | A | T | C | G | A | G | G | T | C | G | C | A | A |
| NC023462.1 | Megaloptera | *Acanthacorydalis orientalis* | A | T | T | C | A | A | C | A | T | C | G | A | G | G | T | C | G | C | A | A |
| KU925864.1 | Megaloptera | *Archichauliodes deceptor* | A | T | T | C | A | A | C | A | T | C | G | A | G | G | T | T | G | C | A | A |
| NC035504.1 | Megaloptera | *Archichauliodes deceptor* | A | T | T | C | A | A | C | A | T | C | G | A | G | G | T | T | G | C | A | A |
| KU302713.1 | Megaloptera | *Chloronia mirifica* | A | T | T | C | A | A | C | A | T | C | G | A | G | G | T | C | G | C | A | A |
| NC033349.1 | Megaloptera | *Chloronia mirifica* | A | T | T | C | A | A | C | A | T | C | G | A | G | G | T | C | G | C | A | A |
| FJ171323.1 | Megaloptera | *Corydalus cornutus* | A | T | T | C | A | A | C | A | T | C | G | A | G | G | T | C | G | C | A | A |
| NC011276.1 | Megaloptera | *Corydalus cornutus* | A | T | T | C | A | A | C | A | T | C | G | A | G | G | T | C | G | C | A | A |
| KJ806318.1 | Megaloptera | *Dysmicohermes ingens* | A | T | T | C | A | A | C | A | T | C | G | A | G | G | T | C | G | C | A | A |
| NC024657.1 | Megaloptera | *Dysmicohermes ingens* | A | T | T | C | A | A | C | A | T | C | G | A | G | G | T | C | G | C | A | A |
| JQ351950.1 | Megaloptera | *Neochauliodes bowringi* | A | T | T | C | A | A | C | A | T | C | G | A | G | G | T | C | G | C | A | A |
| NC023444.1 | Megaloptera | *Neochauliodes bowringi* | A | T | T | C | A | A | C | A | T | C | G | A | G | G | T | C | G | C | A | A |
| KF785799.1 | Megaloptera | *Neochauliodes* | A | T | T | C | A | A | C | A | T | C | G | A | G | G | T | C | G | C | A | A |

| | | | | | | | | | | | | | | | | | | | | | | |
|---|---|---|---|---|---|---|---|---|---|---|---|---|---|---|---|---|---|---|---|---|---|---|
| | | *fraternus* | | | | | | | | | | | | | | | | | | | | |
| NC025282.1 | Megaloptera | *Neochauliodes fraternus* | A | T | T | C | A | A | C | A | T | C | G | A | G | G | T | C | G | C | A | A |
| KX821680.1 | Megaloptera | *Neochauliodes parasparsus* | A | T | T | C | A | A | C | A | T | C | G | A | G | G | T | C | G | C | A | A |
| JX110703.1 | Megaloptera | *Neochauliodes punctatolosus* | A | T | T | C | A | A | C | A | T | C | G | A | G | G | T | C | G | C | A | A |
| MK301249.1 | Megaloptera | *Neochauliodes punctatolosus* | A | T | T | C | A | A | C | A | T | C | G | A | G | G | T | C | G | C | A | A |
| NC018772.1 | Megaloptera | *Neochauliodes punctatolosus* | A | T | T | C | A | A | C | A | T | C | G | A | G | G | T | C | G | C | A | A |
| KF771894.1 | Megaloptera | *Neochauliodes rotundatus* | A | T | T | C | A | A | C | A | T | C | G | A | G | G | T | C | G | C | A | A |
| NC025281.1 | Megaloptera | *Neochauliodes rotundatus* | A | T | T | C | A | A | C | A | T | C | G | A | G | G | T | C | G | C | A | A |
| KP126231.1 | Megaloptera | *Neoneuromus tonkinensis* | A | T | T | C | A | A | C | A | T | C | G | A | G | G | T | C | G | C | A | A |
| NC027852.1 | Megaloptera | *Neoneuromus tonkinensis* | A | T | T | C | A | A | C | A | T | C | G | A | G | G | T | C | G | C | A | A |
| KP126232.1 | Megaloptera | *Nevromus exterior* | A | T | T | C | A | A | C | A | T | C | G | A | G | G | T | C | G | C | A | A |
| NC027851.1 | Megaloptera | *Nevromus exterior* | A | T | T | C | A | A | C | A | T | C | G | A | G | G | T | C | G | C | A | A |
| KY230493.1 | Megaloptera | *Protochauliodes* | A | T | T | C | A | A | C | A | T | C | G | A | G | G | T | C | G | C | A | A |

| | | | | | | | | | | | | | | | | | | | | | | |
|---|---|---|---|---|---|---|---|---|---|---|---|---|---|---|---|---|---|---|---|---|---|---|
| | | *biconicus* | | | | | | | | | | | | | | | | | | | | |
| EU526394.1 | Megaloptera | *Protohermes concolorus* | A | T | T | C | A | A | C | A | T | C | G | A | G | G | T | C | G | C | A | A |
| NC011524.1 | Megaloptera | *Protohermes concolorus* | A | T | T | C | A | A | C | A | T | C | G | A | G | G | T | C | G | C | A | A |
| FJ859905.1 | Megaloptera | *Sialis hamata* | A | T | T | C | A | A | C | A | T | C | G | A | G | G | T | C | G | C | A | A |
| NC013256.1 | Megaloptera | *Sialis hamata* | A | T | T | C | A | A | C | A | T | C | G | A | G | G | T | C | G | C | A | A |
| KT876911.1 | Megaloptera | *Sialis* sp. | A | T | T | C | A | A | C | A | T | C | G | A | G | G | T | C | G | C | A | A |
| KT425084.1 | Megaloptera | *Stenosialis australiensis* | A | T | T | C | A | A | C | A | T | C | G | A | G | G | T | C | G | C | A | A |
| KY587199.1 | Neuroptera | *Abachrysa eureka* | A | T | T | C | A | A | C | A | T | C | G | A | G | G | T | C | G | C | A | A |
| NC015095.1 | Neuroptera | *Apochrysa matsumurae* | A | T | T | C | A | A | C | A | T | C | G | A | G | G | T | C | G | C | A | A |
| KC758703.1 | Neuroptera | *Ascalohybris subjacens* | A | T | T | C | A | A | C | A | T | C | G | A | G | G | T | C | G | C | A | A |
| NC021428.1 | Neuroptera | *Ascalohybris subjacens* | A | T | T | C | A | A | C | A | T | C | G | A | G | G | T | C | G | C | A | A |
| FJ171324.1 | Neuroptera | *Ascaloptynx appendiculatus* | A | T | T | C | A | A | C | A | T | C | G | A | G | G | T | C | G | C | A | A |
| NC011277.1 | Neuroptera | *Ascaloptynx appendiculatus* | A | T | T | C | A | A | C | A | T | C | G | A | G | G | T | C | G | C | A | A |
| KT425083.1 | Neuroptera | *Balmes birmanus* | A | T | T | C | A | A | C | A | T | C | G | A | G | G | T | C | G | C | A | A |

| | | | | | | | | | | | | | | | | | | | | | | |
|---|---|---|---|---|---|---|---|---|---|---|---|---|---|---|---|---|---|---|---|---|---|---|
| KX369241.1 | Neuroptera | *Bullanga florida* | A | T | T | C | A | A | C | A | T | C | G | A | G | G | T | C | G | C | A | A |
| NC032298.1 | Neuroptera | *Bullanga florida* | A | T | T | C | A | A | C | A | T | C | G | A | G | G | T | C | G | C | A | A |
| KT425069.1 | Neuroptera | *Chasmoptera huttii* | A | T | T | C | A | A | C | A | T | C | G | A | G | G | T | C | G | C | A | A |
| JX033119.1 | Neuroptera | *Chrysopa pallens* | A | T | T | C | A | A | C | A | T | C | G | A | G | G | T | C | G | C | A | A |
| NC019618.1 | Neuroptera | *Chrysopa pallens* | A | T | T | C | A | A | C | A | T | C | G | A | G | G | T | C | G | C | A | A |
| KJ778879.1 | Neuroptera | *Chrysoperla carnea* | A | T | T | C | A | A | C | A | T | C | G | A | G | G | T | C | G | C | A | A |
| KU877169.1 | Neuroptera | *Chrysoperla externa* | A | T | T | C | A | A | C | A | T | C | G | A | G | G | T | C | G | C | A | A |
| NC030341.1 | Neuroptera | *Chrysoperla externa* | A | T | T | C | A | A | C | A | T | C | G | A | G | G | T | C | G | C | A | A |
| KP027407.1 | Neuroptera | *Chrysoperla nipponensis* | A | T | T | C | A | A | C | A | T | C | G | A | G | G | T | C | G | C | A | A |
| NC015093.1 | Neuroptera | *Chrysoperla nipponensis* | A | T | T | C | A | A | C | A | T | C | G | A | G | G | T | C | G | C | A | A |
| KT425088.1 | Neuroptera | *Climacia areolaris* | A | T | T | C | A | A | C | A | T | C | G | A | G | G | T | C | A | C | A | A |
| KT425078.1 | Neuroptera | *Coniopteryx* sp. | A | T | T | C | A | A | C | A | T | C | G | A | G | G | T | C | G | C | A | A |
| KT425068.1 | Neuroptera | *Dendroleon pantherinus* | A | T | T | C | A | A | C | A | T | C | G | A | G | G | T | C | G | C | A | A |
| MK301246.1 | Neuroptera | *Dendroleon pantherinus* | A | T | T | C | A | A | C | A | T | C | G | A | G | G | T | C | G | C | A | A |
| KT425073.1 | Neuroptera | *Dilar* sp. | A | T | T | C | A | A | C | A | T | C | G | A | G | G | T | C | G | C | A | A |
| FJ859906.1 | Neuroptera | *Ditaxis biseriata* | A | T | T | C | A | A | C | A | T | C | G | A | G | G | T | C | G | C | A | A |
| NC013257.1 | Neuroptera | *Ditaxis biseriata* | A | T | T | C | A | A | C | A | T | C | G | A | G | G | T | C | G | C | A | A |
| KT425087.1 | Neuroptera | *Drepanepteryx* | A | T | T | C | A | A | C | A | T | C | G | A | G | G | T | C | G | C | A | A |

| | | | | | | | | | | | | | | | | | | | | | | |
|---|---|---|---|---|---|---|---|---|---|---|---|---|---|---|---|---|---|---|---|---|---|---|
| | | *phalaenoides* | | | | | | | | | | | | | | | | | | | | |
| KF701327.1 | Neuroptera | *Epacanthaclisis banksi* | A | T | T | C | A | A | C | A | T | C | G | A | G | G | T | C | G | C | A | A |
| NC025905.1 | Neuroptera | *Epacanthaclisis banksi* | A | T | T | C | A | A | C | A | T | C | G | A | G | G | T | C | G | C | A | A |
| MF972496.1 | Neuroptera | *Euclimacia badia* | A | T | T | C | A | A | C | A | T | C | G | A | G | G | T | C | G | C | A | A |
| NC039773.1 | Neuroptera | *Euclimacia badia* | A | T | T | C | A | A | C | A | T | C | G | A | G | G | T | C | G | C | A | A |
| KT425080.1 | Neuroptera | *Eumantispa harmandi* | A | T | T | C | A | A | C | A | T | C | G | A | G | G | T | C | G | C | A | A |
| MF972495.1 | Neuroptera | *Eumantispa harmandi* | A | T | T | C | A | A | C | A | T | C | G | A | G | G | T | C | G | C | A | A |
| NC039772.1 | Neuroptera | *Eumantispa harmandi* | A | T | T | C | A | A | C | A | T | C | G | A | G | G | T | C | G | C | A | A |
| KT425072.1 | Neuroptera | *Fontecilla graphicus* | A | T | T | C | A | A | C | A | T | C | G | A | G | G | T | C | G | C | A | A |
| KY364372.1 | Neuroptera | *Gatzara jezoensis* | A | T | T | C | A | A | C | A | T | C | G | A | G | G | T | C | G | C | A | A |
| MK341582.1 | Neuroptera | *Gryposmylus pennyi* | A | T | T | C | A | A | C | A | T | C | G | A | G | G | T | C | G | C | A | A |
| MN852445.1 | Neuroptera | *Hemerobius japonicus* | A | T | T | C | A | A | C | A | T | C | G | A | G | G | T | C | G | C | A | A |
| NC048517.1 | Neuroptera | *Hemerobius japonicus* | A | T | T | C | A | A | C | A | T | C | G | A | G | G | T | C | G | C | A | A |
| KT425077.1 | Neuroptera | *Heterosmylus* sp. | A | T | T | C | A | A | C | A | T | C | G | A | G | G | T | C | G | C | A | A |
| KT425085.1 | Neuroptera | *Inocellia fujiana* | A | T | T | C | A | A | C | A | T | C | G | A | G | G | T | C | G | C | A | A |

| | | | | | | | | | | | | | | | | | | | | | | |
|---|---|---|---|---|---|---|---|---|---|---|---|---|---|---|---|---|---|---|---|---|---|---|
| KY587200.1 | Neuroptera | *Italochrysa insignis* | A | T | T | C | A | A | C | A | T | C | G | A | G | G | T | C | G | T | A | A |
| KY587201.1 | Neuroptera | *Leucochrysa pretiosa* | A | T | T | C | A | A | C | A | T | C | G | A | G | G | T | C | G | C | A | A |
| NC015609.1 | Neuroptera | *Libelloides macaronius* | A | T | T | C | A | A | C | A | T | C | G | A | G | G | T | C | G | C | A | A |
| MF972494.1 | Neuroptera | *Mantispa japonica* | A | T | T | C | A | A | C | A | T | C | G | A | G | G | T | C | G | C | A | A |
| NC039771.1 | Neuroptera | *Mantispa japonica* | A | T | T | C | A | A | C | A | T | C | G | A | G | G | T | C | G | C | A | A |
| MK472073.1 | Neuroptera | *Micromus angulatus* | A | T | T | C | A | A | C | A | T | C | G | A | G | G | T | C | G | C | A | A |
| NC042677.1 | Neuroptera | *Micromus angulatus* | A | T | T | C | A | A | C | A | T | C | G | A | G | G | T | C | G | C | A | A |
| KT425075.1 | Neuroptera | *Micromus* sp. | A | T | T | C | A | A | C | A | T | C | G | A | G | G | T | C | G | C | A | A |
| KT425092.1 | Neuroptera | *Mucroberotha vesicaria* | A | T | T | C | A | A | C | A | T | C | G | A | G | G | T | C | G | C | A | A |
| KT425089.1 | Neuroptera | *Myiodactylus osmyloides* | A | T | T | C | A | A | C | A | T | C | G | A | G | G | T | C | G | C | A | A |
| MT118664.1 | Neuroptera | *Myrmeleon formicarius* | A | T | T | C | A | A | C | A | T | C | G | A | G | G | T | C | G | C | A | A |
| NC047285.1 | Neuroptera | *Myrmeleon formicarius* | A | T | T | C | A | A | C | A | T | C | G | A | G | G | T | C | G | C | A | A |
| KJ461323.1 | Neuroptera | *Myrmeleon immanis* | A | T | T | C | A | A | C | A | T | C | G | A | G | G | T | C | G | C | A | A |
| NC024826.1 | Neuroptera | *Myrmeleon immanis* | A | T | T | C | A | A | C | A | T | C | G | A | G | G | T | C | G | C | A | A |
| KT425071.1 | Neuroptera | *Nallachius americanus* | A | T | T | C | A | A | C | A | T | C | G | A | G | G | T | C | G | C | A | A |

| | | | | | | | | | | | | | | | | | | | | | | |
|---|---|---|---|---|---|---|---|---|---|---|---|---|---|---|---|---|---|---|---|---|---|---|
| KT425086.1 | Neuroptera | *Negha inflata* | A | T | T | C | A | A | C | A | T | C | G | A | G | G | T | C | G | C | A | A |
| KT425079.1 | Neuroptera | *Nemoptera coa* | A | T | T | C | A | A | C | A | T | C | G | A | G | G | T | C | G | C | A | A |
| KR078257.1 | Neuroptera | *Neuronema laminatum* | A | T | T | C | A | A | C | A | T | C | G | A | G | G | T | C | G | C | A | A |
| NC028153.1 | Neuroptera | *Neuronema laminatum* | A | T | T | C | A | A | C | A | T | C | G | A | G | G | T | C | G | C | A | A |
| KT425074.1 | Neuroptera | *Nevrorthus apatelios* | A | T | T | C | A | A | C | A | T | C | G | A | G | G | T | C | G | C | A | A |
| KT425076.1 | Neuroptera | *Nipponeurorthus fuscinervis* | A | T | T | C | A | A | C | A | T | C | G | A | G | G | T | C | G | C | A | A |
| KP264629.1 | Neuroptera | *Nothancyla verreauxi* | A | T | T | C | A | A | C | A | T | C | G | A | G | G | T | C | G | C | A | A |
| KP264630.1 | Neuroptera | *Nothochrysa* sp. | A | T | T | C | A | A | C | A | T | C | G | A | G | G | T | C | G | C | A | A |
| KJ461322.1 | Neuroptera | *Nymphes myrmeleonoides* | A | T | T | C | A | A | C | A | T | C | G | A | G | G | T | C | G | C | A | A |
| NC024825.1 | Neuroptera | *Nymphes myrmeleonoides* | A | T | T | C | A | A | C | A | T | C | G | A | G | G | T | C | G | C | A | A |
| KT425090.1 | Neuroptera | *Oliarces clara* | A | T | T | C | A | A | C | A | T | C | G | A | G | G | T | C | G | C | A | A |
| MN818868.1 | Neuroptera | *Osmylus atomatus* | A | T | T | C | A | A | C | A | T | C | G | A | G | G | T | C | G | C | A | A |
| MN818867.1 | Neuroptera | *Osmylus fulvicephalus* | A | T | T | C | A | A | C | A | T | C | G | A | G | G | T | C | G | C | A | A |
| MN617021.1 | Neuroptera | *Osmylus lucalatus* | A | T | T | C | A | A | C | A | T | C | G | A | G | G | T | C | G | C | A | A |

| | | | | | | | | | | | | | | | | | | | | | | |
|---|---|---|---|---|---|---|---|---|---|---|---|---|---|---|---|---|---|---|---|---|---|---|
| MN818870.1 | Neuroptera | *Osmylus* sp. | A | T | T | C | A | A | C | A | T | C | G | A | G | G | T | C | G | C | A | A |
| MN818869.1 | Neuroptera | *Osmylus zheanus* | A | T | T | C | A | A | C | A | T | C | G | A | G | G | T | C | G | C | A | A |
| KY587202.1 | Neuroptera | *Parankylopteryx sp.* | A | T | T | C | A | A | C | A | T | C | G | A | G | G | T | C | G | C | A | A |
| KT425091.1 | Neuroptera | *Podallea sp.* | A | T | T | C | A | A | C | A | T | C | G | A | G | G | T | C | G | C | A | A |
| FJ171325.1 | Neuroptera | *Polystoechotes punctatus* | A | T | T | C | A | A | C | A | T | C | G | A | G | G | T | C | G | C | A | A |
| NC011278.1 | Neuroptera | *Polystoechotes punctatus* | A | T | T | C | A | A | C | A | T | C | G | A | G | G | T | C | G | C | A | A |
| KT425082.1 | Neuroptera | *Psychopsis coelivaga* | A | T | T | C | A | A | C | A | T | C | G | A | G | G | T | C | G | C | A | A |
| KF626446.1 | Neuroptera | *Rapisma xizangense* | A | T | T | C | A | A | C | A | T | C | G | A | G | G | T | C | G | C | A | A |
| NC023362.1 | Neuroptera | *Rapisma xizangense* | A | T | T | C | A | A | C | A | T | C | G | A | G | G | T | C | G | C | A | A |
| KF626447.1 | Neuroptera | *Rapisma zayuanum* | A | T | T | C | A | A | C | A | T | C | G | A | G | G | T | C | G | C | A | A |
| NC023363.1 | Neuroptera | *Rapisma zayuanum* | A | T | T | C | A | A | C | A | T | C | G | A | G | G | T | C | G | C | A | A |
| KT425067.1 | Neuroptera | *Semidalis aleyrodiformis* | A | T | T | C | A | A | C | A | T | C | G | A | G | G | T | C | G | C | A | A |
| KT425070.1 | Neuroptera | *Sisyra nigra* | A | T | T | C | A | A | C | A | T | C | G | A | G | G | T | C | A | C | A | A |
| KT425081.1 | Neuroptera | *Stenobiella* sp. | A | T | T | C | A | A | C | A | T | C | G | A | G | G | T | C | G | C | A | A |
| MH361300.1 | Neuroptera | *Suhpalacsa longialata* | A | T | T | C | A | A | C | A | T | C | G | A | G | G | T | C | G | C | A | A |

| | | | | | | | | | | | | | | | | | | | | | | |
|---|---|---|---|---|---|---|---|---|---|---|---|---|---|---|---|---|---|---|---|---|---|---|
| NC039948.1 | Neuroptera | *Suhpalacsa longialata* | A | T | T | C | A | A | C | A | T | C | G | A | G | G | T | C | G | C | A | A |
| MK301247.1 | Neuroptera | *Suhpalacsa* sp. | A | T | T | C | A | A | C | A | T | C | G | A | G | G | T | C | G | C | A | A |
| MK408756.1 | Neuroptera | *Thaumatosmylus hainanus* | A | T | T | C | A | A | C | A | T | C | G | A | G | G | T | C | G | C | A | A |
| NC046578.1 | Neuroptera | *Thaumatosmylus hainanus* | A | T | T | C | A | A | C | A | T | C | G | A | G | G | T | C | G | C | A | A |
| KC515397.1 | Neuroptera | *Thyridosmylus langii* | A | T | T | C | A | A | C | A | T | C | G | A | G | G | T | C | G | C | A | A |
| NC021415.1 | Neuroptera | *Thyridosmylus langii* | A | T | T | C | A | A | C | A | T | C | G | A | G | G | T | C | G | C | A | A |
| MH020192.1 | Coleoptera | *Abscondita anceyi* | A | T | C | C | A | A | C | A | T | C | G | A | G | G | T | C | G | C | A | A |
| NC039706.1 | Coleoptera | *Abscondita anceyi* | A | T | C | C | A | A | C | A | T | C | G | A | G | G | T | C | G | C | A | A |
| MK122952.1 | Coleoptera | *Abscondita chinensis* | A | T | C | C | A | A | C | A | T | C | G | A | G | G | T | C | G | C | A | A |
| MK292092.1 | Coleoptera | *Abscondita terminalis* | A | T | C | C | A | A | C | A | T | C | G | A | G | G | T | C | G | C | A | A |
| NC044776.1 | Coleoptera | *Abscondita terminalis* | A | T | C | C | A | A | C | A | T | C | G | A | G | G | T | C | G | C | A | A |
| KT876878.1 | Coleoptera | *Acilius sulcatus* | A | T | C | C | A | A | C | A | T | C | G | A | G | G | T | C | G | C | A | A |
| KX758085.1 | Coleoptera | *Aquatica ficta* | A | T | C | C | A | A | C | A | T | C | G | A | G | G | T | C | G | C | A | A |
| NC035060.1 | Coleoptera | *Aquatica ficta* | A | T | C | C | A | A | C | A | T | C | G | A | G | G | T | C | G | C | A | A |
| NC035755.1 | Coleoptera | *Aquatica lateralis* | A | T | C | C | A | A | C | A | T | C | G | A | G | G | T | C | G | C | A | A |
| KF667531.1 | Coleoptera | *Aquatica leii* | A | T | C | C | A | A | C | A | T | C | G | A | G | G | T | C | G | C | A | A |

| | | | | | | | | | | | | | | | | | | | | | | |
|---|---|---|---|---|---|---|---|---|---|---|---|---|---|---|---|---|---|---|---|---|---|---|
| NC025276.1 | Coleoptera | *Aquatica leii* | A | T | C | C | A | A | C | A | T | C | G | A | G | G | T | C | G | C | A | A |
| KX758086.1 | Coleoptera | *Aquatica wuhana* | A | T | C | C | A | A | C | A | T | C | G | A | G | G | T | C | G | C | A | A |
| NC035061.1 | Coleoptera | *Aquatica wuhana* | A | T | C | C | A | A | C | A | T | C | G | A | G | G | T | C | G | C | A | A |
| KX229747.1 | Coleoptera | *Asymmetricata circumdata* | A | T | C | C | A | A | C | A | T | C | G | A | G | G | T | C | G | C | A | A |
| MK292113.1 | Coleoptera | *Asymmetricata circumdata* | A | T | C | C | A | A | C | A | T | C | G | A | G | G | T | C | G | C | A | A |
| NC032062.1 | Coleoptera | *Asymmetricata circumdata* | A | T | C | C | A | A | C | A | T | C | G | A | G | G | T | C | G | C | A | A |
| KJ922151.1 | Coleoptera | *Bicellonychia lividipennis* | A | T | C | C | A | A | C | A | T | C | G | A | G | G | T | C | G | C | A | A |
| NC030060.1 | Coleoptera | *Bicellonychia lividipennis* | A | T | C | C | A | A | C | A | T | C | G | A | G | G | T | C | G | C | A | A |
| KT876885.1 | Coleoptera | *Colymbetes fuscus* | A | T | C | C | A | A | C | A | T | C | G | A | G | G | T | C | G | C | A | A |
| MK292114.1 | Coleoptera | *Curtos bilineatus* | A | T | C | C | A | A | C | A | T | C | G | A | G | G | T | C | G | C | A | A |
| NC044789.1 | Coleoptera | *Curtos bilineatus* | A | T | C | C | A | A | C | A | T | C | G | A | G | G | T | C | G | C | A | A |
| MK609965.1 | Coleoptera | *Curtos costipennis* | A | T | C | C | A | A | C | A | T | C | G | A | G | G | T | C | G | C | A | A |
| MH651351.1 | Coleoptera | *Diaphanes citrinus* | A | T | C | C | A | A | C | A | T | C | G | A | G | G | T | C | G | C | A | A |
| MK292116.1 | Coleoptera | *Diaphanes mendax* | A | T | C | C | A | A | C | A | T | C | G | A | G | G | T | C | G | C | A | A |
| NC044791.1 | Coleoptera | *Diaphanes mendax* | A | T | C | C | A | A | C | A | T | C | G | A | G | G | T | C | G | C | A | A |
| MK292094.1 | Coleoptera | *Diaphanes nubilus* | A | T | C | C | A | A | C | A | T | C | G | A | G | G | T | C | G | C | A | A |
| NC044787.1 | Coleoptera | *Diaphanes nubilus* | A | T | C | C | A | A | C | A | T | C | G | A | G | G | T | C | G | C | A | A |

| | | | | | | | | | | | | | | | | | | | | | | |
|---|---|---|---|---|---|---|---|---|---|---|---|---|---|---|---|---|---|---|---|---|---|---|
| MK292118.1 | Coleoptera | *Diaphanes pectinealis* | A | T | C | C | A | A | C | A | T | C | G | A | G | G | T | C | G | C | A | A |
| NC044793.1 | Coleoptera | *Diaphanes pectinealis* | A | T | C | C | A | A | C | A | T | C | G | A | G | G | T | C | G | C | A | A |
| MK292095.1 | Coleoptera | *Diaphanes* sp. | A | T | C | C | A | A | C | A | T | C | G | A | G | G | T | C | G | C | A | A |
| MG242622.1 | Coleoptera | *Ellychnia corrusca* | A | T | C | C | A | A | C | A | T | C | G | A | G | G | T | C | G | C | A | A |
| MK292112.1 | Coleoptera | *Emeia pseudosauteri* | A | T | C | C | A | A | C | A | T | C | G | A | G | G | T | C | G | C | A | A |
| NC044778.1 | Coleoptera | *Emeia pseudosauteri* | A | T | C | C | A | A | C | A | T | C | G | A | G | G | T | C | G | C | A | A |
| KT876897.1 | Coleoptera | *Hydroporus erythrocephalus* | A | T | C | C | A | A | C | A | T | C | G | A | G | G | T | C | G | C | A | A |
| KT876896.1 | Coleoptera | *Hydroporus obscurus* | A | T | C | C | A | A | C | A | T | C | G | A | G | G | T | C | G | C | A | A |
| KT876901.1 | Coleoptera | *Hygrotus confluens* | A | T | C | C | A | A | C | A | T | C | G | A | G | G | T | C | G | C | A | A |
| KT876900.1 | Coleoptera | *Hygrotus impressopunctatus* | A | T | C | C | A | A | C | A | T | C | G | A | G | G | T | C | G | C | A | A |
| KT876899.1 | Coleoptera | *Hygrotus inaequalis* | A | T | C | C | A | A | C | A | T | C | G | A | G | G | T | C | G | C | A | A |
| KX087299.1 | Coleoptera | *Hygrotus nigrolineatus* | A | T | C | C | A | A | C | A | T | C | G | A | G | G | T | C | G | C | A | A |
| MH427718.1 | Coleoptera | *Inflata indica* | A | T | C | C | A | A | C | A | T | C | G | A | G | G | T | C | G | C | A | A |
| NC039700.1 | Coleoptera | *Inflata indica* | A | T | C | C | A | A | C | A | T | C | G | A | G | G | T | C | G | C | A | A |

| | | | | | | | | | | | | | | | | | | | | | | |
|---|---|---|---|---|---|---|---|---|---|---|---|---|---|---|---|---|---|---|---|---|---|---|
| MK292091.1 | Coleoptera | *Lamprigera yunnana* | A | T | C | C | A | A | C | A | T | C | G | A | G | G | T | C | G | C | A | A |
| NC044786.1 | Coleoptera | *Lamprigera yunnana* | A | T | C | C | A | A | C | A | T | C | G | A | G | G | T | C | G | C | A | A |
| KX087302.1 | Coleoptera | *Lampyris noctiluca* | A | T | C | C | A | A | C | A | T | C | G | A | G | G | T | C | G | C | A | A |
| MG912994.1 | Coleoptera | *Limbodessus palmulaoides* | A | T | C | C | A | A | C | A | T | C | G | A | G | G | T | C | G | C | A | A |
| NC037749.1 | Coleoptera | *Limbodessus palmulaoides* | A | T | C | C | A | A | C | A | T | C | G | A | G | G | T | C | G | C | A | A |
| JQ034416.1 | Coleoptera | *Limnichidae* sp. | A | T | C | C | A | A | C | A | T | C | G | A | G | G | T | C | G | C | A | A |
| KT876902.1 | Coleoptera | *Liopterus haemorrhoidalis* | A | T | C | C | A | A | C | A | T | C | G | A | G | G | T | C | G | C | A | A |
| NC022472.1 | Coleoptera | *Luciola cruciata* | A | T | C | C | A | A | C | A | T | C | G | A | G | G | T | C | G | C | A | A |
| MG770613.1 | Coleoptera | *Luciola curtithorax* | A | T | C | C | A | A | C | A | T | C | G | A | G | G | T | C | G | C | A | A |
| NC038225.1 | Coleoptera | *Luciola curtithorax* | A | T | C | C | A | A | C | A | T | C | G | A | G | G | T | C | G | C | A | A |
| KP313820.1 | Coleoptera | *Luciola substriata* | A | T | C | C | A | A | C | A | T | C | G | A | G | G | T | C | G | C | A | A |
| NC027176.1 | Coleoptera | *Luciola substriata* | A | T | C | C | A | A | C | A | T | C | G | A | G | G | T | C | G | C | A | A |
| FJ859901.1 | Coleoptera | *Macrogyrus oblongus* | A | T | C | C | A | A | C | A | T | C | G | A | G | G | T | C | G | C | A | A |
| NC013249.1 | Coleoptera | *Macrogyrus oblongus* | A | T | C | C | A | A | C | A | T | C | G | A | G | G | T | C | G | C | A | A |
| MG912995.1 | Coleoptera | *Paroster* | A | T | C | C | A | A | C | A | T | C | G | A | G | G | T | C | G | C | A | A |

| | | | | | | | | | | | | | | | | | | | | | | |
|---|---|---|---|---|---|---|---|---|---|---|---|---|---|---|---|---|---|---|---|---|---|---|
| | | *macrosturtensis* | | | | | | | | | | | | | | | | | | | | |
| NC037750.1 | Coleoptera | *Paroster macrosturtensis* | A | T | C | C | A | A | C | A | T | C | G | A | G | G | T | C | G | C | A | A |
| MG912996.1 | Coleoptera | *Paroster mesosturtensis* | A | T | C | C | A | A | C | A | T | C | G | A | G | G | T | C | G | C | A | A |
| NC037751.1 | Coleoptera | *Paroster mesosturtensis* | A | T | C | C | A | A | C | A | T | C | G | A | G | G | T | C | G | C | A | A |
| MG912997.1 | Coleoptera | *Paroster microsturtensis* | A | T | C | C | A | A | C | A | T | C | G | A | G | G | T | C | G | C | A | A |
| NC037752.1 | Coleoptera | *Paroster microsturtensis* | A | T | C | C | A | A | C | A | T | C | G | A | G | G | T | C | G | C | A | A |
| KY778696.1 | Coleoptera | *Photinus pyralis* | A | T | C | C | A | A | C | A | T | C | G | A | G | G | T | C | G | C | A | A |
| MF686051.1 | Coleoptera | *Pteroptyx maipo* | A | T | C | C | A | A | C | A | T | C | G | A | G | G | T | C | G | C | A | A |
| NC036353.1 | Coleoptera | *Pteroptyx maipo* | A | T | C | C | A | A | C | A | T | C | G | A | G | G | T | C | G | C | A | A |
| MK292115.1 | Coleoptera | *Pyrocoelia praetexta* | A | T | C | C | A | A | C | A | T | C | G | A | G | G | T | C | G | C | A | A |
| NC044790.1 | Coleoptera | *Pyrocoelia praetexta* | A | T | C | C | A | A | C | A | T | C | G | A | G | G | T | C | G | C | A | A |
| AF452048.1 | Coleoptera | *Pyrocoelia rufa* | A | T | C | C | A | A | C | A | T | C | G | A | G | G | T | C | G | C | A | A |
| NC003970.1 | Coleoptera | *Pyrocoelia rufa* | A | T | C | C | A | A | C | A | T | C | G | A | G | G | T | C | G | C | A | A |
| MK292117.1 | Coleoptera | *Pyrocoelia thibetana* | A | T | C | C | A | A | C | A | T | C | G | A | G | G | T | C | G | C | A | A |
| NC044792.1 | Coleoptera | *Pyrocoelia thibetana* | A | T | C | C | A | A | C | A | T | C | G | A | G | G | T | C | G | C | A | A |
| MK292111.1 | Coleoptera | *Vesta saturnalis* | A | T | C | C | A | A | C | A | T | C | G | A | G | G | T | C | G | C | A | A |
| NC044788.1 | Coleoptera | *Vesta saturnalis* | A | T | C | C | A | A | C | A | T | C | G | A | G | G | T | C | G | C | A | A |

| | | | | | | | | | | | | | | | | | | | | | | |
|---|---|---|---|---|---|---|---|---|---|---|---|---|---|---|---|---|---|---|---|---|---|---|
| MF680449.1 | Trichoptera | *Anabolia bimaculata* | A | T | C | C | A | A | C | A | T | C | G | A | G | G | T | C | G | C | A | A |
| NC036004.1 | Trichoptera | *Anabolia bimaculata* | A | T | C | C | A | A | C | A | T | C | G | A | G | G | T | C | G | C | A | A |
| KF756944.1 | Trichoptera | *Apatania* sp. | A | T | C | C | A | A | C | A | T | C | G | A | G | G | T | C | G | C | A | A |
| MT410856.1 | Trichoptera | *Beraea pullata* | A | T | C | C | A | A | C | A | T | C | G | A | G | G | T | C | G | C | A | A |
| MG669126.1 | Trichoptera | *Cheumatopsyche analis* | A | T | C | C | A | A | C | A | T | C | G | A | G | G | T | C | G | C | A | A |
| NC036955.1 | Trichoptera | *Cheumatopsyche analis* | A | T | C | C | A | A | C | A | T | C | G | A | G | G | T | C | G | C | A | A |
| KX385010.1 | Trichoptera | *Cheumatopsyche brevilineata* | A | T | C | C | A | A | C | A | T | C | G | A | G | G | T | C | G | C | A | A |
| NC043769.1 | Trichoptera | *Cheumatopsyche brevilineata* | A | T | C | C | A | A | C | A | T | C | G | A | G | G | T | C | G | C | A | A |
| MG669125.1 | Trichoptera | *Cheumatopsyche campyla* | A | T | C | C | A | A | C | A | T | C | G | A | G | G | T | C | G | C | A | A |
| NC036954.1 | Trichoptera | *Cheumatopsyche campyla* | A | T | C | C | A | A | C | A | T | C | G | A | G | G | T | C | G | C | A | A |
| MG669123.1 | Trichoptera | *Cheumatopsyche speciosa* | A | T | C | C | A | A | C | A | T | C | G | A | G | G | T | C | G | C | A | A |
| NC036952.1 | Trichoptera | *Cheumatopsyche speciosa* | A | T | C | C | A | A | C | A | T | C | G | A | G | G | T | C | G | C | A | A |
| KF717094.1 | Trichoptera | *Eubasilissa regina* | A | T | C | C | A | A | C | A | T | C | G | A | G | G | T | C | G | C | A | A |
| KF756943.1 | Trichoptera | *Eubasilissa regina* | A | T | C | C | A | A | C | A | T | C | G | A | G | G | T | C | G | C | A | A |

| | | | | | | | | | | | | | | | | | | | | | | |
|---|---|---|---|---|---|---|---|---|---|---|---|---|---|---|---|---|---|---|---|---|---|---|
| NC023374.1 | Trichoptera | *Eubasilissa regina* | A | T | C | C | A | A | C | A | T | C | G | A | G | G | T | C | G | C | A | A |
| KX385011.1 | Trichoptera | *Hydatophylax nigrovittatus* | A | T | C | C | A | A | C | A | T | C | G | A | G | G | T | C | G | C | A | A |
| NC043770.1 | Trichoptera | *Hydatophylax nigrovittatus* | A | T | C | C | A | A | C | A | T | C | G | A | G | G | T | C | G | C | A | A |
| KF717095.1 | Trichoptera | *Hydromanicus wulaianus* | A | T | C | C | A | A | C | A | T | C | G | A | G | G | T | C | G | C | A | A |
| NC036156.1 | Trichoptera | *Hydromanicus wulaianus* | A | T | C | C | A | A | C | A | T | C | G | A | G | G | T | C | G | C | A | A |
| MT628580.1 | Trichoptera | *Hydropsyche angustipennis* | A | T | C | C | A | A | C | A | T | C | G | A | G | G | T | C | G | C | A | A |
| MG669122.1 | Trichoptera | *Hydropsyche orris* | A | T | C | C | A | A | C | A | T | C | G | A | G | G | T | C | G | C | A | A |
| NC036951.1 | Trichoptera | *Hydropsyche orris* | A | T | C | C | A | A | C | A | T | C | G | A | G | G | T | C | G | C | A | A |
| KT876876.1 | Trichoptera | *Hydropsyche pellucidula* | A | T | C | C | A | A | C | A | T | C | G | A | G | G | T | C | G | C | A | A |
| NC029246.1 | Trichoptera | *Hydropsyche pellucidula* | A | T | C | C | A | A | C | A | T | C | G | A | G | G | T | C | G | C | A | A |
| MG669121.1 | Trichoptera | *Hydropsyche simulans* | A | T | C | C | A | A | C | A | T | C | G | A | G | G | T | C | G | C | A | A |
| NC036950.1 | Trichoptera | *Hydropsyche simulans* | A | T | C | C | A | A | C | A | T | C | G | A | G | G | T | C | G | C | A | A |
| MK077681.1 | Trichoptera | *Limnephilus* | A | T | C | C | A | A | C | A | T | C | G | A | G | G | T | C | G | C | A | A |

| | | | | | | | | | | | | | | | | | | | | | | |
|---|---|---|---|---|---|---|---|---|---|---|---|---|---|---|---|---|---|---|---|---|---|---|
| | | *hyalinus* | | | | | | | | | | | | | | | | | | | | |
| NC044710.1 | Trichoptera | *Limnephilus hyalinus* | A | T | C | C | A | A | C | A | T | C | G | A | G | G | T | C | G | C | A | A |
| MG980616.1 | Trichoptera | *Phryganea cinerea* | A | T | C | C | A | A | C | A | T | C | G | A | G | G | T | C | G | C | A | A |
| NC039714.1 | Trichoptera | *Phryganea cinerea* | A | T | C | C | A | A | C | A | T | C | G | A | G | G | T | C | G | C | A | A |
| KX385012.1 | Trichoptera | *Phryganopsyche latipennis* | A | T | C | C | A | A | C | A | T | C | G | A | G | G | T | C | G | C | A | A |
| NC043771.1 | Trichoptera | *Phryganopsyche latipennis* | A | T | C | C | A | A | C | A | T | C | G | A | G | G | T | C | G | C | A | A |
| MG669124.1 | Trichoptera | *Potamyia flava* | A | T | C | C | A | A | C | A | T | C | G | A | G | G | T | C | G | C | A | C |
| NC036953.1 | Trichoptera | *Potamyia flava* | A | T | C | C | A | A | C | A | T | C | G | A | G | G | T | C | G | C | A | C |
| KP455290.1 | Trichoptera | *Sericostoma personatum* | A | T | C | C | A | A | C | A | T | C | G | A | G | G | T | C | G | C | A | A |
| KP455291.1 | Trichoptera | *Thremma gallicum* | A | T | C | C | A | A | C | A | T | C | G | A | G | G | T | C | G | C | A | A |
| AY072044.1 | Diptera | *Aedes albopictus* | A | T | C | C | A | A | C | A | T | C | G | A | G | G | T | C | G | C | A | A |
| NC030249.1 | Diptera | *Anopheles bellator* | A | T | C | C | A | A | C | A | T | C | G | A | G | G | T | C | G | C | A | A |
| NC030717.1 | Diptera | *Anopheles janconnae* | A | T | C | C | A | A | C | A | T | C | G | A | G | G | T | C | G | C | A | A |
| HQ335345.1 | Diptera | *Anopheles oryzalimnetes* | A | T | C | C | A | A | C | A | T | C | G | A | G | G | T | C | G | C | A | A |
| MF381722.1 | Diptera | *Culex bilineatus* | A | T | C | C | A | A | C | A | T | C | G | A | G | G | T | C | G | C | A | A |
| MF381718.1 | Diptera | *Culex lygrus* | A | T | C | C | A | A | C | A | T | C | G | A | G | G | T | C | G | C | A | A |

| | | | | | | | | | | | | | | | | | | | | | | |
|---|---|---|---|---|---|---|---|---|---|---|---|---|---|---|---|---|---|---|---|---|---|---|
| MH316119.1 | Diptera | *Lutzia halifaxii* | A | T | C | C | A | A | C | A | T | C | G | A | G | G | T | C | G | C | A | A |
| MN626442.1 | Diptera | *Ochlerotatus taeniorhynchus* | A | T | C | C | A | A | C | A | T | C | G | A | G | G | T | C | G | C | A | A |
| KP721463.1 | Diptera | *Ochlerotatus vigilax* | A | T | C | C | A | A | C | A | T | C | G | A | G | G | T | C | G | C | A | A |
| MF957171.1 | Diptera | *Sabethes belisarioi* | A | T | C | C | A | A | C | A | T | C | G | A | G | G | T | C | G | C | A | A |
| KP793690.1 | Diptera | *Simulium aureohirtum* | A | T | C | C | A | A | C | A | T | C | G | A | G | G | T | C | G | C | A | A |
| NC029753.1 | Diptera | *Simulium aureohirtum* | A | T | C | C | A | A | C | A | T | C | G | A | G | G | T | C | G | C | A | A |
| MH729190.1 | Diptera | *Simulium maculatum* | A | T | C | C | A | A | C | A | T | C | G | A | G | G | T | C | G | C | A | A |
| NC040120.1 | Diptera | *Simulium maculatum* | A | T | C | C | A | A | C | A | T | C | G | A | G | G | T | C | G | C | A | A |
| KU252587.1 | Diptera | *Simulium variegatum* | A | T | C | C | A | A | C | A | T | C | G | A | G | G | T | C | G | C | A | A |
| NC033348.1 | Diptera | *Simulium variegatum* | A | T | C | C | A | A | C | A | T | C | G | A | G | G | T | C | G | C | A | A |
| JN861743.1 | Diptera | *Tipula abdominalis* | A | T | C | C | A | A | C | A | T | C | G | A | G | G | T | C | G | C | A | A |
| KT970065.1 | Diptera | *Tipula cockerelliana* | A | T | C | C | A | A | C | A | T | C | G | A | G | G | T | C | G | C | A | A |
| NC030520.1 | Diptera | *Tipula cockerelliana* | A | T | C | C | A | A | C | A | T | C | G | A | G | G | T | C | G | C | A | A |
| MK575491.1 | Diptera | *Uranotaenia geometrica* | A | T | C | C | A | A | C | A | T | C | G | A | G | G | T | C | G | C | A | A |

| | | | | | | | | | | | | | | | | | | | | |
|---|---|---|---|---|---|---|---|---|---|---|---|---|---|---|---|---|---|---|---|---|
| A | 96.2 | 0.0 | 0.0 | 0.0 | 100.0 | 100.0 | 0.0 | 100.0 | 0.0 | 0.0 | 0.0 | 99.6 | 0.0 | 0.0 | 0.0 | 0.0 | 2.3 | 0.0 | 100.0 | 99.6 |
| T | 0.0 | 99.3 | 32.5 | 0.0 | 0.0 | 0.0 | 0.0 | 0.0 | 100.0 | 0.0 | 0.0 | 0.0 | 0.0 | 0.0 | 100.0 | 2.0 | 0.0 | 1.1 | 0.0 | 0.0 |
| G | 3.8 | 0.0 | 0.0 | 0.0 | 0.0 | 0.0 | 0.0 | 0.0 | 0.0 | 0.0 | 100.0 | 0.4 | 100.0 | 100.0 | 0.0 | 0.0 | 97.7 | 0.0 | 0.0 | 0.0 |
| C | 0.0 | 0.7 | 67.5 | 100.0 | 0.0 | 0.0 | 100.0 | 0.0 | 0.0 | 100.0 | 0.0 | 0.0 | 0.0 | 0.0 | 0.0 | 98.0 | 0.0 | 98.9 | 0.0 | 0.4 |

Table S4. Nucleotide sequences of universal primer (AQdb-16S_F) and homologous region of 557 sequences of insects used for development in this study.

| | | | | | | | | | | | | | | | | | | | | | |
|---|---|---|---|---|---|---|---|---|---|---|---|---|---|---|---|---|---|---|---|---|---|
| AQdb-16S_F | | | | T | R | A | C | Y | G | T | R | C | A | A | A | G | G | T | A | G | C |
| | | | A | 0 | 93 | 557 | 0 | 0 | 0 | 0 | 77 | 0 | 515 | 555 | 557 | 0 | 0 | 0 | 557 | 0 | 0 |
| | | | T | 555 | 0 | 0 | 2 | 273 | 0 | 557 | 0 | 0 | 11 | 0 | 0 | 2 | 0 | 557 | 0 | 0 | 0 |
| | | | G | 0 | 464 | 0 | 0 | 0 | 557 | 0 | 480 | 0 | 31 | 2 | 0 | 555 | 557 | 0 | 0 | 557 | 0 |
| | | | C | 0 | 0 | 0 | 555 | 284 | 0 | 0 | 0 | 557 | 0 | 0 | 0 | 0 | 0 | 0 | 0 | 0 | 557 |
| GenBank Acc. No. | Order | Species | | | | | | | | | | | | | | | | | | | |
| MN481418.1 | Diplura | *Campodea augens* | | T | A | A | C | T | G | T | A | C | A | A | A | G | G | T | A | G | C |
| DQ529236.1 | Diplura | *Campodea fragilis* | | T | A | A | C | T | G | T | A | C | A | A | A | G | G | T | A | G | C |
| NC008233.1 | Diplura | *Campodea fragilis* | | T | A | A | C | T | G | T | A | C | A | A | A | G | G | T | A | G | C |
| DQ529237.1 | Diplura | *Campodea lubbocki* | | T | A | A | C | T | G | T | A | C | A | A | A | G | G | T | A | G | C |
| NC008234.1 | Diplura | *Campodea lubbocki* | | T | A | A | C | T | G | T | A | C | A | A | A | G | G | T | A | G | C |
| AY771989.1 | Diplura | *Japyx solifugus* | | T | G | A | C | T | G | T | A | C | A | A | A | G | G | T | A | G | C |
| NC007214.1 | Diplura | *Japyx solifugus* | | T | G | A | C | T | G | T | A | C | A | A | A | G | G | T | A | G | C |
| JN990601.1 | Diplura | *Lepidocampa weberi* | | T | G | A | C | T | G | T | G | C | A | A | A | G | G | T | A | G | C |
| NC022675.1 | Diplura | *Lepidocampa weberi* | | T | G | A | C | T | G | T | G | C | A | A | A | G | G | T | A | G | C |
| JN990600.1 | Diplura | *Occasjapyx japonicus* | | T | G | A | C | T | G | T | A | C | A | A | A | G | G | T | A | G | C |
| NC022674.1 | Diplura | *Occasjapyx japonicus* | | T | G | A | C | T | G | T | A | C | A | A | A | G | G | T | A | G | C |
| JN990598.1 | Diplura | *Octostigma sinensis* | | T | G | A | C | T | G | T | A | C | T | A | A | G | G | T | A | G | C |
| NC022672.1 | Diplura | *Octostigma sinensis* | | T | G | A | C | T | G | T | A | C | T | A | A | G | G | T | A | G | C |
| JN990599.1 | Diplura | *Parajapyx emeryanus* | | T | G | A | C | C | G | T | A | C | A | A | A | G | G | T | A | G | C |

| | | | | | | | | | | | | | | | | | | | | |
|---|---|---|---|---|---|---|---|---|---|---|---|---|---|---|---|---|---|---|---|---|
| NC022673.1 | Diplura | *Parajapyx emeryanus* | T | G | A | C | C | G | T | A | C | A | A | A | G | G | T | A | G | C |
| AY072044.1 | Diptera | *Aedes albopictus* | T | G | A | C | T | G | T | G | C | G | A | A | G | G | T | A | G | C |
| NC030249.1 | Diptera | *Anopheles bellator* | T | G | A | C | C | G | T | G | C | G | A | A | G | G | T | A | G | C |
| NC030717.1 | Diptera | *Anopheles janconnae* | T | G | A | C | C | G | T | G | C | G | A | A | G | G | T | A | G | C |
| HQ335345.1 | Diptera | *Anopheles oryzalimnetes* | T | G | A | C | C | G | T | G | C | G | A | A | G | G | T | A | G | C |
| MF381722.1 | Diptera | *Culex bilineatus* | T | G | A | C | T | G | T | G | C | G | A | A | G | G | T | A | G | C |
| MF381718.1 | Diptera | *Culex lygrus* | T | G | A | C | T | G | T | G | C | G | A | A | G | G | T | A | G | C |
| MH316119.1 | Diptera | *Lutzia halifaxii* | T | G | A | C | T | G | T | G | C | G | A | A | G | G | T | A | G | C |
| MN626442.1 | Diptera | *Ochlerotatus taeniorhynchus* | T | G | A | C | T | G | T | G | C | G | A | A | G | G | T | A | G | C |
| KP721463.1 | Diptera | *Ochlerotatus vigilax* | T | G | A | C | T | G | T | G | C | G | A | A | G | G | T | A | G | C |
| MF957171.1 | Diptera | *Sabethes belisarioi* | T | G | A | C | T | G | T | G | C | A | A | A | G | G | T | A | G | C |
| KP793690.1 | Diptera | *Simulium aureohirtum* | T | G | A | C | T | G | T | G | C | A | A | A | G | G | T | A | G | C |
| NC029753.1 | Diptera | *Simulium aureohirtum* | T | G | A | C | T | G | T | G | C | A | A | A | G | G | T | A | G | C |
| MH729190.1 | Diptera | *Simulium maculatum* | T | G | A | C | T | G | T | G | C | A | A | A | G | G | T | A | G | C |
| NC040120.1 | Diptera | *Simulium maculatum* | T | G | A | C | T | G | T | G | C | A | A | A | G | G | T | A | G | C |
| KU252587.1 | Diptera | *Simulium variegatum* | T | G | A | C | T | G | T | G | C | A | A | A | G | G | T | A | G | C |
| NC033348.1 | Diptera | *Simulium variegatum* | T | G | A | C | T | G | T | G | C | A | A | A | G | G | T | A | G | C |
| JN861743.1 | Diptera | *Tipula abdominalis* | T | G | A | C | C | G | T | G | C | A | A | A | G | G | T | A | G | C |
| KT970065.1 | Diptera | *Tipula cockerelliana* | T | G | A | C | C | G | T | G | C | A | A | A | G | G | T | A | G | C |
| NC030520.1 | Diptera | *Tipula cockerelliana* | T | G | A | C | C | G | T | G | C | A | A | A | G | G | T | A | G | C |
| MK575491.1 | Diptera | *Uranotaenia geometrica* | T | G | A | C | C | G | T | G | C | G | A | A | G | G | T | A | G | C |
| KJ754500.1 | Archaeognatha | *Allopsontus* sp. | T | G | A | C | C | G | T | G | C | A | A | A | G | G | T | A | G | C |

| | | | | | | | | | | | | | | | | | | | | |
|---|---|---|---|---|---|---|---|---|---|---|---|---|---|---|---|---|---|---|---|---|
| KJ754501.1 | Archaeognatha | *Allopsontus* sp. | T | G | A | C | C | G | T | G | C | A | A | A | G | G | T | A | G | C |
| AY793551.1 | Archaeognatha | *Nesomachilis australica* | T | G | A | C | C | G | T | G | C | A | A | A | G | G | T | A | G | C |
| NC006895.1 | Archaeognatha | *Nesomachilis australica* | T | G | A | C | C | G | T | G | C | A | A | A | G | G | T | A | G | C |
| KJ754502.1 | Archaeognatha | *Pedetontinus luanchuanensis* | T | G | A | C | C | G | T | G | C | A | A | A | G | G | T | A | G | C |
| EU621793.1 | Archaeognatha | *Pedetontus silvestrii* | T | G | A | C | C | G | T | G | C | A | A | A | G | G | T | A | G | C |
| NC011717.1 | Archaeognatha | *Pedetontus silvestrii* | T | G | A | C | C | G | T | G | C | A | A | A | G | G | T | A | G | C |
| KJ754503.1 | Archaeognatha | *Petrobiellus* sp. | T | G | A | C | C | G | T | G | C | A | A | A | G | G | T | A | G | C |
| KJ754504.1 | Archaeognatha | *Petrobiellus* sp. | T | G | A | C | C | G | T | G | C | A | A | A | G | G | T | A | G | C |
| AY956355.1 | Archaeognatha | *Petrobius brevistylis* | T | A | A | C | C | G | T | G | C | A | G | A | T | G | T | A | G | C |
| NC007688.1 | Archaeognatha | *Petrobius brevistylis* | T | A | A | C | C | G | T | G | C | A | G | A | T | G | T | A | G | C |
| JX308221.1 | Archaeognatha | *Songmachilis xinxiangensis* | T | G | A | C | C | G | T | G | C | A | A | A | G | G | T | A | G | C |
| NC021384.1 | Archaeognatha | *Songmachilis xinxiangensis* | T | G | A | C | C | G | T | G | C | A | A | A | G | G | T | A | G | C |
| EU016193.1 | Archaeognatha | *Trigoniophthalmus alternatus* | T | G | A | C | T | G | T | G | C | A | A | A | G | G | T | A | G | C |
| NC010532.1 | Archaeognatha | *Trigoniophthalmus alternatus* | T | G | A | C | T | G | T | G | C | A | A | A | G | G | T | A | G | C |
| EU084035.1 | Zygentoma | *Atelura formicaria* | T | G | A | C | C | G | T | G | C | A | A | A | G | G | T | A | G | C |
| NC011197.1 | Zygentoma | *Atelura formicaria* | T | G | A | C | C | G | T | G | C | A | A | A | G | G | T | A | G | C |
| MK301436.1 | Zygentoma | *Ctenolepisma villosa* | T | G | A | C | C | G | T | G | C | G | A | A | G | G | T | A | G | C |
| NC046478.1 | Zygentoma | *Ctenolepisma villosa* | T | G | A | C | C | G | T | G | C | G | A | A | G | G | T | A | G | C |
| MT108230.1 | Zygentoma | *Lepisma saccharina* | T | G | A | C | C | G | T | G | C | A | A | A | G | G | T | A | G | C |
| NC047445.1 | Zygentoma | *Lepisma saccharina* | T | G | A | C | C | G | T | G | C | A | A | A | G | G | T | A | G | C |
| AY639935.1 | Zygentoma | *Thermobia domestica* | T | G | A | C | C | G | T | G | C | G | A | A | G | G | T | A | G | C |
| NC006080.1 | Zygentoma | *Thermobia domestica* | T | G | A | C | C | G | T | G | C | G | A | A | G | G | T | A | G | C |

| | | | | | | | | | | | | | | | | | | | | |
|---|---|---|---|---|---|---|---|---|---|---|---|---|---|---|---|---|---|---|---|---|
| MK951657.1 | Zygentoma | *Thermobia* sp. | T | G | A | C | C | G | T | G | C | A | A | A | G | G | T | A | G | C |
| AY191994.1 | Zygentoma | *Tricholepidion gertschi* | T | G | A | C | C | G | T | G | C | A | A | A | G | G | T | A | G | C |
| NC005437.1 | Zygentoma | *Tricholepidion gertschi* | T | G | A | C | C | G | T | G | C | A | A | A | G | G | T | A | G | C |
| GU479735.1 | Ephemeroptera | *Alainites yixiani* | T | G | A | C | T | G | T | G | C | A | A | A | G | G | T | A | G | C |
| GU936204.1 | Ephemeroptera | *Baetis* sp. | T | G | A | C | T | G | T | G | C | A | A | A | G | G | T | A | G | C |
| MT628575.1 | Ephemeroptera | *Caenis robusta* | T | G | A | C | C | G | T | G | C | A | A | A | G | G | T | A | G | C |
| GQ502451.1 | Ephemeroptera | *Caenis* sp. | T | G | A | C | C | G | T | G | C | A | A | A | G | G | T | A | G | C |
| MG910499.1 | Ephemeroptera | *Caenis* sp. | T | G | A | C | C | G | T | G | C | A | A | A | G | G | T | A | G | C |
| MN356096.1 | Ephemeroptera | *Caenis* sp. | T | G | A | C | C | G | T | G | C | A | A | A | G | G | T | A | G | C |
| MN807287.1 | Ephemeroptera | *Choroterpides apiculata* | T | G | A | C | C | G | T | G | C | A | A | A | G | G | T | A | G | C |
| MG870104.1 | Ephemeroptera | *Epeorus herklotsi* | T | G | A | C | C | G | T | G | C | A | A | A | G | G | T | A | G | C |
| MH752075.1 | Ephemeroptera | *Epeorus herklotsi* | T | G | A | C | C | G | T | G | C | A | A | A | G | G | T | A | G | C |
| NC039612.1 | Ephemeroptera | *Epeorus herklotsi* | T | G | A | C | C | G | T | G | C | A | A | A | G | G | T | A | G | C |
| KJ493406.1 | Ephemeroptera | *Epeorus* sp. | T | G | A | C | C | G | T | G | C | A | A | A | G | G | T | A | G | C |
| KM244708.1 | Ephemeroptera | *Epeorus* sp. | T | G | A | C | C | G | T | G | C | A | A | A | G | G | T | A | G | C |
| MK951674.1 | Ephemeroptera | *Epeorus* sp. | T | G | A | C | C | G | T | G | C | A | A | A | G | G | T | A | G | C |
| EU591678.1 | Ephemeroptera | *Ephemera orientalis* | T | G | A | C | C | G | T | G | C | A | A | A | G | G | T | A | G | C |
| NC012645.1 | Ephemeroptera | *Ephemera orientalis* | T | G | A | C | C | G | T | G | C | A | A | A | G | G | T | A | G | C |
| MK951659.1 | Ephemeroptera | *Ephemera* sp. | T | G | A | C | C | G | T | G | C | A | A | A | G | G | T | A | G | C |
| KM244691.1 | Ephemeroptera | *Ephemerella* sp. | T | G | A | C | C | G | T | G | C | A | A | A | G | G | T | A | G | C |
| MT274127.1 | Ephemeroptera | *Ephemerella* sp. | T | G | A | C | C | G | T | G | C | A | A | A | G | G | T | A | G | C |
| NC050279.1 | Ephemeroptera | *Ephemerella* sp. | T | G | A | C | C | G | T | G | C | A | A | A | G | G | T | A | G | C |

| | | | | | | | | | | | | | | | | | | | | |
|---|---|---|---|---|---|---|---|---|---|---|---|---|---|---|---|---|---|---|---|---|
| GU936203.1 | Ephemeroptera | *Habrophlebiodes zijinensis* | T | G | A | C | T | G | T | G | C | A | A | A | G | G | T | A | G | C |
| HM143892.1 | Ephemeroptera | *Isonychia ignota* | T | G | A | C | C | G | T | G | C | A | A | A | G | G | T | A | G | C |
| MH119135.1 | Ephemeroptera | *Isonychia kiangsinensis* | T | G | A | C | C | G | T | G | C | A | A | A | G | G | T | A | G | C |
| NC042163.1 | Ephemeroptera | *Isonychia kiangsinensis* | T | G | A | C | C | G | T | G | C | A | A | A | G | G | T | A | G | C |
| MK951658.1 | Ephemeroptera | *Isonychia* sp. | T | G | A | C | C | G | T | G | C | A | A | A | G | G | T | A | G | C |
| HM004123.1 | Ephemeroptera | *Paegniodes cupulatus* | T | G | A | C | C | G | T | G | C | A | A | A | G | G | T | A | G | C |
| MK951673.1 | Ephemeroptera | *Parafronurus* sp. | T | G | A | C | C | G | T | G | C | A | A | A | G | G | T | A | G | C |
| EU349015.1 | Ephemeroptera | *Parafronurus youi* | T | G | A | C | C | G | T | G | C | A | A | A | G | G | T | A | G | C |
| NC011359.1 | Ephemeroptera | *Parafronurus youi* | T | G | A | C | C | G | T | G | C | A | A | A | G | G | T | A | G | C |
| KM244674.1 | Ephemeroptera | *Potamanthus* sp. | T | G | A | C | C | G | T | G | C | A | A | A | G | G | T | A | G | C |
| MK951670.1 | Ephemeroptera | *Rhithrogena* sp. | T | G | A | C | C | G | T | G | C | A | A | A | G | G | T | A | G | C |
| MT274128.1 | Ephemeroptera | *Serratella* sp. | T | G | A | C | C | G | T | G | C | A | A | A | G | G | T | A | G | C |
| MT274129.1 | Ephemeroptera | *Serratella* sp. | T | G | A | C | C | G | T | G | C | A | A | A | G | G | T | A | G | C |
| NC050280.1 | Ephemeroptera | *Serratella* sp. | T | G | A | C | C | G | T | G | C | A | A | A | G | G | T | A | G | C |
| NC050281.1 | Ephemeroptera | *Serratella* sp. | T | G | A | C | C | G | T | G | C | A | A | A | G | G | T | A | G | C |
| MT274130.1 | Ephemeroptera | *Serratella zapekinae* | T | G | A | C | C | G | T | G | C | A | A | A | G | G | T | A | G | C |
| NC050282.1 | Ephemeroptera | *Serratella zapekinae* | T | G | A | C | C | G | T | G | C | A | A | A | G | G | T | A | G | C |
| FJ606783.1 | Ephemeroptera | *Siphlonurus immanis* | T | G | A | C | C | G | T | G | C | A | A | A | G | G | T | A | G | C |
| NC013822.1 | Ephemeroptera | *Siphlonurus immanis* | T | G | A | C | C | G | T | G | C | A | A | A | G | G | T | A | G | C |
| KM244684.1 | Ephemeroptera | *Siphlonurus* sp. | T | G | A | C | C | G | T | G | C | A | A | A | G | G | T | A | G | C |
| HQ875717.1 | Ephemeroptera | *Siphluriscus chinensis* | T | G | A | C | C | G | T | G | C | A | A | A | G | G | T | A | G | C |
| MT274131.1 | Ephemeroptera | *Torleya grandiforceps* | T | G | A | C | C | G | T | G | C | A | A | A | G | G | T | A | G | C |

| | | | | | | | | | | | | | | | | | | | | |
|---|---|---|---|---|---|---|---|---|---|---|---|---|---|---|---|---|---|---|---|---|
| NC050283.1 | Ephemeroptera | *Torleya grandiforceps* | T | G | A | C | C | G | T | G | C | A | A | A | G | G | T | A | G | C |
| MT274132.1 | Ephemeroptera | *Torleya nepalica* | T | G | A | C | C | G | T | G | C | A | A | A | G | G | T | A | G | C |
| NC050284.1 | Ephemeroptera | *Torleya nepalica* | T | G | A | C | C | G | T | G | C | A | A | A | G | G | T | A | G | C |
| HM067837.1 | Ephemeroptera | *Vietnamella dabieshanensis* | T | G | A | C | C | G | T | G | C | A | A | A | G | G | T | A | G | C |
| KM244655.1 | Ephemeroptera | *Vietnamella* sp. | T | G | A | C | C | G | T | G | C | A | A | A | G | G | T | A | G | C |
| MH751435.1 | Odonata | *Acisoma panorpoides* | T | G | A | C | C | G | T | G | C | A | A | A | G | G | T | A | G | C |
| NC042689.1 | Odonata | *Acisoma panorpoides* | T | G | A | C | C | G | T | G | C | A | A | A | G | G | T | A | G | C |
| KX161841.1 | Odonata | *Anax imperator* | T | G | A | C | C | G | T | G | C | A | A | A | G | G | T | A | G | C |
| NC031821.1 | Odonata | *Anax imperator* | T | G | A | C | C | G | T | G | C | A | A | A | G | G | T | A | G | C |
| KP233805.1 | Odonata | *Atrocalopteryx atrata* | T | G | A | C | C | G | T | G | C | A | A | A | G | G | T | A | G | C |
| NC027181.1 | Odonata | *Atrocalopteryx atrata* | T | G | A | C | C | G | T | G | C | A | A | A | G | G | T | A | G | C |
| MG011692.1 | Odonata | *Atrocalopteryx melli* | T | G | A | C | C | G | T | G | C | A | A | A | G | G | T | A | G | C |
| NC036749.1 | Odonata | *Atrocalopteryx melli* | T | G | A | C | C | G | T | G | C | A | A | A | G | G | T | A | G | C |
| KM658172.1 | Odonata | *Brachythemis contaminata* | T | G | A | C | C | G | T | G | C | A | A | A | G | G | T | A | G | C |
| NC026305.1 | Odonata | *Brachythemis contaminata* | T | G | A | C | C | G | T | G | C | A | A | A | G | G | T | A | G | C |
| EU591677.1 | Odonata | *Davidius lunatus* | T | G | A | C | C | G | T | G | C | A | A | A | G | G | T | A | G | C |
| NC012644.1 | Odonata | *Davidius lunatus* | T | G | A | C | C | G | T | G | C | A | A | A | G | G | T | A | G | C |
| MH751436.1 | Odonata | *Deielia phaon* | T | G | A | C | C | G | T | G | C | A | A | A | G | G | T | A | G | C |
| NC042690.1 | Odonata | *Deielia phaon* | T | G | A | C | C | G | T | G | C | A | A | A | G | G | T | A | G | C |
| MF716899.1 | Odonata | *Enallagma cyathigerum* | T | G | A | C | C | G | T | G | C | A | A | A | G | G | T | A | G | C |
| JX050223.1 | Odonata | *Epiophlebia superstes* | T | G | A | C | C | G | T | G | C | A | A | A | G | G | T | A | G | C |
| NC023232.1 | Odonata | *Epiophlebia superstes* | T | G | A | C | C | G | T | G | C | A | A | A | G | G | T | A | G | C |

| | | | | | | | | | | | | | | | | | | | | |
|---|---|---|---|---|---|---|---|---|---|---|---|---|---|---|---|---|---|---|---|---|
| MK522522.1 | Odonata | *Epophthalmia elegans* | T | G | A | C | C | G | T | G | C | A | A | A | G | G | T | A | G | C |
| NC046756.1 | Odonata | *Epophthalmia elegans* | T | G | A | C | C | G | T | G | C | A | A | A | G | G | T | A | G | C |
| HM126547.1 | Odonata | *Euphaea formosa* | T | G | A | C | C | G | T | G | C | A | A | A | G | G | T | A | G | C |
| NC014493.1 | Odonata | *Euphaea formosa* | T | G | A | C | C | G | T | G | C | A | A | A | G | G | T | A | G | C |
| KM244659.1 | Odonata | *Hydrobasileus croceus* | T | G | A | C | C | G | T | G | C | A | A | A | G | G | T | A | G | C |
| NC025758.1 | Odonata | *Hydrobasileus croceus* | T | G | A | C | C | G | T | G | C | A | A | A | G | G | T | A | G | C |
| KM244673.1 | Odonata | *Ictinogomphus* sp. | T | G | A | C | C | G | T | G | C | A | A | A | G | G | T | A | G | C |
| KU958378.1 | Odonata | *Ischnura elegans* | T | G | A | C | C | G | T | G | C | A | A | A | G | G | T | A | G | C |
| MK951668.1 | Odonata | *Ischnura elegans* | T | G | A | C | C | G | T | G | C | A | A | A | G | G | T | A | G | C |
| NC031824.1 | Odonata | *Ischnura elegans* | T | G | A | C | C | G | T | G | C | A | A | A | G | G | T | A | G | C |
| KC878732.1 | Odonata | *Ischnura pumilio* | T | G | A | C | C | G | T | G | C | A | A | A | G | G | T | A | G | C |
| NC021617.1 | Odonata | *Ischnura pumilio* | T | G | A | C | C | G | T | G | C | A | A | A | G | G | T | A | G | C |
| MF990748.1 | Odonata | *Macromia daimoji* | T | G | A | C | C | G | T | G | C | A | A | A | G | G | T | A | G | C |
| NC041425.1 | Odonata | *Macromia daimoji* | T | G | A | C | C | G | T | G | C | A | A | A | G | G | T | A | G | C |
| MK722304.1 | Odonata | *Matrona basilaris* | T | G | A | C | C | G | T | G | C | A | A | A | G | G | T | A | G | C |
| MK722305.1 | Odonata | *Matrona basilaris* | T | G | A | C | C | G | T | G | C | A | A | A | G | G | T | A | G | C |
| KU958377.1 | Odonata | *Megaloprepus caerulatus* | T | G | A | C | T | G | T | G | C | A | A | A | G | G | T | A | G | C |
| NC031823.1 | Odonata | *Megaloprepus caerulatus* | T | G | A | C | T | G | T | G | C | A | A | A | G | G | T | A | G | C |
| KU871065.1 | Odonata | *Mnais costalis* | T | G | A | C | C | G | T | G | C | A | A | A | G | G | T | A | G | C |
| KY402222.1 | Odonata | *Nannophya pygmaea* | T | G | A | C | C | G | T | G | C | A | A | A | G | G | T | A | G | C |
| NC039410.1 | Odonata | *Nannophya pygmaea* | T | G | A | C | C | G | T | G | C | A | A | A | G | G | T | A | G | C |
| KU361233.1 | Odonata | *Orthetrum chrysis* | T | G | A | C | C | G | T | G | C | A | A | A | G | G | T | A | G | C |

| | | | | | | | | | | | | | | | | | | | | |
|---|---|---|---|---|---|---|---|---|---|---|---|---|---|---|---|---|---|---|---|---|
| NC032048.1 | Odonata | *Orthetrum chrysis* | T | G | A | C | C | G | T | G | C | A | A | A | G | G | T | A | G | C |
| KU361232.1 | Odonata | *Orthetrum glaucum* | T | G | A | C | C | G | T | G | C | A | A | A | G | G | T | A | G | C |
| NC032047.1 | Odonata | *Orthetrum glaucum* | T | G | A | C | C | G | T | G | C | A | A | A | G | G | T | A | G | C |
| MH751437.1 | Odonata | *Orthetrum melania* | T | G | A | C | C | G | T | G | C | A | A | A | G | G | T | A | G | C |
| NC042732.1 | Odonata | *Orthetrum melania* | T | G | A | C | C | G | T | G | C | A | A | A | G | G | T | A | G | C |
| KU361234.1 | Odonata | *Orthetrum sabina* | T | G | A | C | C | G | T | G | C | A | A | A | G | G | T | A | G | C |
| MH751438.1 | Odonata | *Orthetrum sabina* | T | G | A | C | C | G | T | G | C | A | A | A | G | G | T | A | G | C |
| NC032049.1 | Odonata | *Orthetrum sabina* | T | G | A | C | C | G | T | G | C | A | A | A | G | G | T | A | G | C |
| KU361235.1 | Odonata | *Orthetrum testaceum* | T | G | A | C | C | G | T | G | C | A | A | A | G | G | T | A | G | C |
| MH751439.1 | Odonata | *Orthetrum testaceum* | T | G | A | C | C | G | T | G | C | A | A | A | G | G | T | A | G | C |
| NC032050.1 | Odonata | *Orthetrum testaceum* | T | G | A | C | C | G | T | G | C | A | A | A | G | G | T | A | G | C |
| AB126005.1 | Odonata | *Orthetrum triangulare melania* | T | G | A | C | C | G | T | G | C | A | A | A | G | G | T | A | G | C |
| MK951669.1 | Odonata | *Paracercion v-nigrum* | T | G | A | C | C | G | T | G | C | A | A | A | G | G | T | A | G | C |
| KP233804.1 | Odonata | *Platycnemis foliacea* | T | G | A | C | C | G | T | G | C | A | A | A | G | G | T | A | G | C |
| NC027180.1 | Odonata | *Platycnemis foliacea* | T | G | A | C | C | G | T | G | C | A | A | A | G | G | T | A | G | C |
| FJ606784.1 | Odonata | *Pseudolestes mirabilis* | T | G | A | C | C | G | T | G | C | A | A | A | G | G | T | A | G | C |
| NC020636.1 | Odonata | *Pseudolestes mirabilis* | T | G | A | C | C | G | T | G | C | A | A | A | G | G | T | A | G | C |
| MF150044.1 | Odonata | *Psolodesmus mandarinus* | T | G | A | C | C | G | T | G | C | A | A | A | G | G | T | A | G | C |
| NC042442.1 | Odonata | *Psolodesmus mandarinus* | T | G | A | C | C | G | T | G | C | A | A | A | G | G | T | A | G | C |
| MG594801.1 | Odonata | *Somatochlora hineana* | T | G | A | C | C | G | T | G | C | A | A | A | G | G | T | A | G | C |
| MH751440.1 | Odonata | *Tramea virginia* | T | G | A | C | C | G | T | G | C | A | A | A | G | G | T | A | G | C |
| NC042691.1 | Odonata | *Tramea virginia* | T | G | A | C | C | G | T | G | C | A | A | A | G | G | T | A | G | C |

| | | | | | | | | | | | | | | | | | | | | |
|---|---|---|---|---|---|---|---|---|---|---|---|---|---|---|---|---|---|---|---|---|
| MH751441.1 | Odonata | *Trigomphus carus* | T | G | A | C | C | G | T | G | C | A | A | A | G | G | T | A | G | C |
| NC042733.1 | Odonata | *Trigomphus carus* | T | G | A | C | C | G | T | G | C | A | A | A | G | G | T | A | G | C |
| JX050224.1 | Odonata | *Vestalis melania* | T | G | A | C | C | G | T | G | C | A | A | A | G | G | T | A | G | C |
| NC023233.1 | Odonata | *Vestalis melania* | T | G | A | C | C | G | T | G | C | A | A | A | G | G | T | A | G | C |
| KM199685.1 | Plecoptera | *Acroneuria hainana* | T | G | A | C | C | G | T | G | C | A | A | A | G | G | T | A | G | C |
| NC026104.1 | Plecoptera | *Acroneuria hainana* | T | G | A | C | C | G | T | G | C | A | A | A | G | G | T | A | G | C |
| KX091846.1 | Plecoptera | *Agnetina aequalis* | T | G | A | C | C | G | T | G | C | A | A | A | G | G | T | A | G | C |
| MH085446.1 | Plecoptera | *Amphinemura longispina* | T | G | A | C | C | G | T | G | C | A | A | A | G | G | T | A | G | C |
| NC044748.1 | Plecoptera | *Amphinemura longispina* | T | G | A | C | C | G | T | G | C | A | A | A | G | G | T | A | G | C |
| KX091847.1 | Plecoptera | *Amphinemura* sp. | T | G | A | C | C | G | T | G | C | A | A | A | G | G | T | A | G | C |
| MH085447.1 | Plecoptera | *Amphinemura yao* | T | G | A | C | C | G | T | G | C | A | A | A | G | G | T | A | G | C |
| NC044749.1 | Plecoptera | *Amphinemura yao* | T | G | A | C | C | G | T | G | C | A | A | A | G | G | T | A | G | C |
| MK111413.1 | Plecoptera | *Antarctoperla michaelseni* | T | G | A | C | C | G | T | G | C | A | A | A | G | G | T | A | G | C |
| NC042199.1 | Plecoptera | *Antarctoperla michaelseni* | T | G | A | C | C | G | T | G | C | A | A | A | G | G | T | A | G | C |
| KR604721.1 | Plecoptera | *Apteroperla tikumana* | T | G | A | C | C | G | T | G | C | A | A | A | G | G | T | A | G | C |
| NC027698.1 | Plecoptera | *Apteroperla tikumana* | T | G | A | C | C | G | T | G | C | A | A | A | G | G | T | A | G | C |
| MT410838.1 | Plecoptera | *Brachyptera risi* | T | G | A | C | C | G | T | G | C | A | A | A | G | G | T | A | G | C |
| MG677941.1 | Plecoptera | *Calineuria stigmatica* | T | G | A | C | C | G | T | G | C | A | A | A | G | G | T | A | G | C |
| KX094942.1 | Plecoptera | *Capnia zijinshana* | T | G | A | C | C | G | T | G | C | A | A | A | G | G | T | A | G | C |
| NC034661.1 | Plecoptera | *Capnia zijinshana* | T | G | A | C | C | G | T | G | C | A | A | A | G | G | T | A | G | C |
| MG677942.1 | Plecoptera | *Caroperla siveci* | T | G | A | C | C | G | T | G | C | A | A | A | G | G | T | A | G | C |
| MF100783.1 | Plecoptera | *Cerconychia flectospina* | T | G | A | C | C | G | T | G | C | A | A | A | G | G | T | A | G | C |

| | | | | | | | | | | | | | | | | | | | | |
|---|---|---|---|---|---|---|---|---|---|---|---|---|---|---|---|---|---|---|---|---|
| MN419914.1 | Plecoptera | *Claassenia* sp. | T | G | A | C | C | G | T | G | C | T | A | A | G | G | T | A | G | C |
| KC952026.1 | Plecoptera | *Cryptoperla* sp. | T | G | A | C | T | G | T | G | C | A | A | A | G | G | T | A | G | C |
| MK111416.1 | Plecoptera | *Diamphipnoa annulata* | T | G | A | C | C | G | T | G | C | A | A | A | G | G | T | A | G | C |
| NC042205.1 | Plecoptera | *Diamphipnoa annulata* | T | G | A | C | C | G | T | G | C | A | A | A | G | G | T | A | G | C |
| MK111417.1 | Plecoptera | *Diamphipnopsis* sp. | T | G | A | C | C | G | T | G | C | A | A | A | G | G | T | A | G | C |
| KF484757.1 | Plecoptera | *Dinocras cephalotes* | T | G | A | C | C | G | T | G | C | A | A | A | G | G | T | A | G | C |
| NC022843.1 | Plecoptera | *Dinocras cephalotes* | T | G | A | C | C | G | T | G | C | A | A | A | G | G | T | A | G | C |
| MG589787.1 | Plecoptera | *Doddsia occidentalis* | T | G | A | C | C | G | T | G | C | A | A | A | G | G | T | A | G | C |
| MK905888.1 | Plecoptera | *Etrocorema hochii* | T | G | A | C | T | G | T | G | C | A | A | A | G | G | T | A | G | C |
| MK905206.1 | Plecoptera | *Flavoperla* sp. | T | G | A | C | C | G | T | G | C | A | A | A | G | G | T | A | G | C |
| MN419916.1 | Plecoptera | *Flavoperla* sp. | T | G | A | C | C | G | T | G | C | A | A | A | G | G | T | A | G | C |
| MN419915.1 | Plecoptera | *Indonemoura auriformis* | T | G | A | C | C | G | T | G | C | A | A | A | G | G | T | A | G | C |
| MH085448.1 | Plecoptera | *Indonemoura jacobsoni* | T | G | A | C | C | G | T | G | C | A | A | A | G | G | T | A | G | C |
| NC044750.1 | Plecoptera | *Indonemoura jacobsoni* | T | G | A | C | C | G | T | G | C | A | A | A | G | G | T | A | G | C |
| MH085449.1 | Plecoptera | *Indonemoura nohirae* | T | G | A | C | C | G | T | G | C | A | A | A | G | G | T | A | G | C |
| NC044751.1 | Plecoptera | *Indonemoura nohirae* | T | G | A | C | C | G | T | G | C | A | A | A | G | G | T | A | G | C |
| MF716959.1 | Plecoptera | *Isoperla bilineata* | T | G | A | C | C | G | T | G | C | A | A | A | G | G | T | A | G | C |
| NC038190.1 | Plecoptera | *Isoperla bilineata* | T | G | A | C | C | G | T | G | C | A | A | A | G | G | T | A | G | C |
| MG910457.1 | Plecoptera | *Isoperla eximia* | T | G | A | C | C | G | T | G | C | A | A | A | G | G | T | A | G | C |
| NC038167.1 | Plecoptera | *Isoperla eximia* | T | G | A | C | C | G | T | G | C | A | A | A | G | G | T | A | G | C |
| KT186102.1 | Plecoptera | *Kamimuria chungnanshana* | T | G | A | C | C | G | T | G | C | A | A | A | G | G | T | A | G | C |
| NC028076.1 | Plecoptera | *Kamimuria chungnanshana* | T | G | A | C | C | G | T | G | C | A | A | A | G | G | T | A | G | C |

| | | | | | | | | | | | | | | | | | | | | |
|---|---|---|---|---|---|---|---|---|---|---|---|---|---|---|---|---|---|---|---|---|
| MN400755.1 | Plecoptera | *Kamimuria klapaleki* | T | G | A | C | C | G | T | G | C | A | A | A | G | G | T | A | G | C |
| MK111414.1 | Plecoptera | *Klapopteryx armillata* | T | G | A | C | C | G | T | G | C | A | A | A | G | G | T | A | G | C |
| MH374046.1 | Plecoptera | *Lednia tumana* | T | G | A | C | C | G | T | G | C | A | A | A | G | G | T | A | G | C |
| MK568475.1 | Plecoptera | *Leuctra fusca* | T | G | A | C | C | G | T | G | C | A | A | A | G | G | T | A | G | C |
| KP642637.1 | Plecoptera | *Mesocapnia arizonensis* | T | G | A | C | C | G | T | G | C | A | A | A | G | G | T | A | G | C |
| KY568983.1 | Plecoptera | *Mesocapnia daxingana* | T | G | A | C | C | G | T | G | C | A | A | A | G | G | T | A | G | C |
| MH085450.1 | Plecoptera | *Mesonemoura metafiligera* | T | G | A | C | C | G | T | G | C | A | A | A | G | G | T | A | G | C |
| NC044719.1 | Plecoptera | *Mesonemoura metafiligera* | T | G | A | C | C | G | T | G | C | A | A | A | G | G | T | A | G | C |
| MH085451.1 | Plecoptera | *Mesonemoura tritaenia* | T | G | A | C | C | G | T | G | C | A | A | A | G | G | T | A | G | C |
| NC044720.1 | Plecoptera | *Mesonemoura tritaenia* | T | G | A | C | C | G | T | G | C | A | A | A | G | G | T | A | G | C |
| MN096323.1 | Plecoptera | *Microperla geei* | T | G | A | C | C | G | T | G | C | A | A | A | G | G | T | A | G | C |
| MT410776.1 | Plecoptera | *Nemoura avicularis* | T | G | A | C | C | G | T | G | C | A | A | A | G | G | T | A | G | C |
| MT410849.1 | Plecoptera | *Nemoura cinerea* | T | G | A | C | C | G | T | G | C | A | A | A | G | G | T | A | G | C |
| KY940360.1 | Plecoptera | *Nemoura nankinensis* | T | G | A | C | C | G | T | G | C | A | A | A | G | G | T | A | G | C |
| NC034939.1 | Plecoptera | *Nemoura nankinensis* | T | G | A | C | C | G | T | G | C | A | A | A | G | G | T | A | G | C |
| MK290826.1 | Plecoptera | *Nemoura papilla* | T | G | A | C | C | G | T | G | C | A | A | A | G | G | T | A | G | C |
| MK111418.1 | Plecoptera | *Neonemura barrosi* | T | G | A | C | C | G | T | G | C | A | A | A | G | G | T | A | G | C |
| NC042206.1 | Plecoptera | *Neonemura barrosi* | T | G | A | C | C | G | T | G | C | A | A | A | G | G | T | A | G | C |
| KX091858.1 | Plecoptera | *Neoperla ignacsiveci* | T | G | A | C | C | G | T | G | C | A | A | A | G | G | T | A | G | C |
| KX091859.1 | Plecoptera | *Neoperla* sp. | T | G | A | C | C | G | T | G | C | A | A | A | G | G | T | A | G | C |
| MN400756.1 | Plecoptera | *Neoperlops gressitti* | T | G | A | C | C | G | T | G | C | A | A | A | G | G | T | A | G | C |
| MK111415.1 | Plecoptera | *Neuroperla schedingi* | T | G | A | C | C | G | T | G | C | A | A | A | G | G | T | A | G | C |

| | | | | | | | | | | | | | | | | | | | | |
|---|---|---|---|---|---|---|---|---|---|---|---|---|---|---|---|---|---|---|---|---|
| NC042200.1 | Plecoptera | *Neuroperla schedingi* | T | G | A | C | C | G | T | G | C | A | A | A | G | G | T | A | G | C |
| MK686067.1 | Plecoptera | *Niponiella limbatella* | T | G | A | C | C | G | T | G | C | A | A | A | G | G | T | A | G | C |
| MK387068.1 | Plecoptera | *Peltoperlopsis cebuano* | T | G | A | C | C | G | T | G | C | A | A | A | G | G | T | A | G | C |
| MF197377.1 | Plecoptera | *Perlodes* sp. | T | G | A | C | C | G | T | G | C | A | A | A | G | G | T | A | G | C |
| MH085452.1 | Plecoptera | *Protonemura kohnoae* | T | G | A | C | C | G | T | G | C | A | A | A | G | G | T | A | G | C |
| NC044752.1 | Plecoptera | *Protonemura kohnoae* | T | G | A | C | C | G | T | G | C | A | A | A | G | G | T | A | G | C |
| MT410850.1 | Plecoptera | *Protonemura meyeri* | T | G | A | C | C | G | T | G | C | A | A | A | G | G | T | A | G | C |
| NC050322.1 | Plecoptera | *Protonemura meyeri* | T | G | A | C | C | G | T | G | C | A | A | A | G | G | T | A | G | C |
| MH085453.1 | Plecoptera | *Protonemura orbiculata* | T | G | A | C | C | G | T | G | C | A | A | A | G | G | T | A | G | C |
| NC044753.1 | Plecoptera | *Protonemura orbiculata* | T | G | A | C | C | G | T | G | C | A | A | A | G | G | T | A | G | C |
| MG910458.1 | Plecoptera | *Pseudomegarcys japonica* | T | G | A | C | C | G | T | G | C | A | A | A | G | G | T | A | G | C |
| NC038168.1 | Plecoptera | *Pseudomegarcys japonica* | T | G | A | C | C | G | T | G | C | A | A | A | G | G | T | A | G | C |
| KU182360.1 | Plecoptera | *Pteronarcella badia* | T | G | A | C | C | G | T | G | C | A | A | A | G | G | T | A | G | C |
| NC029248.1 | Plecoptera | *Pteronarcella badia* | T | G | A | C | C | G | T | G | C | A | A | A | G | G | T | A | G | C |
| AY687866.1 | Plecoptera | *Pteronarcys princeps* | T | G | A | C | C | G | T | G | C | A | A | A | G | G | T | A | G | C |
| NC006133.1 | Plecoptera | *Pteronarcys princeps* | T | G | A | C | C | G | T | G | C | A | A | A | G | G | T | A | G | C |
| MK111419.1 | Plecoptera | *Rhopalopsole bulbifera* | T | G | A | C | C | G | T | G | C | A | A | A | G | G | T | A | G | C |
| NC042207.1 | Plecoptera | *Rhopalopsole bulbifera* | T | G | A | C | C | G | T | G | C | A | A | A | G | G | T | A | G | C |
| MH510071.1 | Plecoptera | *Scopura longa* | T | G | A | C | C | G | T | G | C | A | A | A | G | G | T | A | G | C |
| NC041105.1 | Plecoptera | *Scopura longa* | T | G | A | C | C | G | T | G | C | A | A | A | G | G | T | A | G | C |
| MK492253.1 | Plecoptera | *Sinacroneuria dabieshana* | T | G | A | C | C | G | T | G | C | A | A | A | G | G | T | A | G | C |
| MF716958.1 | Plecoptera | *Soliperla* sp. | T | G | A | C | C | G | T | G | C | A | A | A | G | G | T | A | G | C |

| | | | | | | | | | | | | | | | | | | | | |
|---|---|---|---|---|---|---|---|---|---|---|---|---|---|---|---|---|---|---|---|---|
| NC038189.1 | Plecoptera | *Soliperla* sp. | T | G | A | C | C | G | T | G | C | A | A | A | G | G | T | A | G | C |
| MH085455.1 | Plecoptera | *Sphaeronemoura acutispina* | T | G | A | C | C | G | T | G | C | A | A | A | G | G | T | A | G | C |
| MH085454.1 | Plecoptera | *Sphaeronemoura grandicauda* | T | G | A | C | C | G | T | G | C | A | A | A | G | G | T | A | G | C |
| NC044754.1 | Plecoptera | *Sphaeronemoura grandicauda* | T | G | A | C | C | G | T | G | C | A | A | A | G | G | T | A | G | C |
| MK111420.1 | Plecoptera | *Sphaeronemoura hainana* | T | G | A | C | C | G | T | G | C | A | A | A | G | G | T | A | G | C |
| NC044755.1 | Plecoptera | *Sphaeronemoura hamistyla* | T | G | A | C | C | G | T | G | C | A | A | A | G | G | T | A | G | C |
| KR088971.1 | Plecoptera | *Styloperla* sp. | T | G | A | C | C | G | T | G | C | A | A | A | G | G | T | A | G | C |
| KX845569.1 | Plecoptera | *Styloperla spinicercia* | T | G | A | C | C | G | T | G | C | A | A | A | G | G | T | A | G | C |
| NC034809.1 | Plecoptera | *Styloperla spinicercia* | T | G | A | C | C | G | T | G | C | A | A | A | G | G | T | A | G | C |
| KX091865.1 | Plecoptera | *Suwallia bimaculata* | T | G | A | C | C | G | T | G | C | A | A | A | G | G | T | A | G | C |
| MN121757.1 | Plecoptera | *Suwallia bimaculata* | T | G | A | C | C | G | T | G | C | A | A | A | G | G | T | A | G | C |
| MF198253.1 | Plecoptera | *Suwallia teleckojensis* | T | G | A | C | C | G | T | G | C | A | A | A | G | G | T | A | G | C |
| NC037754.1 | Plecoptera | *Suwallia teleckojensis* | T | G | A | C | C | G | T | G | C | A | A | A | G | G | T | A | G | C |
| MG589786.1 | Plecoptera | *Taeniopteryx ugola* | T | G | A | C | C | G | T | G | C | A | A | A | G | G | T | A | G | C |
| NC037897.1 | Plecoptera | *Taeniopteryx ugola* | T | G | A | C | C | G | T | G | C | A | A | A | G | G | T | A | G | C |
| KM409708.1 | Plecoptera | *Togoperla* sp. | T | G | A | C | C | G | T | G | C | A | A | A | G | G | T | A | G | C |
| KY522907.1 | Plecoptera | *Zelandoperla fenestrata* | T | G | A | C | C | G | T | G | C | A | A | A | G | G | T | A | G | C |
| NC034997.1 | Plecoptera | *Zelandoperla fenestrata* | T | G | A | C | C | G | T | G | C | A | A | A | G | G | T | A | G | C |
| FJ456939.1 | Heteroptera | *Aphelocheirus ellipsoideus* | T | G | A | C | T | G | T | G | C | A | A | A | G | G | T | A | G | C |
| KX505849.1 | Heteroptera | *Aphelocheirus jendeki* | T | A | A | C | C | G | T | G | C | A | A | A | G | G | T | A | G | C |
| NC037370.1 | Heteroptera | *Aphelocheirus jendeki* | T | A | A | C | C | G | T | G | C | A | A | A | G | G | T | A | G | C |
| FJ456944.1 | Heteroptera | *Aquarius paludum* | T | G | A | C | T | G | T | G | C | A | A | A | G | G | T | A | G | C |

| | | | | | | | | | | | | | | | | | | | | |
|---|---|---|---|---|---|---|---|---|---|---|---|---|---|---|---|---|---|---|---|---|
| NC012841.1 | Heteroptera | *Aquarius paludum* | T | G | A | C | T | G | T | G | C | A | A | A | G | G | T | A | G | C |
| MK251109.1 | Heteroptera | *Diplonychus esakii* | T | A | A | C | C | G | T | G | C | A | A | A | G | G | T | A | G | C |
| FJ456940.1 | Heteroptera | *Diplonychus rusticus* | T | A | A | C | C | G | T | G | C | A | A | A | G | G | T | A | G | C |
| FJ456949.1 | Heteroptera | *Enithares tibialis* | T | A | A | C | C | G | T | G | C | A | A | A | G | G | T | A | G | C |
| NC012819.1 | Heteroptera | *Enithares tibialis* | T | A | A | C | C | G | T | G | C | A | A | A | G | G | T | A | G | C |
| KP400582.1 | Heteroptera | *Entomovelia* sp. | T | A | A | C | T | G | T | A | C | T | A | A | G | G | T | A | G | C |
| MF177288.1 | Heteroptera | *Gigantometra gigas* | T | A | A | C | T | G | T | A | C | G | A | A | G | G | T | A | G | C |
| MF177289.1 | Heteroptera | *Gigantometra gigas* | T | A | A | C | T | G | T | A | C | G | A | A | G | G | T | A | G | C |
| MF177290.1 | Heteroptera | *Gigantometra gigas* | T | A | A | C | T | G | T | A | C | G | A | A | G | G | T | A | G | C |
| MF177291.1 | Heteroptera | *Gigantometra gigas* | T | A | A | C | T | G | T | A | C | G | A | A | G | G | T | A | G | C |
| NC041084.1 | Heteroptera | *Gigantometra gigas* | T | A | A | C | T | G | T | A | C | G | A | A | G | G | T | A | G | C |
| KJ027513.1 | Heteroptera | *Helotrephes semiglobosus semiglobosus* | T | A | A | C | T | G | T | G | C | G | A | A | G | G | T | A | G | C |
| FJ456951.1 | Heteroptera | *Helotrephes* sp. | T | A | A | C | T | G | T | G | C | G | A | A | G | G | T | A | G | C |
| NC012822.1 | Heteroptera | *Helotrephes* sp. | T | A | A | C | T | G | T | G | C | G | A | A | G | G | T | A | G | C |
| FJ456945.1 | Heteroptera | *Hydrometra greeni* | T | A | A | C | C | G | T | A | C | A | A | A | G | G | T | A | G | C |
| NC012842.1 | Heteroptera | *Hydrometra greeni* | T | A | A | C | C | G | T | A | C | A | A | A | G | G | T | A | G | C |
| FJ456947.1 | Heteroptera | *Ilyocoris cimicoides* | T | G | A | C | C | G | T | G | C | A | A | A | G | G | T | A | G | C |
| NC012845.1 | Heteroptera | *Ilyocoris cimicoides* | T | G | A | C | C | G | T | G | C | A | A | A | G | G | T | A | G | C |
| FJ456948.1 | Heteroptera | *Laccotrephes robustus* | T | A | A | C | C | G | T | G | C | A | A | A | G | G | T | A | G | C |
| NC012817.1 | Heteroptera | *Laccotrephes robustus* | T | A | A | C | C | G | T | G | C | A | A | A | G | G | T | A | G | C |
| FJ456946.1 | Heteroptera | *Leptopus* sp. | T | A | A | C | T | G | T | G | C | A | A | A | G | G | T | A | G | C |

| | | | | | | | | | | | | | | | | | | | | |
|---|---|---|---|---|---|---|---|---|---|---|---|---|---|---|---|---|---|---|---|---|
| JQ910985.1 | Heteroptera | *Lethocerus deyrollei* | T | G | A | C | C | G | T | G | C | A | A | A | G | G | T | A | G | C |
| KU237288.1 | Heteroptera | *Lethocerus deyrollei* | T | G | A | C | C | G | T | G | C | A | A | A | G | G | T | A | G | C |
| KM588201.1 | Heteroptera | *Lethocerus indicus* | T | G | A | C | C | G | T | G | C | A | A | A | G | G | T | A | G | C |
| KR072671.1 | Heteroptera | *Lethocerus indicus* | T | G | A | C | C | G | T | G | C | A | A | A | G | G | T | A | G | C |
| NC027194.1 | Heteroptera | *Lethocerus indicus* | T | G | A | C | C | G | T | G | C | A | A | A | G | G | T | A | G | C |
| FJ456942.1 | Heteroptera | *Lycorma delicatula* | T | G | A | C | T | G | T | G | C | T | A | A | G | G | T | A | G | C |
| NC012835.1 | Heteroptera | *Lycorma delicatula* | T | G | A | C | T | G | T | G | C | T | A | A | G | G | T | A | G | C |
| MN027278.1 | Heteroptera | *Metrocoris* sp. | T | A | A | C | T | G | T | A | C | T | A | A | G | G | T | A | G | C |
| MH382835.1 | Heteroptera | *Micronecta sahlbergii* | T | G | A | C | C | G | T | G | C | A | A | A | G | G | T | A | G | C |
| NC039588.1 | Heteroptera | *Micronecta sahlbergii* | T | G | A | C | C | G | T | G | C | A | A | A | G | G | T | A | G | C |
| KT031802.1 | Heteroptera | *Nepa hoffmanni* | T | A | A | C | C | G | T | G | C | A | A | A | G | G | T | A | G | C |
| NC028084.1 | Heteroptera | *Nepa hoffmanni* | T | A | A | C | C | G | T | G | C | A | A | A | G | G | T | A | G | C |
| FJ456943.1 | Heteroptera | *Nerthra indica* | T | A | A | C | C | G | T | A | C | A | A | A | G | G | T | A | G | C |
| NC012838.1 | Heteroptera | *Nerthra indica* | T | A | A | C | C | G | T | A | C | A | A | A | G | G | T | A | G | C |
| KX034086.1 | Heteroptera | *Notonecta chinensis* | T | A | A | C | C | G | T | G | C | A | A | A | G | G | T | A | G | C |
| NC036671.1 | Heteroptera | *Notonecta chinensis* | T | A | A | C | C | G | T | G | C | A | A | A | G | G | T | A | G | C |
| MH780861.1 | Heteroptera | *Notonecta montandoni* | T | A | A | C | C | G | T | G | C | A | A | A | G | G | T | A | G | C |
| NC045886.1 | Heteroptera | *Notonecta montandoni* | T | A | A | C | C | G | T | G | C | A | A | A | G | G | T | A | G | C |
| KM244707.1 | Heteroptera | *Notonectidae* sp. | T | A | A | C | T | G | T | G | C | A | A | A | G | G | T | A | G | C |
| FJ456950.1 | Heteroptera | *Ochterus marginatus* | T | A | A | C | C | G | T | A | C | A | A | A | G | G | T | A | G | C |
| KJ027516.1 | Heteroptera | *Paraplea frontalis* | T | A | A | C | T | G | T | G | C | A | A | A | G | G | T | A | G | C |
| NC028629.1 | Heteroptera | *Paraplea frontalis* | T | A | A | C | T | G | T | G | C | A | A | A | G | G | T | A | G | C |

| | | | | | | | | | | | | | | | | | | | | |
|---|---|---|---|---|---|---|---|---|---|---|---|---|---|---|---|---|---|---|---|---|
| JQ910988.1 | Heteroptera | *Perittopus* sp. | T | A | A | C | T | G | T | A | C | G | A | A | G | G | T | A | G | C |
| KP400583.1 | Heteroptera | *Ptilomera tigrina* | T | A | A | C | T | G | T | A | C | G | A | A | G | G | T | A | G | C |
| MN027271.1 | Heteroptera | *Rhagovelia reitteri* | T | A | A | C | T | G | T | A | C | G | A | A | G | G | T | A | G | C |
| NC044721.1 | Heteroptera | *Rhagovelia reitteri* | T | A | A | C | T | G | T | A | C | G | A | A | G | G | T | A | G | C |
| MN027279.1 | Heteroptera | *Rhagovelia* sp. | T | A | A | C | T | G | T | A | C | G | A | A | G | G | T | A | G | C |
| FJ456941.1 | Heteroptera | *Sigara septemlineata* | T | G | A | C | C | G | T | G | C | A | A | A | G | G | T | A | G | C |
| FJ456952.1 | Heteroptera | *Valentia hoffmanni* | T | A | A | C | C | G | T | A | C | T | A | A | G | G | T | A | G | C |
| NC012823.1 | Heteroptera | *Valentia hoffmanni* | T | A | A | C | C | G | T | A | C | T | A | A | G | G | T | A | G | C |
| KF840564.1 | Megaloptera | *Acanthacorydalis orientalis* | T | G | A | C | T | G | T | G | C | A | A | A | G | G | T | A | G | C |
| NC023462.1 | Megaloptera | *Acanthacorydalis orientalis* | T | G | A | C | T | G | T | G | C | A | A | A | G | G | T | A | G | C |
| KU925864.1 | Megaloptera | *Archichauliodes deceptor* | T | G | A | C | T | G | T | G | C | A | A | A | G | G | T | A | G | C |
| NC035504.1 | Megaloptera | *Archichauliodes deceptor* | T | G | A | C | T | G | T | G | C | A | A | A | G | G | T | A | G | C |
| KU302713.1 | Megaloptera | *Chloronia mirifica* | T | G | A | C | T | G | T | G | C | A | A | A | G | G | T | A | G | C |
| NC033349.1 | Megaloptera | *Chloronia mirifica* | T | G | A | C | T | G | T | G | C | A | A | A | G | G | T | A | G | C |
| FJ171323.1 | Megaloptera | *Corydalus cornutus* | T | G | A | C | T | G | T | G | C | A | A | A | G | G | T | A | G | C |
| NC011276.1 | Megaloptera | *Corydalus cornutus* | T | G | A | C | T | G | T | G | C | A | A | A | G | G | T | A | G | C |
| KJ806318.1 | Megaloptera | *Dysmicohermes ingens* | T | G | A | C | T | G | T | G | C | A | A | A | G | G | T | A | G | C |
| NC024657.1 | Megaloptera | *Dysmicohermes ingens* | T | G | A | C | T | G | T | G | C | A | A | A | G | G | T | A | G | C |
| JQ351950.1 | Megaloptera | *Neochauliodes bowringi* | T | G | A | C | T | G | T | G | C | A | A | A | G | G | T | A | G | C |
| NC023444.1 | Megaloptera | *Neochauliodes bowringi* | T | G | A | C | T | G | T | G | C | A | A | A | G | G | T | A | G | C |
| KF785799.1 | Megaloptera | *Neochauliodes fraternus* | T | G | A | C | T | G | T | G | C | A | A | A | G | G | T | A | G | C |
| NC025282.1 | Megaloptera | *Neochauliodes fraternus* | T | G | A | C | T | G | T | G | C | A | A | A | G | G | T | A | G | C |

| | | | | | | | | | | | | | | | | | | | | |
|---|---|---|---|---|---|---|---|---|---|---|---|---|---|---|---|---|---|---|---|---|
| KX821680.1 | Megaloptera | *Neochauliodes parasparsus* | T | G | A | C | T | G | T | G | C | A | A | A | G | G | T | A | G | C |
| JX110703.1 | Megaloptera | *Neochauliodes punctatolosus* | T | G | A | C | T | G | T | G | C | A | A | A | G | G | T | A | G | C |
| MK301249.1 | Megaloptera | *Neochauliodes punctatolosus* | T | G | A | C | T | G | T | G | C | A | A | A | G | G | T | A | G | C |
| NC018772.1 | Megaloptera | *Neochauliodes punctatolosus* | T | G | A | C | T | G | T | G | C | A | A | A | G | G | T | A | G | C |
| KF771894.1 | Megaloptera | *Neochauliodes rotundatus* | T | G | A | C | T | G | T | G | C | A | A | A | G | G | T | A | G | C |
| NC025281.1 | Megaloptera | *Neochauliodes rotundatus* | T | G | A | C | T | G | T | G | C | A | A | A | G | G | T | A | G | C |
| KP126231.1 | Megaloptera | *Neoneuromus tonkinensis* | T | G | A | C | T | G | T | G | C | A | A | A | G | G | T | A | G | C |
| NC027852.1 | Megaloptera | *Neoneuromus tonkinensis* | T | G | A | C | T | G | T | G | C | A | A | A | G | G | T | A | G | C |
| KP126232.1 | Megaloptera | *Nevromus exterior* | T | G | A | C | T | G | T | G | C | A | A | A | G | G | T | A | G | C |
| NC027851.1 | Megaloptera | *Nevromus exterior* | T | G | A | C | T | G | T | G | C | A | A | A | G | G | T | A | G | C |
| KY230493.1 | Megaloptera | *Protochauliodes biconicus* | T | G | A | C | T | G | T | G | C | A | A | A | G | G | T | A | G | C |
| EU526394.1 | Megaloptera | *Protohermes concolorus* | T | G | A | C | T | G | T | G | C | A | A | A | G | G | T | A | G | C |
| NC011524.1 | Megaloptera | *Protohermes concolorus* | T | G | A | C | T | G | T | G | C | A | A | A | G | G | T | A | G | C |
| FJ859905.1 | Megaloptera | *Sialis hamata* | T | G | A | C | T | G | T | G | C | A | A | A | G | G | T | A | G | C |
| NC013256.1 | Megaloptera | *Sialis hamata* | T | G | A | C | T | G | T | G | C | A | A | A | G | G | T | A | G | C |
| KT876911.1 | Megaloptera | *Sialis* sp. | T | G | A | C | T | G | T | G | C | A | A | A | G | G | T | A | G | C |
| KT425084.1 | Megaloptera | *Stenosialis australiensis* | T | G | A | C | T | G | T | G | C | A | A | A | G | G | T | A | G | C |
| KY587199.1 | Neuroptera | *Abachrysa eureka* | T | G | A | C | T | G | T | G | C | A | A | A | G | G | T | A | G | C |
| NC015095.1 | Neuroptera | *Apochrysa matsumurae* | T | G | A | C | T | G | T | G | C | A | A | A | G | G | T | A | G | C |
| KC758703.1 | Neuroptera | *Ascalohybris subjacens* | T | G | A | C | T | G | T | G | C | A | A | A | G | G | T | A | G | C |
| NC021428.1 | Neuroptera | *Ascalohybris subjacens* | T | G | A | C | T | G | T | G | C | A | A | A | G | G | T | A | G | C |
| FJ171324.1 | Neuroptera | *Ascaloptynx appendiculatus* | T | G | A | C | T | G | T | G | C | A | A | A | G | G | T | A | G | C |

| | | | | | | | | | | | | | | | | | | | | |
|---|---|---|---|---|---|---|---|---|---|---|---|---|---|---|---|---|---|---|---|---|
| NC011277.1 | Neuroptera | *Ascaloptynx appendiculatus* | T | G | A | C | T | G | T | G | C | A | A | A | G | G | T | A | G | C |
| KT425083.1 | Neuroptera | *Balmes birmanus* | T | G | A | C | T | G | T | G | C | A | A | A | G | G | T | A | G | C |
| KX369241.1 | Neuroptera | *Bullanga florida* | T | G | A | C | T | G | T | G | C | A | A | A | G | G | T | A | G | C |
| NC032298.1 | Neuroptera | *Bullanga florida* | T | G | A | C | T | G | T | G | C | A | A | A | G | G | T | A | G | C |
| KT425069.1 | Neuroptera | *Chasmoptera huttii* | T | G | A | C | T | G | T | G | C | A | A | A | G | G | T | A | G | C |
| JX033119.1 | Neuroptera | *Chrysopa pallens* | T | G | A | C | T | G | T | G | C | A | A | A | G | G | T | A | G | C |
| NC019618.1 | Neuroptera | *Chrysopa pallens* | T | G | A | C | T | G | T | G | C | A | A | A | G | G | T | A | G | C |
| KJ778879.1 | Neuroptera | *Chrysoperla carnea* | T | G | A | C | T | G | T | G | C | A | A | A | G | G | T | A | G | C |
| KU877169.1 | Neuroptera | *Chrysoperla externa* | T | G | A | C | T | G | T | G | C | A | A | A | G | G | T | A | G | C |
| NC030341.1 | Neuroptera | *Chrysoperla externa* | T | G | A | C | T | G | T | G | C | A | A | A | G | G | T | A | G | C |
| KP027407.1 | Neuroptera | *Chrysoperla nipponensis* | T | G | A | C | T | G | T | G | C | A | A | A | G | G | T | A | G | C |
| NC015093.1 | Neuroptera | *Chrysoperla nipponensis* | T | G | A | C | T | G | T | G | C | A | A | A | G | G | T | A | G | C |
| KT425088.1 | Neuroptera | *Climacia areolaris* | T | G | A | C | T | G | T | G | C | A | A | A | G | G | T | A | G | C |
| KT425078.1 | Neuroptera | *Coniopteryx* sp. | T | G | A | C | T | G | T | G | C | T | A | A | G | G | T | A | G | C |
| KT425068.1 | Neuroptera | *Dendroleon pantherinus* | T | G | A | C | T | G | T | G | C | A | A | A | G | G | T | A | G | C |
| MK301246.1 | Neuroptera | *Dendroleon pantherinus* | T | G | A | C | T | G | T | G | C | A | A | A | G | G | T | A | G | C |
| KT425073.1 | Neuroptera | *Dilar* sp. | T | G | A | C | T | G | T | G | C | A | A | A | G | G | T | A | G | C |
| FJ859906.1 | Neuroptera | *Ditaxis biseriata* | T | G | A | C | T | G | T | G | C | A | A | A | G | G | T | A | G | C |
| NC013257.1 | Neuroptera | *Ditaxis biseriata* | T | G | A | C | T | G | T | G | C | A | A | A | G | G | T | A | G | C |
| KT425087.1 | Neuroptera | *Drepanepteryx phalaenoides* | T | G | A | C | T | G | T | G | C | A | A | A | G | G | T | A | G | C |
| KF701327.1 | Neuroptera | *Epacanthaclisis banksi* | T | G | A | C | T | G | T | G | C | A | A | A | G | G | T | A | G | C |
| NC025905.1 | Neuroptera | *Epacanthaclisis banksi* | T | G | A | C | T | G | T | G | C | A | A | A | G | G | T | A | G | C |

| | | | | | | | | | | | | | | | | | | | | |
|---|---|---|---|---|---|---|---|---|---|---|---|---|---|---|---|---|---|---|---|---|
| MF972496.1 | Neuroptera | *Euclimacia badia* | T | G | A | C | T | G | T | G | C | A | A | A | G | G | T | A | G | C |
| NC039773.1 | Neuroptera | *Euclimacia badia* | T | G | A | C | T | G | T | G | C | A | A | A | G | G | T | A | G | C |
| KT425080.1 | Neuroptera | *Eumantispa harmandi* | T | G | A | C | T | G | T | G | C | A | A | A | G | G | T | A | G | C |
| MF972495.1 | Neuroptera | *Eumantispa harmandi* | T | G | A | C | T | G | T | G | C | A | A | A | G | G | T | A | G | C |
| NC039772.1 | Neuroptera | *Eumantispa harmandi* | T | G | A | C | T | G | T | G | C | A | A | A | G | G | T | A | G | C |
| KT425072.1 | Neuroptera | *Fontecilla graphicus* | T | G | A | C | T | G | T | G | C | A | A | A | G | G | T | A | G | C |
| KY364372.1 | Neuroptera | *Gatzara jezoensis* | T | G | A | C | T | G | T | G | C | A | A | A | G | G | T | A | G | C |
| MK341582.1 | Neuroptera | *Gryposmylus pennyi* | T | G | A | C | T | G | T | G | C | A | A | A | G | G | T | A | G | C |
| MN852445.1 | Neuroptera | *Hemerobius japonicus* | T | G | A | C | T | G | T | G | C | A | A | A | G | G | T | A | G | C |
| NC048517.1 | Neuroptera | *Hemerobius japonicus* | T | G | A | C | T | G | T | G | C | A | A | A | G | G | T | A | G | C |
| KT425077.1 | Neuroptera | *Heterosmylus* sp. | T | G | A | C | T | G | T | G | C | A | A | A | G | G | T | A | G | C |
| KT425085.1 | Neuroptera | *Inocellia fujiana* | T | G | A | C | T | G | T | G | C | A | A | A | G | G | T | A | G | C |
| KY587200.1 | Neuroptera | *Italochrysa insignis* | T | G | A | C | T | G | T | G | C | A | A | A | G | G | T | A | G | C |
| KY587201.1 | Neuroptera | *Leucochrysa pretiosa* | T | G | A | C | T | G | T | G | C | A | A | A | G | G | T | A | G | C |
| NC015609.1 | Neuroptera | *Libelloides macaronius* | T | G | A | C | T | G | T | G | C | A | A | A | G | G | T | A | G | C |
| MF972494.1 | Neuroptera | *Mantispa japonica* | T | G | A | C | T | G | T | G | C | A | A | A | G | G | T | A | G | C |
| NC039771.1 | Neuroptera | *Mantispa japonica* | T | G | A | C | T | G | T | G | C | A | A | A | G | G | T | A | G | C |
| MK472073.1 | Neuroptera | *Micromus angulatus* | T | G | A | C | T | G | T | G | C | A | A | A | G | G | T | A | G | C |
| NC042677.1 | Neuroptera | *Micromus angulatus* | T | G | A | C | T | G | T | G | C | A | A | A | G | G | T | A | G | C |
| KT425075.1 | Neuroptera | *Micromus* sp. | T | G | A | C | T | G | T | G | C | A | A | A | G | G | T | A | G | C |
| KT425092.1 | Neuroptera | *Mucroberotha vesicaria* | T | G | A | C | T | G | T | G | C | A | A | A | G | G | T | A | G | C |
| KT425089.1 | Neuroptera | *Myiodactylus osmyloides* | T | G | A | C | T | G | T | G | C | A | A | A | G | G | T | A | G | C |

| | | | | | | | | | | | | | | | | | | | | |
|---|---|---|---|---|---|---|---|---|---|---|---|---|---|---|---|---|---|---|---|---|
| MT118664.1 | Neuroptera | *Myrmeleon formicarius* | T | G | A | C | T | G | T | G | C | A | A | A | G | G | T | A | G | C |
| NC047285.1 | Neuroptera | *Myrmeleon formicarius* | T | G | A | C | T | G | T | G | C | A | A | A | G | G | T | A | G | C |
| KJ461323.1 | Neuroptera | *Myrmeleon immanis* | T | G | A | C | T | G | T | G | C | A | A | A | G | G | T | A | G | C |
| NC024826.1 | Neuroptera | *Myrmeleon immanis* | T | G | A | C | T | G | T | G | C | A | A | A | G | G | T | A | G | C |
| KT425071.1 | Neuroptera | *Nallachius americanus* | T | G | A | C | T | G | T | G | C | A | A | A | G | G | T | A | G | C |
| KT425086.1 | Neuroptera | *Negha inflata* | T | G | A | C | T | G | T | G | C | A | A | A | G | G | T | A | G | C |
| KT425079.1 | Neuroptera | *Nemoptera coa* | T | G | A | C | T | G | T | G | C | A | A | A | G | G | T | A | G | C |
| KR078257.1 | Neuroptera | *Neuronema laminatum* | T | G | A | C | T | G | T | G | C | A | A | A | G | G | T | A | G | C |
| NC028153.1 | Neuroptera | *Neuronema laminatum* | T | G | A | C | T | G | T | G | C | A | A | A | G | G | T | A | G | C |
| KT425074.1 | Neuroptera | *Nevrorthus apatelios* | T | G | A | C | T | G | T | G | C | A | A | A | G | G | T | A | G | C |
| KT425076.1 | Neuroptera | *Nipponeurorthus fuscinervis* | T | G | A | C | T | G | T | G | C | A | A | A | G | G | T | A | G | C |
| KP264629.1 | Neuroptera | *Nothancyla verreauxi* | T | G | A | C | T | G | T | G | C | A | A | A | G | G | T | A | G | C |
| KP264630.1 | Neuroptera | *Nothochrysa* sp. | T | G | A | C | T | G | T | G | C | A | A | A | G | G | T | A | G | C |
| KJ461322.1 | Neuroptera | *Nymphes myrmeleonoides* | T | G | A | C | T | G | T | G | C | A | A | A | G | G | T | A | G | C |
| NC024825.1 | Neuroptera | *Nymphes myrmeleonoides* | T | G | A | C | T | G | T | G | C | A | A | A | G | G | T | A | G | C |
| KT425090.1 | Neuroptera | *Oliarces clara* | T | G | A | C | T | G | T | G | C | A | A | A | G | G | T | A | G | C |
| MN818868.1 | Neuroptera | *Osmylus atomatus* | T | G | A | C | T | G | T | G | C | A | A | A | G | G | T | A | G | C |
| MN818867.1 | Neuroptera | *Osmylus fulvicephalus* | T | G | A | C | T | G | T | G | C | A | A | A | G | G | T | A | G | C |
| MN617021.1 | Neuroptera | *Osmylus lucalatus* | T | G | A | C | T | G | T | G | C | A | A | A | G | G | T | A | G | C |
| MN818870.1 | Neuroptera | *Osmylus* sp. | T | G | A | C | T | G | T | G | C | A | A | A | G | G | T | A | G | C |
| MN818869.1 | Neuroptera | *Osmylus zheanus* | T | G | A | C | T | G | T | G | C | A | A | A | G | G | T | A | G | C |
| KY587202.1 | Neuroptera | *Parankylopteryx* sp. | T | G | A | C | T | G | T | G | C | A | A | A | G | G | T | A | G | C |

| | | | | | | | | | | | | | | | | | | | | |
|---|---|---|---|---|---|---|---|---|---|---|---|---|---|---|---|---|---|---|---|---|
| KT425091.1 | Neuroptera | *Podallea* sp. | T | G | A | C | T | G | T | G | C | A | A | A | G | G | T | A | G | C |
| FJ171325.1 | Neuroptera | *Polystoechotes punctatus* | T | G | A | C | T | G | T | G | C | A | A | A | G | G | T | A | G | C |
| NC011278.1 | Neuroptera | *Polystoechotes punctatus* | T | G | A | C | T | G | T | G | C | A | A | A | G | G | T | A | G | C |
| KT425082.1 | Neuroptera | *Psychopsis coelivaga* | T | G | A | C | T | G | T | G | C | A | A | A | G | G | T | A | G | C |
| KF626446.1 | Neuroptera | *Rapisma xizangense* | T | G | A | C | T | G | T | G | C | A | A | A | G | G | T | A | G | C |
| NC023362.1 | Neuroptera | *Rapisma xizangense* | T | G | A | C | T | G | T | G | C | A | A | A | G | G | T | A | G | C |
| KF626447.1 | Neuroptera | *Rapisma zayuanum* | T | G | A | C | T | G | T | G | C | A | A | A | G | G | T | A | G | C |
| NC023363.1 | Neuroptera | *Rapisma zayuanum* | T | G | A | C | T | G | T | G | C | A | A | A | G | G | T | A | G | C |
| KT425067.1 | Neuroptera | *Semidalis aleyrodiformis* | T | G | A | C | T | G | T | G | C | T | A | A | G | G | T | A | G | C |
| KT425070.1 | Neuroptera | *Sisyra nigra* | T | G | A | C | T | G | T | G | C | A | A | A | G | G | T | A | G | C |
| KT425081.1 | Neuroptera | *Stenobiella* sp. | T | G | A | C | T | G | T | G | C | A | A | A | G | G | T | A | G | C |
| MH361300.1 | Neuroptera | *Suhpalacsa longialata* | T | G | A | C | T | G | T | G | C | A | A | A | G | G | T | A | G | C |
| NC039948.1 | Neuroptera | *Suhpalacsa longialata* | T | G | A | C | T | G | T | G | C | A | A | A | G | G | T | A | G | C |
| MK301247.1 | Neuroptera | *Suhpalacsa* sp. | T | G | A | C | T | G | T | G | C | A | A | A | G | G | T | A | G | C |
| MK408756.1 | Neuroptera | *Thaumatosmylus hainanus* | T | G | A | C | T | G | T | G | C | A | A | A | G | G | T | A | G | C |
| NC046578.1 | Neuroptera | *Thaumatosmylus hainanus* | T | G | A | C | T | G | T | G | C | A | A | A | G | G | T | A | G | C |
| KC515397.1 | Neuroptera | *Thyridosmylus langii* | T | G | A | C | T | G | T | G | C | A | A | A | G | G | T | A | G | C |
| NC021415.1 | Neuroptera | *Thyridosmylus langii* | T | G | A | C | T | G | T | G | C | A | A | A | G | G | T | A | G | C |
| MH020192.1 | Coleoptera | *Abscondita anceyi* | T | G | A | C | T | G | T | G | C | A | A | A | G | G | T | A | G | C |
| NC039706.1 | Coleoptera | *Abscondita anceyi* | T | G | A | C | T | G | T | G | C | A | A | A | G | G | T | A | G | C |
| MK122952.1 | Coleoptera | *Abscondita chinensis* | T | G | A | C | T | G | T | G | C | A | A | A | G | G | T | A | G | C |
| MK292092.1 | Coleoptera | *Abscondita terminalis* | T | G | A | C | T | G | T | G | C | A | A | A | G | G | T | A | G | C |

| | | | | | | | | | | | | | | | | | | | | |
|---|---|---|---|---|---|---|---|---|---|---|---|---|---|---|---|---|---|---|---|---|
| NC044776.1 | Coleoptera | *Abscondita terminalis* | T | G | A | C | T | G | T | G | C | A | A | A | G | G | T | A | G | C |
| KT876878.1 | Coleoptera | *Acilius sulcatus* | T | G | A | C | T | G | T | G | C | A | A | A | G | G | T | A | G | C |
| KX758085.1 | Coleoptera | *Aquatica ficta* | T | G | A | C | T | G | T | G | C | A | A | A | G | G | T | A | G | C |
| NC035060.1 | Coleoptera | *Aquatica ficta* | T | G | A | C | T | G | T | G | C | A | A | A | G | G | T | A | G | C |
| NC035755.1 | Coleoptera | *Aquatica lateralis* | T | G | A | C | T | G | T | G | C | A | A | A | G | G | T | A | G | C |
| KF667531.1 | Coleoptera | *Aquatica leii* | T | G | A | C | T | G | T | G | C | A | A | A | G | G | T | A | G | C |
| NC025276.1 | Coleoptera | *Aquatica leii* | T | G | A | C | T | G | T | G | C | A | A | A | G | G | T | A | G | C |
| KX758086.1 | Coleoptera | *Aquatica wuhana* | T | G | A | C | T | G | T | G | C | A | A | A | G | G | T | A | G | C |
| NC035061.1 | Coleoptera | *Aquatica wuhana* | T | G | A | C | T | G | T | G | C | A | A | A | G | G | T | A | G | C |
| KX229747.1 | Coleoptera | *Asymmetricata circumdata* | T | G | A | C | T | G | T | A | C | A | A | A | G | G | T | A | G | C |
| MK292113.1 | Coleoptera | *Asymmetricata circumdata* | T | G | A | C | T | G | T | A | C | A | A | A | G | G | T | A | G | C |
| NC032062.1 | Coleoptera | *Asymmetricata circumdata* | T | G | A | C | T | G | T | A | C | A | A | A | G | G | T | A | G | C |
| KJ922151.1 | Coleoptera | *Bicellonychia lividipennis* | T | A | A | C | T | G | T | G | C | A | A | A | G | G | T | A | G | C |
| NC030060.1 | Coleoptera | *Bicellonychia lividipennis* | T | A | A | C | T | G | T | G | C | A | A | A | G | G | T | A | G | C |
| KT876885.1 | Coleoptera | *Colymbetes fuscus* | T | G | A | C | T | G | T | G | C | A | A | A | G | G | T | A | G | C |
| MK292114.1 | Coleoptera | *Curtos bilineatus* | T | G | A | C | T | G | T | A | C | A | A | A | G | G | T | A | G | C |
| NC044789.1 | Coleoptera | *Curtos bilineatus* | T | G | A | C | T | G | T | A | C | A | A | A | G | G | T | A | G | C |
| MK609965.1 | Coleoptera | *Curtos costipennis* | T | G | A | C | T | G | T | G | C | A | A | A | G | G | T | A | G | C |
| MH651351.1 | Coleoptera | *Diaphanes citrinus* | T | G | A | C | T | G | T | G | C | A | A | A | G | G | T | A | G | C |
| MK292116.1 | Coleoptera | *Diaphanes mendax* | T | G | A | C | T | G | T | G | C | G | A | A | G | G | T | A | G | C |
| NC044791.1 | Coleoptera | *Diaphanes mendax* | T | G | A | C | T | G | T | G | C | G | A | A | G | G | T | A | G | C |
| MK292094.1 | Coleoptera | *Diaphanes nubilus* | T | G | A | C | T | G | T | A | C | A | A | A | G | G | T | A | G | C |

| | | | | | | | | | | | | | | | | | | | | |
|---|---|---|---|---|---|---|---|---|---|---|---|---|---|---|---|---|---|---|---|---|
| NC044787.1 | Coleoptera | *Diaphanes nubilus* | T | G | A | C | T | G | T | A | C | A | A | A | G | G | T | A | G | C |
| MK292118.1 | Coleoptera | *Diaphanes pectinealis* | T | G | A | C | T | G | T | A | C | A | A | A | G | G | T | A | G | C |
| NC044793.1 | Coleoptera | *Diaphanes pectinealis* | T | G | A | C | T | G | T | A | C | A | A | A | G | G | T | A | G | C |
| MK292095.1 | Coleoptera | *Diaphanes* sp. | T | G | A | C | T | G | T | G | C | A | A | A | G | G | T | A | G | C |
| MG242622.1 | Coleoptera | *Ellychnia corrusca* | T | A | A | C | T | G | T | G | C | G | A | A | G | G | T | A | G | C |
| MK292112.1 | Coleoptera | *Emeia pseudosauteri* | T | A | A | C | T | G | T | G | C | A | A | A | G | G | T | A | G | C |
| NC044778.1 | Coleoptera | *Emeia pseudosauteri* | T | A | A | C | T | G | T | G | C | A | A | A | G | G | T | A | G | C |
| KT876897.1 | Coleoptera | *Hydroporus erythrocephalus* | T | G | A | C | T | G | T | G | C | A | A | A | G | G | T | A | G | C |
| KT876896.1 | Coleoptera | *Hydroporus obscurus* | T | G | A | C | T | G | T | G | C | A | A | A | G | G | T | A | G | C |
| KT876901.1 | Coleoptera | *Hygrotus confluens* | T | G | A | C | T | G | T | G | C | A | A | A | G | G | T | A | G | C |
| KT876900.1 | Coleoptera | *Hygrotus impressopunctatus* | T | G | A | C | T | G | T | G | C | A | A | A | G | G | T | A | G | C |
| KT876899.1 | Coleoptera | *Hygrotus inaequalis* | T | G | A | C | T | G | T | G | C | A | A | A | G | G | T | A | G | C |
| KX087299.1 | Coleoptera | *Hygrotus nigrolineatus* | T | G | A | C | T | G | T | G | C | A | A | A | G | G | T | A | G | C |
| MH427718.1 | Coleoptera | *Inflata indica* | T | G | A | C | T | G | T | G | C | A | A | A | G | G | T | A | G | C |
| NC039700.1 | Coleoptera | *Inflata indica* | T | G | A | C | T | G | T | G | C | A | A | A | G | G | T | A | G | C |
| MK292091.1 | Coleoptera | *Lamprigera yunnana* | T | A | A | C | T | G | T | G | C | A | A | A | G | G | T | A | G | C |
| NC044786.1 | Coleoptera | *Lamprigera yunnana* | T | A | A | C | T | G | T | G | C | A | A | A | G | G | T | A | G | C |
| KX087302.1 | Coleoptera | *Lampyris noctiluca* | T | G | A | C | T | G | T | G | C | A | A | A | G | G | T | A | G | C |
| MG912994.1 | Coleoptera | *Limbodessus palmulaoides* | T | G | A | C | T | G | T | G | C | A | A | A | G | G | T | A | G | C |
| NC037749.1 | Coleoptera | *Limbodessus palmulaoides* | T | G | A | C | T | G | T | G | C | A | A | A | G | G | T | A | G | C |
| JQ034416.1 | Coleoptera | *Limnichidae* sp. | T | G | A | C | T | G | T | G | C | A | A | A | G | G | T | A | G | C |
| KT876902.1 | Coleoptera | *Liopterus haemorrhoidalis* | T | G | A | C | T | G | T | G | C | A | A | A | G | G | T | A | G | C |

| | | | | | | | | | | | | | | | | | | | | |
|---|---|---|---|---|---|---|---|---|---|---|---|---|---|---|---|---|---|---|---|---|
| NC022472.1 | Coleoptera | *Luciola cruciata* | T | G | A | C | T | G | T | G | C | A | A | A | G | G | T | A | G | C |
| MG770613.1 | Coleoptera | *Luciola curtithorax* | T | G | A | C | T | G | T | G | C | A | A | A | G | G | T | A | G | C |
| NC038225.1 | Coleoptera | *Luciola curtithorax* | T | G | A | C | T | G | T | G | C | A | A | A | G | G | T | A | G | C |
| KP313820.1 | Coleoptera | *Luciola substriata* | T | G | A | C | T | G | T | G | C | A | A | A | G | G | T | A | G | C |
| NC027176.1 | Coleoptera | *Luciola substriata* | T | G | A | C | T | G | T | G | C | A | A | A | G | G | T | A | G | C |
| FJ859901.1 | Coleoptera | *Macrogyrus oblongus* | T | A | A | C | T | G | T | G | C | A | A | A | G | G | T | A | G | C |
| NC013249.1 | Coleoptera | *Macrogyrus oblongus* | T | A | A | C | T | G | T | G | C | A | A | A | G | G | T | A | G | C |
| MG912995.1 | Coleoptera | *Paroster macrosturtensis* | T | G | A | C | T | G | T | G | C | A | A | A | G | G | T | A | G | C |
| NC037750.1 | Coleoptera | *Paroster macrosturtensis* | T | G | A | C | T | G | T | G | C | A | A | A | G | G | T | A | G | C |
| MG912996.1 | Coleoptera | *Paroster mesosturtensis* | T | G | A | C | T | G | T | G | C | A | A | A | G | G | T | A | G | C |
| NC037751.1 | Coleoptera | *Paroster mesosturtensis* | T | G | A | C | T | G | T | G | C | A | A | A | G | G | T | A | G | C |
| MG912997.1 | Coleoptera | *Paroster microsturtensis* | T | G | A | C | T | G | T | G | C | A | A | A | G | G | T | A | G | C |
| NC037752.1 | Coleoptera | *Paroster microsturtensis* | T | G | A | C | T | G | T | G | C | A | A | A | G | G | T | A | G | C |
| KY778696.1 | Coleoptera | *Photinus pyralis* | T | A | A | C | T | G | T | G | C | G | A | A | G | G | T | A | G | C |
| MF686051.1 | Coleoptera | *Pteroptyx maipo* | T | G | A | C | T | G | T | G | C | A | A | A | G | G | T | A | G | C |
| NC036353.1 | Coleoptera | *Pteroptyx maipo* | T | G | A | C | T | G | T | G | C | A | A | A | G | G | T | A | G | C |
| MK292115.1 | Coleoptera | *Pyrocoelia praetexta* | T | G | A | C | T | G | T | G | C | A | A | A | G | G | T | A | G | C |
| NC044790.1 | Coleoptera | *Pyrocoelia praetexta* | T | G | A | C | T | G | T | G | C | A | A | A | G | G | T | A | G | C |
| AF452048.1 | Coleoptera | *Pyrocoelia rufa* | T | G | A | C | T | G | T | G | C | A | A | A | G | G | T | A | G | C |
| NC003970.1 | Coleoptera | *Pyrocoelia rufa* | T | G | A | C | T | G | T | G | C | A | A | A | G | G | T | A | G | C |
| MK292117.1 | Coleoptera | *Pyrocoelia thibetana* | T | G | A | C | T | G | T | G | C | A | A | A | G | G | T | A | G | C |
| NC044792.1 | Coleoptera | *Pyrocoelia thibetana* | T | G | A | C | T | G | T | G | C | A | A | A | G | G | T | A | G | C |

| | | | | | | | | | | | | | | | | | | | | |
|---|---|---|---|---|---|---|---|---|---|---|---|---|---|---|---|---|---|---|---|---|
| MK292111.1 | Coleoptera | *Vesta saturnalis* | T | G | A | C | T | G | T | A | C | A | A | A | G | G | T | A | G | C |
| NC044788.1 | Coleoptera | *Vesta saturnalis* | T | G | A | C | T | G | T | A | C | A | A | A | G | G | T | A | G | C |
| MF680449.1 | Trichoptera | *Anabolia bimaculata* | T | A | A | C | C | G | T | A | C | A | A | A | G | G | T | A | G | C |
| NC036004.1 | Trichoptera | *Anabolia bimaculata* | T | A | A | C | C | G | T | A | C | A | A | A | G | G | T | A | G | C |
| KF756944.1 | Trichoptera | *Apatania* sp. | T | A | A | C | T | G | T | A | C | A | A | A | G | G | T | A | G | C |
| MT410856.1 | Trichoptera | *Beraea pullata* | T | A | A | T | T | G | T | G | C | A | A | A | G | G | T | A | G | C |
| MG669126.1 | Trichoptera | *Cheumatopsyche analis* | T | A | A | C | T | G | T | A | C | A | A | A | G | G | T | A | G | C |
| NC036955.1 | Trichoptera | *Cheumatopsyche analis* | T | A | A | C | T | G | T | A | C | A | A | A | G | G | T | A | G | C |
| KX385010.1 | Trichoptera | *Cheumatopsyche brevilineata* | T | A | A | C | T | G | T | A | C | A | A | A | G | G | T | A | G | C |
| NC043769.1 | Trichoptera | *Cheumatopsyche brevilineata* | T | A | A | C | T | G | T | A | C | A | A | A | G | G | T | A | G | C |
| MG669125.1 | Trichoptera | *Cheumatopsyche campyla* | T | A | A | C | T | G | T | A | C | A | A | A | G | G | T | A | G | C |
| NC036954.1 | Trichoptera | *Cheumatopsyche campyla* | T | A | A | C | T | G | T | A | C | A | A | A | G | G | T | A | G | C |
| MG669123.1 | Trichoptera | *Cheumatopsyche speciosa* | T | A | A | C | T | G | T | A | C | A | A | A | G | G | T | A | G | C |
| NC036952.1 | Trichoptera | *Cheumatopsyche speciosa* | T | A | A | C | T | G | T | A | C | A | A | A | G | G | T | A | G | C |
| KF717094.1 | Trichoptera | *Eubasilissa regina* | T | A | A | C | C | G | T | A | C | A | A | A | G | G | T | A | G | C |
| KF756943.1 | Trichoptera | *Eubasilissa regina* | T | A | A | C | C | G | T | A | C | A | A | A | G | G | T | A | G | C |
| NC023374.1 | Trichoptera | *Eubasilissa regina* | T | A | A | C | C | G | T | A | C | A | A | A | G | G | T | A | G | C |
| KX385011.1 | Trichoptera | *Hydatophylax nigrovittatus* | T | A | A | C | C | G | T | A | C | A | A | A | G | G | T | A | G | C |
| NC043770.1 | Trichoptera | *Hydatophylax nigrovittatus* | T | A | A | C | C | G | T | A | C | A | A | A | G | G | T | A | G | C |
| KF717095.1 | Trichoptera | *Hydromanicus wulaianus* | T | A | A | C | T | G | T | A | C | A | A | A | G | G | T | A | G | C |
| NC036156.1 | Trichoptera | *Hydromanicus wulaianus* | T | A | A | C | T | G | T | A | C | A | A | A | G | G | T | A | G | C |
| MT628580.1 | Trichoptera | *Hydropsyche angustipennis* | T | A | A | C | T | G | T | A | C | A | A | A | G | G | T | A | G | C |

| | | | | | | | | | | | | | | | | | | | | | |
|---|---|---|---|---|---|---|---|---|---|---|---|---|---|---|---|---|---|---|---|---|---|
| MG669122.1 | Trichoptera | *Hydropsyche orris* | | ? | A | A | C | T | G | T | A | C | A | A | A | G | G | T | A | G | C |
| NC036951.1 | Trichoptera | *Hydropsyche orris* | | ? | A | A | C | T | G | T | A | C | A | A | A | G | G | T | A | G | C |
| KT876876.1 | Trichoptera | *Hydropsyche pellucidula* | | T | A | A | C | T | G | T | A | C | A | A | A | G | G | T | A | G | C |
| NC029246.1 | Trichoptera | *Hydropsyche pellucidula* | | T | A | A | C | T | G | T | A | C | A | A | A | G | G | T | A | G | C |
| MG669121.1 | Trichoptera | *Hydropsyche simulans* | | T | A | A | C | T | G | T | A | C | A | A | A | G | G | T | A | G | C |
| NC036950.1 | Trichoptera | *Hydropsyche simulans* | | T | A | A | C | T | G | T | A | C | A | A | A | G | G | T | A | G | C |
| MK077681.1 | Trichoptera | *Limnephilus hyalinus* | | T | A | A | C | C | G | T | A | C | A | A | A | G | G | T | A | G | C |
| NC044710.1 | Trichoptera | *Limnephilus hyalinus* | | T | A | A | C | C | G | T | A | C | A | A | A | G | G | T | A | G | C |
| MG980616.1 | Trichoptera | *Phryganea cinerea* | | T | A | A | C | C | G | T | A | C | A | A | A | G | G | T | A | G | C |
| NC039714.1 | Trichoptera | *Phryganea cinerea* | | T | A | A | C | C | G | T | A | C | A | A | A | G | G | T | A | G | C |
| KX385012.1 | Trichoptera | *Phryganopsyche latipennis* | | T | A | A | C | C | G | T | A | C | A | A | A | G | G | T | A | G | C |
| NC043771.1 | Trichoptera | *Phryganopsyche latipennis* | | T | A | A | C | C | G | T | A | C | A | A | A | G | G | T | A | G | C |
| MG669124.1 | Trichoptera | *Potamyia flava* | | T | A | A | C | T | G | T | A | C | A | A | A | G | G | T | A | G | C |
| NC036953.1 | Trichoptera | *Potamyia flava* | | T | A | A | C | T | G | T | A | C | A | A | A | G | G | T | A | G | C |
| KP455290.1 | Trichoptera | *Sericostoma personatum* | | T | A | A | T | T | G | T | G | C | A | A | A | G | G | T | A | G | C |
| KP455291.1 | Trichoptera | *Thremma gallicum* | | T | A | A | C | C | G | T | A | C | A | A | A | G | G | T | A | G | C |
| | | | A | 0.0 | 16.7 | 100.0 | 0.0 | 0.0 | 0.0 | 0.0 | 13.8 | 0.0 | 92.5 | 99.6 | 100.0 | 0.0 | 0.0 | 0.0 | 100.0 | 0.0 | 0.0 |
| | | | T | 100.0 | 0.0 | 0.0 | 0.4 | 49.0 | 0.0 | 100.0 | 0.0 | 0.0 | 2.0 | 0.0 | 0.0 | 0.4 | 0.0 | 100.0 | 0.0 | 0.0 | 0.0 |
| | | | G | 0.0 | 83.3 | 0.0 | 0.0 | 0.0 | 100.0 | 0.0 | 86.2 | 0.0 | 5.6 | 0.4 | 0.0 | 99.6 | 100.0 | 0.0 | 0.0 | 100.0 | 0.0 |
| | | | C | 0.0 | 0.0 | 0.0 | 99.6 | 51.0 | 0.0 | 0.0 | 0.0 | 100.0 | 0.0 | 0.0 | 0.0 | 0.0 | 0.0 | 0.0 | 0.0 | 0.0 | 100.0 |

Table S5. Nucleotide sequences of universal primer (AQdb-16S_R) and homologous region of 558 sequences of insects used for development in this study.

| AQdb-16S_R | | | C | C | G | G | T | Y | T | R | A | A | C | T | C | A | R | A | T | C | A | T | G | T |
|---|---|---|---|---|---|---|---|---|---|---|---|---|---|---|---|---|---|---|---|---|---|---|---|---|
| | | A | 0 | 0 | 0 | 19 | 0 | 0 | 0 | 55 | 557 | 557 | 0 | 0 | 0 | 557 | 179 | 557 | 0 | 0 | 557 | 0 | 0 | 0 |
| | | T | 0 | 0 | 0 | 0 | 555 | 197 | 557 | 3 | 0 | 0 | 0 | 557 | 0 | 0 | 0 | 0 | 557 | 0 | 0 | 554 | 0 | 557 |
| | | G | 0 | 0 | 557 | 538 | 0 | 0 | 0 | 499 | 0 | 0 | 0 | 0 | 0 | 0 | 376 | 0 | 0 | 0 | 0 | 0 | 557 | 0 |
| | | C | 557 | 557 | 0 | 0 | 2 | 360 | 0 | 0 | 0 | 0 | 557 | 0 | 557 | 0 | 2 | 0 | 0 | 557 | 0 | 3 | 0 | 0 |
| GenBank Acc. No. | Order | Species | | | | | | | | | | | | | | | | | | | | | | |
| MN481418.1 | Diplura | *Campodea augens* | C | C | G | G | T | C | T | G | A | A | C | T | C | A | A | A | T | C | A | T | G | T |
| DQ529236.1 | Diplura | *Campodea fragilis* | C | C | G | G | T | C | T | G | A | A | C | T | C | A | A | A | T | C | A | T | G | T |
| NC008233.1 | Diplura | *Campodea fragilis* | C | C | G | G | T | C | T | G | A | A | C | T | C | A | A | A | T | C | A | T | G | T |
| DQ529237.1 | Diplura | *Campodea lubbocki* | C | C | G | G | T | C | T | G | A | A | C | T | C | A | C | A | T | C | A | T | G | T |
| NC008234.1 | Diplura | *Campodea lubbocki* | C | C | G | G | T | C | T | G | A | A | C | T | C | A | C | A | T | C | A | T | G | T |
| AY771989.1 | Diplura | *Japyx solifugus* | C | C | G | G | T | C | T | G | A | A | C | T | C | A | G | A | T | C | A | T | G | T |
| NC007214.1 | Diplura | *Japyx solifugus* | C | C | G | G | T | C | T | G | A | A | C | T | C | A | G | A | T | C | A | T | G | T |
| JN990601.1 | Diplura | *Lepidocampa weberi* | C | C | G | A | T | C | T | G | A | A | C | T | C | A | A | A | T | C | A | T | G | T |
| NC022675.1 | Diplura | *Lepidocampa weberi* | C | C | G | A | T | C | T | G | A | A | C | T | C | A | A | A | T | C | A | T | G | T |

| | | | | | | | | | | | | | | | | | | | | | | | | |
|---|---|---|---|---|---|---|---|---|---|---|---|---|---|---|---|---|---|---|---|---|---|---|---|---|
| JN990600.1 | Diplura | *Occasjapyx japonicus* | C | C | G | G | T | C | T | G | A | A | C | T | C | A | G | A | T | C | A | T | G | T |
| NC022674.1 | Diplura | *Occasjapyx japonicus* | C | C | G | G | T | C | T | G | A | A | C | T | C | A | G | A | T | C | A | T | G | T |
| JN990598.1 | Diplura | *Octostigma sinensis* | C | C | G | G | T | C | T | G | A | A | C | T | C | A | A | A | T | C | A | T | G | T |
| NC022672.1 | Diplura | *Octostigma sinensis* | C | C | G | G | T | C | T | G | A | A | C | T | C | A | A | A | T | C | A | T | G | T |
| JN990599.1 | Diplura | *Parajapyx emeryanus* | C | C | G | G | T | C | T | G | A | A | C | T | C | A | G | A | T | C | A | T | G | T |
| NC022673.1 | Diplura | *Parajapyx emeryanus* | C | C | G | G | T | C | T | G | A | A | C | T | C | A | G | A | T | C | A | T | G | T |
| KJ754500.1 | Archaeognatha | *Allopsontus* sp. | C | C | G | A | T | T | T | G | A | A | C | T | C | A | G | A | T | C | A | T | G | T |
| KJ754501.1 | Archaeognatha | *Allopsontus* sp. | C | C | G | G | T | C | T | G | A | A | C | T | C | A | G | A | T | C | A | T | G | T |
| AY793551.1 | Archaeognatha | *Nesomachilis australica* | C | C | G | G | T | C | T | G | A | A | C | T | C | A | G | A | T | C | A | T | G | T |
| NC006895.1 | Archaeognatha | *Nesomachilis australica* | C | C | G | G | T | C | T | G | A | A | C | T | C | A | G | A | T | C | A | T | G | T |
| KJ754502.1 | Archaeognatha | *Pedetontinus luanchuanensis* | C | C | G | G | T | T | T | G | A | A | C | T | C | A | G | A | T | C | A | T | G | T |
| EU621793.1 | Archaeognatha | *Pedetontus silvestrii* | C | C | G | G | T | C | T | G | A | A | C | T | C | A | G | A | T | C | A | T | G | T |

| | | | | | | | | | | | | | | | | | | | | | | | | |
|---|---|---|---|---|---|---|---|---|---|---|---|---|---|---|---|---|---|---|---|---|---|---|---|---|
| NC011717.1 | Archaeognatha | *Pedetontus silvestrii* | C | C | G | G | T | C | T | G | A | A | C | T | C | A | G | A | T | C | A | T | G | T |
| KJ754503.1 | Archaeognatha | *Petrobiellus* sp. | C | C | G | G | T | C | T | G | A | A | C | T | C | A | G | A | T | C | A | T | G | T |
| KJ754504.1 | Archaeognatha | *Petrobiellus* sp. | C | C | G | G | T | C | T | G | A | A | C | T | C | A | G | A | T | C | A | T | G | T |
| AY956355.1 | Archaeognatha | *Petrobius brevistylis* | C | C | G | G | T | T | T | G | A | A | C | T | C | A | G | A | T | C | A | T | G | T |
| NC007688.1 | Archaeognatha | *Petrobius brevistylis* | C | C | G | G | T | T | T | G | A | A | C | T | C | A | G | A | T | C | A | T | G | T |
| JX308221.1 | Archaeognatha | *Songmachilis xinxiangensis* | C | C | G | G | T | T | T | G | A | A | C | T | C | A | G | A | T | C | A | T | G | T |
| NC021384.1 | Archaeognatha | *Songmachilis xinxiangensis* | C | C | G | G | T | T | T | G | A | A | C | T | C | A | G | A | T | C | A | T | G | T |
| EU016193.1 | Archaeognatha | *Trigoniophthalmus alternatus* | C | C | G | G | T | C | T | G | A | A | C | T | C | A | G | A | T | C | A | T | G | T |
| NC010532.1 | Archaeognatha | *Trigoniophthalmus alternatus* | C | C | G | G | T | C | T | G | A | A | C | T | C | A | G | A | T | C | A | T | G | T |
| EU084035.1 | Zygentoma | *Atelura formicaria* | C | C | G | G | T | T | T | G | A | A | C | T | C | A | A | A | T | C | A | T | G | T |
| NC011197.1 | Zygentoma | *Atelura formicaria* | C | C | G | G | T | T | T | G | A | A | C | T | C | A | A | A | T | C | A | T | G | T |
| MK301436.1 | Zygentoma | *Ctenolepisma villosa* | C | C | G | A | T | C | T | G | A | A | C | T | C | A | G | A | T | C | A | T | G | T |
| NC046478.1 | Zygentoma | *Ctenolepisma villosa* | C | C | G | A | T | C | T | G | A | A | C | T | C | A | G | A | T | C | A | T | G | T |

| | | | | | | | | | | | | | | | | | | | | | | | | |
|---|---|---|---|---|---|---|---|---|---|---|---|---|---|---|---|---|---|---|---|---|---|---|---|---|
| MT108230.1 | Zygentoma | *Lepisma saccharina* | C | C | G | A | T | C | T | G | A | A | C | T | C | A | G | A | T | C | A | T | G | T |
| NC047445.1 | Zygentoma | *Lepisma saccharina* | C | C | G | A | T | C | T | G | A | A | C | T | C | A | G | A | T | C | A | T | G | T |
| AY639935.1 | Zygentoma | *Thermobia domestica* | C | C | G | A | T | C | T | G | A | A | C | T | C | A | G | A | T | C | A | T | G | T |
| NC006080.1 | Zygentoma | *Thermobia domestica* | C | C | G | A | T | C | T | G | A | A | C | T | C | A | G | A | T | C | A | T | G | T |
| MK951657.1 | Zygentoma | *Thermobia* sp. | C | C | G | A | T | C | T | G | A | A | C | T | C | A | G | A | T | C | A | T | G | T |
| AY191994.1 | Zygentoma | *Tricholepidion gertschi* | C | C | G | G | T | T | T | G | A | A | C | T | C | A | G | A | T | C | A | T | G | T |
| NC005437.1 | Zygentoma | *Tricholepidion gertschi* | C | C | G | G | T | T | T | G | A | A | C | T | C | A | G | A | T | C | A | T | G | T |
| GU479735.1 | Ephemeroptera | *Alainites yixiani* | C | C | G | A | T | T | T | G | A | A | C | T | C | A | G | A | T | C | A | T | G | T |
| GU936204.1 | Ephemeroptera | *Baetis* sp. | C | C | G | A | T | T | T | G | A | A | C | T | C | A | G | A | T | C | A | T | G | T |
| MT628575.1 | Ephemeroptera | *Caenis robusta* | C | C | G | G | T | C | T | G | A | A | C | T | C | A | G | A | T | C | A | T | G | T |
| GQ502451.1 | Ephemeroptera | *Caenis* sp. | C | C | G | G | T | C | T | G | A | A | C | T | C | A | G | A | T | C | A | T | G | T |
| MG910499.1 | Ephemeroptera | *Caenis* sp. | C | C | G | G | T | C | T | G | A | A | C | T | C | A | G | A | T | C | A | T | G | T |
| MN356096.1 | Ephemeroptera | *Caenis* sp. | C | C | G | G | T | C | T | G | A | A | C | T | C | A | G | A | T | C | A | T | G | T |
| MN807287.1 | Ephemeroptera | *Choroterpides apiculata* | C | C | G | G | T | T | T | G | A | A | C | T | C | A | G | A | T | C | A | T | G | T |
| MG870104.1 | Ephemeroptera | *Epeorus herklotsi* | C | C | G | G | T | C | T | G | A | A | C | T | C | A | G | A | T | C | A | T | G | T |

| | | | | | | | | | | | | | | | | | | | | | | | | |
|---|---|---|---|---|---|---|---|---|---|---|---|---|---|---|---|---|---|---|---|---|---|---|---|---|
| MH752075.1 | Ephemeroptera | *Epeorus herklotsi* | C | C | G | G | T | C | T | G | A | A | C | T | C | A | G | A | T | C | A | T | G | T |
| NC039612.1 | Ephemeroptera | *Epeorus herklotsi* | C | C | G | G | T | C | T | G | A | A | C | T | C | A | G | A | T | C | A | T | G | T |
| KJ493406.1 | Ephemeroptera | *Epeorus* sp. | C | C | G | G | T | C | T | G | A | A | C | T | C | A | G | A | T | C | A | T | G | T |
| KM244708.1 | Ephemeroptera | *Epeorus* sp. | C | C | G | G | T | C | T | G | A | A | C | T | C | A | G | A | T | C | A | T | G | T |
| MK951674.1 | Ephemeroptera | *Epeorus* sp. | C | C | G | G | T | C | T | G | A | A | C | T | C | A | G | A | T | C | A | T | G | T |
| EU591678.1 | Ephemeroptera | *Ephemera orientalis* | C | C | G | G | T | C | T | G | A | A | C | T | C | A | G | A | T | C | A | T | G | T |
| NC012645.1 | Ephemeroptera | *Ephemera orientalis* | C | C | G | G | T | C | T | G | A | A | C | T | C | A | G | A | T | C | A | T | G | T |
| MK951659.1 | Ephemeroptera | *Ephemera* sp. | C | C | G | G | T | C | T | G | A | A | C | T | C | A | G | A | T | C | A | T | G | T |
| KM244691.1 | Ephemeroptera | *Ephemerella* sp. | C | C | G | G | T | C | T | G | A | A | C | T | C | A | G | A | T | C | A | T | G | T |
| MT274127.1 | Ephemeroptera | *Ephemerella* sp. | C | C | G | G | T | C | T | G | A | A | C | T | C | A | G | A | T | C | A | T | G | T |
| NC050279.1 | Ephemeroptera | *Ephemerella* sp. | C | C | G | G | T | C | T | G | A | A | C | T | C | A | G | A | T | C | A | T | G | T |
| GU936203.1 | Ephemeroptera | *Habrophlebiodes zijinensis* | C | C | G | G | T | T | T | G | A | A | C | T | C | A | G | A | T | C | A | T | G | T |
| HM143892.1 | Ephemeroptera | *Isonychia ignota* | C | C | G | G | T | C | T | G | A | A | C | T | C | A | G | A | T | C | A | T | G | T |
| MH119135.1 | Ephemeroptera | *Isonychia kiangsinensis* | C | C | G | G | T | C | T | G | A | A | C | T | C | A | G | A | T | C | A | T | G | T |
| NC042163.1 | Ephemeroptera | *Isonychia kiangsinensis* | C | C | G | G | T | C | T | G | A | A | C | T | C | A | G | A | T | C | A | T | G | T |
| MK951658.1 | Ephemeroptera | *Isonychia* sp. | C | C | G | G | T | C | T | G | A | A | C | T | C | A | G | A | T | C | A | T | G | T |
| HM004123.1 | Ephemeroptera | *Paegniodes* | C | C | G | G | T | C | T | G | A | A | C | T | C | A | G | A | T | C | A | T | G | T |

| | | | | | | | | | | | | | | | | | | | | | | | | |
|---|---|---|---|---|---|---|---|---|---|---|---|---|---|---|---|---|---|---|---|---|---|---|---|---|
| | | *cupulatus* | | | | | | | | | | | | | | | | | | | | | | |
| MK951673.1 | Ephemeroptera | *Parafronurus* sp. | C | C | G | G | T | C | T | G | A | A | C | T | C | A | G | A | T | C | A | T | G | T |
| EU349015.1 | Ephemeroptera | *Parafronurus youi* | C | C | G | G | T | C | T | G | A | A | C | T | C | A | G | A | T | C | A | T | G | T |
| NC011359.1 | Ephemeroptera | *Parafronurus youi* | C | C | G | G | T | C | T | G | A | A | C | T | C | A | G | A | T | C | A | T | G | T |
| KM244674.1 | Ephemeroptera | *Potamanthus* sp. | C | C | G | G | T | C | T | G | A | A | C | T | C | A | G | A | T | C | A | T | G | T |
| MK951670.1 | Ephemeroptera | *Rhithrogena* sp. | C | C | G | G | T | C | T | G | A | A | C | T | C | A | G | A | T | C | A | T | G | T |
| MT274128.1 | Ephemeroptera | *Serratella* sp. | C | C | G | G | T | C | T | G | A | A | C | T | C | A | G | A | T | C | A | T | G | T |
| MT274129.1 | Ephemeroptera | *Serratella* sp. | C | C | G | G | T | C | T | G | A | A | C | T | C | A | G | A | T | C | A | T | G | T |
| NC050280.1 | Ephemeroptera | *Serratella* sp. | C | C | G | G | T | C | T | G | A | A | C | T | C | A | G | A | T | C | A | T | G | T |
| NC050281.1 | Ephemeroptera | *Serratella* sp. | C | C | G | G | T | C | T | G | A | A | C | T | C | A | G | A | T | C | A | T | G | T |
| MT274130.1 | Ephemeroptera | *Serratella zapekinae* | C | C | G | G | T | C | T | G | A | A | C | T | C | A | G | A | T | C | A | T | G | T |
| NC050282.1 | Ephemeroptera | *Serratella zapekinae* | C | C | G | G | T | C | T | G | A | A | C | T | C | A | G | A | T | C | A | T | G | T |
| FJ606783.1 | Ephemeroptera | *Siphlonurus immanis* | C | C | G | G | T | C | T | G | A | A | C | T | C | A | G | A | T | C | A | T | G | T |
| NC013822.1 | Ephemeroptera | *Siphlonurus immanis* | C | C | G | G | T | C | T | G | A | A | C | T | C | A | G | A | T | C | A | T | G | T |
| KM244684.1 | Ephemeroptera | *Siphlonurus* sp. | C | C | G | G | T | C | T | G | A | A | C | T | C | A | G | A | T | C | A | T | G | T |
| HQ875717.1 | Ephemeroptera | *Siphluriscus chinensis* | C | C | G | G | T | C | T | G | A | A | C | T | C | A | G | A | T | C | A | T | G | T |
| MT274131.1 | Ephemeroptera | *Torleya* | C | C | G | G | T | C | T | G | A | A | C | T | C | A | G | A | T | C | A | T | G | T |

| | | | | | | | | | | | | | | | | | | | | | | | | |
|---|---|---|---|---|---|---|---|---|---|---|---|---|---|---|---|---|---|---|---|---|---|---|---|---|
| | | *grandiforceps* | | | | | | | | | | | | | | | | | | | | | | |
| NC050283.1 | Ephemeroptera | *Torleya grandiforceps* | C | C | G | G | T | C | T | G | A | A | C | T | C | A | G | A | T | C | A | T | G | T |
| MT274132.1 | Ephemeroptera | *Torleya nepalica* | C | C | G | G | T | C | T | G | A | A | C | T | C | A | G | A | T | C | A | T | G | T |
| NC050284.1 | Ephemeroptera | *Torleya nepalica* | C | C | G | G | T | C | T | G | A | A | C | T | C | A | G | A | T | C | A | T | G | T |
| HM067837.1 | Ephemeroptera | *Vietnamella dabieshanensis* | C | C | G | G | T | C | T | G | A | A | C | T | C | A | G | A | T | C | A | T | G | T |
| KM244655.1 | Ephemeroptera | *Vietnamella* sp. | C | C | G | G | T | C | T | G | A | A | C | T | C | A | G | A | T | C | A | T | G | T |
| MH751435.1 | Odonata | *Acisoma panorpoides* | C | C | G | G | T | C | T | G | A | A | C | T | C | A | G | A | T | C | A | T | G | T |
| NC042689.1 | Odonata | *Acisoma panorpoides* | C | C | G | G | T | C | T | G | A | A | C | T | C | A | G | A | T | C | A | T | G | T |
| KX161841.1 | Odonata | *Anax imperator* | C | C | G | G | T | T | T | G | A | A | C | T | C | A | G | A | T | C | A | T | G | T |
| NC031821.1 | Odonata | *Anax imperator* | C | C | G | G | T | T | T | G | A | A | C | T | C | A | G | A | T | C | A | T | G | T |
| KP233805.1 | Odonata | *Atrocalopteryx atrata* | C | C | G | G | T | C | T | G | A | A | C | T | C | A | G | A | T | C | A | T | G | T |
| NC027181.1 | Odonata | *Atrocalopteryx atrata* | C | C | G | G | T | C | T | G | A | A | C | T | C | A | G | A | T | C | A | T | G | T |
| MG011692.1 | Odonata | *Atrocalopteryx melli* | C | C | G | G | T | C | T | G | A | A | C | T | C | A | G | A | T | C | A | T | G | T |
| NC036749.1 | Odonata | *Atrocalopteryx melli* | C | C | G | G | T | C | T | G | A | A | C | T | C | A | G | A | T | C | A | T | G | T |

| | | | | | | | | | | | | | | | | | | | | | | | | |
|---|---|---|---|---|---|---|---|---|---|---|---|---|---|---|---|---|---|---|---|---|---|---|---|---|
| KM658172.1 | Odonata | *Brachythemis contaminata* | C | C | G | G | T | C | T | G | A | A | C | T | C | A | G | A | T | C | A | T | G | T |
| NC026305.1 | Odonata | *Brachythemis contaminata* | C | C | G | G | T | C | T | G | A | A | C | T | C | A | G | A | T | C | A | T | G | T |
| EU591677.1 | Odonata | *Davidius lunatus* | C | C | G | G | T | T | T | G | A | A | C | T | C | A | G | A | T | C | A | T | G | T |
| NC012644.1 | Odonata | *Davidius lunatus* | C | C | G | G | T | T | T | G | A | A | C | T | C | A | G | A | T | C | A | T | G | T |
| MH751436.1 | Odonata | *Deielia phaon* | C | C | G | G | T | T | T | G | A | A | C | T | C | A | G | A | T | C | A | T | G | T |
| NC042690.1 | Odonata | *Deielia phaon* | C | C | G | G | T | T | T | G | A | A | C | T | C | A | G | A | T | C | A | T | G | T |
| MF716899.1 | Odonata | *Enallagma cyathigerum* | C | C | G | G | T | C | T | G | A | A | C | T | C | A | G | A | T | C | A | T | G | T |
| JX050223.1 | Odonata | *Epiophlebia superstes* | C | C | G | G | T | T | T | G | A | A | C | T | C | A | G | A | T | C | A | T | G | T |
| NC023232.1 | Odonata | *Epiophlebia superstes* | C | C | G | G | T | T | T | G | A | A | C | T | C | A | G | A | T | C | A | T | G | T |
| MK522522.1 | Odonata | *Epophthalmia elegans* | C | C | G | G | T | C | T | G | A | A | C | T | C | A | G | A | T | C | A | T | G | T |
| NC046756.1 | Odonata | *Epophthalmia elegans* | C | C | G | G | T | C | T | G | A | A | C | T | C | A | G | A | T | C | A | T | G | T |
| HM126547.1 | Odonata | *Euphaea formosa* | C | C | G | G | T | C | T | G | A | A | C | T | C | A | G | A | T | C | A | T | G | T |
| NC014493.1 | Odonata | *Euphaea formosa* | C | C | G | G | T | C | T | G | A | A | C | T | C | A | G | A | T | C | A | T | G | T |
| KM244659.1 | Odonata | *Hydrobasileus croceus* | C | C | G | G | T | C | T | G | A | A | C | T | C | A | G | A | T | C | A | T | G | T |

| | | | | | | | | | | | | | | | | | | | | | | |
|---|---|---|---|---|---|---|---|---|---|---|---|---|---|---|---|---|---|---|---|---|---|---|
| NC025758.1 | Odonata | *Hydrobasileus croceus* | C | C | G | G | T | C | T | G | A | A | C | T | C | A | G | A | T | C | A | T | G | T |
| KM244673.1 | Odonata | *Ictinogomphus* sp. | C | C | G | G | T | T | T | G | A | A | C | T | C | A | G | A | T | C | A | T | G | T |
| KU958378.1 | Odonata | *Ischnura elegans* | C | C | G | G | T | C | T | G | A | A | C | T | C | A | G | A | T | C | A | T | G | T |
| MK951668.1 | Odonata | *Ischnura elegans* | C | C | G | G | T | C | T | G | A | A | C | T | C | A | G | A | T | C | A | T | G | T |
| NC031824.1 | Odonata | *Ischnura elegans* | C | C | G | G | T | C | T | G | A | A | C | T | C | A | G | A | T | C | A | T | G | T |
| KC878732.1 | Odonata | *Ischnura pumilio* | C | C | G | G | T | C | T | G | A | A | C | T | C | A | G | A | T | C | A | T | G | T |
| NC021617.1 | Odonata | *Ischnura pumilio* | C | C | G | G | T | C | T | G | A | A | C | T | C | A | G | A | T | C | A | T | G | T |
| MF990748.1 | Odonata | *Macromia daimoji* | C | C | G | G | T | C | T | G | A | A | C | T | C | A | G | A | T | C | A | T | G | T |
| NC041425.1 | Odonata | *Macromia daimoji* | C | C | G | G | T | C | T | G | A | A | C | T | C | A | G | A | T | C | A | T | G | T |
| MK722304.1 | Odonata | *Matrona basilaris* | C | C | G | G | T | C | T | G | A | A | C | T | C | A | G | A | T | C | A | T | G | T |
| MK722305.1 | Odonata | *Matrona basilaris* | C | C | G | G | T | C | T | G | A | A | C | T | C | A | G | A | T | C | A | T | G | T |
| KU958377.1 | Odonata | *Megaloprepus caerulatus* | C | C | G | G | T | C | T | G | A | A | C | T | C | A | G | A | T | C | A | T | G | T |
| NC031823.1 | Odonata | *Megaloprepus caerulatus* | C | C | G | G | T | C | T | G | A | A | C | T | C | A | G | A | T | C | A | T | G | T |
| KU871065.1 | Odonata | *Mnais costalis* | C | C | G | G | T | C | T | G | A | A | C | T | C | A | G | A | T | C | A | T | G | T |
| KY402222.1 | Odonata | *Nannophya pygmaea* | C | C | G | G | T | C | T | G | A | A | C | T | C | A | G | A | T | C | A | T | G | T |
| NC039410.1 | Odonata | *Nannophya pygmaea* | C | C | G | G | T | C | T | G | A | A | C | T | C | A | G | A | T | C | A | T | G | T |
| KU361233.1 | Odonata | *Orthetrum chrysis* | C | C | G | G | T | C | T | G | A | A | C | T | C | A | G | A | T | C | A | T | G | T |

| | | | | | | | | | | | | | | | | | | | | | | | | |
|---|---|---|---|---|---|---|---|---|---|---|---|---|---|---|---|---|---|---|---|---|---|---|---|---|
| NC032048.1 | Odonata | *Orthetrum chrysis* | C | C | G | G | T | C | T | G | A | A | C | T | C | A | G | A | T | C | A | T | G | T |
| KU361232.1 | Odonata | *Orthetrum glaucum* | C | C | G | G | T | C | T | G | A | A | C | T | C | A | G | A | T | C | A | T | G | T |
| NC032047.1 | Odonata | *Orthetrum glaucum* | C | C | G | G | T | C | T | G | A | A | C | T | C | A | G | A | T | C | A | T | G | T |
| MH751437.1 | Odonata | *Orthetrum melania* | C | C | G | G | T | C | T | G | A | A | C | T | C | A | G | A | T | C | A | T | G | T |
| NC042732.1 | Odonata | *Orthetrum melania* | C | C | G | G | T | C | T | G | A | A | C | T | C | A | G | A | T | C | A | T | G | T |
| KU361234.1 | Odonata | *Orthetrum sabina* | C | C | G | G | T | C | T | G | A | A | C | T | C | A | G | A | T | C | A | T | G | T |
| MH751438.1 | Odonata | *Orthetrum sabina* | C | C | G | G | T | C | T | G | A | A | C | T | C | A | G | A | T | C | A | T | G | T |
| NC032049.1 | Odonata | *Orthetrum sabina* | C | C | G | G | T | C | T | G | A | A | C | T | C | A | G | A | T | C | A | T | G | T |
| KU361235.1 | Odonata | *Orthetrum testaceum* | C | C | G | G | T | C | T | G | A | A | C | T | C | A | G | A | T | C | A | T | G | T |
| MH751439.1 | Odonata | *Orthetrum testaceum* | C | C | G | G | T | C | T | G | A | A | C | T | C | A | G | A | T | C | A | T | G | T |
| NC032050.1 | Odonata | *Orthetrum testaceum* | C | C | G | G | T | C | T | G | A | A | C | T | C | A | G | A | T | C | A | T | G | T |
| AB126005.1 | Odonata | *Orthetrum triangulare melania* | C | C | G | G | T | C | T | G | A | A | C | T | C | A | G | A | T | C | A | T | G | T |
| MK951669.1 | Odonata | *Paracercion v-nigrum* | C | C | G | G | T | C | T | G | A | A | C | T | C | A | G | A | T | C | A | T | G | T |
| KP233804.1 | Odonata | *Platycnemis* | C | C | G | G | T | C | T | G | A | A | C | T | C | A | G | A | T | C | A | T | G | T |

| | | | | | | | | | | | | | | | | | | | | | | | | |
|---|---|---|---|---|---|---|---|---|---|---|---|---|---|---|---|---|---|---|---|---|---|---|---|---|
| | | *foliacea* | | | | | | | | | | | | | | | | | | | | | | |
| NC027180.1 | Odonata | *Platycnemis foliacea* | C | C | G | G | T | C | T | G | A | A | C | T | C | A | G | A | T | C | A | T | G | T |
| FJ606784.1 | Odonata | *Pseudolestes mirabilis* | C | C | G | G | T | C | T | G | A | A | C | T | C | A | G | A | T | C | A | T | G | T |
| NC020636.1 | Odonata | *Pseudolestes mirabilis* | C | C | G | G | T | C | T | G | A | A | C | T | C | A | G | A | T | C | A | T | G | T |
| MF150044.1 | Odonata | *Psolodesmus mandarinus* | C | C | G | G | T | C | T | G | A | A | C | T | C | A | G | A | T | C | A | T | G | T |
| NC042442.1 | Odonata | *Psolodesmus mandarinus* | C | C | G | G | T | C | T | G | A | A | C | T | C | A | G | A | T | C | A | T | G | T |
| MG594801.1 | Odonata | *Somatochlora hineana* | C | C | G | G | T | C | T | G | A | A | C | T | C | A | G | A | T | C | A | T | G | T |
| MH751440.1 | Odonata | *Tramea virginia* | C | C | G | G | T | C | T | G | A | A | C | T | C | A | G | A | T | C | A | T | G | T |
| NC042691.1 | Odonata | *Tramea virginia* | C | C | G | G | T | C | T | G | A | A | C | T | C | A | G | A | T | C | A | T | G | T |
| MH751441.1 | Odonata | *Trigomphus carus* | C | C | G | G | T | C | T | G | A | A | C | T | C | A | G | A | T | C | A | T | G | T |
| NC042733.1 | Odonata | *Trigomphus carus* | C | C | G | G | T | C | T | G | A | A | C | T | C | A | G | A | T | C | A | T | G | T |
| JX050224.1 | Odonata | *Vestalis melania* | C | C | G | G | T | C | T | G | A | A | C | T | C | A | G | A | T | C | A | T | G | T |
| NC023233.1 | Odonata | *Vestalis melania* | C | C | G | G | T | C | T | G | A | A | C | T | C | A | G | A | T | C | A | T | G | T |
| KM199685.1 | Plecoptera | *Acroneuria hainana* | C | C | G | G | T | C | T | G | A | A | C | T | C | A | G | A | T | C | A | T | G | T |
| NC026104.1 | Plecoptera | *Acroneuria* | C | C | G | G | T | C | T | G | A | A | C | T | C | A | G | A | T | C | A | T | G | T |

| | | | | | | | | | | | | | | | | | | | | | | | | |
|---|---|---|---|---|---|---|---|---|---|---|---|---|---|---|---|---|---|---|---|---|---|---|---|---|
| | | *hainana* | | | | | | | | | | | | | | | | | | | | | | |
| KX091846.1 | Plecoptera | *Agnetina aequalis* | C | C | G | G | T | C | T | G | A | A | C | T | C | A | G | A | T | C | A | T | G | T |
| MH085446.1 | Plecoptera | *Amphinemura longispina* | C | C | G | G | T | C | T | G | A | A | C | T | C | A | G | A | T | C | A | T | G | T |
| NC044748.1 | Plecoptera | *Amphinemura longispina* | C | C | G | G | T | C | T | G | A | A | C | T | C | A | G | A | T | C | A | T | G | T |
| KX091847.1 | Plecoptera | *Amphinemura* sp. | C | C | G | G | T | C | T | G | A | A | C | T | C | A | G | A | T | C | A | T | G | T |
| MH085447.1 | Plecoptera | *Amphinemura yao* | C | C | G | G | T | C | T | G | A | A | C | T | C | A | G | A | T | C | A | T | G | T |
| NC044749.1 | Plecoptera | *Amphinemura yao* | C | C | G | G | T | C | T | G | A | A | C | T | C | A | G | A | T | C | A | T | G | T |
| MK111413.1 | Plecoptera | *Antarctoperla michaelseni* | C | C | G | G | T | C | T | G | A | A | C | T | C | A | G | A | T | C | A | T | G | T |
| NC042199.1 | Plecoptera | *Antarctoperla michaelseni* | C | C | G | G | T | C | T | G | A | A | C | T | C | A | G | A | T | C | A | T | G | T |
| KR604721.1 | Plecoptera | *Apteroperla tikumana* | C | C | G | G | T | C | T | G | A | A | C | T | C | A | G | A | T | C | A | T | G | T |
| NC027698.1 | Plecoptera | *Apteroperla tikumana* | C | C | G | G | T | C | T | G | A | A | C | T | C | A | G | A | T | C | A | T | G | T |
| MT410838.1 | Plecoptera | *Brachyptera risi* | C | C | G | G | T | C | T | G | A | A | C | T | C | A | G | A | T | C | A | T | G | T |
| MG677941.1 | Plecoptera | *Calineuria stigmatica* | C | C | G | G | T | C | T | G | A | A | C | T | C | A | G | A | T | C | A | T | G | T |
| KX094942.1 | Plecoptera | *Capnia zijinshana* | C | C | G | G | T | C | T | G | A | A | C | T | C | A | G | A | T | C | A | T | G | T |
| NC034661.1 | Plecoptera | *Capnia zijinshana* | C | C | G | G | T | C | T | G | A | A | C | T | C | A | G | A | T | C | A | T | G | T |

| | | | | | | | | | | | | | | | | | | | | | | | | |
|---|---|---|---|---|---|---|---|---|---|---|---|---|---|---|---|---|---|---|---|---|---|---|---|---|
| MG677942.1 | Plecoptera | *Caroperla siveci* | C | C | G | G | T | C | T | G | A | A | C | T | C | A | G | A | T | C | A | T | G | T |
| MF100783.1 | Plecoptera | *Cerconychia flectospina* | C | C | G | G | T | C | T | G | A | A | C | T | C | A | G | A | T | C | A | T | G | T |
| MN419914.1 | Plecoptera | *Claassenia* sp. | C | C | G | G | T | C | T | G | A | A | C | T | C | A | G | A | T | C | A | T | G | T |
| KC952026.1 | Plecoptera | *Cryptoperla* sp. | C | C | G | G | T | C | T | G | A | A | C | T | C | A | G | A | T | C | A | T | G | T |
| MK111416.1 | Plecoptera | *Diamphipnoa annulata* | C | C | G | G | T | C | T | G | A | A | C | T | C | A | G | A | T | C | A | T | G | T |
| NC042205.1 | Plecoptera | *Diamphipnoa annulata* | C | C | G | G | T | C | T | G | A | A | C | T | C | A | G | A | T | C | A | T | G | T |
| MK111417.1 | Plecoptera | *Diamphipnopsis* sp. | C | C | G | G | T | C | T | G | A | A | C | T | C | A | G | A | T | C | A | T | G | T |
| KF484757.1 | Plecoptera | *Dinocras cephalotes* | C | C | G | G | T | C | T | G | A | A | C | T | C | A | G | A | T | C | A | T | G | T |
| NC022843.1 | Plecoptera | *Dinocras cephalotes* | C | C | G | G | T | C | T | G | A | A | C | T | C | A | G | A | T | C | A | T | G | T |
| MG589787.1 | Plecoptera | *Doddsia occidentalis* | C | C | G | G | T | C | T | G | A | A | C | T | C | A | G | A | T | C | A | T | G | T |
| MK905888.1 | Plecoptera | *Etrocorema hochii* | C | C | G | G | T | C | T | G | A | A | C | T | C | A | G | A | T | C | A | T | G | T |
| MK905206.1 | Plecoptera | *Flavoperla* sp. | C | C | G | G | T | C | T | G | A | A | C | T | C | A | G | A | T | C | A | C | G | T |
| MN419916.1 | Plecoptera | *Flavoperla* sp. | C | C | G | G | T | C | T | G | A | A | C | T | C | A | G | A | T | C | A | T | G | T |
| MN419915.1 | Plecoptera | *Indonemoura auriformis* | C | C | G | G | T | C | T | G | A | A | C | T | C | A | G | A | T | C | A | T | G | T |

| | | | | | | | | | | | | | | | | | | | | | | |
|---|---|---|---|---|---|---|---|---|---|---|---|---|---|---|---|---|---|---|---|---|---|---|
| MH085448.1 | Plecoptera | *Indonemoura jacobsoni* | C | C | G | G | T | C | T | G | A | A | C | T | C | A | G | A | T | C | A | T | G | T |
| NC044750.1 | Plecoptera | *Indonemoura jacobsoni* | C | C | G | G | T | C | T | G | A | A | C | T | C | A | G | A | T | C | A | T | G | T |
| MH085449.1 | Plecoptera | *Indonemoura nohirae* | C | C | G | G | T | C | T | G | A | A | C | T | C | A | G | A | T | C | A | T | G | T |
| NC044751.1 | Plecoptera | *Indonemoura nohirae* | C | C | G | G | T | C | T | G | A | A | C | T | C | A | G | A | T | C | A | T | G | T |
| MF716959.1 | Plecoptera | *Isoperla bilineata* | C | C | G | G | T | C | T | G | A | A | C | T | C | A | G | A | T | C | A | T | G | T |
| NC038190.1 | Plecoptera | *Isoperla bilineata* | C | C | G | G | T | C | T | G | A | A | C | T | C | A | G | A | T | C | A | T | G | T |
| MG910457.1 | Plecoptera | *Isoperla eximia* | C | C | G | G | T | C | T | G | A | A | C | T | C | A | G | A | T | C | A | T | G | T |
| NC038167.1 | Plecoptera | *Isoperla eximia* | C | C | G | G | T | C | T | G | A | A | C | T | C | A | G | A | T | C | A | T | G | T |
| KT186102.1 | Plecoptera | *Kamimuria chungnanshana* | C | C | G | G | T | C | T | G | A | A | C | T | C | A | G | A | T | C | A | T | G | T |
| NC028076.1 | Plecoptera | *Kamimuria chungnanshana* | C | C | G | G | T | C | T | G | A | A | C | T | C | A | G | A | T | C | A | T | G | T |
| MN400755.1 | Plecoptera | *Kamimuria klapaleki* | C | C | G | G | T | C | T | G | A | A | C | T | C | A | G | A | T | C | A | T | G | T |
| MK111414.1 | Plecoptera | *Klapopteryx armillata* | C | C | G | G | T | C | T | G | A | A | C | T | C | A | G | A | T | C | A | T | G | T |
| MH374046.1 | Plecoptera | *Lednia tumana* | C | C | G | G | T | C | T | G | A | A | C | T | C | A | G | A | T | C | A | T | G | T |
| MK568475.1 | Plecoptera | *Leuctra fusca* | C | C | G | G | T | C | T | G | A | A | C | T | C | A | G | A | T | C | A | T | G | T |

| | | | | | | | | | | | | | | | | | | | | | | | | |
|---|---|---|---|---|---|---|---|---|---|---|---|---|---|---|---|---|---|---|---|---|---|---|---|---|
| KP642637.1 | Plecoptera | *Mesocapnia arizonensis* | C | C | G | G | T | C | T | G | A | A | C | T | C | A | G | A | T | C | A | T | G | T |
| KY568983.1 | Plecoptera | *Mesocapnia daxingana* | C | C | G | G | T | C | T | G | A | A | C | T | C | A | G | A | T | C | A | T | G | T |
| MH085450.1 | Plecoptera | *Mesonemoura metafiligera* | C | C | G | G | T | C | T | G | A | A | C | T | C | A | G | A | T | C | A | T | G | T |
| NC044719.1 | Plecoptera | *Mesonemoura metafiligera* | C | C | G | G | T | C | T | G | A | A | C | T | C | A | G | A | T | C | A | T | G | T |
| MH085451.1 | Plecoptera | *Mesonemoura tritaenia* | C | C | G | G | T | C | T | G | A | A | C | T | C | A | G | A | T | C | A | T | G | T |
| NC044720.1 | Plecoptera | *Mesonemoura tritaenia* | C | C | G | G | T | C | T | G | A | A | C | T | C | A | G | A | T | C | A | T | G | T |
| MN096323.1 | Plecoptera | *Microperla geei* | C | C | G | G | T | C | T | G | A | A | C | T | C | A | G | A | T | C | A | T | G | T |
| MT410776.1 | Plecoptera | *Nemoura avicularis* | C | C | G | G | T | C | T | G | A | A | C | T | C | A | G | A | T | C | A | T | G | T |
| MT410849.1 | Plecoptera | *Nemoura cinerea* | C | C | G | G | T | C | T | G | A | A | C | T | C | A | G | A | T | C | A | T | G | T |
| KY940360.1 | Plecoptera | *Nemoura nankinensis* | C | C | G | G | T | C | T | G | A | A | C | T | C | A | G | A | T | C | A | T | G | T |
| NC034939.1 | Plecoptera | *Nemoura nankinensis* | C | C | G | G | T | C | T | G | A | A | C | T | C | A | G | A | T | C | A | T | G | T |
| MK290826.1 | Plecoptera | *Nemoura papilla* | C | C | G | G | T | C | T | G | A | A | C | T | C | A | G | A | T | C | A | T | G | T |
| MK111418.1 | Plecoptera | *Neonemura* | C | C | G | G | T | C | T | G | A | A | C | T | C | A | G | A | T | C | A | T | G | T |

| | | | | | | | | | | | | | | | | | | | | | | | | |
|---|---|---|---|---|---|---|---|---|---|---|---|---|---|---|---|---|---|---|---|---|---|---|---|---|
| | | *barrosi* | | | | | | | | | | | | | | | | | | | | | | |
| NC042206.1 | Plecoptera | *Neonemura barrosi* | C | C | G | G | T | C | T | G | A | A | C | T | C | A | G | A | T | C | A | T | G | T |
| KX091858.1 | Plecoptera | *Neoperla ignacsiveci* | C | C | G | G | T | C | T | G | A | A | C | T | C | A | G | A | T | C | A | T | G | T |
| KX091859.1 | Plecoptera | *Neoperla* sp. | C | C | G | G | T | C | T | G | A | A | C | T | C | A | G | A | T | C | A | T | G | T |
| MN400756.1 | Plecoptera | *Neoperlops gressitti* | C | C | G | G | T | C | T | G | A | A | C | T | C | A | G | A | T | C | A | T | G | T |
| MK111415.1 | Plecoptera | *Neuroperla schedingi* | C | C | G | G | T | C | T | G | A | A | C | T | C | A | G | A | T | C | A | T | G | T |
| NC042200.1 | Plecoptera | *Neuroperla schedingi* | C | C | G | G | T | C | T | G | A | A | C | T | C | A | G | A | T | C | A | T | G | T |
| MK686067.1 | Plecoptera | *Niponiella limbatella* | C | C | G | G | T | C | T | G | A | A | C | T | C | A | G | A | T | C | A | T | G | T |
| MK387068.1 | Plecoptera | *Peltoperlopsis cebuano* | C | C | G | G | T | C | T | G | A | A | C | T | C | A | G | A | T | C | A | T | G | T |
| MF197377.1 | Plecoptera | *Perlodes* sp. | C | C | G | G | T | C | T | G | A | A | C | T | C | A | G | A | T | C | A | T | G | T |
| MH085452.1 | Plecoptera | *Protonemura kohnoae* | C | C | G | G | T | C | T | G | A | A | C | T | C | A | G | A | T | C | A | T | G | T |
| NC044752.1 | Plecoptera | *Protonemura kohnoae* | C | C | G | G | T | C | T | G | A | A | C | T | C | A | G | A | T | C | A | T | G | T |
| MT410850.1 | Plecoptera | *Protonemura* | C | C | G | G | T | C | T | G | A | A | C | T | C | A | G | A | T | C | A | T | G | T |

| | | | | | | | | | | | | | | | | | | | | | | | | |
|---|---|---|---|---|---|---|---|---|---|---|---|---|---|---|---|---|---|---|---|---|---|---|---|---|
| | | *meyeri* | | | | | | | | | | | | | | | | | | | | | | |
| NC050322.1 | Plecoptera | *Protonemura meyeri* | C | C | G | G | T | C | T | G | A | A | C | T | C | A | G | A | T | C | A | T | G | T |
| MH085453.1 | Plecoptera | *Protonemura orbiculata* | C | C | G | G | T | C | T | G | A | A | C | T | C | A | G | A | T | C | A | T | G | T |
| NC044753.1 | Plecoptera | *Protonemura orbiculata* | C | C | G | G | T | C | T | G | A | A | C | T | C | A | G | A | T | C | A | T | G | T |
| MG910458.1 | Plecoptera | *Pseudomegarcys japonica* | C | C | G | G | T | C | T | G | A | A | C | T | C | A | G | A | T | C | A | T | G | T |
| NC038168.1 | Plecoptera | *Pseudomegarcys japonica* | C | C | G | G | T | C | T | G | A | A | C | T | C | A | G | A | T | C | A | T | G | T |
| KU182360.1 | Plecoptera | *Pteronarcella badia* | C | C | G | G | T | C | T | G | A | A | C | T | C | A | G | A | T | C | A | T | G | T |
| NC029248.1 | Plecoptera | *Pteronarcella badia* | C | C | G | G | T | C | T | G | A | A | C | T | C | A | G | A | T | C | A | T | G | T |
| AY687866.1 | Plecoptera | *Pteronarcys princeps* | C | C | G | G | T | C | T | G | A | A | C | T | C | A | G | A | T | C | A | T | G | T |
| NC006133.1 | Plecoptera | *Pteronarcys princeps* | C | C | G | G | T | C | T | G | A | A | C | T | C | A | G | A | T | C | A | T | G | T |
| MK111419.1 | Plecoptera | *Rhopalopsole bulbifera* | C | C | G | G | T | C | T | G | A | A | C | T | C | A | G | A | T | C | A | T | G | T |
| NC042207.1 | Plecoptera | *Rhopalopsole* | C | C | G | G | T | C | T | G | A | A | C | T | C | A | G | A | T | C | A | T | G | T |

| | | | | | | | | | | | | | | | | | | | | | | | | |
|---|---|---|---|---|---|---|---|---|---|---|---|---|---|---|---|---|---|---|---|---|---|---|---|---|
| | | *bulbifera* | | | | | | | | | | | | | | | | | | | | | | |
| MH510071.1 | Plecoptera | *Scopura longa* | C | C | G | G | T | C | T | G | A | A | C | T | C | A | G | A | T | C | A | T | G | T |
| NC041105.1 | Plecoptera | *Scopura longa* | C | C | G | G | T | C | T | G | A | A | C | T | C | A | G | A | T | C | A | T | G | T |
| MK492253.1 | Plecoptera | *Sinacroneuria dabieshana* | C | C | G | G | T | C | T | G | A | A | C | T | C | A | G | A | T | C | A | T | G | T |
| MF716958.1 | Plecoptera | *Soliperla* sp. | C | C | G | G | T | C | T | G | A | A | C | T | C | A | G | A | T | C | A | T | G | T |
| NC038189.1 | Plecoptera | *Soliperla* sp. | C | C | G | G | T | C | T | G | A | A | C | T | C | A | G | A | T | C | A | T | G | T |
| MH085455.1 | Plecoptera | *Sphaeronemoura acutispina* | C | C | G | G | T | C | T | G | A | A | C | T | C | A | G | A | T | C | A | T | G | T |
| MH085454.1 | Plecoptera | *Sphaeronemoura grandicauda* | C | C | G | G | T | C | T | G | A | A | C | T | C | A | G | A | T | C | A | T | G | T |
| NC044754.1 | Plecoptera | *Sphaeronemoura grandicauda* | C | C | G | G | T | C | T | G | A | A | C | T | C | A | G | A | T | C | A | T | G | T |
| MK111420.1 | Plecoptera | *Sphaeronemoura hainana* | C | C | G | G | T | C | T | G | A | A | C | T | C | A | G | A | T | C | A | T | G | T |
| NC044755.1 | Plecoptera | *Sphaeronemoura hamistyla* | C | C | G | G | T | C | T | G | A | A | C | T | C | A | G | A | T | C | A | T | G | T |
| KR088971.1 | Plecoptera | *Styloperla* sp. | C | C | G | G | T | C | T | G | A | A | C | T | C | A | G | A | T | C | A | T | G | T |
| KX845569.1 | Plecoptera | *Styloperla spinicercia* | C | C | G | G | T | C | T | G | A | A | C | T | C | A | G | A | T | C | A | T | G | T |
| NC034809.1 | Plecoptera | *Styloperla spinicercia* | C | C | G | G | T | C | T | G | A | A | C | T | C | A | G | A | T | C | A | T | G | T |

| | | | | | | | | | | | | | | | | | | | | | | |
|---|---|---|---|---|---|---|---|---|---|---|---|---|---|---|---|---|---|---|---|---|---|---|
| KX091865.1 | Plecoptera | *Suwallia bimaculata* | C | C | G | G | T | C | T | G | A | A | C | T | C | A | G | A | T | C | A | T | G | T |
| MN121757.1 | Plecoptera | *Suwallia bimaculata* | C | C | G | G | T | C | T | G | A | A | C | T | C | A | G | A | T | C | A | T | G | T |
| MF198253.1 | Plecoptera | *Suwallia teleckojensis* | C | C | G | G | T | C | T | G | A | A | C | T | C | A | G | A | T | C | A | T | G | T |
| NC037754.1 | Plecoptera | *Suwallia teleckojensis* | C | C | G | G | T | C | T | G | A | A | C | T | C | A | G | A | T | C | A | T | G | T |
| MG589786.1 | Plecoptera | *Taeniopteryx ugola* | C | C | G | G | T | C | T | G | A | A | C | T | C | A | G | A | T | C | A | T | G | T |
| NC037897.1 | Plecoptera | *Taeniopteryx ugola* | C | C | G | G | T | C | T | G | A | A | C | T | C | A | G | A | T | C | A | T | G | T |
| KM409708.1 | Plecoptera | *Togoperla* sp. | C | C | G | G | T | C | T | G | A | A | C | T | C | A | G | A | T | C | A | T | G | T |
| KY522907.1 | Plecoptera | *Zelandoperla fenestrata* | C | C | G | G | T | C | T | G | A | A | C | T | C | A | G | A | T | C | A | T | G | T |
| NC034997.1 | Plecoptera | *Zelandoperla fenestrata* | C | C | G | G | T | C | T | G | A | A | C | T | C | A | G | A | T | C | A | T | G | T |
| FJ456939.1 | Heteroptera | *Aphelocheirus ellipsoideus* | C | C | G | G | T | C | T | G | A | A | C | T | C | A | A | A | T | C | A | T | G | T |
| KX505849.1 | Heteroptera | *Aphelocheirus jendeki* | C | C | G | G | T | C | T | G | A | A | C | T | C | A | A | A | T | C | A | T | G | T |
| NC037370.1 | Heteroptera | *Aphelocheirus* | C | C | G | G | T | C | T | G | A | A | C | T | C | A | A | A | T | C | A | T | G | T |

| | | | | | | | | | | | | | | | | | | | | | | | | |
|---|---|---|---|---|---|---|---|---|---|---|---|---|---|---|---|---|---|---|---|---|---|---|---|---|
| | | *jendeki* | | | | | | | | | | | | | | | | | | | | | | |
| FJ456944.1 | Heteroptera | *Aquarius paludum* | C | C | G | G | T | C | T | G | A | A | C | T | C | A | A | A | T | C | A | T | G | T |
| NC012841.1 | Heteroptera | *Aquarius paludum* | C | C | G | G | T | C | T | G | A | A | C | T | C | A | A | A | T | C | A | T | G | T |
| MK251109.1 | Heteroptera | *Diplonychus esakii* | C | C | G | G | T | C | T | G | A | A | C | T | C | A | A | A | T | C | A | T | G | T |
| FJ456940.1 | Heteroptera | *Diplonychus rusticus* | C | C | G | G | T | C | T | G | A | A | C | T | C | A | A | A | T | C | A | T | G | T |
| FJ456949.1 | Heteroptera | *Enithares tibialis* | C | C | G | G | T | C | T | T | A | A | C | T | C | A | A | A | T | C | A | T | G | T |
| NC012819.1 | Heteroptera | *Enithares tibialis* | C | C | G | G | T | C | T | T | A | A | C | T | C | A | A | A | T | C | A | T | G | T |
| KP400582.1 | Heteroptera | *Entomovelia* sp. | C | C | G | G | T | C | T | G | A | A | C | T | C | A | A | A | T | C | A | T | G | T |
| MF177288.1 | Heteroptera | *Gigantometra gigas* | C | C | G | G | T | C | T | G | A | A | C | T | C | A | A | A | T | C | A | T | G | T |
| MF177289.1 | Heteroptera | *Gigantometra gigas* | C | C | G | G | T | C | T | G | A | A | C | T | C | A | A | A | T | C | A | T | G | T |
| MF177290.1 | Heteroptera | *Gigantometra gigas* | C | C | G | G | T | C | T | G | A | A | C | T | C | A | A | A | T | C | A | T | G | T |
| MF177291.1 | Heteroptera | *Gigantometra gigas* | C | C | G | G | T | C | T | G | A | A | C | T | C | A | A | A | T | C | A | T | G | T |
| NC041084.1 | Heteroptera | *Gigantometra gigas* | C | C | G | G | T | C | T | G | A | A | C | T | C | A | A | A | T | C | A | T | G | T |
| KJ027513.1 | Heteroptera | *Helotrephes semiglobosus semiglobosus* | C | C | G | G | T | C | T | G | A | A | C | T | C | A | A | A | T | C | A | T | G | T |

| | | | | | | | | | | | | | | | | | | | | | | | |
|---|---|---|---|---|---|---|---|---|---|---|---|---|---|---|---|---|---|---|---|---|---|---|---|
| FJ456951.1 | Heteroptera | *Helotrephes* sp. | C | C | G | G | T | C | T | G | A | A | C | T | C | A | A | A | T | C | A | T | G | T |
| NC012822.1 | Heteroptera | *Helotrephes* sp. | C | C | G | G | T | C | T | G | A | A | C | T | C | A | A | A | T | C | A | T | G | T |
| FJ456945.1 | Heteroptera | *Hydrometra greeni* | C | C | G | G | T | T | T | G | A | A | C | T | C | A | A | A | T | C | A | T | G | T |
| NC012842.1 | Heteroptera | *Hydrometra greeni* | C | C | G | G | T | T | T | G | A | A | C | T | C | A | A | A | T | C | A | T | G | T |
| FJ456947.1 | Heteroptera | *Ilyocoris cimicoides* | C | C | G | G | T | T | T | G | A | A | C | T | C | A | A | A | T | C | A | T | G | T |
| NC012845.1 | Heteroptera | *Ilyocoris cimicoides* | C | C | G | G | T | T | T | G | A | A | C | T | C | A | A | A | T | C | A | T | G | T |
| FJ456948.1 | Heteroptera | *Laccotrephes robustus* | C | C | G | G | T | C | T | G | A | A | C | T | C | A | A | A | T | C | A | T | G | T |
| NC012817.1 | Heteroptera | *Laccotrephes robustus* | C | C | G | G | T | C | T | G | A | A | C | T | C | A | A | A | T | C | A | T | G | T |
| FJ456946.1 | Heteroptera | *Leptopus* sp. | C | C | G | G | T | C | T | G | A | A | C | T | C | A | G | A | T | C | A | T | G | T |
| JQ910985.1 | Heteroptera | *Lethocerus deyrollei* | C | C | G | G | T | C | T | G | A | A | C | T | C | A | A | A | T | C | A | T | G | T |
| KU237288.1 | Heteroptera | *Lethocerus deyrollei* | C | C | G | G | T | C | T | G | A | A | C | T | C | A | A | A | T | C | A | T | G | T |
| KM588201.1 | Heteroptera | *Lethocerus indicus* | C | C | G | G | T | C | T | G | A | A | C | T | C | A | A | A | T | C | A | T | G | T |
| KR072671.1 | Heteroptera | *Lethocerus indicus* | C | C | G | G | T | C | T | G | A | A | C | T | C | A | A | A | T | C | A | T | G | T |
| NC027194.1 | Heteroptera | *Lethocerus indicus* | C | C | G | G | T | C | T | G | A | A | C | T | C | A | A | A | T | C | A | T | G | T |
| FJ456942.1 | Heteroptera | *Lycorma delicatula* | C | C | G | G | C | T | T | G | A | A | C | T | C | A | G | A | T | C | A | T | G | T |
| NC012835.1 | Heteroptera | *Lycorma delicatula* | C | C | G | G | C | T | T | G | A | A | C | T | C | A | G | A | T | C | A | T | G | T |

| | | | | | | | | | | | | | | | | | | | | | | | | |
|---|---|---|---|---|---|---|---|---|---|---|---|---|---|---|---|---|---|---|---|---|---|---|---|---|
| MN027278.1 | Heteroptera | *Metrocoris* sp. | C | C | G | G | T | C | T | G | A | A | C | T | C | A | A | A | T | C | A | T | G | T |
| MH382835.1 | Heteroptera | *Micronecta sahlbergii* | C | C | G | G | T | T | T | G | A | A | C | T | C | A | A | A | T | C | A | T | G | T |
| NC039588.1 | Heteroptera | *Micronecta sahlbergii* | C | C | G | G | T | T | T | G | A | A | C | T | C | A | A | A | T | C | A | T | G | T |
| KT031802.1 | Heteroptera | *Nepa hoffmanni* | C | C | G | G | T | T | T | G | A | A | C | T | C | A | A | A | T | C | A | T | G | T |
| NC028084.1 | Heteroptera | *Nepa hoffmanni* | C | C | G | G | T | T | T | G | A | A | C | T | C | A | A | A | T | C | A | T | G | T |
| FJ456943.1 | Heteroptera | *Nerthra indica* | C | C | G | G | T | T | T | G | A | A | C | T | C | A | A | A | T | C | A | T | G | T |
| NC012838.1 | Heteroptera | *Nerthra indica* | C | C | G | G | T | T | T | G | A | A | C | T | C | A | A | A | T | C | A | T | G | T |
| KX034086.1 | Heteroptera | *Notonecta chinensis* | C | C | G | G | T | T | T | G | A | A | C | T | C | A | A | A | T | C | A | T | G | T |
| NC036671.1 | Heteroptera | *Notonecta chinensis* | C | C | G | G | T | T | T | G | A | A | C | T | C | A | A | A | T | C | A | T | G | T |
| MH780861.1 | Heteroptera | *Notonecta montandoni* | C | C | G | G | T | T | T | G | A | A | C | T | C | A | A | A | T | C | A | T | G | T |
| NC045886.1 | Heteroptera | *Notonecta montandoni* | C | C | G | G | T | T | T | G | A | A | C | T | C | A | A | A | T | C | A | T | G | T |
| KM244707.1 | Heteroptera | *Notonectidae* sp. | C | C | G | G | T | C | T | T | A | A | C | T | C | A | A | A | T | C | A | T | G | T |
| FJ456950.1 | Heteroptera | *Ochterus marginatus* | C | C | G | G | T | T | T | G | A | A | C | T | C | A | A | A | T | C | A | T | G | T |
| KJ027516.1 | Heteroptera | *Paraplea frontalis* | C | C | G | G | T | T | T | G | A | A | C | T | C | A | A | A | T | C | A | T | G | T |
| NC028629.1 | Heteroptera | *Paraplea frontalis* | C | C | G | G | T | T | T | G | A | A | C | T | C | A | A | A | T | C | A | T | G | T |

| | | | | | | | | | | | | | | | | | | | | | | | | |
|---|---|---|---|---|---|---|---|---|---|---|---|---|---|---|---|---|---|---|---|---|---|---|---|---|
| JQ910988.1 | Heteroptera | *Perittopus* sp. | C | C | G | G | T | C | T | G | A | A | C | T | C | A | A | A | T | C | A | T | G | T |
| KP400583.1 | Heteroptera | *Ptilomera tigrina* | C | C | G | G | T | C | T | G | A | A | C | T | C | A | A | A | T | C | A | T | G | T |
| MN027271.1 | Heteroptera | *Rhagovelia reitteri* | C | C | G | G | T | C | T | G | A | A | C | T | C | A | A | A | T | C | A | T | G | T |
| NC044721.1 | Heteroptera | *Rhagovelia reitteri* | C | C | G | G | T | C | T | G | A | A | C | T | C | A | A | A | T | C | A | T | G | T |
| MN027279.1 | Heteroptera | *Rhagovelia* sp. | C | C | G | G | T | C | T | G | A | A | C | T | C | A | A | A | T | C | A | T | G | T |
| FJ456941.1 | Heteroptera | *Sigara septemlineata* | C | C | G | G | T | C | T | G | A | A | C | T | C | A | A | A | T | C | A | T | G | T |
| FJ456952.1 | Heteroptera | *Valentia hoffmanni* | C | C | G | G | T | C | T | G | A | A | C | T | C | A | G | A | T | C | A | T | G | T |
| NC012823.1 | Heteroptera | *Valentia hoffmanni* | C | C | G | G | T | C | T | G | A | A | C | T | C | A | G | A | T | C | A | T | G | T |
| KF840564.1 | Megaloptera | *Acanthacorydalis orientalis* | C | C | G | G | T | T | T | G | A | A | C | T | C | A | A | A | T | C | A | T | G | T |
| NC023462.1 | Megaloptera | *Acanthacorydalis orientalis* | C | C | G | G | T | T | T | G | A | A | C | T | C | A | A | A | T | C | A | T | G | T |
| KU925864.1 | Megaloptera | *Archichauliodes deceptor* | C | C | G | G | T | T | T | G | A | A | C | T | C | A | A | A | T | C | A | T | G | T |
| NC035504.1 | Megaloptera | *Archichauliodes deceptor* | C | C | G | G | T | T | T | G | A | A | C | T | C | A | A | A | T | C | A | T | G | T |
| KU302713.1 | Megaloptera | *Chloronia mirifica* | C | C | G | G | T | T | T | G | A | A | C | T | C | A | A | A | T | C | A | T | G | T |
| NC033349.1 | Megaloptera | *Chloronia mirifica* | C | C | G | G | T | T | T | G | A | A | C | T | C | A | A | A | T | C | A | T | G | T |
| FJ171323.1 | Megaloptera | *Corydalus cornutus* | C | C | G | G | T | T | T | G | A | A | C | T | C | A | A | A | T | C | A | T | G | T |
| NC011276.1 | Megaloptera | *Corydalus* | C | C | G | G | T | T | T | G | A | A | C | T | C | A | A | A | T | C | A | T | G | T |

| | | | | | | | | | | | | | | | | | | | | | | | |
|---|---|---|---|---|---|---|---|---|---|---|---|---|---|---|---|---|---|---|---|---|---|---|---|
| | | *cornutus* | | | | | | | | | | | | | | | | | | | | | |
| KJ806318.1 | Megaloptera | *Dysmicohermes ingens* | C | C | G | G | T | T | T | G | A | A | C | T | C | A | A | A | T | C | A | T | G | T |
| NC024657.1 | Megaloptera | *Dysmicohermes ingens* | C | C | G | G | T | T | T | G | A | A | C | T | C | A | A | A | T | C | A | T | G | T |
| JQ351950.1 | Megaloptera | *Neochauliodes bowringi* | C | C | G | G | T | T | T | G | A | A | C | T | C | A | A | A | T | C | A | T | G | T |
| NC023444.1 | Megaloptera | *Neochauliodes bowringi* | C | C | G | G | T | T | T | G | A | A | C | T | C | A | A | A | T | C | A | T | G | T |
| KF785799.1 | Megaloptera | *Neochauliodes fraternus* | C | C | G | G | T | T | T | G | A | A | C | T | C | A | A | A | T | C | A | T | G | T |
| NC025282.1 | Megaloptera | *Neochauliodes fraternus* | C | C | G | G | T | T | T | G | A | A | C | T | C | A | A | A | T | C | A | T | G | T |
| KX821680.1 | Megaloptera | *Neochauliodes parasparsus* | C | C | G | G | T | T | T | G | A | A | C | T | C | A | A | A | T | C | A | T | G | T |
| JX110703.1 | Megaloptera | *Neochauliodes punctatolosus* | C | C | G | G | T | T | T | G | A | A | C | T | C | A | A | A | T | C | A | T | G | T |
| MK301249.1 | Megaloptera | *Neochauliodes punctatolosus* | C | C | G | G | T | T | T | G | A | A | C | T | C | A | A | A | T | C | A | T | G | T |
| NC018772.1 | Megaloptera | *Neochauliodes punctatolosus* | C | C | G | G | T | T | T | G | A | A | C | T | C | A | A | A | T | C | A | T | G | T |
| KF771894.1 | Megaloptera | *Neochauliodes* | C | C | G | G | T | T | T | G | A | A | C | T | C | A | A | A | T | C | A | T | G | T |

| | | | | | | | | | | | | | | | | | | | | | | | | |
|---|---|---|---|---|---|---|---|---|---|---|---|---|---|---|---|---|---|---|---|---|---|---|---|---|
| | | *rotundatus* | | | | | | | | | | | | | | | | | | | | | | |
| NC025281.1 | Megaloptera | *Neochauliodes rotundatus* | C | C | G | G | T | T | T | G | A | A | C | T | C | A | A | A | T | C | A | T | G | T |
| KP126231.1 | Megaloptera | *Neoneuromus tonkinensis* | C | C | G | G | T | T | T | G | A | A | C | T | C | A | A | A | T | C | A | T | G | T |
| NC027852.1 | Megaloptera | *Neoneuromus tonkinensis* | C | C | G | G | T | T | T | G | A | A | C | T | C | A | A | A | T | C | A | T | G | T |
| KP126232.1 | Megaloptera | *Nevromus exterior* | C | C | G | G | T | T | T | G | A | A | C | T | C | A | A | A | T | C | A | T | G | T |
| NC027851.1 | Megaloptera | *Nevromus exterior* | C | C | G | G | T | T | T | G | A | A | C | T | C | A | A | A | T | C | A | T | G | T |
| KY230493.1 | Megaloptera | *Protochauliodes biconicus* | C | C | G | G | T | T | T | G | A | A | C | T | C | A | A | A | T | C | A | T | G | T |
| EU526394.1 | Megaloptera | *Protohermes concolorus* | C | C | G | G | T | T | T | G | A | A | C | T | C | A | A | A | T | C | A | T | G | T |
| NC011524.1 | Megaloptera | *Protohermes concolorus* | C | C | G | G | T | T | T | G | A | A | C | T | C | A | A | A | T | C | A | T | G | T |
| FJ859905.1 | Megaloptera | *Sialis hamata* | C | C | G | A | T | T | T | G | A | A | C | T | C | A | A | A | T | C | A | T | G | T |
| NC013256.1 | Megaloptera | *Sialis hamata* | C | C | G | A | T | T | T | G | A | A | C | T | C | A | A | A | T | C | A | T | G | T |
| KT876911.1 | Megaloptera | *Sialis* sp. | C | C | G | A | T | T | T | G | A | A | C | T | C | A | A | A | T | C | A | T | G | T |
| KT425084.1 | Megaloptera | *Stenosialis australiensis* | C | C | G | A | T | T | T | G | A | A | C | T | C | A | A | A | T | C | A | T | G | T |
| KY587199.1 | Neuroptera | *Abachrysa eureka* | C | C | G | G | T | C | T | G | A | A | C | T | C | A | A | A | T | C | A | T | G | T |
| NC015095.1 | Neuroptera | *Apochrysa* | C | C | G | G | T | C | T | G | A | A | C | T | C | A | A | A | T | C | A | T | G | T |

| | | | | | | | | | | | | | | | | | | | | | | | | |
|---|---|---|---|---|---|---|---|---|---|---|---|---|---|---|---|---|---|---|---|---|---|---|---|---|
| | | *matsumurae* | | | | | | | | | | | | | | | | | | | | | | |
| KC758703.1 | Neuroptera | *Ascalohybris subjacens* | C | C | G | G | T | C | T | G | A | A | C | T | C | A | A | A | T | C | A | T | G | T |
| NC021428.1 | Neuroptera | *Ascalohybris subjacens* | C | C | G | G | T | C | T | G | A | A | C | T | C | A | A | A | T | C | A | T | G | T |
| FJ171324.1 | Neuroptera | *Ascaloptynx appendiculatus* | C | C | G | G | T | C | T | G | A | A | C | T | C | A | A | A | T | C | A | T | G | T |
| NC011277.1 | Neuroptera | *Ascaloptynx appendiculatus* | C | C | G | G | T | C | T | G | A | A | C | T | C | A | A | A | T | C | A | T | G | T |
| KT425083.1 | Neuroptera | *Balmes birmanus* | C | C | G | G | T | C | T | G | A | A | C | T | C | A | A | A | T | C | A | T | G | T |
| KX369241.1 | Neuroptera | *Bullanga florida* | C | C | G | G | T | C | T | G | A | A | C | T | C | A | A | A | T | C | A | T | G | T |
| NC032298.1 | Neuroptera | *Bullanga florida* | C | C | G | G | T | C | T | G | A | A | C | T | C | A | A | A | T | C | A | T | G | T |
| KT425069.1 | Neuroptera | *Chasmoptera huttii* | C | C | G | G | T | C | T | G | A | A | C | T | C | A | A | A | T | C | A | T | G | T |
| JX033119.1 | Neuroptera | *Chrysopa pallens* | C | C | G | G | T | C | T | G | A | A | C | T | C | A | A | A | T | C | A | T | G | T |
| NC019618.1 | Neuroptera | *Chrysopa pallens* | C | C | G | G | T | C | T | G | A | A | C | T | C | A | A | A | T | C | A | T | G | T |
| KJ778879.1 | Neuroptera | *Chrysoperla carnea* | C | C | G | G | T | C | T | G | A | A | C | T | C | A | A | A | T | C | A | T | G | T |
| KU877169.1 | Neuroptera | *Chrysoperla externa* | C | C | G | G | T | C | T | G | A | A | C | T | C | A | A | A | T | C | A | T | G | T |
| NC030341.1 | Neuroptera | *Chrysoperla externa* | C | C | G | G | T | C | T | G | A | A | C | T | C | A | A | A | T | C | A | T | G | T |
| KP027407.1 | Neuroptera | *Chrysoperla* | C | C | G | G | T | C | T | G | A | A | C | T | C | A | A | A | T | C | A | T | G | T |

| | | | | | | | | | | | | | | | | | | | | | | | |
|---|---|---|---|---|---|---|---|---|---|---|---|---|---|---|---|---|---|---|---|---|---|---|---|
| | | *nipponensis* | | | | | | | | | | | | | | | | | | | | | |
| NC015093.1 | Neuroptera | *Chrysoperla nipponensis* | C | C | G | G | T | C | T | G | A | A | C | T | C | A | A | A | T | C | A | T | G | T |
| KT425088.1 | Neuroptera | *Climacia areolaris* | C | C | G | G | T | C | T | G | A | A | C | T | C | A | A | A | T | C | A | T | G | T |
| KT425078.1 | Neuroptera | *Coniopteryx* sp. | C | C | G | G | T | C | T | G | A | A | C | T | C | A | A | A | T | C | A | T | G | T |
| KT425068.1 | Neuroptera | *Dendroleon pantherinus* | C | C | G | G | T | C | T | G | A | A | C | T | C | A | A | A | T | C | A | T | G | T |
| MK301246.1 | Neuroptera | *Dendroleon pantherinus* | C | C | G | G | T | C | T | G | A | A | C | T | C | A | A | A | T | C | A | T | G | T |
| KT425073.1 | Neuroptera | *Dilar* sp. | C | C | G | G | T | C | T | G | A | A | C | T | C | A | A | A | T | C | A | T | G | T |
| FJ859906.1 | Neuroptera | *Ditaxis biseriata* | C | C | G | G | T | T | T | G | A | A | C | T | C | A | A | A | T | C | A | T | G | T |
| NC013257.1 | Neuroptera | *Ditaxis biseriata* | C | C | G | G | T | T | T | G | A | A | C | T | C | A | A | A | T | C | A | T | G | T |
| KT425087.1 | Neuroptera | *Drepanepteryx phalaenoides* | C | C | G | G | T | C | T | G | A | A | C | T | C | A | A | A | T | C | A | T | G | T |
| KF701327.1 | Neuroptera | *Epacanthaclisis banksi* | C | C | G | G | T | C | T | G | A | A | C | T | C | A | A | A | T | C | A | T | G | T |
| NC025905.1 | Neuroptera | *Epacanthaclisis banksi* | C | C | G | G | T | C | T | G | A | A | C | T | C | A | A | A | T | C | A | T | G | T |
| MF972496.1 | Neuroptera | *Euclimacia badia* | C | C | G | G | T | T | T | G | A | A | C | T | C | A | A | A | T | C | A | T | G | T |
| NC039773.1 | Neuroptera | *Euclimacia badia* | C | C | G | G | T | T | T | G | A | A | C | T | C | A | A | A | T | C | A | T | G | T |
| KT425080.1 | Neuroptera | *Eumantispa harmandi* | C | C | G | G | T | T | T | G | A | A | C | T | C | A | A | A | T | C | A | T | G | T |

| | | | | | | | | | | | | | | | | | | | | | | | | |
|---|---|---|---|---|---|---|---|---|---|---|---|---|---|---|---|---|---|---|---|---|---|---|---|---|
| MF972495.1 | Neuroptera | *Eumantispa harmandi* | C | C | G | G | T | T | T | G | A | A | C | T | C | A | A | A | T | C | A | T | G | T |
| NC039772.1 | Neuroptera | *Eumantispa harmandi* | C | C | G | G | T | T | T | G | A | A | C | T | C | A | A | A | T | C | A | T | G | T |
| KT425072.1 | Neuroptera | *Fontecilla graphicus* | C | C | G | G | T | C | T | G | A | A | C | T | C | A | A | A | T | C | A | T | G | T |
| KY364372.1 | Neuroptera | *Gatzara jezoensis* | C | C | G | G | T | C | T | G | A | A | C | T | C | A | A | A | T | C | A | T | G | T |
| MK341582.1 | Neuroptera | *Gryposmylus pennyi* | C | C | G | G | T | C | T | G | A | A | C | T | C | A | A | A | T | C | A | T | G | T |
| MN852445.1 | Neuroptera | *Hemerobius japonicus* | C | C | G | G | T | C | T | G | A | A | C | T | C | A | A | A | T | C | A | T | G | T |
| NC048517.1 | Neuroptera | *Hemerobius japonicus* | C | C | G | G | T | C | T | G | A | A | C | T | C | A | A | A | T | C | A | T | G | T |
| KT425077.1 | Neuroptera | *Heterosmylus* sp. | C | C | G | G | T | C | T | G | A | A | C | T | C | A | A | A | T | C | A | T | G | T |
| KT425085.1 | Neuroptera | *Inocellia fujiana* | C | C | G | G | T | T | T | G | A | A | C | T | C | A | A | A | T | C | A | T | G | T |
| KY587200.1 | Neuroptera | *Italochrysa insignis* | C | C | G | G | T | C | T | G | A | A | C | T | C | A | A | A | T | C | A | T | G | T |
| KY587201.1 | Neuroptera | *Leucochrysa pretiosa* | C | C | G | G | T | C | T | G | A | A | C | T | C | A | A | A | T | C | A | T | G | T |
| NC015609.1 | Neuroptera | *Libelloides macaronius* | C | C | G | G | T | C | T | G | A | A | C | T | C | A | A | A | T | C | A | T | G | T |
| MF972494.1 | Neuroptera | *Mantispa japonica* | C | C | G | G | T | T | T | G | A | A | C | T | C | A | A | A | T | C | A | T | G | T |

| | | | | | | | | | | | | | | | | | | | | | | | | |
|---|---|---|---|---|---|---|---|---|---|---|---|---|---|---|---|---|---|---|---|---|---|---|---|---|
| NC039771.1 | Neuroptera | *Mantispa japonica* | C | C | G | G | T | T | T | G | A | A | C | T | C | A | A | A | T | C | A | T | G | T |
| MK472073.1 | Neuroptera | *Micromus angulatus* | C | C | G | G | T | C | T | G | A | A | C | T | C | A | A | A | T | C | A | T | G | T |
| NC042677.1 | Neuroptera | *Micromus angulatus* | C | C | G | G | T | C | T | G | A | A | C | T | C | A | A | A | T | C | A | T | G | T |
| KT425075.1 | Neuroptera | *Micromus* sp. | C | C | G | G | T | C | T | G | A | A | C | T | C | A | A | A | T | C | A | T | G | T |
| KT425092.1 | Neuroptera | *Mucroberotha vesicaria* | C | C | G | G | T | C | T | G | A | A | C | T | C | A | A | A | T | C | A | T | G | T |
| KT425089.1 | Neuroptera | *Myiodactylus osmyloides* | C | C | G | G | T | C | T | G | A | A | C | T | C | A | A | A | T | C | A | T | G | T |
| MT118664.1 | Neuroptera | *Myrmeleon formicarius* | C | C | G | G | T | C | T | G | A | A | C | T | C | A | A | A | T | C | A | T | G | T |
| NC047285.1 | Neuroptera | *Myrmeleon formicarius* | C | C | G | G | T | C | T | G | A | A | C | T | C | A | A | A | T | C | A | T | G | T |
| KJ461323.1 | Neuroptera | *Myrmeleon immanis* | C | C | G | G | T | C | T | G | A | A | C | T | C | A | A | A | T | C | A | T | G | T |
| NC024826.1 | Neuroptera | *Myrmeleon immanis* | C | C | G | G | T | C | T | G | A | A | C | T | C | A | A | A | T | C | A | T | G | T |
| KT425071.1 | Neuroptera | *Nallachius americanus* | C | C | G | G | T | C | T | G | A | A | C | T | C | A | A | A | T | C | A | T | G | T |
| KT425086.1 | Neuroptera | *Negha inflata* | C | C | G | G | T | T | T | G | A | A | C | T | C | A | A | A | T | C | A | T | G | T |
| KT425079.1 | Neuroptera | *Nemoptera coa* | C | C | G | G | T | C | T | G | A | A | C | T | C | A | A | A | T | C | A | T | G | T |

| | | | | | | | | | | | | | | | | | | | | | | |
|---|---|---|---|---|---|---|---|---|---|---|---|---|---|---|---|---|---|---|---|---|---|---|
| KR078257.1 | Neuroptera | *Neuronema laminatum* | C | C | G | G | T | C | T | G | A | A | C | T | C | A | A | A | T | C | A | T | G | T |
| NC028153.1 | Neuroptera | *Neuronema laminatum* | C | C | G | G | T | C | T | G | A | A | C | T | C | A | A | A | T | C | A | T | G | T |
| KT425074.1 | Neuroptera | *Nevrorthus apatelios* | C | C | G | G | T | C | T | G | A | A | C | T | C | A | A | A | T | C | A | T | G | T |
| KT425076.1 | Neuroptera | *Nipponeurorthus fuscinervis* | C | C | G | G | T | C | T | G | A | A | C | T | C | A | A | A | T | C | A | T | G | T |
| KP264629.1 | Neuroptera | *Nothancyla verreauxi* | C | C | G | G | T | C | T | G | A | A | C | T | C | A | A | A | T | C | A | T | G | T |
| KP264630.1 | Neuroptera | *Nothochrysa* sp. | C | C | G | G | T | C | T | G | A | A | C | T | C | A | A | A | T | C | A | T | G | T |
| KJ461322.1 | Neuroptera | *Nymphes myrmeleonoides* | C | C | G | G | T | C | T | G | A | A | C | T | C | A | A | A | T | C | A | T | G | T |
| NC024825.1 | Neuroptera | *Nymphes myrmeleonoides* | C | C | G | G | T | C | T | G | A | A | C | T | C | A | A | A | T | C | A | T | G | T |
| KT425090.1 | Neuroptera | *Oliarces clara* | C | C | G | G | T | C | T | G | A | A | C | T | C | A | A | A | T | C | A | T | G | T |
| MN818868.1 | Neuroptera | *Osmylus atomatus* | C | C | G | G | T | C | T | G | A | A | C | T | C | A | A | A | T | C | A | T | G | T |
| MN818867.1 | Neuroptera | *Osmylus fulvicephalus* | C | C | G | G | T | C | T | G | A | A | C | T | C | A | A | A | T | C | A | T | G | T |
| MN617021.1 | Neuroptera | *Osmylus lucalatus* | C | C | G | G | T | C | T | G | A | A | C | T | C | A | A | A | T | C | A | T | G | T |
| MN818870.1 | Neuroptera | *Osmylus* sp. | C | C | G | G | T | C | T | G | A | A | C | T | C | A | A | A | T | C | A | T | G | T |
| MN818869.1 | Neuroptera | *Osmylus zheanus* | C | C | G | G | T | C | T | G | A | A | C | T | C | A | A | A | T | C | A | T | G | T |

| | | | | | | | | | | | | | | | | | | | | | | | | |
|---|---|---|---|---|---|---|---|---|---|---|---|---|---|---|---|---|---|---|---|---|---|---|---|---|
| KY587202.1 | Neuroptera | *Parankylopteryx* sp. | C | C | G | G | T | C | T | G | A | A | C | T | C | A | A | A | T | C | A | T | G | T |
| KT425091.1 | Neuroptera | *Podallea* sp. | C | C | G | G | T | T | T | G | A | A | C | T | C | A | A | A | T | C | A | T | G | T |
| FJ171325.1 | Neuroptera | *Polystoechotes punctatus* | C | C | G | G | T | C | T | G | A | A | C | T | C | A | A | A | T | C | A | T | G | T |
| NC011278.1 | Neuroptera | *Polystoechotes punctatus* | C | C | G | G | T | C | T | G | A | A | C | T | C | A | A | A | T | C | A | T | G | T |
| KT425082.1 | Neuroptera | *Psychopsis coelivaga* | C | C | G | G | T | C | T | G | A | A | C | T | C | A | A | A | T | C | A | T | G | T |
| KF626446.1 | Neuroptera | *Rapisma xizangense* | C | C | G | G | T | T | T | G | A | A | C | T | C | A | A | A | T | C | A | T | G | T |
| NC023362.1 | Neuroptera | *Rapisma xizangense* | C | C | G | G | T | T | T | G | A | A | C | T | C | A | A | A | T | C | A | T | G | T |
| KF626447.1 | Neuroptera | *Rapisma zayuanum* | C | C | G | G | T | T | T | G | A | A | C | T | C | A | A | A | T | C | A | T | G | T |
| NC023363.1 | Neuroptera | *Rapisma zayuanum* | C | C | G | G | T | T | T | G | A | A | C | T | C | A | A | A | T | C | A | T | G | T |
| KT425067.1 | Neuroptera | *Semidalis aleyrodiformis* | C | C | G | G | T | C | T | G | A | A | C | T | C | A | A | A | T | C | A | T | G | T |
| KT425070.1 | Neuroptera | *Sisyra nigra* | C | C | G | G | T | C | T | G | A | A | C | T | C | A | A | A | T | C | A | T | G | T |
| KT425081.1 | Neuroptera | *Stenobiella* sp. | C | C | G | G | T | T | T | G | A | A | C | T | C | A | A | A | T | C | A | T | G | T |
| MH361300.1 | Neuroptera | *Suhpalacsa* | C | C | G | G | T | C | T | G | A | A | C | T | C | A | A | A | T | C | A | T | G | T |

| | | | | | | | | | | | | | | | | | | | | | | | | |
|---|---|---|---|---|---|---|---|---|---|---|---|---|---|---|---|---|---|---|---|---|---|---|---|---|
| | | *longialata* | | | | | | | | | | | | | | | | | | | | | | |
| NC039948.1 | Neuroptera | *Suhpalacsa longialata* | C | C | G | G | T | C | T | G | A | A | C | T | C | A | A | A | T | C | A | T | G | T |
| MK301247.1 | Neuroptera | *Suhpalacsa* sp. | C | C | G | G | T | C | T | G | A | A | C | T | C | A | A | A | T | C | A | T | G | T |
| MK408756.1 | Neuroptera | *Thaumatosmylus hainanus* | C | C | G | G | T | C | T | G | A | A | C | T | C | A | A | A | T | C | A | T | G | T |
| NC046578.1 | Neuroptera | *Thaumatosmylus hainanus* | C | C | G | G | T | C | T | G | A | A | C | T | C | A | A | A | T | C | A | T | G | T |
| KC515397.1 | Neuroptera | *Thyridosmylus langii* | C | C | G | G | T | C | T | G | A | A | C | T | C | A | A | A | T | C | A | T | G | T |
| NC021415.1 | Neuroptera | *Thyridosmylus langii* | C | C | G | G | T | C | T | G | A | A | C | T | C | A | A | A | T | C | A | T | G | T |
| MH020192.1 | Coleoptera | *Abscondita anceyi* | C | C | G | G | T | T | T | A | A | A | C | T | C | A | G | A | T | C | A | T | G | T |
| NC039706.1 | Coleoptera | *Abscondita anceyi* | C | C | G | G | T | T | T | A | A | A | C | T | C | A | G | A | T | C | A | T | G | T |
| MK122952.1 | Coleoptera | *Abscondita chinensis* | C | C | G | G | T | T | T | A | A | A | C | T | C | A | G | A | T | C | A | T | G | T |
| MK292092.1 | Coleoptera | *Abscondita terminalis* | C | C | G | G | T | T | T | A | A | A | C | T | C | A | G | A | T | C | A | T | G | T |
| NC044776.1 | Coleoptera | *Abscondita terminalis* | C | C | G | G | T | T | T | A | A | A | C | T | C | A | G | A | T | C | A | T | G | T |
| KT876878.1 | Coleoptera | *Acilius sulcatus* | C | C | G | G | T | C | T | G | A | A | C | T | C | A | G | A | T | C | A | T | G | T |
| KX758085.1 | Coleoptera | *Aquatica ficta* | C | C | G | G | T | T | T | A | A | A | C | T | C | A | G | A | T | C | A | T | G | T |

| | | | | | | | | | | | | | | | | | | | | | | | | |
|---|---|---|---|---|---|---|---|---|---|---|---|---|---|---|---|---|---|---|---|---|---|---|---|---|
| NC035060.1 | Coleoptera | *Aquatica ficta* | C | C | G | G | T | T | T | A | A | A | C | T | C | A | G | A | T | C | A | T | G | T |
| NC035755.1 | Coleoptera | *Aquatica lateralis* | C | C | G | G | T | T | T | A | A | A | C | T | C | A | G | A | T | C | A | T | G | T |
| KF667531.1 | Coleoptera | *Aquatica leii* | C | C | G | G | T | T | T | A | A | A | C | T | C | A | G | A | T | C | A | T | G | T |
| NC025276.1 | Coleoptera | *Aquatica leii* | C | C | G | G | T | T | T | A | A | A | C | T | C | A | G | A | T | C | A | T | G | T |
| KX758086.1 | Coleoptera | *Aquatica wuhana* | C | C | G | G | T | T | T | A | A | A | C | T | C | A | G | A | T | C | A | T | G | T |
| NC035061.1 | Coleoptera | *Aquatica wuhana* | C | C | G | G | T | T | T | A | A | A | C | T | C | A | G | A | T | C | A | T | G | T |
| KX229747.1 | Coleoptera | *Asymmetricata circumdata* | C | C | G | G | T | T | T | A | A | A | C | T | C | A | G | A | T | C | A | T | G | T |
| MK292113.1 | Coleoptera | *Asymmetricata circumdata* | C | C | G | G | T | T | T | A | A | A | C | T | C | A | G | A | T | C | A | T | G | T |
| NC032062.1 | Coleoptera | *Asymmetricata circumdata* | C | C | G | G | T | T | T | A | A | A | C | T | C | A | G | A | T | C | A | T | G | T |
| KJ922151.1 | Coleoptera | *Bicellonychia lividipennis* | C | C | G | G | T | T | T | A | A | A | C | T | C | A | G | A | T | C | A | T | G | T |
| NC030060.1 | Coleoptera | *Bicellonychia lividipennis* | C | C | G | G | T | T | T | A | A | A | C | T | C | A | G | A | T | C | A | T | G | T |
| KT876885.1 | Coleoptera | *Colymbetes fuscus* | C | C | G | G | T | C | T | G | A | A | C | T | C | A | G | A | T | C | A | T | G | T |
| MK292114.1 | Coleoptera | *Curtos bilineatus* | C | C | G | G | T | T | T | A | A | A | C | T | C | A | G | A | T | C | A | T | G | T |
| NC044789.1 | Coleoptera | *Curtos bilineatus* | C | C | G | G | T | T | T | A | A | A | C | T | C | A | G | A | T | C | A | T | G | T |
| MK609965.1 | Coleoptera | *Curtos costipennis* | C | C | G | G | T | T | T | A | A | A | C | T | C | A | G | A | T | C | A | T | G | T |
| MH651351.1 | Coleoptera | *Diaphanes citrinus* | C | C | G | G | T | T | T | A | A | A | C | T | C | A | G | A | T | C | A | T | G | T |

| | | | | | | | | | | | | | | | | | | | | | | | | |
|---|---|---|---|---|---|---|---|---|---|---|---|---|---|---|---|---|---|---|---|---|---|---|---|---|
| MK292116.1 | Coleoptera | *Diaphanes mendax* | C | C | G | A | T | T | T | A | A | A | C | T | C | A | G | A | T | C | A | T | G | T |
| NC044791.1 | Coleoptera | *Diaphanes mendax* | C | C | G | A | T | T | T | A | A | A | C | T | C | A | G | A | T | C | A | T | G | T |
| MK292094.1 | Coleoptera | *Diaphanes nubilus* | C | C | G | G | T | T | T | A | A | A | C | T | C | A | G | A | T | C | A | T | G | T |
| NC044787.1 | Coleoptera | *Diaphanes nubilus* | C | C | G | G | T | T | T | A | A | A | C | T | C | A | G | A | T | C | A | T | G | T |
| MK292118.1 | Coleoptera | *Diaphanes pectinealis* | C | C | G | G | T | T | T | A | A | A | C | T | C | A | G | A | T | C | A | T | G | T |
| NC044793.1 | Coleoptera | *Diaphanes pectinealis* | C | C | G | G | T | T | T | A | A | A | C | T | C | A | G | A | T | C | A | T | G | T |
| MK292095.1 | Coleoptera | *Diaphanes* sp. | C | C | G | G | T | T | T | A | A | A | C | T | C | A | G | A | T | C | A | T | G | T |
| MG242622.1 | Coleoptera | *Ellychnia corrusca* | C | C | G | G | T | T | T | A | A | A | C | T | C | A | G | A | T | C | A | T | G | T |
| MK292112.1 | Coleoptera | *Emeia pseudosauteri* | C | C | G | G | T | T | T | A | A | A | C | T | C | A | G | A | T | C | A | T | G | T |
| NC044778.1 | Coleoptera | *Emeia pseudosauteri* | C | C | G | G | T | T | T | A | A | A | C | T | C | A | G | A | T | C | A | T | G | T |
| KT876897.1 | Coleoptera | *Hydroporus erythrocephalus* | C | C | G | G | T | C | T | A | A | A | C | T | C | A | G | A | T | C | A | T | G | T |
| KT876896.1 | Coleoptera | *Hydroporus obscurus* | C | C | G | G | T | C | T | A | A | A | C | T | C | A | G | A | T | C | A | T | G | T |
| KT876901.1 | Coleoptera | *Hygrotus confluens* | C | C | G | G | T | C | T | G | A | A | C | T | C | A | G | A | T | C | A | T | G | T |

| | | | | | | | | | | | | | | | | | | | | | | | | |
|---|---|---|---|---|---|---|---|---|---|---|---|---|---|---|---|---|---|---|---|---|---|---|---|---|
| KT876900.1 | Coleoptera | *Hygrotus impressopunctatus* | C | C | G | G | T | C | T | G | A | A | C | T | C | A | G | A | T | C | A | T | G | T |
| KT876899.1 | Coleoptera | *Hygrotus inaequalis* | C | C | G | G | T | C | T | G | A | A | C | T | C | A | G | A | T | C | A | T | G | T |
| KX087299.1 | Coleoptera | *Hygrotus nigrolineatus* | C | C | G | G | T | C | T | G | A | A | C | T | C | A | G | A | T | C | A | T | G | T |
| MH427718.1 | Coleoptera | *Inflata indica* | C | C | G | G | T | T | T | A | A | A | C | T | C | A | G | A | T | C | A | T | G | T |
| NC039700.1 | Coleoptera | *Inflata indica* | C | C | G | G | T | T | T | A | A | A | C | T | C | A | G | A | T | C | A | T | G | T |
| MK292091.1 | Coleoptera | *Lamprigera yunnana* | C | C | G | G | T | T | T | A | A | A | C | T | C | A | G | A | T | C | A | T | G | T |
| NC044786.1 | Coleoptera | *Lamprigera yunnana* | C | C | G | G | T | T | T | A | A | A | C | T | C | A | G | A | T | C | A | T | G | T |
| KX087302.1 | Coleoptera | *Lampyris noctiluca* | C | C | G | G | T | T | T | A | A | A | C | T | C | A | G | A | T | C | A | T | G | T |
| MG912994.1 | Coleoptera | *Limbodessus palmulaoides* | C | C | G | G | T | T | T | G | A | A | C | T | C | A | G | A | T | C | A | T | G | T |
| NC037749.1 | Coleoptera | *Limbodessus palmulaoides* | C | C | G | G | T | T | T | G | A | A | C | T | C | A | G | A | T | C | A | T | G | T |
| JQ034416.1 | Coleoptera | *Limnichidae* sp. | C | C | G | G | T | T | T | A | A | A | C | T | C | A | G | A | T | C | A | T | G | T |
| KT876902.1 | Coleoptera | *Liopterus haemorrhoidalis* | C | C | G | G | T | C | T | G | A | A | C | T | C | A | G | A | T | C | A | T | G | T |
| NC022472.1 | Coleoptera | *Luciola cruciata* | C | C | G | A | T | T | T | A | A | A | C | T | C | A | G | A | T | C | A | T | G | T |
| MG770613.1 | Coleoptera | *Luciola curtithorax* | C | C | G | G | T | T | T | A | A | A | C | T | C | A | G | A | T | C | A | T | G | T |

| | | | | | | | | | | | | | | | | | | | | | | | | |
|---|---|---|---|---|---|---|---|---|---|---|---|---|---|---|---|---|---|---|---|---|---|---|---|---|
| NC038225.1 | Coleoptera | *Luciola curtithorax* | C | C | G | G | T | T | T | A | A | A | C | T | C | A | G | A | T | C | A | T | G | T |
| KP313820.1 | Coleoptera | *Luciola substriata* | C | C | G | G | T | T | T | A | A | A | C | T | C | A | G | A | T | C | A | C | G | T |
| NC027176.1 | Coleoptera | *Luciola substriata* | C | C | G | G | T | T | T | A | A | A | C | T | C | A | G | A | T | C | A | C | G | T |
| FJ859901.1 | Coleoptera | *Macrogyrus oblongus* | C | C | G | G | T | C | T | G | A | A | C | T | C | A | G | A | T | C | A | T | G | T |
| NC013249.1 | Coleoptera | *Macrogyrus oblongus* | C | C | G | G | T | C | T | G | A | A | C | T | C | A | G | A | T | C | A | T | G | T |
| MG912995.1 | Coleoptera | *Paroster macrosturtensis* | C | C | G | G | T | C | T | G | A | A | C | T | C | A | G | A | T | C | A | T | G | T |
| NC037750.1 | Coleoptera | *Paroster macrosturtensis* | C | C | G | G | T | C | T | G | A | A | C | T | C | A | G | A | T | C | A | T | G | T |
| MG912996.1 | Coleoptera | *Paroster mesosturtensis* | C | C | G | G | T | C | T | G | A | A | C | T | C | A | G | A | T | C | A | T | G | T |
| NC037751.1 | Coleoptera | *Paroster mesosturtensis* | C | C | G | G | T | C | T | G | A | A | C | T | C | A | G | A | T | C | A | T | G | T |
| MG912997.1 | Coleoptera | *Paroster microsturtensis* | C | C | G | G | T | C | T | G | A | A | C | T | C | A | G | A | T | C | A | T | G | T |
| NC037752.1 | Coleoptera | *Paroster microsturtensis* | C | C | G | G | T | C | T | G | A | A | C | T | C | A | G | A | T | C | A | T | G | T |
| KY778696.1 | Coleoptera | *Photinus pyralis* | C | C | G | G | T | T | T | A | A | A | C | T | C | A | G | A | T | C | A | T | G | T |
| MF686051.1 | Coleoptera | *Pteroptyx maipo* | C | C | G | G | T | T | T | A | A | A | C | T | C | A | G | A | T | C | A | T | G | T |
| NC036353.1 | Coleoptera | *Pteroptyx maipo* | C | C | G | G | T | T | T | A | A | A | C | T | C | A | G | A | T | C | A | T | G | T |

| | | | | | | | | | | | | | | | | | | | | | | | | |
|---|---|---|---|---|---|---|---|---|---|---|---|---|---|---|---|---|---|---|---|---|---|---|---|---|
| MK292115.1 | Coleoptera | *Pyrocoelia praetexta* | C | C | G | G | T | T | T | A | A | A | C | T | C | A | G | A | T | C | A | T | G | T |
| NC044790.1 | Coleoptera | *Pyrocoelia praetexta* | C | C | G | G | T | T | T | A | A | A | C | T | C | A | G | A | T | C | A | T | G | T |
| AF452048.1 | Coleoptera | *Pyrocoelia rufa* | C | C | G | G | T | T | T | A | A | A | C | T | C | A | G | A | T | C | A | T | G | T |
| NC003970.1 | Coleoptera | *Pyrocoelia rufa* | C | C | G | G | T | T | T | A | A | A | C | T | C | A | G | A | T | C | A | T | G | T |
| MK292117.1 | Coleoptera | *Pyrocoelia thibetana* | C | C | G | G | T | T | T | A | A | A | C | T | C | A | G | A | T | C | A | T | G | T |
| NC044792.1 | Coleoptera | *Pyrocoelia thibetana* | C | C | G | G | T | T | T | A | A | A | C | T | C | A | G | A | T | C | A | T | G | T |
| MK292111.1 | Coleoptera | *Vesta saturnalis* | C | C | G | G | T | T | T | A | A | A | C | T | C | A | G | A | T | C | A | T | G | T |
| NC044788.1 | Coleoptera | *Vesta saturnalis* | C | C | G | G | T | T | T | A | A | A | C | T | C | A | G | A | T | C | A | T | G | T |
| MF680449.1 | Trichoptera | *Anabolia bimaculata* | C | C | G | G | T | T | T | G | A | A | C | T | C | A | G | A | T | C | A | T | G | T |
| NC036004.1 | Trichoptera | *Anabolia bimaculata* | C | C | G | G | T | T | T | G | A | A | C | T | C | A | G | A | T | C | A | T | G | T |
| KF756944.1 | Trichoptera | *Apatania* sp. | C | C | G | G | T | T | T | G | A | A | C | T | C | A | G | A | T | C | A | T | G | T |
| MT410856.1 | Trichoptera | *Beraea pullata* | C | C | G | G | T | T | T | G | A | A | C | T | C | A | G | A | T | C | A | T | G | T |
| MG669126.1 | Trichoptera | *Cheumatopsyche analis* | C | C | G | G | T | T | T | G | A | A | C | T | C | A | G | A | T | C | A | T | G | T |
| NC036955.1 | Trichoptera | *Cheumatopsyche analis* | C | C | G | G | T | T | T | G | A | A | C | T | C | A | G | A | T | C | A | T | G | T |

| | | | | | | | | | | | | | | | | | | | | | | | | |
|---|---|---|---|---|---|---|---|---|---|---|---|---|---|---|---|---|---|---|---|---|---|---|---|---|
| KX385010.1 | Trichoptera | *Cheumatopsyche brevilineata* | C | C | G | G | T | T | T | G | A | A | C | T | C | A | G | A | T | C | A | T | G | T |
| NC043769.1 | Trichoptera | *Cheumatopsyche brevilineata* | C | C | G | G | T | T | T | G | A | A | C | T | C | A | G | A | T | C | A | T | G | T |
| MG669125.1 | Trichoptera | *Cheumatopsyche campyla* | C | C | G | G | T | T | T | G | A | A | C | T | C | A | G | A | T | C | A | T | G | T |
| NC036954.1 | Trichoptera | *Cheumatopsyche campyla* | C | C | G | G | T | T | T | G | A | A | C | T | C | A | G | A | T | C | A | T | G | T |
| MG669123.1 | Trichoptera | *Cheumatopsyche speciosa* | C | C | G | G | T | T | T | G | A | A | C | T | C | A | G | A | T | C | A | T | G | T |
| NC036952.1 | Trichoptera | *Cheumatopsyche speciosa* | C | C | G | G | T | T | T | G | A | A | C | T | C | A | G | A | T | C | A | T | G | T |
| KF717094.1 | Trichoptera | *Eubasilissa regina* | C | C | G | G | T | T | T | G | A | A | C | T | C | A | G | A | T | C | A | T | G | T |
| KF756943.1 | Trichoptera | *Eubasilissa regina* | C | C | G | G | T | T | T | G | A | A | C | T | C | A | G | A | T | C | A | T | G | T |
| NC023374.1 | Trichoptera | *Eubasilissa regina* | C | C | G | G | T | T | T | G | A | A | C | T | C | A | G | A | T | C | A | T | G | T |
| KX385011.1 | Trichoptera | *Hydatophylax nigrovittatus* | C | C | G | G | T | T | T | G | A | A | C | T | C | A | G | A | T | C | A | T | G | T |
| NC043770.1 | Trichoptera | *Hydatophylax nigrovittatus* | C | C | G | G | T | T | T | G | A | A | C | T | C | A | G | A | T | C | A | T | G | T |
| KF717095.1 | Trichoptera | *Hydromanicus wulaianus* | C | C | G | G | T | T | T | G | A | A | C | T | C | A | G | A | T | C | A | T | G | T |
| NC036156.1 | Trichoptera | *Hydromanicus* | C | C | G | G | T | T | T | G | A | A | C | T | C | A | G | A | T | C | A | T | G | T |

| | | | | | | | | | | | | | | | | | | | | | | | | |
|---|---|---|---|---|---|---|---|---|---|---|---|---|---|---|---|---|---|---|---|---|---|---|---|---|
| | | *wulaianus* | | | | | | | | | | | | | | | | | | | | | | |
| MT628580.1 | Trichoptera | *Hydropsyche angustipennis* | C | C | G | G | T | T | T | G | A | A | C | T | C | A | G | A | T | C | A | T | G | T |
| MG669122.1 | Trichoptera | *Hydropsyche orris* | C | C | G | G | T | T | T | G | A | A | C | T | C | A | G | A | T | C | A | T | G | T |
| NC036951.1 | Trichoptera | *Hydropsyche orris* | C | C | G | G | T | T | T | G | A | A | C | T | C | A | G | A | T | C | A | T | G | T |
| KT876876.1 | Trichoptera | *Hydropsyche pellucidula* | C | C | G | G | T | T | T | G | A | A | C | T | C | A | G | A | T | C | A | T | G | T |
| NC029246.1 | Trichoptera | *Hydropsyche pellucidula* | C | C | G | G | T | T | T | G | A | A | C | T | C | A | G | A | T | C | A | T | G | T |
| MG669121.1 | Trichoptera | *Hydropsyche simulans* | C | C | G | G | T | T | T | G | A | A | C | T | C | A | G | A | T | C | A | T | G | T |
| NC036950.1 | Trichoptera | *Hydropsyche simulans* | C | C | G | G | T | T | T | G | A | A | C | T | C | A | G | A | T | C | A | T | G | T |
| MK077681.1 | Trichoptera | *Limnephilus hyalinus* | C | C | G | G | T | T | T | G | A | A | C | T | C | A | G | A | T | C | A | T | G | T |
| NC044710.1 | Trichoptera | *Limnephilus hyalinus* | C | C | G | G | T | T | T | G | A | A | C | T | C | A | G | A | T | C | A | T | G | T |
| MG980616.1 | Trichoptera | *Phryganea cinerea* | C | C | G | G | T | T | T | G | A | A | C | T | C | A | G | A | T | C | A | T | G | T |
| NC039714.1 | Trichoptera | *Phryganea cinerea* | C | C | G | G | T | T | T | G | A | A | C | T | C | A | G | A | T | C | A | T | G | T |
| KX385012.1 | Trichoptera | *Phryganopsyche* | C | C | G | G | T | C | T | G | A | A | C | T | C | A | G | A | T | C | A | T | G | T |

| | | | | | | | | | | | | | | | | | | | | | | | | |
|---|---|---|---|---|---|---|---|---|---|---|---|---|---|---|---|---|---|---|---|---|---|---|---|---|
| | | *latipennis* | | | | | | | | | | | | | | | | | | | | | | |
| NC043771.1 | Trichoptera | *Phryganopsyche latipennis* | C | C | G | G | T | C | T | G | A | A | C | T | C | A | G | A | T | C | A | T | G | T |
| MG669124.1 | Trichoptera | *Potamyia flava* | C | C | G | G | T | T | T | G | A | A | C | T | C | A | G | A | T | C | A | T | G | T |
| NC036953.1 | Trichoptera | *Potamyia flava* | C | C | G | G | T | T | T | G | A | A | C | T | C | A | G | A | T | C | A | T | G | T |
| KP455290.1 | Trichoptera | *Sericostoma personatum* | C | C | G | G | T | T | T | G | A | A | C | T | C | A | G | A | T | C | A | T | G | T |
| KP455291.1 | Trichoptera | *Thremma gallicum* | C | C | G | G | T | T | T | G | A | A | C | T | C | A | G | A | T | C | A | T | G | T |
| AY072044.1 | Diptera | *Aedes albopictus* | C | C | G | G | T | T | T | G | A | A | C | T | C | A | G | A | T | C | A | T | G | T |
| NC030249.1 | Diptera | *Anopheles bellator* | C | C | G | G | T | T | T | G | A | A | C | T | C | A | G | A | T | C | A | T | G | T |
| NC030717.1 | Diptera | *Anopheles janconnae* | C | C | G | G | T | T | T | G | A | A | C | T | C | A | G | A | T | C | A | T | G | T |
| HQ335345.1 | Diptera | *Anopheles oryzalimnetes* | C | C | G | G | T | T | T | G | A | A | C | T | C | A | G | A | T | C | A | T | G | T |
| MF381722.1 | Diptera | *Culex bilineatus* | C | C | G | G | T | T | T | G | A | A | C | T | C | A | G | A | T | C | A | T | G | T |
| MF381718.1 | Diptera | *Culex lygrus* | C | C | G | G | T | T | T | G | A | A | C | T | C | A | G | A | T | C | A | T | G | T |
| MH316119.1 | Diptera | *Lutzia halifaxii* | C | C | G | G | T | T | T | G | A | A | C | T | C | A | G | A | T | C | A | T | G | T |
| MN626442.1 | Diptera | *Ochlerotatus taeniorhynchus* | C | C | G | G | T | T | T | G | A | A | C | T | C | A | G | A | T | C | A | T | G | T |
| KP721463.1 | Diptera | *Ochlerotatus vigilax* | C | C | G | G | T | T | T | G | A | A | C | T | C | A | G | A | T | C | A | T | G | T |
| MF957171.1 | Diptera | *Sabethes* | C | C | G | G | T | T | T | G | A | A | C | T | C | A | G | A | T | C | A | T | G | T |

| | | | | | | | | | | | | | | | | | | | | | | | | |
|---|---|---|---|---|---|---|---|---|---|---|---|---|---|---|---|---|---|---|---|---|---|---|---|---|
| | | *belisarioi* | | | | | | | | | | | | | | | | | | | | | | |
| KP793690.1 | Diptera | *Simulium aureohirtum* | C | C | G | G | T | T | T | G | A | A | C | T | C | A | G | A | T | C | A | T | G | T |
| NC029753.1 | Diptera | *Simulium aureohirtum* | C | C | G | G | T | T | T | G | A | A | C | T | C | A | G | A | T | C | A | T | G | T |
| MH729190.1 | Diptera | *Simulium maculatum* | C | C | G | G | T | T | T | G | A | A | C | T | C | A | G | A | T | C | A | T | G | T |
| NC040120.1 | Diptera | *Simulium maculatum* | C | C | G | G | T | T | T | G | A | A | C | T | C | A | G | A | T | C | A | T | G | T |
| KU252587.1 | Diptera | *Simulium variegatum* | C | C | G | G | T | T | T | G | A | A | C | T | C | A | G | A | T | C | A | T | G | T |
| NC033348.1 | Diptera | *Simulium variegatum* | C | C | G | G | T | T | T | G | A | A | C | T | C | A | G | A | T | C | A | T | G | T |
| JN861743.1 | Diptera | *Tipula abdominalis* | C | C | G | G | T | T | T | G | A | A | C | T | C | A | G | A | T | C | A | T | G | T |
| KT970065.1 | Diptera | *Tipula cockerelliana* | C | C | G | G | T | C | T | G | A | A | C | T | C | A | G | A | T | C | A | T | G | T |
| NC030520.1 | Diptera | *Tipula cockerelliana* | C | C | G | G | T | C | T | G | A | A | C | T | C | A | G | A | T | C | A | T | G | T |
| MK575491.1 | Diptera | *Uranotaenia geometrica* | C | C | G | G | T | T | T | G | A | A | C | T | C | A | G | A | T | C | A | T | G | T |
| | | A | 0.0 | 0.0 | 0.0 | 3.4 | 0.0 | 0.0 | 0.0 | 9.9 | 100.0 | 100.0 | 0.0 | 0.0 | 0.0 | 100.0 | 32.1 | 100.0 | 0.0 | 0.0 | 100.0 | 0.0 | 0.0 | 0.0 |

| | | | | | | | | | | | | | | | | | | | | | | |
|---|---|---|---|---|---|---|---|---|---|---|---|---|---|---|---|---|---|---|---|---|---|---|
| T | 0.0 | 0.0 | 0.0 | 0.0 | 99.6 | 35.4 | 100.0 | 0.5 | 0.0 | 0.0 | 0.0 | 100.0 | 0.0 | 0.0 | 0.0 | 0.0 | 100.0 | 0.0 | 0.0 | 99.5 | 0.0 | 100.0 |
| G | 0.0 | 0.0 | 100.0 | 96.6 | 0.0 | 0.0 | 0.0 | 89.6 | 0.0 | 0.0 | 0.0 | 0.0 | 0.0 | 0.0 | 67.5 | 0.0 | 0.0 | 0.0 | 0.0 | 0.0 | 100.0 | 0.0 |
| C | 100.0 | 100.0 | 0.0 | 0.0 | 0.4 | 64.6 | 0.0 | 0.0 | 0.0 | 0.0 | 100.0 | 0.0 | 100.0 | 0.0 | 0.4 | 0.0 | 0.0 | 100.0 | 0.0 | 0.5 | 0.0 | 0.0 |

Table S6. Nucleotide sequences of universal primer (MtInsects-12S_HemHol_F) and homologous region of 491 sequences of insects used for development in this study.

| | | | | | | | | | | | | | | | | | | | | | | |
|---|---|---|---|---|---|---|---|---|---|---|---|---|---|---|---|---|---|---|---|---|---|---|
| MtInsects-12S_HemHol_F | | | | G | T | G | C | C | A | G | C | H | D | Y | Y | G | C | G | G | T | T | A |
| | | | A | 0 | 0 | 0 | 0 | 0 | 491 | 0 | 0 | 267 | 141 | 0 | 1 | 0 | 0 | 0 | 0 | 0 | 2 | 491 |
| | | | T | 0 | 491 | 0 | 0 | 0 | 0 | 0 | 0 | 49 | 113 | 143 | 149 | 0 | 0 | 0 | 0 | 491 | 487 | 0 |
| | | | G | 491 | 0 | 491 | 0 | 0 | 0 | 491 | 0 | 2 | 237 | 0 | 0 | 489 | 0 | 491 | 491 | 0 | 2 | 0 |
| | | | C | 0 | 0 | 0 | 491 | 491 | 0 | 0 | 491 | 173 | 0 | 348 | 341 | 0 | 491 | 0 | 0 | 0 | 0 | 0 |
| GenBank Acc. No. | Order | Species | | | | | | | | | | | | | | | | | | | | |
| MN481418.1 | Diplura | *Campodea augens* | | G | T | G | C | C | A | G | C | C | A | C | C | G | C | G | G | T | T | A |
| DQ529236.1 | Diplura | *Campodea fragilis* | | G | T | G | C | C | A | G | C | C | A | C | C | G | C | G | G | T | T | A |
| NC008233.1 | Diplura | *Campodea fragilis* | | G | T | G | C | C | A | G | C | C | A | C | C | G | C | G | G | T | T | A |
| DQ529237.1 | Diplura | *Campodea lubbocki* | | G | T | G | C | C | A | G | C | C | A | T | C | G | C | G | G | T | T | A |
| NC008234.1 | Diplura | *Campodea lubbocki* | | G | T | G | C | C | A | G | C | C | A | T | C | G | C | G | G | T | T | A |
| AY771989.1 | Diplura | *Japyx solifugus* | | G | T | G | C | C | A | G | C | C | G | C | C | G | C | G | G | T | T | A |
| NC007214.1 | Diplura | *Japyx solifugus* | | G | T | G | C | C | A | G | C | C | G | C | C | G | C | G | G | T | T | A |
| JN990601.1 | Diplura | *Lepidocampa weberi* | | G | T | G | C | C | A | G | C | C | G | C | T | G | C | G | G | T | T | A |
| NC022675.1 | Diplura | *Lepidocampa weberi* | | G | T | G | C | C | A | G | C | C | G | C | T | G | C | G | G | T | T | A |
| JN990600.1 | Diplura | *Occasjapyx japonicus* | | G | T | G | C | C | A | G | C | C | G | C | C | G | C | G | G | T | T | A |
| NC022674.1 | Diplura | *Occasjapyx japonicus* | | G | T | G | C | C | A | G | C | C | G | C | C | G | C | G | G | T | T | A |
| JN990599.1 | Diplura | *Parajapyx emeryanus* | | G | T | G | C | C | A | G | C | C | G | C | C | G | C | G | G | T | T | A |
| NC022673.1 | Diplura | *Parajapyx emeryanus* | | G | T | G | C | C | A | G | C | C | G | C | C | G | C | G | G | T | T | A |
| AY072044.1 | Diptera | *Aedes albopictus* | | G | T | G | C | C | A | G | C | T | A | C | C | G | C | G | G | T | T | A |

| | | | | | | | | | | | | | | | | | | | | |
|---|---|---|---|---|---|---|---|---|---|---|---|---|---|---|---|---|---|---|---|---|---|
| NC030249.1 | Diptera | *Anopheles bellator* | G | T | G | C | C | A | G | C | T | A | C | T | G | C | G | G | T | T | A |
| NC030717.1 | Diptera | *Anopheles janconnae* | G | T | G | C | C | A | G | C | T | A | C | T | G | C | G | G | T | T | A |
| MF381592.1 | Diptera | *Anopheles marajoara* | G | T | G | C | C | A | G | C | T | A | C | T | G | C | G | G | T | T | A |
| HQ335345.1 | Diptera | *Anopheles oryzalimnetes* | G | T | G | C | C | A | G | C | T | A | C | T | G | C | G | G | T | T | A |
| MF381722.1 | Diptera | *Culex bilineatus* | G | T | G | C | C | A | G | C | T | A | C | C | G | C | G | G | T | T | A |
| MF381718.1 | Diptera | *Culex lygrus* | G | T | G | C | C | A | G | C | T | A | C | C | G | C | G | G | T | T | A |
| MH316119.1 | Diptera | *Lutzia halifaxii* | G | T | G | C | C | A | G | C | T | A | C | C | G | C | G | G | T | T | A |
| MN626442.1 | Diptera | *Ochlerotatus taeniorhynchus* | G | T | G | C | C | A | G | C | T | A | C | C | G | C | G | G | T | T | A |
| KP721463.1 | Diptera | *Ochlerotatus vigilax* | G | T | G | C | C | A | G | C | T | A | C | C | G | C | G | G | T | T | A |
| MF957171.1 | Diptera | *Sabethes belisarioi* | G | T | G | C | C | A | G | C | T | A | C | C | G | C | G | G | T | T | A |
| KP793690.1 | Diptera | *Simulium aureohirtum* | G | T | G | C | C | A | G | C | A | A | C | C | G | C | G | G | T | T | A |
| NC029753.1 | Diptera | *Simulium aureohirtum* | G | T | G | C | C | A | G | C | A | A | C | C | G | C | G | G | T | T | A |
| MH729190.1 | Diptera | *Simulium maculatum* | G | T | G | C | C | A | G | C | A | A | C | C | G | C | G | G | T | T | A |
| NC040120.1 | Diptera | *Simulium maculatum* | G | T | G | C | C | A | G | C | A | A | C | C | G | C | G | G | T | T | A |
| MT410847.1 | Diptera | *Simulium noelleri* | G | T | G | C | C | A | G | C | A | A | C | C | G | C | G | G | T | T | A |
| NC050320.1 | Diptera | *Simulium noelleri* | G | T | G | C | C | A | G | C | A | A | C | C | G | C | G | G | T | T | A |
| KU252587.1 | Diptera | *Simulium variegatum* | G | T | G | C | C | A | G | C | A | A | C | C | G | C | G | G | T | T | A |
| NC033348.1 | Diptera | *Simulium variegatum* | G | T | G | C | C | A | G | C | A | A | C | C | G | C | G | G | T | T | A |
| NC044661.1 | Diptera | *Trichoprosopon pallidiventer* | G | T | G | C | C | A | G | C | T | A | C | C | G | C | G | G | T | T | A |
| MK575491.1 | Diptera | *Uranotaenia geometrica* | G | T | G | C | C | A | G | C | T | A | C | T | G | C | G | G | T | T | A |

| | | | | | | | | | | | | | | | | | | | | | |
|---|---|---|---|---|---|---|---|---|---|---|---|---|---|---|---|---|---|---|---|---|---|
| MK575492.1 | Diptera | *Wyeomyia confusa* | G | T | G | C | C | A | G | C | T | A | C | C | G | C | G | G | T | T | A |
| KJ754500.1 | Archaeognatha | *Allopsontus* sp. | G | T | G | C | C | A | G | C | A | T | C | C | G | C | G | G | T | T | A |
| KJ754501.1 | Archaeognatha | *Allopsontus* sp. | G | T | G | C | C | A | G | C | A | T | C | C | G | C | G | G | T | T | A |
| AY793551.1 | Archaeognatha | *Nesomachilis australica* | G | T | G | C | C | A | G | C | A | G | C | C | G | C | G | G | T | T | A |
| NC006895.1 | Archaeognatha | *Nesomachilis australica* | G | T | G | C | C | A | G | C | A | G | C | C | G | C | G | G | T | T | A |
| EU621793.1 | Archaeognatha | *Pedetontus silvestrii* | G | T | G | C | C | A | G | C | A | G | C | C | G | C | G | G | T | T | A |
| NC011717.1 | Archaeognatha | *Pedetontus silvestrii* | G | T | G | C | C | A | G | C | A | G | C | C | G | C | G | G | T | T | A |
| KJ754503.1 | Archaeognatha | *Petrobiellus* sp. | G | T | G | C | C | A | G | C | T | G | C | C | G | C | G | G | T | A | A |
| AY956355.1 | Archaeognatha | *Petrobius brevistylis* | G | T | G | C | C | A | G | C | A | T | C | C | G | C | G | G | T | T | A |
| NC007688.1 | Archaeognatha | *Petrobius brevistylis* | G | T | G | C | C | A | G | C | A | T | C | C | G | C | G | G | T | T | A |
| JX308221.1 | Archaeognatha | *Songmachilis xinxiangensis* | G | T | G | C | C | A | G | C | A | G | C | T | G | C | G | G | T | T | A |
| NC021384.1 | Archaeognatha | *Songmachilis xinxiangensis* | G | T | G | C | C | A | G | C | A | G | C | T | G | C | G | G | T | T | A |
| EU016193.1 | Archaeognatha | *Trigoniophthalmus alternatus* | G | T | G | C | C | A | G | C | T | T | T | C | G | C | G | G | T | T | A |
| NC010532.1 | Archaeognatha | *Trigoniophthalmus alternatus* | G | T | G | C | C | A | G | C | T | T | T | C | G | C | G | G | T | T | A |
| EU084035.1 | Zygentoma | *Atelura formicaria* | G | T | G | C | C | A | G | C | A | G | C | C | G | C | G | G | T | T | A |
| NC011197.1 | Zygentoma | *Atelura formicaria* | G | T | G | C | C | A | G | C | A | G | C | C | G | C | G | G | T | T | A |
| MK301436.1 | Zygentoma | *Ctenolepisma villosa* | G | T | G | C | C | A | G | C | T | G | C | C | G | C | G | G | T | T | A |
| NC046478.1 | Zygentoma | *Ctenolepisma villosa* | G | T | G | C | C | A | G | C | T | G | C | C | G | C | G | G | T | T | A |

| | | | | | | | | | | | | | | | | | | | | |
|---|---|---|---|---|---|---|---|---|---|---|---|---|---|---|---|---|---|---|---|---|---|
| MT108230.1 | Zygentoma | *Lepisma saccharina* | G | T | G | C | C | A | G | C | T | G | C | C | G | C | G | G | T | T | A |
| NC047445.1 | Zygentoma | *Lepisma saccharina* | G | T | G | C | C | A | G | C | T | G | C | C | G | C | G | G | T | T | A |
| AY639935.1 | Zygentoma | *Thermobia domestica* | G | T | G | C | C | A | G | C | C | G | C | C | G | C | G | G | T | G | A |
| NC006080.1 | Zygentoma | *Thermobia domestica* | G | T | G | C | C | A | G | C | C | G | C | C | G | C | G | G | T | G | A |
| MK951657.1 | Zygentoma | *Thermobia* sp. | G | T | G | C | C | A | G | C | T | G | C | C | G | C | G | G | T | T | A |
| AY191994.1 | Zygentoma | *Tricholepidion gertschi* | G | T | G | C | C | A | G | C | T | G | C | C | G | C | G | G | T | T | A |
| NC005437.1 | Zygentoma | *Tricholepidion gertschi* | G | T | G | C | C | A | G | C | T | G | C | C | G | C | G | G | T | T | A |
| GU479735.1 | Ephemeroptera | *Alainites yixiani* | G | T | G | C | C | A | G | C | T | G | C | C | G | C | G | G | T | T | A |
| GU936204.1 | Ephemeroptera | *Baetis* sp. | G | T | G | C | C | A | G | C | T | G | C | C | G | C | G | G | T | T | A |
| GQ502451.1 | Ephemeroptera | *Caenis* sp. | G | T | G | C | C | A | G | C | A | G | C | C | G | C | G | G | T | T | A |
| MG910499.1 | Ephemeroptera | *Caenis* sp. | G | T | G | C | C | A | G | C | A | G | C | C | G | C | G | G | T | T | A |
| MN356096.1 | Ephemeroptera | *Caenis* sp. | G | T | G | C | C | A | G | C | A | G | C | C | G | C | G | G | T | T | A |
| MN807287.1 | Ephemeroptera | *Choroterpides apiculata* | G | T | G | C | C | A | G | C | A | G | T | T | G | C | G | G | T | T | A |
| MG870104.1 | Ephemeroptera | *Epeorus herklotsi* | G | T | G | C | C | A | G | C | C | G | C | C | G | C | G | G | T | T | A |
| MH752075.1 | Ephemeroptera | *Epeorus herklotsi* | G | T | G | C | C | A | G | C | C | G | C | C | G | C | G | G | T | T | A |
| NC039612.1 | Ephemeroptera | *Epeorus herklotsi* | G | T | G | C | C | A | G | C | C | G | C | C | G | C | G | G | T | T | A |
| KJ493406.1 | Ephemeroptera | *Epeorus* sp. | G | T | G | C | C | A | G | C | T | G | C | C | G | C | G | G | T | T | A |
| KM244708.1 | Ephemeroptera | *Epeorus* sp. | G | T | G | C | C | A | G | C | C | G | C | C | G | C | G | G | T | T | A |
| MK951674.1 | Ephemeroptera | *Epeorus* sp. | G | T | G | C | C | A | G | C | C | G | C | C | G | C | G | G | T | T | A |
| EU591678.1 | Ephemeroptera | *Ephemera orientalis* | G | T | G | C | C | A | G | C | C | A | C | C | G | C | G | G | T | T | A |
| NC012645.1 | Ephemeroptera | *Ephemera orientalis* | G | T | G | C | C | A | G | C | C | A | C | C | G | C | G | G | T | T | A |
| MK951659.1 | Ephemeroptera | *Ephemera* sp. | G | T | G | C | C | A | G | C | C | A | C | C | G | C | G | G | T | T | A |

| | | | | | | | | | | | | | | | | | | | | | |
|---|---|---|---|---|---|---|---|---|---|---|---|---|---|---|---|---|---|---|---|---|---|
| MT274127.1 | Ephemeroptera | *Ephemerella* sp. | G | T | G | C | C | A | G | C | G | A | C | C | G | C | G | G | T | T | A |
| NC050279.1 | Ephemeroptera | *Ephemerella* sp. | G | T | G | C | C | A | G | C | G | A | C | C | G | C | G | G | T | T | A |
| HM143892.1 | Ephemeroptera | *Isonychia ignota* | G | T | G | C | C | A | G | C | C | G | C | C | G | C | G | G | T | T | A |
| MH119135.1 | Ephemeroptera | *Isonychia kiangsinensis* | G | T | G | C | C | A | G | C | C | G | C | C | G | C | G | G | T | T | A |
| NC042163.1 | Ephemeroptera | *Isonychia kiangsinensis* | G | T | G | C | C | A | G | C | C | G | C | C | G | C | G | G | T | T | A |
| MK951658.1 | Ephemeroptera | *Isonychia* sp. | G | T | G | C | C | A | G | C | C | G | C | C | G | C | G | G | T | T | A |
| HM004123.1 | Ephemeroptera | *Paegniodes cupulatus* | G | T | G | C | C | A | G | C | C | G | C | C | G | C | G | G | T | T | A |
| MK951673.1 | Ephemeroptera | *Parafronurus* sp. | G | T | G | C | C | A | G | C | C | G | C | C | G | C | G | G | T | T | A |
| EU349015.1 | Ephemeroptera | *Parafronurus youi* | G | T | G | C | C | A | G | C | C | G | C | C | G | C | G | G | T | T | A |
| NC011359.1 | Ephemeroptera | *Parafronurus youi* | G | T | G | C | C | A | G | C | C | G | C | C | G | C | G | G | T | T | A |
| KM244674.1 | Ephemeroptera | *Potamanthus* sp. | G | T | G | C | C | A | G | C | C | A | C | C | G | C | G | G | T | T | A |
| MK951670.1 | Ephemeroptera | *Rhithrogena* sp. | G | T | G | C | C | A | G | C | C | G | C | C | G | C | G | G | T | T | A |
| MT274128.1 | Ephemeroptera | *Serratella* sp. | G | T | G | C | C | A | G | C | C | A | C | C | G | C | G | G | T | T | A |
| MT274129.1 | Ephemeroptera | *Serratella* sp. | G | T | G | C | C | A | G | C | A | A | C | C | G | C | G | G | T | T | A |
| NC050280.1 | Ephemeroptera | *Serratella* sp. | G | T | G | C | C | A | G | C | C | A | C | C | G | C | G | G | T | T | A |
| NC050281.1 | Ephemeroptera | *Serratella* sp. | G | T | G | C | C | A | G | C | A | A | C | C | G | C | G | G | T | T | A |
| MT274130.1 | Ephemeroptera | *Serratella zapekinae* | G | T | G | C | C | A | G | C | C | A | C | C | G | C | G | G | T | T | A |
| NC050282.1 | Ephemeroptera | *Serratella zapekinae* | G | T | G | C | C | A | G | C | C | A | C | C | G | C | G | G | T | T | A |
| FJ606783.1 | Ephemeroptera | *Siphlonurus immanis* | G | T | G | C | C | A | G | C | C | A | C | C | G | C | G | G | T | T | A |
| NC013822.1 | Ephemeroptera | *Siphlonurus immanis* | G | T | G | C | C | A | G | C | C | A | C | C | G | C | G | G | T | T | A |
| KM244684.1 | Ephemeroptera | *Siphlonurus* sp. | G | T | G | C | C | A | G | C | C | G | C | C | G | C | G | G | T | T | A |
| HQ875717.1 | Ephemeroptera | *Siphluriscus chinensis* | G | T | G | C | C | A | G | C | T | A | C | C | G | C | G | G | T | T | A |

| | | | | | | | | | | | | | | | | | | | | | |
|---|---|---|---|---|---|---|---|---|---|---|---|---|---|---|---|---|---|---|---|---|---|
| MT274131.1 | Ephemeroptera | *Torleya grandiforceps* | G | T | G | C | C | A | G | C | C | A | C | C | G | C | G | G | T | T | A |
| NC050283.1 | Ephemeroptera | *Torleya grandiforceps* | G | T | G | C | C | A | G | C | C | A | C | C | G | C | G | G | T | T | A |
| MT274132.1 | Ephemeroptera | *Torleya nepalica* | G | T | G | C | C | A | G | C | C | A | C | C | G | C | G | G | T | T | A |
| NC050284.1 | Ephemeroptera | *Torleya nepalica* | G | T | G | C | C | A | G | C | C | A | C | C | G | C | G | G | T | T | A |
| HM067837.1 | Ephemeroptera | *Vietnamella dabieshanensis* | G | T | G | C | C | A | G | C | C | A | C | C | G | C | G | G | T | T | A |
| KM244655.1 | Ephemeroptera | *Vietnamella* sp. | G | T | G | C | C | A | G | C | C | A | C | C | G | C | G | G | T | T | A |
| MH751435.1 | Odonata | *Acisoma panorpoides* | G | T | G | C | C | A | G | C | C | A | C | C | G | C | G | G | T | T | A |
| NC042689.1 | Odonata | *Acisoma panorpoides* | G | T | G | C | C | A | G | C | C | A | C | C | G | C | G | G | T | T | A |
| KX161841.1 | Odonata | *Anax imperator* | G | T | G | C | C | A | G | C | C | G | C | C | G | C | G | G | T | T | A |
| NC031821.1 | Odonata | *Anax imperator* | G | T | G | C | C | A | G | C | C | G | C | C | G | C | G | G | T | T | A |
| KP233805.1 | Odonata | *Atrocalopteryx atrata* | G | T | G | C | C | A | G | C | A | G | C | C | G | C | G | G | T | T | A |
| NC027181.1 | Odonata | *Atrocalopteryx atrata* | G | T | G | C | C | A | G | C | A | G | C | C | G | C | G | G | T | T | A |
| KM658172.1 | Odonata | *Brachythemis contaminata* | G | T | G | C | C | A | G | C | C | A | C | C | G | C | G | G | T | T | A |
| NC026305.1 | Odonata | *Brachythemis contaminata* | G | T | G | C | C | A | G | C | C | A | C | C | G | C | G | G | T | T | A |
| EU591677.1 | Odonata | *Davidius lunatus* | G | T | G | C | C | A | G | C | C | A | C | C | G | C | G | G | T | T | A |
| NC012644.1 | Odonata | *Davidius lunatus* | G | T | G | C | C | A | G | C | C | A | C | C | G | C | G | G | T | T | A |
| MH751436.1 | Odonata | *Deielia phaon* | G | T | G | C | C | A | G | C | C | A | C | C | G | C | G | G | T | T | A |
| NC042690.1 | Odonata | *Deielia phaon* | G | T | G | C | C | A | G | C | C | A | C | C | G | C | G | G | T | T | A |
| MF716899.1 | Odonata | *Enallagma cyathigerum* | G | T | G | C | C | A | G | C | A | T | C | C | G | C | G | G | T | T | A |

| | | | | | | | | | | | | | | | | | | | | | |
|---|---|---|---|---|---|---|---|---|---|---|---|---|---|---|---|---|---|---|---|---|---|
| JX050223.1 | Odonata | *Epiophlebia superstes* | G | T | G | C | C | A | G | C | C | A | C | C | G | C | G | G | T | T | A |
| NC023232.1 | Odonata | *Epiophlebia superstes* | G | T | G | C | C | A | G | C | C | A | C | C | G | C | G | G | T | T | A |
| MK522522.1 | Odonata | *Epophthalmia elegans* | G | T | G | C | C | A | G | C | C | G | C | C | G | C | G | G | T | T | A |
| NC046756.1 | Odonata | *Epophthalmia elegans* | G | T | G | C | C | A | G | C | C | G | C | C | G | C | G | G | T | T | A |
| HM126547.1 | Odonata | *Euphaea formosa* | G | T | G | C | C | A | G | C | A | G | T | C | K | C | G | G | T | T | A |
| NC014493.1 | Odonata | *Euphaea formosa* | G | T | G | C | C | A | G | C | A | G | T | C | K | C | G | G | T | T | A |
| KM244659.1 | Odonata | *Hydrobasileus croceus* | G | T | G | C | C | A | G | C | C | G | C | C | G | C | G | G | T | T | A |
| NC025758.1 | Odonata | *Hydrobasileus croceus* | G | T | G | C | C | A | G | C | C | G | C | C | G | C | G | G | T | T | A |
| KU958378.1 | Odonata | *Ischnura elegans* | G | T | G | C | C | A | G | C | A | T | C | C | G | C | G | G | T | T | A |
| MK951668.1 | Odonata | *Ischnura elegans* | G | T | G | C | C | A | G | C | A | T | C | C | G | C | G | G | T | T | A |
| NC031824.1 | Odonata | *Ischnura elegans* | G | T | G | C | C | A | G | C | A | T | C | C | G | C | G | G | T | T | A |
| KC878732.1 | Odonata | *Ischnura pumilio* | G | T | G | C | C | A | G | C | A | T | C | C | G | C | G | G | T | T | A |
| NC021617.1 | Odonata | *Ischnura pumilio* | G | T | G | C | C | A | G | C | A | T | C | C | G | C | G | G | T | T | A |
| MF990748.1 | Odonata | *Macromia daimoji* | G | T | G | C | C | A | G | C | C | G | C | C | G | C | G | G | T | T | A |
| NC041425.1 | Odonata | *Macromia daimoji* | G | T | G | C | C | A | G | C | C | G | C | C | G | C | G | G | T | T | A |
| MK722304.1 | Odonata | *Matrona basilaris* | G | T | G | C | C | A | G | C | A | G | C | C | G | C | G | G | T | T | A |
| MK722305.1 | Odonata | *Matrona basilaris* | G | T | G | C | C | A | G | C | A | G | C | C | G | C | G | G | T | T | A |
| KU958377.1 | Odonata | *Megaloprepus caerulatus* | G | T | G | C | C | A | G | C | A | T | C | C | G | C | G | G | T | T | A |
| NC031823.1 | Odonata | *Megaloprepus caerulatus* | G | T | G | C | C | A | G | C | A | T | C | C | G | C | G | G | T | T | A |
| KU871065.1 | Odonata | *Mnais costalis* | G | T | G | C | C | A | G | C | A | G | C | C | G | C | G | G | T | T | A |
| KY402222.1 | Odonata | *Nannophya pygmaea* | G | T | G | C | C | A | G | C | C | A | C | C | G | C | G | G | T | T | A |
| NC039410.1 | Odonata | *Nannophya pygmaea* | G | T | G | C | C | A | G | C | C | A | C | C | G | C | G | G | T | T | A |

| | | | | | | | | | | | | | | | | | | | | |
|---|---|---|---|---|---|---|---|---|---|---|---|---|---|---|---|---|---|---|---|---|---|
| KU361233.1 | Odonata | *Orthetrum chrysis* | G | T | G | C | C | A | G | C | C | A | C | C | G | C | G | G | T | T | A |
| NC032048.1 | Odonata | *Orthetrum chrysis* | G | T | G | C | C | A | G | C | C | A | C | C | G | C | G | G | T | T | A |
| KU361232.1 | Odonata | *Orthetrum glaucum* | G | T | G | C | C | A | G | C | C | A | C | C | G | C | G | G | T | T | A |
| NC032047.1 | Odonata | *Orthetrum glaucum* | G | T | G | C | C | A | G | C | C | A | C | C | G | C | G | G | T | T | A |
| MH751437.1 | Odonata | *Orthetrum melania* | G | T | G | C | C | A | G | C | C | A | C | C | G | C | G | G | T | T | A |
| NC042732.1 | Odonata | *Orthetrum melania* | G | T | G | C | C | A | G | C | C | A | C | C | G | C | G | G | T | T | A |
| KU361234.1 | Odonata | *Orthetrum sabina* | G | T | G | C | C | A | G | C | C | A | C | C | G | C | G | G | T | T | A |
| MH751438.1 | Odonata | *Orthetrum sabina* | G | T | G | C | C | A | G | C | C | A | C | C | G | C | G | G | T | T | A |
| NC032049.1 | Odonata | *Orthetrum sabina* | G | T | G | C | C | A | G | C | C | A | C | C | G | C | G | G | T | T | A |
| KU361235.1 | Odonata | *Orthetrum testaceum* | G | T | G | C | C | A | G | C | C | A | C | C | G | C | G | G | T | T | A |
| MH751439.1 | Odonata | *Orthetrum testaceum* | G | T | G | C | C | A | G | C | C | A | C | C | G | C | G | G | T | T | A |
| NC032050.1 | Odonata | *Orthetrum testaceum* | G | T | G | C | C | A | G | C | C | A | C | C | G | C | G | G | T | T | A |
| MK951669.1 | Odonata | *Paracercion v-nigrum* | G | T | G | C | C | A | G | C | A | G | C | C | G | C | G | G | T | T | A |
| KP233804.1 | Odonata | *Platycnemis foliacea* | G | T | G | C | C | A | G | C | A | A | C | C | G | C | G | G | T | T | A |
| NC027180.1 | Odonata | *Platycnemis foliacea* | G | T | G | C | C | A | G | C | A | A | C | C | G | C | G | G | T | T | A |
| FJ606784.1 | Odonata | *Pseudolestes mirabilis* | G | T | G | C | C | A | G | C | A | A | C | C | G | C | G | G | T | T | A |
| NC020636.1 | Odonata | *Pseudolestes mirabilis* | G | T | G | C | C | A | G | C | A | A | C | C | G | C | G | G | T | T | A |
| MF150044.1 | Odonata | *Psolodesmus mandarinus* | G | T | G | C | C | A | G | C | A | G | T | C | G | C | G | G | T | T | A |
| NC042442.1 | Odonata | *Psolodesmus mandarinus* | G | T | G | C | C | A | G | C | A | G | T | C | G | C | G | G | T | T | A |
| MG594801.1 | Odonata | *Somatochlora hineana* | G | T | G | C | C | A | G | C | T | G | C | C | G | C | G | G | T | T | A |

| | | | | | | | | | | | | | | | | | | | | | |
|---|---|---|---|---|---|---|---|---|---|---|---|---|---|---|---|---|---|---|---|---|---|
| MH751440.1 | Odonata | *Tramea virginia* | G | T | G | C | C | A | G | C | C | A | C | C | G | C | G | G | T | T | A |
| NC042691.1 | Odonata | *Tramea virginia* | G | T | G | C | C | A | G | C | C | A | C | C | G | C | G | G | T | T | A |
| MH751441.1 | Odonata | *Trigomphus carus* | G | T | G | C | C | A | G | C | C | A | C | C | G | C | G | G | T | T | A |
| NC042733.1 | Odonata | *Trigomphus carus* | G | T | G | C | C | A | G | C | C | A | C | C | G | C | G | G | T | T | A |
| JX050224.1 | Odonata | *Vestalis melania* | G | T | G | C | C | A | G | C | A | G | T | C | G | C | G | G | T | T | A |
| NC023233.1 | Odonata | *Vestalis melania* | G | T | G | C | C | A | G | C | A | G | T | C | G | C | G | G | T | T | A |
| KM199685.1 | Plecoptera | *Acroneuria hainana* | G | T | G | C | C | A | G | C | C | G | C | C | G | C | G | G | T | T | A |
| NC026104.1 | Plecoptera | *Acroneuria hainana* | G | T | G | C | C | A | G | C | C | G | C | C | G | C | G | G | T | T | A |
| KX091846.1 | Plecoptera | *Agnetina aequalis* | G | T | G | C | C | A | G | C | A | G | C | C | G | C | G | G | T | T | A |
| MH085446.1 | Plecoptera | *Amphinemura longispina* | G | T | G | C | C | A | G | C | C | G | C | C | G | C | G | G | T | T | A |
| NC044748.1 | Plecoptera | *Amphinemura longispina* | G | T | G | C | C | A | G | C | C | G | C | C | G | C | G | G | T | T | A |
| KX091847.1 | Plecoptera | *Amphinemura* sp. | G | T | G | C | C | A | G | C | C | G | C | C | G | C | G | G | T | T | A |
| MH085447.1 | Plecoptera | *Amphinemura yao* | G | T | G | C | C | A | G | C | C | G | C | C | G | C | G | G | T | T | A |
| NC044749.1 | Plecoptera | *Amphinemura yao* | G | T | G | C | C | A | G | C | C | G | C | C | G | C | G | G | T | T | A |
| MK111413.1 | Plecoptera | *Antarctoperla michaelseni* | G | T | G | C | C | A | G | C | A | G | C | C | G | C | G | G | T | T | A |
| NC042199.1 | Plecoptera | *Antarctoperla michaelseni* | G | T | G | C | C | A | G | C | A | G | C | C | G | C | G | G | T | T | A |
| KR604721.1 | Plecoptera | *Apteroperla tikumana* | G | T | G | C | C | A | G | C | C | G | C | C | G | C | G | G | T | T | A |
| NC027698.1 | Plecoptera | *Apteroperla tikumana* | G | T | G | C | C | A | G | C | C | G | C | C | G | C | G | G | T | T | A |
| MT410838.1 | Plecoptera | *Brachyptera risi* | G | T | G | C | C | A | G | C | C | G | C | C | G | C | G | G | T | T | A |
| MG677941.1 | Plecoptera | *Calineuria stigmatica* | G | T | G | C | C | A | G | C | C | G | C | C | G | C | G | G | T | T | A |

| | | | | | | | | | | | | | | | | | | | | | |
|---|---|---|---|---|---|---|---|---|---|---|---|---|---|---|---|---|---|---|---|---|---|
| KX094942.1 | Plecoptera | *Capnia zijinshana* | G | T | G | C | C | A | G | C | C | G | C | C | G | C | G | G | T | T | A |
| NC034661.1 | Plecoptera | *Capnia zijinshana* | G | T | G | C | C | A | G | C | C | G | C | C | G | C | G | G | T | T | A |
| MG677942.1 | Plecoptera | *Caroperla siveci* | G | T | G | C | C | A | G | C | T | A | C | C | G | C | G | G | T | T | A |
| MF100783.1 | Plecoptera | *Cerconychia flectospina* | G | T | G | C | C | A | G | C | T | G | C | C | G | C | G | G | T | T | A |
| MN419914.1 | Plecoptera | *Claassenia* sp. | G | T | G | C | C | A | G | C | C | G | C | C | G | C | G | G | T | T | A |
| MK111416.1 | Plecoptera | *Diamphipnoa annulata* | G | T | G | C | C | A | G | C | T | G | C | C | G | C | G | G | T | T | A |
| NC042205.1 | Plecoptera | *Diamphipnoa annulata* | G | T | G | C | C | A | G | C | T | G | C | C | G | C | G | G | T | T | A |
| MK111417.1 | Plecoptera | *Diamphipnopsis sp.* | G | T | G | C | C | A | G | C | T | G | C | C | G | C | G | G | T | T | A |
| KF484757.1 | Plecoptera | *Dinocras cephalotes* | G | T | G | C | C | A | G | C | A | G | C | C | G | C | G | G | T | T | A |
| NC022843.1 | Plecoptera | *Dinocras cephalotes* | G | T | G | C | C | A | G | C | A | G | C | C | G | C | G | G | T | T | A |
| MG589787.1 | Plecoptera | *Doddsia occidentalis* | G | T | G | C | C | A | G | C | C | G | C | C | G | C | G | G | T | T | A |
| MK905888.1 | Plecoptera | *Etrocorema hochii* | G | T | G | C | C | A | G | C | T | G | C | C | G | C | G | G | T | T | A |
| MK905206.1 | Plecoptera | *Flavoperla* sp. | G | T | G | C | C | A | G | C | C | A | C | C | G | C | G | G | T | T | A |
| MN419916.1 | Plecoptera | *Flavoperla* sp. | G | T | G | C | C | A | G | C | C | A | C | C | G | C | G | G | T | T | A |
| MN419915.1 | Plecoptera | *Indonemoura auriformis* | G | T | G | C | C | A | G | C | C | G | C | C | G | C | G | G | T | T | A |
| MH085448.1 | Plecoptera | *Indonemoura jacobsoni* | G | T | G | C | C | A | G | C | C | G | C | C | G | C | G | G | T | T | A |
| NC044750.1 | Plecoptera | *Indonemoura jacobsoni* | G | T | G | C | C | A | G | C | C | G | C | C | G | C | G | G | T | T | A |
| MH085449.1 | Plecoptera | *Indonemoura nohirae* | G | T | G | C | C | A | G | C | T | G | C | C | G | C | G | G | T | T | A |
| NC044751.1 | Plecoptera | *Indonemoura nohirae* | G | T | G | C | C | A | G | C | T | G | C | C | G | C | G | G | T | T | A |
| MF716959.1 | Plecoptera | *Isoperla bilineata* | G | T | G | C | C | A | G | C | C | G | C | C | G | C | G | G | T | T | A |
| NC038190.1 | Plecoptera | *Isoperla bilineata* | G | T | G | C | C | A | G | C | C | G | C | C | G | C | G | G | T | T | A |
| MG910457.1 | Plecoptera | *Isoperla eximia* | G | T | G | C | C | A | G | C | C | G | C | C | G | C | G | G | T | T | A |

| | | | | | | | | | | | | | | | | | | | | |
|---|---|---|---|---|---|---|---|---|---|---|---|---|---|---|---|---|---|---|---|---|
| NC038167.1 | Plecoptera | *Isoperla eximia* | G | T | G | C | C | A | G | C | C | G | C | C | G | C | G | G | T | T | A |
| KT186102.1 | Plecoptera | *Kamimuria chungnanshana* | G | T | G | C | C | A | G | C | C | G | C | C | G | C | G | G | T | T | A |
| NC028076.1 | Plecoptera | *Kamimuria chungnanshana* | G | T | G | C | C | A | G | C | C | G | C | C | G | C | G | G | T | T | A |
| MN400755.1 | Plecoptera | *Kamimuria klapaleki* | G | T | G | C | C | A | G | C | C | G | C | C | G | C | G | G | T | T | A |
| KC894944.1 | Plecoptera | *Kamimuria wangi* | G | T | G | C | C | A | G | C | C | G | C | C | G | C | G | G | T | T | A |
| MK111414.1 | Plecoptera | *Klapopteryx armillata* | G | T | G | C | C | A | G | C | A | G | C | C | G | C | G | G | T | T | A |
| KP642637.1 | Plecoptera | *Mesocapnia arizonensis* | G | T | G | C | C | A | G | C | C | G | C | C | G | C | G | G | T | T | A |
| KY568983.1 | Plecoptera | *Mesocapnia daxingana* | G | T | G | C | C | A | G | C | C | G | C | C | G | C | G | G | T | T | A |
| MH085450.1 | Plecoptera | *Mesonemoura metafiligera* | G | T | G | C | C | A | G | C | C | G | C | C | G | C | G | G | T | T | A |
| NC044719.1 | Plecoptera | *Mesonemoura metafiligera* | G | T | G | C | C | A | G | C | C | G | C | C | G | C | G | G | T | T | A |
| MH085451.1 | Plecoptera | *Mesonemoura tritaenia* | G | T | G | C | C | A | G | C | C | G | C | C | G | C | G | G | T | T | A |
| NC044720.1 | Plecoptera | *Mesonemoura tritaenia* | G | T | G | C | C | A | G | C | C | G | C | C | G | C | G | G | T | T | A |
| MN096323.1 | Plecoptera | *Microperla geei* | G | T | G | C | C | A | G | C | T | G | C | C | G | C | G | G | T | T | A |
| MT410776.1 | Plecoptera | *Nemoura avicularis* | G | T | G | C | C | A | G | C | C | G | C | C | G | C | G | G | T | T | A |
| MT410849.1 | Plecoptera | *Nemoura cinerea* | G | T | G | C | C | A | G | C | C | G | C | C | G | C | G | G | T | T | A |
| KY940360.1 | Plecoptera | *Nemoura nankinensis* | G | T | G | C | C | A | G | C | C | G | C | C | G | C | G | G | T | T | A |
| NC034939.1 | Plecoptera | *Nemoura nankinensis* | G | T | G | C | C | A | G | C | C | G | C | C | G | C | G | G | T | T | A |
| MK290826.1 | Plecoptera | *Nemoura papilla* | G | T | G | C | C | A | G | C | C | G | C | C | G | C | G | G | T | T | A |

| | | | | | | | | | | | | | | | | | | | | | |
|---|---|---|---|---|---|---|---|---|---|---|---|---|---|---|---|---|---|---|---|---|---|
| MK111418.1 | Plecoptera | *Neonemura barrosi* | G | T | G | C | C | A | G | C | C | G | C | C | G | C | G | G | T | T | A |
| NC042206.1 | Plecoptera | *Neonemura barrosi* | G | T | G | C | C | A | G | C | C | G | C | C | G | C | G | G | T | T | A |
| KX091858.1 | Plecoptera | *Neoperla ignacsiveci* | G | T | G | C | C | A | G | C | T | G | C | C | G | C | G | G | T | T | A |
| KX091859.1 | Plecoptera | *Neoperla* sp. | G | T | G | C | C | A | G | C | A | G | C | C | G | C | G | G | T | T | A |
| MN400756.1 | Plecoptera | *Neoperlops gressitti* | G | T | G | C | C | A | G | C | T | G | C | C | G | C | G | G | T | T | A |
| MK111415.1 | Plecoptera | *Neuroperla schedingi* | G | T | G | C | C | A | G | C | T | G | C | C | G | C | G | G | T | T | A |
| NC042200.1 | Plecoptera | *Neuroperla schedingi* | G | T | G | C | C | A | G | C | T | G | C | C | G | C | G | G | T | T | A |
| MK686067.1 | Plecoptera | *Niponiella limbatella* | G | T | G | C | C | A | G | C | C | G | C | C | G | C | G | G | T | T | A |
| MK387068.1 | Plecoptera | *Peltoperlopsis cebuano* | G | T | G | C | C | A | G | C | T | G | C | C | G | C | G | G | T | T | A |
| MF197377.1 | Plecoptera | *Perlodes* sp. | G | T | G | C | C | A | G | C | C | G | C | C | G | C | G | G | T | T | A |
| MH085452.1 | Plecoptera | *Protonemura kohnoae* | G | T | G | C | C | A | G | C | C | G | C | C | G | C | G | G | T | T | A |
| NC044752.1 | Plecoptera | *Protonemura kohnoae* | G | T | G | C | C | A | G | C | C | G | C | C | G | C | G | G | T | T | A |
| MT410850.1 | Plecoptera | *Protonemura meyeri* | G | T | G | C | C | A | G | C | C | G | C | C | G | C | G | G | T | T | A |
| NC050322.1 | Plecoptera | *Protonemura meyeri* | G | T | G | C | C | A | G | C | C | G | C | C | G | C | G | G | T | T | A |
| MH085453.1 | Plecoptera | *Protonemura orbiculata* | G | T | G | C | C | A | G | C | C | G | C | C | G | C | G | G | T | T | A |
| NC044753.1 | Plecoptera | *Protonemura orbiculata* | G | T | G | C | C | A | G | C | C | G | C | C | G | C | G | G | T | T | A |
| MG910458.1 | Plecoptera | *Pseudomegarcys japonica* | G | T | G | C | C | A | G | C | C | G | C | C | G | C | G | G | T | T | A |
| NC038168.1 | Plecoptera | *Pseudomegarcys japonica* | G | T | G | C | C | A | G | C | C | G | C | C | G | C | G | G | T | T | A |
| KU182360.1 | Plecoptera | *Pteronarcella badia* | G | T | G | C | C | A | G | C | C | G | C | C | G | C | G | G | T | T | A |
| NC029248.1 | Plecoptera | *Pteronarcella badia* | G | T | G | C | C | A | G | C | C | G | C | C | G | C | G | G | T | T | A |

| | | | | | | | | | | | | | | | | | | | | | |
|---|---|---|---|---|---|---|---|---|---|---|---|---|---|---|---|---|---|---|---|---|---|
| AY687866.1 | Plecoptera | *Pteronarcys princeps* | G | T | G | C | C | A | G | C | C | G | C | C | G | C | G | G | T | T | A |
| NC006133.1 | Plecoptera | *Pteronarcys princeps* | G | T | G | C | C | A | G | C | C | G | C | C | G | C | G | G | T | T | A |
| MK111419.1 | Plecoptera | *Rhopalopsole bulbifera* | G | T | G | C | C | A | G | C | A | G | C | C | G | C | G | G | T | T | A |
| NC042207.1 | Plecoptera | *Rhopalopsole bulbifera* | G | T | G | C | C | A | G | C | A | G | C | C | G | C | G | G | T | T | A |
| MH510071.1 | Plecoptera | *Scopura longa* | G | T | G | C | C | A | G | C | C | G | C | C | G | C | G | G | T | T | A |
| NC041105.1 | Plecoptera | *Scopura longa* | G | T | G | C | C | A | G | C | C | G | C | C | G | C | G | G | T | T | A |
| MK492253.1 | Plecoptera | *Sinacroneuria dabieshana* | G | T | G | C | C | A | G | C | C | G | C | C | G | C | G | G | T | T | A |
| MF716958.1 | Plecoptera | *Soliperla* sp. | G | T | G | C | C | A | G | C | T | G | C | C | G | C | G | G | T | T | A |
| NC038189.1 | Plecoptera | *Soliperla* sp. | G | T | G | C | C | A | G | C | T | G | C | C | G | C | G | G | T | T | A |
| MH085455.1 | Plecoptera | *Sphaeronemoura acutispina* | G | T | G | C | C | A | G | C | C | G | C | C | G | C | G | G | T | T | A |
| MH085454.1 | Plecoptera | *Sphaeronemoura grandicauda* | G | T | G | C | C | A | G | C | C | G | C | C | G | C | G | G | T | T | A |
| NC044754.1 | Plecoptera | *Sphaeronemoura grandicauda* | G | T | G | C | C | A | G | C | C | G | C | C | G | C | G | G | T | T | A |
| MK111420.1 | Plecoptera | *Sphaeronemoura hainana* | G | T | G | C | C | A | G | C | C | G | C | C | G | C | G | G | T | T | A |
| NC044755.1 | Plecoptera | *Sphaeronemoura hamistyla* | G | T | G | C | C | A | G | C | C | G | C | C | G | C | G | G | T | T | A |
| KR088971.1 | Plecoptera | *Styloperla* sp. | G | T | G | C | C | A | G | C | T | G | C | C | G | C | G | G | T | T | A |
| KX845569.1 | Plecoptera | *Styloperla spinicercia* | G | T | G | C | C | A | G | C | T | G | C | C | G | C | G | G | T | T | A |

| | | | | | | | | | | | | | | | | | | | | | |
|---|---|---|---|---|---|---|---|---|---|---|---|---|---|---|---|---|---|---|---|---|---|
| NC034809.1 | Plecoptera | *Styloperla spinicercia* | G | T | G | C | C | A | G | C | T | G | C | C | G | C | G | G | T | T | A |
| KX091865.1 | Plecoptera | *Suwallia bimaculata* | G | T | G | C | C | A | G | C | C | A | C | C | G | C | G | G | T | T | A |
| MN121757.1 | Plecoptera | *Suwallia bimaculata* | G | T | G | C | C | A | G | C | C | A | C | C | G | C | G | G | T | T | A |
| MF198253.1 | Plecoptera | *Suwallia teleckojensis* | G | T | G | C | C | A | G | C | C | A | C | C | G | C | G | G | T | T | A |
| NC037754.1 | Plecoptera | *Suwallia teleckojensis* | G | T | G | C | C | A | G | C | C | A | C | C | G | C | G | G | T | T | A |
| KM216826.1 | Plecoptera | *Sweltsa longistyla* | G | T | G | C | C | A | G | C | C | A | C | C | G | C | G | G | T | T | A |
| MG589786.1 | Plecoptera | *Taeniopteryx ugola* | G | T | G | C | C | A | G | C | C | G | C | C | G | C | G | G | T | T | A |
| NC037897.1 | Plecoptera | *Taeniopteryx ugola* | G | T | G | C | C | A | G | C | C | G | C | C | G | C | G | G | T | T | A |
| KM409708.1 | Plecoptera | *Togoperla* sp. | G | T | G | C | C | A | G | C | C | G | C | C | G | C | G | G | T | T | A |
| KY522907.1 | Plecoptera | *Zelandoperla fenestrata* | G | T | G | C | C | A | G | C | A | G | C | C | G | C | G | G | T | T | A |
| NC034997.1 | Plecoptera | *Zelandoperla fenestrata* | G | T | G | C | C | A | G | C | A | G | C | C | G | C | G | G | T | T | A |
| FJ456939.1 | Heteroptera | *Aphelocheirus ellipsoideus* | G | T | G | C | C | A | G | C | A | G | T | C | G | C | G | G | T | T | A |
| KX505849.1 | Heteroptera | *Aphelocheirus jendeki* | G | T | G | C | C | A | G | C | A | G | T | C | G | C | G | G | T | T | A |
| NC037370.1 | Heteroptera | *Aphelocheirus jendeki* | G | T | G | C | C | A | G | C | A | G | T | C | G | C | G | G | T | T | A |
| FJ456944.1 | Heteroptera | *Aquarius paludum* | G | T | G | C | C | A | G | C | C | G | T | C | G | C | G | G | T | T | A |
| NC012841.1 | Heteroptera | *Aquarius paludum* | G | T | G | C | C | A | G | C | C | G | T | C | G | C | G | G | T | T | A |
| MK251109.1 | Heteroptera | *Diplonychus esakii* | G | T | G | C | C | A | G | C | A | T | T | C | G | C | G | G | T | T | A |
| FJ456940.1 | Heteroptera | *Diplonychus rusticus* | G | T | G | C | C | A | G | C | A | T | T | C | G | C | G | G | T | T | A |
| FJ456949.1 | Heteroptera | *Enithares tibialis* | G | T | G | C | C | A | G | C | A | G | T | C | G | C | G | G | T | T | A |
| NC012819.1 | Heteroptera | *Enithares tibialis* | G | T | G | C | C | A | G | C | A | G | T | C | G | C | G | G | T | T | A |
| KP400582.1 | Heteroptera | *Entomovelia* sp. | G | T | G | C | C | A | G | C | A | G | T | C | G | C | G | G | T | T | A |

| | | | | | | | | | | | | | | | | | | | | |
|---|---|---|---|---|---|---|---|---|---|---|---|---|---|---|---|---|---|---|---|---|---|
| MF177288.1 | Heteroptera | *Gigantometra gigas* | G | T | G | C | C | A | G | C | C | G | T | C | G | C | G | G | T | T | A |
| MF177289.1 | Heteroptera | *Gigantometra gigas* | G | T | G | C | C | A | G | C | C | G | T | C | G | C | G | G | T | T | A |
| MF177290.1 | Heteroptera | *Gigantometra gigas* | G | T | G | C | C | A | G | C | C | G | T | C | G | C | G | G | T | T | A |
| NC041084.1 | Heteroptera | *Gigantometra gigas* | G | T | G | C | C | A | G | C | C | G | T | C | G | C | G | G | T | T | A |
| KJ027513.1 | Heteroptera | *Helotrephes semiglobosus semiglobosus* | G | T | G | C | C | A | G | C | A | G | T | C | G | C | G | G | T | T | A |
| FJ456951.1 | Heteroptera | *Helotrephes* sp. | G | T | G | C | C | A | G | C | A | G | T | C | G | C | G | G | T | T | A |
| NC012822.1 | Heteroptera | *Helotrephes* sp. | G | T | G | C | C | A | G | C | A | G | T | C | G | C | G | G | T | T | A |
| FJ456945.1 | Heteroptera | *Hydrometra greeni* | G | T | G | C | C | A | G | C | A | T | C | C | G | C | G | G | T | T | A |
| NC012842.1 | Heteroptera | *Hydrometra greeni* | G | T | G | C | C | A | G | C | A | T | C | C | G | C | G | G | T | T | A |
| FJ456947.1 | Heteroptera | *Ilyocoris cimicoides* | G | T | G | C | C | A | G | C | A | G | T | C | G | C | G | G | T | T | A |
| NC012845.1 | Heteroptera | *Ilyocoris cimicoides* | G | T | G | C | C | A | G | C | A | G | T | C | G | C | G | G | T | T | A |
| FJ456948.1 | Heteroptera | *Laccotrephes robustus* | G | T | G | C | C | A | G | C | A | G | T | C | G | C | G | G | T | T | A |
| NC012817.1 | Heteroptera | *Laccotrephes robustus* | G | T | G | C | C | A | G | C | A | G | T | C | G | C | G | G | T | T | A |
| FJ456946.1 | Heteroptera | *Leptopus* sp. | G | T | G | C | C | A | G | C | A | G | T | C | G | C | G | G | T | T | A |
| JQ910985.1 | Heteroptera | *Lethocerus deyrollei* | G | T | G | C | C | A | G | C | A | G | C | C | G | C | G | G | T | T | A |
| KU237288.1 | Heteroptera | *Lethocerus deyrollei* | G | T | G | C | C | A | G | C | A | G | C | C | G | C | G | G | T | T | A |
| KM588201.1 | Heteroptera | *Lethocerus indicus* | G | T | G | C | C | A | G | C | A | G | C | C | G | C | G | G | T | T | A |
| KR072671.1 | Heteroptera | *Lethocerus indicus* | G | T | G | C | C | A | G | C | A | G | C | C | G | C | G | G | T | T | A |
| NC027194.1 | Heteroptera | *Lethocerus indicus* | G | T | G | C | C | A | G | C | A | G | C | C | G | C | G | G | T | T | A |
| MN027272.1 | Heteroptera | *Limnogonus hypoleucus* | G | T | G | C | C | A | G | C | C | G | T | C | G | C | G | G | T | T | A |

| | | | | | | | | | | | | | | | | | | | | | |
|---|---|---|---|---|---|---|---|---|---|---|---|---|---|---|---|---|---|---|---|---|---|
| NC044722.1 | Heteroptera | *Limnogonus hypoleucus* | G | T | G | C | C | A | G | C | C | G | T | C | G | C | G | G | T | T | A |
| MN027275.1 | Heteroptera | *Limnogonus intermedius* | G | T | G | C | C | A | G | C | C | G | T | C | G | C | G | G | T | T | A |
| NC044724.1 | Heteroptera | *Limnogonus intermedius* | G | T | G | C | C | A | G | C | C | G | T | C | G | C | G | G | T | T | A |
| MN027273.1 | Heteroptera | *Limnogonus poissoni* | G | T | G | C | C | A | G | C | C | G | T | C | G | C | G | G | T | T | A |
| MN027274.1 | Heteroptera | *Limnogonus poissoni* | G | T | G | C | C | A | G | C | C | G | T | C | G | C | G | G | T | T | A |
| MN027276.1 | Heteroptera | *Limnogonus poissoni* | G | T | G | C | C | A | G | C | C | G | T | C | G | C | G | G | T | T | A |
| NC044723.1 | Heteroptera | *Limnogonus poissoni* | G | T | G | C | C | A | G | C | C | G | T | C | G | C | G | G | T | T | A |
| FJ456942.1 | Heteroptera | *Lycorma delicatula* | G | T | G | C | C | A | G | C | A | T | T | T | G | C | G | G | T | T | A |
| NC012835.1 | Heteroptera | *Lycorma delicatula* | G | T | G | C | C | A | G | C | A | T | T | T | G | C | G | G | T | T | A |
| MN027278.1 | Heteroptera | *Metrocoris* sp. | G | T | G | C | C | A | G | C | T | G | T | C | G | C | G | G | T | T | A |
| MH382835.1 | Heteroptera | *Micronecta sahlbergii* | G | T | G | C | C | A | G | C | A | G | T | C | G | C | G | G | T | T | A |
| NC039588.1 | Heteroptera | *Micronecta sahlbergii* | G | T | G | C | C | A | G | C | A | G | T | C | G | C | G | G | T | T | A |
| KT031802.1 | Heteroptera | *Nepa hoffmanni* | G | T | G | C | C | A | G | C | A | G | C | C | G | C | G | G | T | T | A |
| NC028084.1 | Heteroptera | *Nepa hoffmanni* | G | T | G | C | C | A | G | C | A | G | C | C | G | C | G | G | T | T | A |
| FJ456943.1 | Heteroptera | *Nerthra indica* | G | T | G | C | C | A | G | C | A | T | T | C | G | C | G | G | T | T | A |
| NC012838.1 | Heteroptera | *Nerthra indica* | G | T | G | C | C | A | G | C | A | T | T | C | G | C | G | G | T | T | A |
| KX034086.1 | Heteroptera | *Notonecta chinensis* | G | T | G | C | C | A | G | C | A | G | T | C | G | C | G | G | T | T | A |
| NC036671.1 | Heteroptera | *Notonecta chinensis* | G | T | G | C | C | A | G | C | A | G | T | C | G | C | G | G | T | T | A |
| MH780861.1 | Heteroptera | *Notonecta montandoni* | G | T | G | C | C | A | G | C | A | G | T | C | G | C | G | G | T | T | A |
| NC045886.1 | Heteroptera | *Notonecta montandoni* | G | T | G | C | C | A | G | C | A | G | T | C | G | C | G | G | T | T | A |
| KM244707.1 | Heteroptera | *Notonectidae* sp. | G | T | G | C | C | A | G | C | A | G | T | C | G | C | G | G | T | T | A |
| FJ456950.1 | Heteroptera | *Ochterus marginatus* | G | T | G | C | C | A | G | C | A | G | T | T | G | C | G | G | T | T | A |

| | | | | | | | | | | | | | | | | | | | | |
|---|---|---|---|---|---|---|---|---|---|---|---|---|---|---|---|---|---|---|---|---|
| JQ910988.1 | Heteroptera | *Perittopus* sp. | G | T | G | C | C | A | G | C | C | G | T | C | G | C | G | G | T | T | A |
| KP400583.1 | Heteroptera | *Ptilomera tigrina* | G | T | G | C | C | A | G | C | C | G | T | C | G | C | G | G | T | T | A |
| MN027271.1 | Heteroptera | *Rhagovelia reitteri* | G | T | G | C | C | A | G | C | C | G | T | C | G | C | G | G | T | T | A |
| NC044721.1 | Heteroptera | *Rhagovelia reitteri* | G | T | G | C | C | A | G | C | C | G | T | C | G | C | G | G | T | T | A |
| MN027279.1 | Heteroptera | *Rhagovelia* sp. | G | T | G | C | C | A | G | C | C | G | T | C | G | C | G | G | T | T | A |
| FJ456941.1 | Heteroptera | *Sigara septemlineata* | G | T | G | C | C | A | G | C | A | G | T | C | G | C | G | G | T | T | A |
| MN027277.1 | Heteroptera | *Trepobates* sp. | G | T | G | C | C | A | G | C | C | G | T | C | G | C | G | G | T | T | A |
| FJ456952.1 | Heteroptera | *Valentia hoffmanni* | G | T | G | C | C | A | G | C | A | G | C | C | G | C | G | G | T | T | A |
| NC012823.1 | Heteroptera | *Valentia hoffmanni* | G | T | G | C | C | A | G | C | A | G | C | C | G | C | G | G | T | T | A |
| KF840564.1 | Megaloptera | *Acanthacorydalis orientalis* | G | T | G | C | C | A | G | C | A | T | C | C | G | C | G | G | T | T | A |
| NC023462.1 | Megaloptera | *Acanthacorydalis orientalis* | G | T | G | C | C | A | G | C | A | T | C | C | G | C | G | G | T | T | A |
| KU925864.1 | Megaloptera | *Archichauliodes deceptor* | G | T | G | C | C | A | G | C | A | G | C | T | G | C | G | G | T | T | A |
| NC035504.1 | Megaloptera | *Archichauliodes deceptor* | G | T | G | C | C | A | G | C | A | G | C | T | G | C | G | G | T | T | A |
| KU302713.1 | Megaloptera | *Chloronia mirifica* | G | T | G | C | C | A | G | C | A | T | C | C | G | C | G | G | T | T | A |
| NC033349.1 | Megaloptera | *Chloronia mirifica* | G | T | G | C | C | A | G | C | A | T | C | C | G | C | G | G | T | T | A |
| FJ171323.1 | Megaloptera | *Corydalus cornutus* | G | T | G | C | C | A | G | C | A | T | C | C | G | C | G | G | T | T | A |
| NC011276.1 | Megaloptera | *Corydalus cornutus* | G | T | G | C | C | A | G | C | A | T | C | C | G | C | G | G | T | T | A |
| KJ806318.1 | Megaloptera | *Dysmicohermes ingens* | G | T | G | C | C | A | G | C | A | T | C | C | G | C | G | G | T | T | A |
| NC024657.1 | Megaloptera | *Dysmicohermes ingens* | G | T | G | C | C | A | G | C | A | T | C | C | G | C | G | G | T | T | A |
| JQ351950.1 | Megaloptera | *Neochauliodes bowringi* | G | T | G | C | C | A | G | C | A | T | C | C | G | C | G | G | T | T | A |

| | | | | | | | | | | | | | | | | | | | | | |
|---|---|---|---|---|---|---|---|---|---|---|---|---|---|---|---|---|---|---|---|---|---|
| NC023444.1 | Megaloptera | *Neochauliodes bowringi* | G | T | G | C | C | A | G | C | A | T | C | C | G | C | G | G | T | T | A |
| KF785799.1 | Megaloptera | *Neochauliodes fraternus* | G | T | G | C | C | A | G | C | A | T | C | C | G | C | G | G | T | T | A |
| NC025282.1 | Megaloptera | *Neochauliodes fraternus* | G | T | G | C | C | A | G | C | A | T | C | C | G | C | G | G | T | T | A |
| KX821680.1 | Megaloptera | *Neochauliodes parasparsus* | G | T | G | C | C | A | G | C | A | T | C | C | G | C | G | G | T | T | A |
| JX110703.1 | Megaloptera | *Neochauliodes punctatolosus* | G | T | G | C | C | A | G | C | A | T | C | C | G | C | G | G | T | T | A |
| NC018772.1 | Megaloptera | *Neochauliodes punctatolosus* | G | T | G | C | C | A | G | C | A | T | C | C | G | C | G | G | T | T | A |
| KF771894.1 | Megaloptera | *Neochauliodes rotundatus* | G | T | G | C | C | A | G | C | A | T | C | C | G | C | G | G | T | T | A |
| NC025281.1 | Megaloptera | *Neochauliodes rotundatus* | G | T | G | C | C | A | G | C | A | T | C | C | G | C | G | G | T | T | A |
| KP126231.1 | Megaloptera | *Neoneuromus tonkinensis* | G | T | G | C | C | A | G | C | A | T | C | C | G | C | G | G | T | T | A |
| NC027852.1 | Megaloptera | *Neoneuromus tonkinensis* | G | T | G | C | C | A | G | C | A | T | C | C | G | C | G | G | T | T | A |
| KP126232.1 | Megaloptera | *Nevromus exterior* | G | T | G | C | C | A | G | C | A | T | C | C | G | C | G | G | T | T | A |
| NC027851.1 | Megaloptera | *Nevromus exterior* | G | T | G | C | C | A | G | C | A | T | C | C | G | C | G | G | T | T | A |
| EU526394.1 | Megaloptera | *Protohermes concolorus* | G | T | G | C | C | A | G | C | A | T | C | C | G | C | G | G | T | T | A |
| NC011524.1 | Megaloptera | *Protohermes concolorus* | G | T | G | C | C | A | G | C | A | T | C | C | G | C | G | G | T | T | A |
| FJ859905.1 | Megaloptera | *Sialis hamata* | G | T | G | C | C | A | G | C | A | T | C | C | G | C | G | G | T | T | A |

| | | | | | | | | | | | | | | | | | | | | |
|---|---|---|---|---|---|---|---|---|---|---|---|---|---|---|---|---|---|---|---|---|
| NC013256.1 | Megaloptera | *Sialis hamata* | G | T | G | C | C | A | G | C | A | T | C | C | G | C | G | G | T | T | A |
| KT876911.1 | Megaloptera | *Sialis* sp. | G | T | G | C | C | A | G | C | A | T | C | C | G | C | G | G | T | T | A |
| KY587199.1 | Neuroptera | *Abachrysa eureka* | G | T | G | C | C | A | G | C | A | T | C | T | G | C | G | G | T | T | A |
| NC015095.1 | Neuroptera | *Apochrysa matsumurae* | G | T | G | C | C | A | G | C | A | T | C | T | G | C | G | G | T | T | A |
| KC758703.1 | Neuroptera | *Ascalohybris subjacens* | G | T | G | C | C | A | G | C | A | T | C | T | G | C | G | G | T | T | A |
| NC021428.1 | Neuroptera | *Ascalohybris subjacens* | G | T | G | C | C | A | G | C | A | T | C | T | G | C | G | G | T | T | A |
| FJ171324.1 | Neuroptera | *Ascaloptynx appendiculatus* | G | T | G | C | C | A | G | C | A | T | C | T | G | C | G | G | T | T | A |
| NC011277.1 | Neuroptera | *Ascaloptynx appendiculatus* | G | T | G | C | C | A | G | C | A | T | C | T | G | C | G | G | T | T | A |
| KT425083.1 | Neuroptera | *Balmes birmanus* | G | T | G | C | C | A | G | C | A | T | C | T | G | C | G | G | T | T | A |
| KX369241.1 | Neuroptera | *Bullanga florida* | G | T | G | C | C | A | G | C | A | T | T | T | G | C | G | G | T | T | A |
| NC032298.1 | Neuroptera | *Bullanga florida* | G | T | G | C | C | A | G | C | A | T | T | T | G | C | G | G | T | T | A |
| KT425069.1 | Neuroptera | *Chasmoptera huttii* | G | T | G | C | C | A | G | C | A | T | C | T | G | C | G | G | T | T | A |
| JX033119.1 | Neuroptera | *Chrysopa pallens* | G | T | G | C | C | A | G | C | A | T | C | T | G | C | G | G | T | T | A |
| NC019618.1 | Neuroptera | *Chrysopa pallens* | G | T | G | C | C | A | G | C | A | T | C | T | G | C | G | G | T | T | A |
| KU877169.1 | Neuroptera | *Chrysoperla externa* | G | T | G | C | C | A | G | C | A | T | C | T | G | C | G | G | T | T | A |
| NC030341.1 | Neuroptera | *Chrysoperla externa* | G | T | G | C | C | A | G | C | A | T | C | T | G | C | G | G | T | T | A |
| KP027407.1 | Neuroptera | *Chrysoperla nipponensis* | G | T | G | C | C | A | G | C | A | T | C | T | G | C | G | G | T | T | A |
| NC015093.1 | Neuroptera | *Chrysoperla nipponensis* | G | T | G | C | C | A | G | C | A | T | C | T | G | C | G | G | T | T | A |
| MK301246.1 | Neuroptera | *Dendroleon pantherinus* | G | T | G | C | C | A | G | C | A | T | T | C | G | C | G | G | T | T | A |
| KT425073.1 | Neuroptera | *Dilar* sp. | G | T | G | C | C | A | G | C | A | T | T | T | G | C | G | G | T | T | A |

| | | | | | | | | | | | | | | | | | | | | | |
|---|---|---|---|---|---|---|---|---|---|---|---|---|---|---|---|---|---|---|---|---|---|
| FJ859906.1 | Neuroptera | *Ditaxis biseriata* | G | T | G | C | C | A | G | C | A | T | C | T | G | C | G | G | T | T | A |
| NC013257.1 | Neuroptera | *Ditaxis biseriata* | G | T | G | C | C | A | G | C | A | T | C | T | G | C | G | G | T | T | A |
| KT425087.1 | Neuroptera | *Drepanepteryx phalaenoides* | G | T | G | C | C | A | G | C | A | T | C | T | G | C | G | G | T | T | A |
| KF701327.1 | Neuroptera | *Epacanthaclisis banksi* | G | T | G | C | C | A | G | C | A | T | C | T | G | C | G | G | T | T | A |
| NC025905.1 | Neuroptera | *Epacanthaclisis banksi* | G | T | G | C | C | A | G | C | A | T | C | T | G | C | G | G | T | T | A |
| MF972496.1 | Neuroptera | *Euclimacia badia* | G | T | G | C | C | A | G | C | A | T | C | T | G | C | G | G | T | T | A |
| NC039773.1 | Neuroptera | *Euclimacia badia* | G | T | G | C | C | A | G | C | A | T | C | T | G | C | G | G | T | T | A |
| MF972495.1 | Neuroptera | *Eumantispa harmandi* | G | T | G | C | C | A | G | C | A | T | C | T | G | C | G | G | T | T | A |
| NC039772.1 | Neuroptera | *Eumantispa harmandi* | G | T | G | C | C | A | G | C | A | T | C | T | G | C | G | G | T | T | A |
| KY364372.1 | Neuroptera | *Gatzara jezoensis* | G | T | G | C | C | A | G | C | A | T | T | C | G | C | G | G | T | T | A |
| MK341582.1 | Neuroptera | *Gryposmylus pennyi* | G | T | G | C | C | A | G | C | A | T | C | T | G | C | G | G | T | T | A |
| MN852445.1 | Neuroptera | *Hemerobius japonicus* | G | T | G | C | C | A | G | C | A | T | C | T | G | C | G | G | T | T | A |
| NC048517.1 | Neuroptera | *Hemerobius japonicus* | G | T | G | C | C | A | G | C | A | T | C | T | G | C | G | G | T | T | A |
| KY587200.1 | Neuroptera | *Italochrysa insignis* | G | T | G | C | C | A | G | C | A | T | C | T | G | C | G | G | T | T | A |
| NC015609.1 | Neuroptera | *Libelloides macaronius* | G | T | G | C | C | A | G | C | A | T | C | T | G | C | G | G | T | T | A |
| MF972494.1 | Neuroptera | *Mantispa japonica* | G | T | G | C | C | A | G | C | A | T | C | T | G | C | G | G | T | T | A |
| NC039771.1 | Neuroptera | *Mantispa japonica* | G | T | G | C | C | A | G | C | A | T | C | T | G | C | G | G | T | T | A |
| MK472073.1 | Neuroptera | *Micromus angulatus* | G | T | G | C | C | A | G | C | A | T | C | T | G | C | G | G | T | T | A |
| NC042677.1 | Neuroptera | *Micromus angulatus* | G | T | G | C | C | A | G | C | A | T | C | T | G | C | G | G | T | T | A |
| MT118664.1 | Neuroptera | *Myrmeleon formicarius* | G | T | G | C | C | A | G | C | A | T | C | T | G | C | G | G | T | T | A |
| NC047285.1 | Neuroptera | *Myrmeleon formicarius* | G | T | G | C | C | A | G | C | A | T | C | T | G | C | G | G | T | T | A |

| | | | | | | | | | | | | | | | | | | | | | |
|---|---|---|---|---|---|---|---|---|---|---|---|---|---|---|---|---|---|---|---|---|---|
| KJ461323.1 | Neuroptera | *Myrmeleon immanis* | G | T | G | C | C | A | G | C | A | T | C | T | G | C | G | G | T | T | A |
| NC024826.1 | Neuroptera | *Myrmeleon immanis* | G | T | G | C | C | A | G | C | A | T | C | T | G | C | G | G | T | T | A |
| KR078257.1 | Neuroptera | *Neuronema laminatum* | G | T | G | C | C | A | G | C | A | T | C | T | G | C | G | G | T | T | A |
| NC028153.1 | Neuroptera | *Neuronema laminatum* | G | T | G | C | C | A | G | C | A | T | C | T | G | C | G | G | T | T | A |
| KJ461322.1 | Neuroptera | *Nymphes myrmeleonoides* | G | T | G | C | C | A | G | C | A | T | C | T | G | C | G | G | T | T | A |
| NC024825.1 | Neuroptera | *Nymphes myrmeleonoides* | G | T | G | C | C | A | G | C | A | T | C | T | G | C | G | G | T | T | A |
| KT425090.1 | Neuroptera | *Oliarces clara* | G | T | G | C | C | A | G | C | A | T | C | T | G | C | G | G | T | T | A |
| MN818868.1 | Neuroptera | *Osmylus atomatus* | G | T | G | C | C | A | G | C | A | T | C | T | G | C | G | G | T | T | A |
| MN818867.1 | Neuroptera | *Osmylus fulvicephalus* | G | T | G | C | C | A | G | C | A | T | C | T | G | C | G | G | T | T | A |
| MN617021.1 | Neuroptera | *Osmylus lucalatus* | G | T | G | C | C | A | G | C | A | T | C | T | G | C | G | G | T | T | A |
| MN818870.1 | Neuroptera | *Osmylus* sp. | G | T | G | C | C | A | G | C | A | T | C | T | G | C | G | G | T | T | A |
| MN818869.1 | Neuroptera | *Osmylus zheanus* | G | T | G | C | C | A | G | C | A | T | C | T | G | C | G | G | T | T | A |
| FJ171325.1 | Neuroptera | *Polystoechotes punctatus* | G | T | G | C | C | A | G | C | A | T | C | T | G | C | G | G | T | T | A |
| NC011278.1 | Neuroptera | *Polystoechotes punctatus* | G | T | G | C | C | A | G | C | A | T | C | T | G | C | G | G | T | T | A |
| KF626446.1 | Neuroptera | *Rapisma xizangense* | G | T | G | C | C | A | G | C | A | T | T | T | G | C | G | G | T | T | A |
| NC023362.1 | Neuroptera | *Rapisma xizangense* | G | T | G | C | C | A | G | C | A | T | T | T | G | C | G | G | T | T | A |
| KF626447.1 | Neuroptera | *Rapisma zayuanum* | G | T | G | C | C | A | G | C | A | T | T | T | G | C | G | G | T | T | A |
| NC023363.1 | Neuroptera | *Rapisma zayuanum* | G | T | G | C | C | A | G | C | A | T | T | T | G | C | G | G | T | T | A |

| | | | | | | | | | | | | | | | | | | | | | |
|---|---|---|---|---|---|---|---|---|---|---|---|---|---|---|---|---|---|---|---|---|---|
| KT425067.1 | Neuroptera | *Semidalis aleyrodiformis* | G | T | G | C | C | A | G | C | A | T | C | T | G | C | G | G | T | T | A |
| MH361300.1 | Neuroptera | *Suhpalacsa longialata* | G | T | G | C | C | A | G | C | A | T | C | T | G | C | G | G | T | T | A |
| NC039948.1 | Neuroptera | *Suhpalacsa longialata* | G | T | G | C | C | A | G | C | A | T | C | T | G | C | G | G | T | T | A |
| MK301247.1 | Neuroptera | *Suhpalacsa* sp. | G | T | G | C | C | A | G | C | A | T | C | T | G | C | G | G | T | T | A |
| MK408756.1 | Neuroptera | *Thaumatosmylus hainanus* | G | T | G | C | C | A | G | C | A | T | C | T | G | C | G | G | T | T | A |
| NC046578.1 | Neuroptera | *Thaumatosmylus hainanus* | G | T | G | C | C | A | G | C | A | T | C | T | G | C | G | G | T | T | A |
| KC515397.1 | Neuroptera | *Thyridosmylus langii* | G | T | G | C | C | A | G | C | A | T | C | T | G | C | G | G | T | T | A |
| NC021415.1 | Neuroptera | *Thyridosmylus langii* | G | T | G | C | C | A | G | C | A | T | C | T | G | C | G | G | T | T | A |
| MH020192.1 | Coleoptera | *Abscondita anceyi* | G | T | G | C | C | A | G | C | A | A | T | T | G | C | G | G | T | T | A |
| NC039706.1 | Coleoptera | *Abscondita anceyi* | G | T | G | C | C | A | G | C | A | A | T | T | G | C | G | G | T | T | A |
| MK122952.1 | Coleoptera | *Abscondita chinensis* | G | T | G | C | C | A | G | C | A | A | T | T | G | C | G | G | T | T | A |
| MK292092.1 | Coleoptera | *Abscondita terminalis* | G | T | G | C | C | A | G | C | A | A | T | T | G | C | G | G | T | T | A |
| NC044776.1 | Coleoptera | *Abscondita terminalis* | G | T | G | C | C | A | G | C | A | A | T | T | G | C | G | G | T | T | A |
| KT876878.1 | Coleoptera | *Acilius sulcatus* | G | T | G | C | C | A | G | C | A | G | T | T | G | C | G | G | T | T | A |
| KX758085.1 | Coleoptera | *Aquatica ficta* | G | T | G | C | C | A | G | C | A | A | T | T | G | C | G | G | T | T | A |
| NC035060.1 | Coleoptera | *Aquatica ficta* | G | T | G | C | C | A | G | C | A | A | T | T | G | C | G | G | T | T | A |
| NC035755.1 | Coleoptera | *Aquatica lateralis* | G | T | G | C | C | A | G | C | A | A | T | T | G | C | G | G | T | T | A |
| KF667531.1 | Coleoptera | *Aquatica leii* | G | T | G | C | C | A | G | C | A | A | T | T | G | C | G | G | T | T | A |
| NC025276.1 | Coleoptera | *Aquatica leii* | G | T | G | C | C | A | G | C | A | A | T | T | G | C | G | G | T | T | A |
| KX758086.1 | Coleoptera | *Aquatica wuhana* | G | T | G | C | C | A | G | C | A | A | T | T | G | C | G | G | T | T | A |

| | | | | | | | | | | | | | | | | | | | | | |
|---|---|---|---|---|---|---|---|---|---|---|---|---|---|---|---|---|---|---|---|---|---|
| NC035061.1 | Coleoptera | *Aquatica wuhana* | G | T | G | C | C | A | G | C | A | A | T | T | G | C | G | G | T | T | A |
| KX229747.1 | Coleoptera | *Asymmetricata circumdata* | G | T | G | C | C | A | G | C | A | A | T | T | G | C | G | G | T | T | A |
| MK292113.1 | Coleoptera | *Asymmetricata circumdata* | G | T | G | C | C | A | G | C | A | A | T | T | G | C | G | G | T | T | A |
| NC032062.1 | Coleoptera | *Asymmetricata circumdata* | G | T | G | C | C | A | G | C | A | A | T | T | G | C | G | G | T | T | A |
| KJ922151.1 | Coleoptera | *Bicellonychia lividipennis* | G | T | G | C | C | A | G | C | A | A | T | T | G | C | G | G | T | T | A |
| NC030060.1 | Coleoptera | *Bicellonychia lividipennis* | G | T | G | C | C | A | G | C | A | A | T | T | G | C | G | G | T | T | A |
| KT876885.1 | Coleoptera | *Colymbetes fuscus* | G | T | G | C | C | A | G | C | A | G | T | T | G | C | G | G | T | T | A |
| MK292114.1 | Coleoptera | *Curtos bilineatus* | G | T | G | C | C | A | G | C | A | A | T | T | G | C | G | G | T | T | A |
| NC044789.1 | Coleoptera | *Curtos bilineatus* | G | T | G | C | C | A | G | C | A | A | T | T | G | C | G | G | T | T | A |
| MK609965.1 | Coleoptera | *Curtos costipennis* | G | T | G | C | C | A | G | C | A | A | T | T | G | C | G | G | T | T | A |
| MH651351.1 | Coleoptera | *Diaphanes citrinus* | G | T | G | C | C | A | G | C | A | A | T | T | G | C | G | G | T | T | A |
| MK292116.1 | Coleoptera | *Diaphanes mendax* | G | T | G | C | C | A | G | C | A | A | T | T | G | C | G | G | T | T | A |
| NC044791.1 | Coleoptera | *Diaphanes mendax* | G | T | G | C | C | A | G | C | A | A | T | T | G | C | G | G | T | T | A |
| MK292094.1 | Coleoptera | *Diaphanes nubilus* | G | T | G | C | C | A | G | C | A | A | T | T | G | C | G | G | T | T | A |
| NC044787.1 | Coleoptera | *Diaphanes nubilus* | G | T | G | C | C | A | G | C | A | A | T | T | G | C | G | G | T | T | A |
| MK292118.1 | Coleoptera | *Diaphanes pectinealis* | G | T | G | C | C | A | G | C | A | A | T | T | G | C | G | G | T | T | A |
| NC044793.1 | Coleoptera | *Diaphanes pectinealis* | G | T | G | C | C | A | G | C | A | A | T | T | G | C | G | G | T | T | A |
| MK292095.1 | Coleoptera | *Diaphanes* sp. | G | T | G | C | C | A | G | C | A | A | T | T | G | C | G | G | T | T | A |
| MG242622.1 | Coleoptera | *Ellychnia corrusca* | G | T | G | C | C | A | G | C | A | A | T | T | G | C | G | G | T | T | A |

| | | | | | | | | | | | | | | | | | | | | | |
|---|---|---|---|---|---|---|---|---|---|---|---|---|---|---|---|---|---|---|---|---|---|
| MK292112.1 | Coleoptera | *Emeia pseudosauteri* | G | T | G | C | C | A | G | C | A | A | T | T | G | C | G | G | T | T | A |
| NC044778.1 | Coleoptera | *Emeia pseudosauteri* | G | T | G | C | C | A | G | C | A | A | T | T | G | C | G | G | T | T | A |
| KT876897.1 | Coleoptera | *Hydroporus erythrocephalus* | G | T | G | C | C | A | G | C | A | G | T | T | G | C | G | G | T | T | A |
| KT876896.1 | Coleoptera | *Hydroporus obscurus* | G | T | G | C | C | A | G | C | A | G | T | T | G | C | G | G | T | T | A |
| KT876901.1 | Coleoptera | *Hygrotus confluens* | G | T | G | C | C | A | G | C | A | G | T | T | G | C | G | G | T | T | A |
| KT876900.1 | Coleoptera | *Hygrotus impressopunctatus* | G | T | G | C | C | A | G | C | A | G | T | T | G | C | G | G | T | T | A |
| KT876899.1 | Coleoptera | *Hygrotus inaequalis* | G | T | G | C | C | A | G | C | A | G | T | T | G | C | G | G | T | T | A |
| KX087299.1 | Coleoptera | *Hygrotus nigrolineatus* | G | T | G | C | C | A | G | C | A | G | T | T | G | C | G | G | T | T | A |
| MH427718.1 | Coleoptera | *Inflata indica* | G | T | G | C | C | A | G | C | A | A | T | T | G | C | G | G | T | T | A |
| NC039700.1 | Coleoptera | *Inflata indica* | G | T | G | C | C | A | G | C | A | A | T | T | G | C | G | G | T | T | A |
| MK292091.1 | Coleoptera | *Lamprigera yunnana* | G | T | G | C | C | A | G | C | A | A | T | T | G | C | G | G | T | T | A |
| NC044786.1 | Coleoptera | *Lamprigera yunnana* | G | T | G | C | C | A | G | C | A | A | T | T | G | C | G | G | T | T | A |
| KX087302.1 | Coleoptera | *Lampyris noctiluca* | G | T | G | C | C | A | G | C | A | A | T | T | G | C | G | G | T | T | A |
| MG912994.1 | Coleoptera | *Limbodessus palmulaoides* | G | T | G | C | C | A | G | C | A | G | T | T | G | C | G | G | T | T | A |
| NC037749.1 | Coleoptera | *Limbodessus palmulaoides* | G | T | G | C | C | A | G | C | A | G | T | T | G | C | G | G | T | T | A |
| KT876902.1 | Coleoptera | *Liopterus haemorrhoidalis* | G | T | G | C | C | A | G | C | A | G | T | A | G | C | G | G | T | T | A |
| NC022472.1 | Coleoptera | *Luciola cruciata* | G | T | G | C | C | A | G | C | A | A | T | T | G | C | G | G | T | T | A |

| | | | | | | | | | | | | | | | | | | | | |
|---|---|---|---|---|---|---|---|---|---|---|---|---|---|---|---|---|---|---|---|---|
| MG770613.1 | Coleoptera | *Luciola curtithorax* | G | T | G | C | C | A | G | C | A | A | T | T | G | C | G | G | T | T | A |
| NC038225.1 | Coleoptera | *Luciola curtithorax* | G | T | G | C | C | A | G | C | A | A | T | T | G | C | G | G | T | T | A |
| KP313820.1 | Coleoptera | *Luciola substriata* | G | T | G | C | C | A | G | C | A | A | T | T | G | C | G | G | T | T | A |
| NC027176.1 | Coleoptera | *Luciola substriata* | G | T | G | C | C | A | G | C | A | A | T | T | G | C | G | G | T | T | A |
| FJ859901.1 | Coleoptera | *Macrogyrus oblongus* | G | T | G | C | C | A | G | C | A | G | T | T | G | C | G | G | T | T | A |
| NC013249.1 | Coleoptera | *Macrogyrus oblongus* | G | T | G | C | C | A | G | C | A | G | T | T | G | C | G | G | T | T | A |
| MG912995.1 | Coleoptera | *Paroster macrosturtensis* | G | T | G | C | C | A | G | C | A | G | T | T | G | C | G | G | T | T | A |
| NC037750.1 | Coleoptera | *Paroster macrosturtensis* | G | T | G | C | C | A | G | C | A | G | T | T | G | C | G | G | T | T | A |
| MG912996.1 | Coleoptera | *Paroster mesosturtensis* | G | T | G | C | C | A | G | C | A | G | T | T | G | C | G | G | T | T | A |
| NC037751.1 | Coleoptera | *Paroster mesosturtensis* | G | T | G | C | C | A | G | C | A | G | T | T | G | C | G | G | T | T | A |
| MG912997.1 | Coleoptera | *Paroster microsturtensis* | G | T | G | C | C | A | G | C | A | G | T | T | G | C | G | G | T | T | A |
| NC037752.1 | Coleoptera | *Paroster microsturtensis* | G | T | G | C | C | A | G | C | A | G | T | T | G | C | G | G | T | T | A |
| KY778696.1 | Coleoptera | *Photinus pyralis* | G | T | G | C | C | A | G | C | A | A | T | T | G | C | G | G | T | T | A |
| MF686051.1 | Coleoptera | *Pteroptyx maipo* | G | T | G | C | C | A | G | C | A | A | T | T | G | C | G | G | T | T | A |
| NC036353.1 | Coleoptera | *Pteroptyx maipo* | G | T | G | C | C | A | G | C | A | A | T | T | G | C | G | G | T | T | A |
| MK292115.1 | Coleoptera | *Pyrocoelia praetexta* | G | T | G | C | C | A | G | C | A | A | T | T | G | C | G | G | T | T | A |
| NC044790.1 | Coleoptera | *Pyrocoelia praetexta* | G | T | G | C | C | A | G | C | A | A | T | T | G | C | G | G | T | T | A |
| AF452048.1 | Coleoptera | *Pyrocoelia rufa* | G | T | G | C | C | A | G | C | A | A | T | T | G | C | G | G | T | T | A |
| NC003970.1 | Coleoptera | *Pyrocoelia rufa* | G | T | G | C | C | A | G | C | A | A | T | T | G | C | G | G | T | T | A |
| MK292117.1 | Coleoptera | *Pyrocoelia thibetana* | G | T | G | C | C | A | G | C | A | A | T | T | G | C | G | G | T | T | A |
| NC044792.1 | Coleoptera | *Pyrocoelia thibetana* | G | T | G | C | C | A | G | C | A | A | T | T | G | C | G | G | T | T | A |
| MK292111.1 | Coleoptera | *Vesta saturnalis* | G | T | G | C | C | A | G | C | A | A | T | T | G | C | G | G | T | T | A |

| | | | | | | | | | | | | | | | | | | | | |
|---|---|---|---|---|---|---|---|---|---|---|---|---|---|---|---|---|---|---|---|---|---|
| NC044788.1 | Coleoptera | *Vesta saturnalis* | G | T | G | C | C | A | G | C | A | A | T | T | G | C | G | G | T | T | A |
| MT584157.1 | Trichoptera | *Lype reducta* | G | T | G | C | C | A | G | C | A | G | T | C | G | C | G | G | T | A | A |
| MT410854.1 | Trichoptera | *Molanna angustata* | G | T | G | C | C | A | G | C | A | A | T | T | G | C | G | G | T | T | A |
| | | | 0.0 | 0.0 | 0.0 | 0.0 | 0.0 | 100.0 | 0.0 | 0.0 | 54.4 | 28.7 | 0.0 | 0.2 | 0.0 | 0.0 | 0.0 | 0.0 | 0.0 | 0.4 | 100.0 |
| | | | 0.0 | 100.0 | 0.0 | 0.0 | 0.0 | 0.0 | 0.0 | 0.0 | 10.0 | 23.0 | 29.1 | 30.3 | 0.0 | 0.0 | 0.0 | 0.0 | 100.0 | 99.2 | 0.0 |
| | | | 100.0 | 0.0 | 100.0 | 0.0 | 0.0 | 0.0 | 100.0 | 0.0 | 0.4 | 48.3 | 0.0 | 0.0 | 100.0 | 0.0 | 100.0 | 100.0 | 0.0 | 0.4 | 0.0 |
| | | | 0.0 | 0.0 | 0.0 | 100.0 | 100.0 | 0.0 | 0.0 | 100.0 | 35.2 | 0.0 | 70.9 | 69.5 | 0.0 | 100.0 | 0.0 | 0.0 | 0.0 | 0.0 | 0.0 |

Table S7. Nucleotide sequences of universal primer (MtInsects-12S-Hem_R) and homologous region of 265 sequences of insects used for development in this study.

| | | | | | | | | | | | | | | | | | | | | | | | |
|---|---|---|---|---|---|---|---|---|---|---|---|---|---|---|---|---|---|---|---|---|---|---|---|
| MtInsects-12S-Hem_R | | | | H | A | T | A | R | D | R | G | G | G | T | M | T | C | T | A | A | T | C | C |
| | | | A | 218 | 261 | 0 | 265 | 235 | 52 | 253 | 0 | 0 | 0 | 0 | 253 | 0 | 0 | 0 | 265 | 265 | 0 | 0 | 0 |
| | | | T | 37 | 2 | 265 | 0 | 0 | 204 | 0 | 0 | 0 | 0 | 265 | 0 | 265 | 0 | 265 | 0 | 0 | 265 | 0 | 0 |
| | | | G | 3 | 2 | 0 | 0 | 30 | 7 | 12 | 265 | 265 | 265 | 0 | 0 | 0 | 0 | 0 | 0 | 0 | 0 | 0 | 0 |
| | | | C | 7 | 0 | 0 | 0 | 0 | 2 | 0 | 0 | 0 | 0 | 0 | 12 | 0 | 265 | 0 | 0 | 0 | 0 | 265 | 265 |
| GenBank Acc. No. | Order | Species | | | | | | | | | | | | | | | | | | | | | |
| GQ502451.1 | Ephemoroptera | *Caenis* sp. | | A | A | T | A | A | T | A | G | G | G | T | A | T | C | T | A | A | T | C | C |
| MG910499.1 | Ephemoroptera | *Caenis* sp. | | A | A | T | A | A | T | A | G | G | G | T | A | T | C | T | A | A | T | C | C |
| MN356096.1 | Ephemoroptera | *Caenis* sp. | | T | A | T | A | A | T | A | G | G | G | T | A | T | C | T | A | A | T | C | C |
| MN807287.1 | Ephemoroptera | *Choroterpides apiculata* | | T | A | T | A | A | T | A | G | G | G | T | A | T | C | T | A | A | T | C | C |
| MG870104.1 | Ephemoroptera | *Epeorus herklotsi* | | A | A | T | A | A | T | G | G | G | G | T | A | T | C | T | A | A | T | C | C |
| MH752075.1 | Ephemoroptera | *Epeorus herklotsi* | | A | A | T | A | A | T | G | G | G | G | T | A | T | C | T | A | A | T | C | C |
| NC039612.1 | Ephemoroptera | *Epeorus herklotsi* | | A | A | T | A | A | T | G | G | G | G | T | A | T | C | T | A | A | T | C | C |
| KJ493406.1 | Ephemoroptera | *Epeorus* sp. | | A | A | T | A | G | T | G | G | G | G | T | A | T | C | T | A | A | T | C | C |
| KM244708.1 | Ephemoroptera | *Epeorus* sp. | | A | A | T | A | G | T | G | G | G | G | T | A | T | C | T | A | A | T | C | C |
| MK951674.1 | Ephemoroptera | *Epeorus* sp. | | A | A | T | A | A | T | A | G | G | G | T | A | T | C | T | A | A | T | C | C |
| EU591678.1 | Ephemoroptera | *Ephemera orientalis* | | T | A | T | A | G | T | A | G | G | G | T | A | T | C | T | A | A | T | C | C |
| NC012645.1 | Ephemoroptera | *Ephemera* | | T | A | T | A | G | T | A | G | G | G | T | A | T | C | T | A | A | T | C | C |

| | | | | | | | | | | | | | | | | | | | | | | |
|---|---|---|---|---|---|---|---|---|---|---|---|---|---|---|---|---|---|---|---|---|---|---|
| | | *orientalis* | | | | | | | | | | | | | | | | | | | | |
| MK951659.1 | Ephemoroptera | *Ephemera* sp. | A | A | T | A | A | T | A | G | G | G | T | A | T | C | T | A | A | T | C | C |
| MT274127.1 | Ephemoroptera | *Ephemerella* sp. | C | A | T | A | G | T | A | G | G | G | T | C | T | C | T | A | A | T | C | C |
| NC050279.1 | Ephemoroptera | *Ephemerella* sp. | C | A | T | A | G | T | A | G | G | G | T | C | T | C | T | A | A | T | C | C |
| GU936203.1 | Ephemoroptera | *Habrophlebiodes zijinensis* | T | A | T | A | A | T | A | G | G | G | T | A | T | C | T | A | A | T | C | C |
| HM143892.1 | Ephemoroptera | *Isonychia ignota* | A | A | T | A | A | T | G | G | G | G | T | A | T | C | T | A | A | T | C | C |
| MH119135.1 | Ephemoroptera | *Isonychia kiangsinensis* | A | A | T | A | A | T | G | G | G | G | T | A | T | C | T | A | A | T | C | C |
| NC042163.1 | Ephemoroptera | *Isonychia kiangsinensis* | A | A | T | A | A | T | G | G | G | G | T | A | T | C | T | A | A | T | C | C |
| MK951658.1 | Ephemoroptera | *Isonychia* sp. | A | A | T | A | A | T | G | G | G | G | T | A | T | C | T | A | A | T | C | C |
| HM004123.1 | Ephemoroptera | *Paegniodes cupulatus* | A | A | T | A | G | T | A | G | G | G | T | A | T | C | T | A | A | T | C | C |
| MK951673.1 | Ephemoroptera | *Parafronurus sp.* | G | A | T | A | A | T | A | G | G | G | T | A | T | C | T | A | A | T | C | C |
| EU349015.1 | Ephemoroptera | *Parafronurus youi* | A | A | T | A | A | T | A | G | G | G | T | A | T | C | T | A | A | T | C | C |
| NC011359.1 | Ephemoroptera | *Parafronurus youi* | A | A | T | A | A | T | A | G | G | G | T | A | T | C | T | A | A | T | C | C |
| KM244674.1 | Ephemoroptera | *Potamanthus* sp. | A | A | T | A | A | T | A | G | G | G | T | A | T | C | T | A | A | T | C | C |
| MK951670.1 | Ephemoroptera | *Rhithrogena* sp. | A | A | T | A | A | T | A | G | G | G | T | A | T | C | T | A | A | T | C | C |
| MT274128.1 | Ephemoroptera | *Serratella* sp. | C | A | T | A | G | T | A | G | G | G | T | C | T | C | T | A | A | T | C | C |
| MT274129.1 | Ephemoroptera | *Serratella* sp. | T | T | T | A | G | T | A | G | G | G | T | C | T | C | T | A | A | T | C | C |
| NC050280.1 | Ephemoroptera | *Serratella* sp. | C | A | T | A | G | T | A | G | G | G | T | C | T | C | T | A | A | T | C | C |

| | | | | | | | | | | | | | | | | | | | | | | |
|---|---|---|---|---|---|---|---|---|---|---|---|---|---|---|---|---|---|---|---|---|---|---|
| NC050281.1 | Ephemoroptera | *Serratella* sp. | T | T | T | A | G | T | A | G | G | G | T | C | T | C | T | A | A | T | C | C |
| MT274130.1 | Ephemoroptera | *Serratella zapekinae* | A | A | T | A | G | T | A | G | G | G | T | C | T | C | T | A | A | T | C | C |
| NC050282.1 | Ephemoroptera | *Serratella zapekinae* | A | A | T | A | G | T | A | G | G | G | T | C | T | C | T | A | A | T | C | C |
| FJ606783.1 | Ephemoroptera | *Siphlonurus immanis* | A | A | T | A | G | T | A | G | G | G | T | A | T | C | T | A | A | T | C | C |
| NC013822.1 | Ephemoroptera | *Siphlonurus immanis* | A | A | T | A | G | T | A | G | G | G | T | A | T | C | T | A | A | T | C | C |
| KM244684.1 | Ephemoroptera | *Siphlonurus* sp. | A | A | T | A | G | T | A | G | G | G | T | A | T | C | T | A | A | T | C | C |
| HQ875717.1 | Ephemoroptera | *Siphluriscus chinensis* | A | A | T | A | A | T | G | G | G | G | T | A | T | C | T | A | A | T | C | C |
| MT274131.1 | Ephemoroptera | *Torleya grandiforceps* | G | A | T | A | G | T | A | G | G | G | T | C | T | C | T | A | A | T | C | C |
| NC050283.1 | Ephemoroptera | *Torleya grandiforceps* | G | A | T | A | G | T | A | G | G | G | T | C | T | C | T | A | A | T | C | C |
| MT274132.1 | Ephemoroptera | *Torleya nepalica* | A | G | T | A | G | T | A | G | G | G | T | C | T | C | T | A | A | T | C | C |
| NC050284.1 | Ephemoroptera | *Torleya nepalica* | A | G | T | A | G | T | A | G | G | G | T | C | T | C | T | A | A | T | C | C |
| HM067837.1 | Ephemoroptera | *Vietnamella dabieshanensis* | A | A | T | A | A | T | A | G | G | G | T | A | T | C | T | A | A | T | C | C |
| KM244655.1 | Ephemoroptera | *Vietnamella* sp. | A | A | T | A | G | T | A | G | G | G | T | A | T | C | T | A | A | T | C | C |
| MH751435.1 | Odonata | *Acisoma* | A | A | T | A | G | A | A | G | G | G | T | A | T | C | T | A | A | T | C | C |

| | | | | | | | | | | | | | | | | | | | | | | |
|---|---|---|---|---|---|---|---|---|---|---|---|---|---|---|---|---|---|---|---|---|---|---|
| | | *panorpoides* | | | | | | | | | | | | | | | | | | | | |
| NC042689.1 | Odonata | *Acisoma panorpoides* | A | A | T | A | G | A | A | G | G | G | T | A | T | C | T | A | A | T | C | C |
| KX161841.1 | Odonata | *Anax imperator* | A | A | T | A | A | A | A | G | G | G | T | A | T | C | T | A | A | T | C | C |
| NC031821.1 | Odonata | *Anax imperator* | A | A | T | A | A | A | A | G | G | G | T | A | T | C | T | A | A | T | C | C |
| KP233805.1 | Odonata | *Atrocalopteryx atrata* | T | A | T | A | A | A | A | G | G | G | T | A | T | C | T | A | A | T | C | C |
| NC027181.1 | Odonata | *Atrocalopteryx atrata* | T | A | T | A | A | A | A | G | G | G | T | A | T | C | T | A | A | T | C | C |
| KM658172.1 | Odonata | *Brachythemis contaminata* | A | A | T | A | A | A | A | G | G | G | T | A | T | C | T | A | A | T | C | C |
| NC026305.1 | Odonata | *Brachythemis contaminata* | A | A | T | A | A | A | A | G | G | G | T | A | T | C | T | A | A | T | C | C |
| EU591677.1 | Odonata | *Davidius lunatus* | A | A | T | A | A | A | A | G | G | G | T | A | T | C | T | A | A | T | C | C |
| NC012644.1 | Odonata | *Davidius lunatus* | A | A | T | A | A | A | A | G | G | G | T | A | T | C | T | A | A | T | C | C |
| MH751436.1 | Odonata | *Deielia phaon* | A | A | T | A | A | A | A | G | G | G | T | A | T | C | T | A | A | T | C | C |
| NC042690.1 | Odonata | *Deielia phaon* | A | A | T | A | A | A | A | G | G | G | T | A | T | C | T | A | A | T | C | C |
| MF716899.1 | Odonata | *Enallagma cyathigerum* | T | A | T | A | A | A | A | G | G | G | T | A | T | C | T | A | A | T | C | C |
| JX050223.1 | Odonata | *Epiophlebia superstes* | A | A | T | A | A | A | A | G | G | G | T | A | T | C | T | A | A | T | C | C |
| NC023232.1 | Odonata | *Epiophlebia* | A | A | T | A | A | A | A | G | G | G | T | A | T | C | T | A | A | T | C | C |

| | | | | | | | | | | | | | | | | | | | | | | |
|---|---|---|---|---|---|---|---|---|---|---|---|---|---|---|---|---|---|---|---|---|---|---|
| | | *superstes* | | | | | | | | | | | | | | | | | | | | |
| MK522522.1 | Odonata | *Epophthalmia elegans* | A | A | T | A | A | A | A | G | G | G | T | A | T | C | T | A | A | T | C | C |
| NC046756.1 | Odonata | *Epophthalmia elegans* | A | A | T | A | A | A | A | G | G | G | T | A | T | C | T | A | A | T | C | C |
| HM126547.1 | Odonata | *Euphaea formosa* | T | A | T | A | A | A | A | G | G | G | T | A | T | C | T | A | A | T | C | C |
| NC014493.1 | Odonata | *Euphaea formosa* | T | A | T | A | A | A | A | G | G | G | T | A | T | C | T | A | A | T | C | C |
| KM244659.1 | Odonata | *Hydrobasileus croceus* | A | A | T | A | A | A | A | G | G | G | T | A | T | C | T | A | A | T | C | C |
| NC025758.1 | Odonata | *Hydrobasileus croceus* | A | A | T | A | A | A | A | G | G | G | T | A | T | C | T | A | A | T | C | C |
| KU958378.1 | Odonata | *Ischnura elegans* | T | A | T | A | A | A | A | G | G | G | T | A | T | C | T | A | A | T | C | C |
| MK951668.1 | Odonata | *Ischnura elegans* | T | A | T | A | A | A | A | G | G | G | T | A | T | C | T | A | A | T | C | C |
| NC031824.1 | Odonata | *Ischnura elegans* | T | A | T | A | A | A | A | G | G | G | T | A | T | C | T | A | A | T | C | C |
| KC878732.1 | Odonata | *Ischnura pumilio* | T | A | T | A | A | A | A | G | G | G | T | A | T | C | T | A | A | T | C | C |
| NC021617.1 | Odonata | *Ischnura pumilio* | T | A | T | A | A | A | A | G | G | G | T | A | T | C | T | A | A | T | C | C |
| MF990748.1 | Odonata | *Macromia daimoji* | A | A | T | A | A | A | A | G | G | G | T | A | T | C | T | A | A | T | C | C |
| NC041425.1 | Odonata | *Macromia daimoji* | A | A | T | A | A | A | A | G | G | G | T | A | T | C | T | A | A | T | C | C |
| MK722304.1 | Odonata | *Matrona basilaris* | T | A | T | A | A | G | A | G | G | G | T | A | T | C | T | A | A | T | C | C |
| MK722305.1 | Odonata | *Matrona basilaris* | T | A | T | A | A | G | A | G | G | G | T | A | T | C | T | A | A | T | C | C |
| KU958377.1 | Odonata | *Megaloprepus caerulatus* | T | A | T | A | A | A | A | G | G | G | T | A | T | C | T | A | A | T | C | C |

| | | | | | | | | | | | | | | | | | | | | | | |
|---|---|---|---|---|---|---|---|---|---|---|---|---|---|---|---|---|---|---|---|---|---|---|
| NC031823.1 | Odonata | *Megaloprepus caerulatus* | T | A | T | A | A | A | A | G | G | G | T | A | T | C | T | A | A | T | C | C |
| KU871065.1 | Odonata | *Mnais costalis* | T | A | T | A | A | G | A | G | G | G | T | A | T | C | T | A | A | T | C | C |
| KY402222.1 | Odonata | *Nannophya pygmaea* | A | A | T | A | A | A | A | G | G | G | T | A | T | C | T | A | A | T | C | C |
| NC039410.1 | Odonata | *Nannophya pygmaea* | A | A | T | A | A | A | A | G | G | G | T | A | T | C | T | A | A | T | C | C |
| KU361233.1 | Odonata | *Orthetrum chrysis* | A | A | T | A | A | A | A | G | G | G | T | A | T | C | T | A | A | T | C | C |
| NC032048.1 | Odonata | *Orthetrum chrysis* | A | A | T | A | A | A | A | G | G | G | T | A | T | C | T | A | A | T | C | C |
| KU361232.1 | Odonata | *Orthetrum glaucum* | A | A | T | A | A | A | A | G | G | G | T | A | T | C | T | A | A | T | C | C |
| NC032047.1 | Odonata | *Orthetrum glaucum* | A | A | T | A | A | A | A | G | G | G | T | A | T | C | T | A | A | T | C | C |
| MH751437.1 | Odonata | *Orthetrum melania* | A | A | T | A | A | A | A | G | G | G | T | A | T | C | T | A | A | T | C | C |
| NC042732.1 | Odonata | *Orthetrum melania* | A | A | T | A | A | A | A | G | G | G | T | A | T | C | T | A | A | T | C | C |
| KU361234.1 | Odonata | *Orthetrum sabina* | A | A | T | A | A | A | A | G | G | G | T | A | T | C | T | A | A | T | C | C |
| MH751438.1 | Odonata | *Orthetrum sabina* | A | A | T | A | A | A | A | G | G | G | T | A | T | C | T | A | A | T | C | C |
| NC032049.1 | Odonata | *Orthetrum sabina* | A | A | T | A | A | A | A | G | G | G | T | A | T | C | T | A | A | T | C | C |
| KU361235.1 | Odonata | *Orthetrum testaceum* | A | A | T | A | A | A | A | G | G | G | T | A | T | C | T | A | A | T | C | C |

| | | | | | | | | | | | | | | | | | | | | | | |
|---|---|---|---|---|---|---|---|---|---|---|---|---|---|---|---|---|---|---|---|---|---|---|
| MH751439.1 | Odonata | *Orthetrum testaceum* | A | A | T | A | A | T | A | G | G | G | T | A | T | C | T | A | A | T | C | C |
| NC032050.1 | Odonata | *Orthetrum testaceum* | A | A | T | A | A | A | A | G | G | G | T | A | T | C | T | A | A | T | C | C |
| AB126005.1 | Odonata | *Orthetrum triangulare melania* | A | A | T | A | A | A | A | G | G | G | T | A | T | C | T | A | A | T | C | C |
| MK951669.1 | Odonata | *Paracercion v-nigrum* | A | A | T | A | A | A | A | G | G | G | T | A | T | C | T | A | A | T | C | C |
| KP233804.1 | Odonata | *Platycnemis foliacea* | A | A | T | A | A | A | A | G | G | G | T | A | T | C | T | A | A | T | C | C |
| NC027180.1 | Odonata | *Platycnemis foliacea* | A | A | T | A | A | A | A | G | G | G | T | A | T | C | T | A | A | T | C | C |
| FJ606784.1 | Odonata | *Pseudolestes mirabilis* | A | A | T | A | A | A | A | G | G | G | T | A | T | C | T | A | A | T | C | C |
| NC020636.1 | Odonata | *Pseudolestes mirabilis* | A | A | T | A | A | A | A | G | G | G | T | A | T | C | T | A | A | T | C | C |
| MF150044.1 | Odonata | *Psolodesmus mandarinus* | T | A | T | A | A | G | A | G | G | G | T | A | T | C | T | A | A | T | C | C |
| NC042442.1 | Odonata | *Psolodesmus mandarinus* | T | A | T | A | A | G | A | G | G | G | T | A | T | C | T | A | A | T | C | C |
| MG594801.1 | Odonata | *Somatochlora* | A | A | T | A | A | A | A | G | G | G | T | A | T | C | T | A | A | T | C | C |

| | | | | | | | | | | | | | | | | | | | | | | |
|---|---|---|---|---|---|---|---|---|---|---|---|---|---|---|---|---|---|---|---|---|---|---|
| | | *hineana* | | | | | | | | | | | | | | | | | | | | |
| MH751440.1 | Odonata | *Tramea virginia* | A | A | T | A | A | T | A | G | G | G | T | A | T | C | T | A | A | T | C | C |
| NC042691.1 | Odonata | *Tramea virginia* | A | A | T | A | A | T | A | G | G | G | T | A | T | C | T | A | A | T | C | C |
| MH751441.1 | Odonata | *Trigomphus carus* | A | A | T | A | A | A | A | G | G | G | T | A | T | C | T | A | A | T | C | C |
| NC042733.1 | Odonata | *Trigomphus carus* | A | A | T | A | A | A | A | G | G | G | T | A | T | C | T | A | A | T | C | C |
| JX050224.1 | Odonata | *Vestalis melania* | T | A | T | A | G | G | G | G | G | G | T | A | T | C | T | A | A | T | C | C |
| NC023233.1 | Odonata | *Vestalis melania* | T | A | T | A | G | G | G | G | G | G | T | A | T | C | T | A | A | T | C | C |
| KM199685.1 | Plecoptera | *Acroneuria hainana* | A | A | T | A | G | T | A | G | G | G | T | A | T | C | T | A | A | T | C | C |
| NC026104.1 | Plecoptera | *Acroneuria hainana* | A | A | T | A | G | T | A | G | G | G | T | A | T | C | T | A | A | T | C | C |
| KX091846.1 | Plecoptera | *Agnetina aequalis* | A | A | T | A | A | T | A | G | G | G | T | A | T | C | T | A | A | T | C | C |
| MH085446.1 | Plecoptera | *Amphinemura longispina* | A | A | T | A | A | T | A | G | G | G | T | A | T | C | T | A | A | T | C | C |
| NC044748.1 | Plecoptera | *Amphinemura longispina* | A | A | T | A | A | T | A | G | G | G | T | A | T | C | T | A | A | T | C | C |
| KX091847.1 | Plecoptera | *Amphinemura* sp. | A | A | T | A | A | T | A | G | G | G | T | A | T | C | T | A | A | T | C | C |
| MH085447.1 | Plecoptera | *Amphinemura yao* | A | A | T | A | A | T | A | G | G | G | T | A | T | C | T | A | A | T | C | C |
| NC044749.1 | Plecoptera | *Amphinemura yao* | A | A | T | A | A | T | A | G | G | G | T | A | T | C | T | A | A | T | C | C |
| MK111413.1 | Plecoptera | *Antarctoperla* | A | A | T | A | A | T | A | G | G | G | T | A | T | C | T | A | A | T | C | C |

| | | | | | | | | | | | | | | | | | | | | | |
|---|---|---|---|---|---|---|---|---|---|---|---|---|---|---|---|---|---|---|---|---|---|---|
| | | *michaelseni* | | | | | | | | | | | | | | | | | | | | |
| NC042199.1 | Plecoptera | *Antarctoperla michaelseni* | A | A | T | A | A | T | A | G | G | G | T | A | T | C | T | A | A | T | C | C |
| KR604721.1 | Plecoptera | *Apteroperla tikumana* | A | A | T | A | A | T | A | G | G | G | T | A | T | C | T | A | A | T | C | C |
| NC027698.1 | Plecoptera | *Apteroperla tikumana* | A | A | T | A | A | T | A | G | G | G | T | A | T | C | T | A | A | T | C | C |
| MT410838.1 | Plecoptera | *Brachyptera risi* | A | A | T | A | A | T | A | G | G | G | T | A | T | C | T | A | A | T | C | C |
| MG677941.1 | Plecoptera | *Calineuria stigmatica* | A | A | T | A | A | T | A | G | G | G | T | A | T | C | T | A | A | T | C | C |
| KX094942.1 | Plecoptera | *Capnia zijinshana* | A | A | T | A | A | T | A | G | G | G | T | A | T | C | T | A | A | T | C | C |
| NC034661.1 | Plecoptera | *Capnia zijinshana* | A | A | T | A | A | T | A | G | G | G | T | A | T | C | T | A | A | T | C | C |
| MG677942.1 | Plecoptera | *Caroperla siveci* | A | A | T | A | A | T | A | G | G | G | T | A | T | C | T | A | A | T | C | C |
| MF100783.1 | Plecoptera | *Cerconychia flectospina* | A | A | T | A | A | T | A | G | G | G | T | A | T | C | T | A | A | T | C | C |
| MN419914.1 | Plecoptera | *Claassenia* sp. | A | A | T | A | A | T | A | G | G | G | T | A | T | C | T | A | A | T | C | C |
| KC952026.1 | Plecoptera | *Cryptoperla* sp. | A | A | T | A | A | T | A | G | G | G | T | A | T | C | T | A | A | T | C | C |
| MK111416.1 | Plecoptera | *Diamphipnoa annulata* | A | A | T | A | A | T | A | G | G | G | T | A | T | C | T | A | A | T | C | C |
| NC042205.1 | Plecoptera | *Diamphipnoa annulata* | A | A | T | A | A | T | A | G | G | G | T | A | T | C | T | A | A | T | C | C |
| MK111417.1 | Plecoptera | *Diamphipnopsis* | A | A | T | A | A | T | A | G | G | G | T | A | T | C | T | A | A | T | C | C |

| | | | | | | | | | | | | | | | | | | | | | | |
|---|---|---|---|---|---|---|---|---|---|---|---|---|---|---|---|---|---|---|---|---|---|---|
| | | sp. | | | | | | | | | | | | | | | | | | | | |
| KF484757.1 | Plecoptera | *Dinocras cephalotes* | A | A | T | A | A | T | A | G | G | G | T | A | T | C | T | A | A | T | C | C |
| NC022843.1 | Plecoptera | *Dinocras cephalotes* | A | A | T | A | A | T | A | G | G | G | T | A | T | C | T | A | A | T | C | C |
| MG589787.1 | Plecoptera | *Doddsia occidentalis* | A | A | T | A | G | T | A | G | G | G | T | A | T | C | T | A | A | T | C | C |
| MK905888.1 | Plecoptera | *Etrocorema hochii* | A | A | T | A | A | T | A | G | G | G | T | A | T | C | T | A | A | T | C | C |
| MK905206.1 | Plecoptera | *Flavoperla* sp. | A | A | T | A | A | T | A | G | G | G | T | A | T | C | T | A | A | T | C | C |
| MN419916.1 | Plecoptera | *Flavoperla* sp. | A | A | T | A | A | T | A | G | G | G | T | A | T | C | T | A | A | T | C | C |
| MN419915.1 | Plecoptera | *Indonemoura auriformis* | A | A | T | A | A | T | A | G | G | G | T | A | T | C | T | A | A | T | C | C |
| MH085448.1 | Plecoptera | *Indonemoura jacobsoni* | A | A | T | A | A | T | A | G | G | G | T | A | T | C | T | A | A | T | C | C |
| NC044750.1 | Plecoptera | *Indonemoura jacobsoni* | A | A | T | A | A | T | A | G | G | G | T | A | T | C | T | A | A | T | C | C |
| MH085449.1 | Plecoptera | *Indonemoura nohirae* | A | A | T | A | G | T | A | G | G | G | T | A | T | C | T | A | A | T | C | C |
| NC044751.1 | Plecoptera | *Indonemoura nohirae* | A | A | T | A | G | T | A | G | G | G | T | A | T | C | T | A | A | T | C | C |
| MF716959.1 | Plecoptera | *Isoperla bilineata* | A | A | T | A | A | T | A | G | G | G | T | A | T | C | T | A | A | T | C | C |

| | | | | | | | | | | | | | | | | | | | | | | |
|---|---|---|---|---|---|---|---|---|---|---|---|---|---|---|---|---|---|---|---|---|---|---|
| NC038190.1 | Plecoptera | *Isoperla bilineata* | A | A | T | A | A | T | A | G | G | G | T | A | T | C | T | A | A | T | C | C |
| MG910457.1 | Plecoptera | *Isoperla eximia* | A | A | T | A | A | T | A | G | G | G | T | A | T | C | T | A | A | T | C | C |
| NC038167.1 | Plecoptera | *Isoperla eximia* | A | A | T | A | A | T | A | G | G | G | T | A | T | C | T | A | A | T | C | C |
| KT186102.1 | Plecoptera | *Kamimuria chungnanshana* | A | A | T | A | A | T | A | G | G | G | T | A | T | C | T | A | A | T | C | C |
| NC028076.1 | Plecoptera | *Kamimuria chungnanshana* | A | A | T | A | A | T | A | G | G | G | T | A | T | C | T | A | A | T | C | C |
| MN400755.1 | Plecoptera | *Kamimuria klapaleki* | A | A | T | A | A | T | A | G | G | G | T | A | T | C | T | A | A | T | C | C |
| KC894944.1 | Plecoptera | *Kamimuria wangi* | A | A | T | A | A | T | A | G | G | G | T | A | T | C | T | A | A | T | C | C |
| MK111414.1 | Plecoptera | *Klapopteryx armillata* | A | A | T | A | A | T | A | G | G | G | T | A | T | C | T | A | A | T | C | C |
| KP642637.1 | Plecoptera | *Mesocapnia arizonensis* | A | A | T | A | A | T | A | G | G | G | T | A | T | C | T | A | A | T | C | C |
| KY568983.1 | Plecoptera | *Mesocapnia daxingana* | A | A | T | A | A | T | A | G | G | G | T | A | T | C | T | A | A | T | C | C |
| MH085450.1 | Plecoptera | *Mesonemoura metafiligera* | A | A | T | A | A | T | A | G | G | G | T | A | T | C | T | A | A | T | C | C |
| NC044719.1 | Plecoptera | *Mesonemoura metafiligera* | A | A | T | A | A | T | A | G | G | G | T | A | T | C | T | A | A | T | C | C |
| MH085451.1 | Plecoptera | *Mesonemoura tritaenia* | A | A | T | A | A | T | A | G | G | G | T | A | T | C | T | A | A | T | C | C |

| | | | | | | | | | | | | | | | | | | | | | |
|---|---|---|---|---|---|---|---|---|---|---|---|---|---|---|---|---|---|---|---|---|---|
| NC044720.1 | Plecoptera | *Mesonemoura tritaenia* | A | A | T | A | A | T | A | G | G | G | T | A | T | C | T | A | A | T | C | C |
| MN096323.1 | Plecoptera | *Microperla geei* | A | A | T | A | A | T | A | G | G | G | T | A | T | C | T | A | A | T | C | C |
| MT410776.1 | Plecoptera | *Nemoura avicularis* | A | A | T | A | A | T | A | G | G | G | T | A | T | C | T | A | A | T | C | C |
| MT410849.1 | Plecoptera | *Nemoura cinerea* | A | A | T | A | A | T | A | G | G | G | T | A | T | C | T | A | A | T | C | C |
| KY940360.1 | Plecoptera | *Nemoura nankinensis* | A | A | T | A | A | T | A | G | G | G | T | A | T | C | T | A | A | T | C | C |
| NC034939.1 | Plecoptera | *Nemoura nankinensis* | A | A | T | A | A | T | A | G | G | G | T | A | T | C | T | A | A | T | C | C |
| MK290826.1 | Plecoptera | *Nemoura papilla* | A | A | T | A | A | T | A | G | G | G | T | A | T | C | T | A | A | T | C | C |
| MK111418.1 | Plecoptera | *Neonemura barrosi* | A | A | T | A | A | T | A | G | G | G | T | A | T | C | T | A | A | T | C | C |
| NC042206.1 | Plecoptera | *Neonemura barrosi* | A | A | T | A | A | T | A | G | G | G | T | A | T | C | T | A | A | T | C | C |
| KX091858.1 | Plecoptera | *Neoperla ignacsiveci* | A | A | T | A | A | T | A | G | G | G | T | A | T | C | T | A | A | T | C | C |
| KX091859.1 | Plecoptera | *Neoperla* sp. | A | A | T | A | A | T | A | G | G | G | T | A | T | C | T | A | A | T | C | C |
| MN400756.1 | Plecoptera | *Neoperlops gressitti* | A | A | T | A | A | T | A | G | G | G | T | A | T | C | T | A | A | T | C | C |
| MK111415.1 | Plecoptera | *Neuroperla schedingi* | A | A | T | A | A | T | A | G | G | G | T | A | T | C | T | A | A | T | C | C |

| | | | | | | | | | | | | | | | | | | | | | | |
|---|---|---|---|---|---|---|---|---|---|---|---|---|---|---|---|---|---|---|---|---|---|---|
| NC042200.1 | Plecoptera | *Neuroperla schedingi* | A | A | T | A | A | T | A | G | G | G | T | A | T | C | T | A | A | T | C | C |
| MK686067.1 | Plecoptera | *Niponiella limbatella* | A | A | T | A | A | T | A | G | G | G | T | A | T | C | T | A | A | T | C | C |
| MK387068.1 | Plecoptera | *Peltoperlopsis cebuano* | A | A | T | A | A | T | A | G | G | G | T | A | T | C | T | A | A | T | C | C |
| MF197377.1 | Plecoptera | *Perlodes* sp. | A | A | T | A | A | T | A | G | G | G | T | A | T | C | T | A | A | T | C | C |
| MH085452.1 | Plecoptera | *Protonemura kohnoae* | A | A | T | A | A | T | A | G | G | G | T | A | T | C | T | A | A | T | C | C |
| NC044752.1 | Plecoptera | *Protonemura kohnoae* | A | A | T | A | A | T | A | G | G | G | T | A | T | C | T | A | A | T | C | C |
| MT410850.1 | Plecoptera | *Protonemura meyeri* | A | A | T | A | A | T | A | G | G | G | T | A | T | C | T | A | A | T | C | C |
| NC050322.1 | Plecoptera | *Protonemura meyeri* | A | A | T | A | A | T | A | G | G | G | T | A | T | C | T | A | A | T | C | C |
| MH085453.1 | Plecoptera | *Protonemura orbiculata* | A | A | T | A | A | T | A | G | G | G | T | A | T | C | T | A | A | T | C | C |
| NC044753.1 | Plecoptera | *Protonemura orbiculata* | A | A | T | A | A | T | A | G | G | G | T | A | T | C | T | A | A | T | C | C |
| MG910458.1 | Plecoptera | *Pseudomegarcys japonica* | A | A | T | A | A | T | A | G | G | G | T | A | T | C | T | A | A | T | C | C |
| NC038168.1 | Plecoptera | *Pseudomegarcys* | A | A | T | A | A | T | A | G | G | G | T | A | T | C | T | A | A | T | C | C |

| | | | | | | | | | | | | | | | | | | | | | | |
|---|---|---|---|---|---|---|---|---|---|---|---|---|---|---|---|---|---|---|---|---|---|---|
| | | *japonica* | | | | | | | | | | | | | | | | | | | | |
| KU182360.1 | Plecoptera | *Pteronarcella badia* | A | A | T | A | A | T | A | G | G | G | T | A | T | C | T | A | A | T | C | C |
| NC029248.1 | Plecoptera | *Pteronarcella badia* | A | A | T | A | A | T | A | G | G | G | T | A | T | C | T | A | A | T | C | C |
| MK111419.1 | Plecoptera | *Rhopalopsole bulbifera* | T | A | T | A | A | T | A | G | G | G | T | A | T | C | T | A | A | T | C | C |
| NC042207.1 | Plecoptera | *Rhopalopsole bulbifera* | T | A | T | A | A | T | A | G | G | G | T | A | T | C | T | A | A | T | C | C |
| MH510071.1 | Plecoptera | *Scopura longa* | A | A | T | A | A | T | A | G | G | G | T | A | T | C | T | A | A | T | C | C |
| NC041105.1 | Plecoptera | *Scopura longa* | A | A | T | A | A | T | A | G | G | G | T | A | T | C | T | A | A | T | C | C |
| MK492253.1 | Plecoptera | *Sinacroneuria dabieshana* | A | A | T | A | A | T | A | G | G | G | T | A | T | C | T | A | A | T | C | C |
| MF716958.1 | Plecoptera | *Soliperla* sp. | A | A | T | A | A | T | A | G | G | G | T | A | T | C | T | A | A | T | C | C |
| NC038189.1 | Plecoptera | *Soliperla* sp. | A | A | T | A | A | T | A | G | G | G | T | A | T | C | T | A | A | T | C | C |
| MH085455.1 | Plecoptera | *Sphaeronemoura acutispina* | A | A | T | A | A | T | A | G | G | G | T | A | T | C | T | A | A | T | C | C |
| MH085454.1 | Plecoptera | *Sphaeronemoura grandicauda* | A | A | T | A | A | T | A | G | G | G | T | A | T | C | T | A | A | T | C | C |
| NC044754.1 | Plecoptera | *Sphaeronemoura grandicauda* | A | A | T | A | A | T | A | G | G | G | T | A | T | C | T | A | A | T | C | C |
| MK111420.1 | Plecoptera | *Sphaeronemoura* | A | A | T | A | A | T | A | G | G | G | T | A | T | C | T | A | A | T | C | C |

| | | | | | | | | | | | | | | | | | | | | | | |
|---|---|---|---|---|---|---|---|---|---|---|---|---|---|---|---|---|---|---|---|---|---|---|
| | | *hainana* | | | | | | | | | | | | | | | | | | | | |
| NC044755.1 | Plecoptera | *Sphaeronemoura hamistyla* | A | A | T | A | A | T | A | G | G | G | T | A | T | C | T | A | A | T | C | C |
| KR088971.1 | Plecoptera | *Styloperla* sp. | A | A | T | A | A | T | A | G | G | G | T | A | T | C | T | A | A | T | C | C |
| KX845569.1 | Plecoptera | *Styloperla spinicercia* | A | A | T | A | A | T | A | G | G | G | T | A | T | C | T | A | A | T | C | C |
| NC034809.1 | Plecoptera | *Styloperla spinicercia* | A | A | T | A | A | T | A | G | G | G | T | A | T | C | T | A | A | T | C | C |
| KX091865.1 | Plecoptera | *Suwallia bimaculata* | A | A | T | A | A | T | A | G | G | G | T | A | T | C | T | A | A | T | C | C |
| MN121757.1 | Plecoptera | *Suwallia bimaculata* | A | A | T | A | A | T | A | G | G | G | T | A | T | C | T | A | A | T | C | C |
| MF198253.1 | Plecoptera | *Suwallia teleckojensis* | A | A | T | A | A | T | A | G | G | G | T | A | T | C | T | A | A | T | C | C |
| NC037754.1 | Plecoptera | *Suwallia teleckojensis* | A | A | T | A | A | T | A | G | G | G | T | A | T | C | T | A | A | T | C | C |
| MG589786.1 | Plecoptera | *Taeniopteryx ugola* | A | A | T | A | A | T | A | G | G | G | T | A | T | C | T | A | A | T | C | C |
| NC037897.1 | Plecoptera | *Taeniopteryx ugola* | A | A | T | A | A | T | A | G | G | G | T | A | T | C | T | A | A | T | C | C |
| KM409708.1 | Plecoptera | *Togoperla* sp. | A | A | T | A | A | T | A | G | G | G | T | A | T | C | T | A | A | T | C | C |
| KY522907.1 | Plecoptera | *Zelandoperla* | A | A | T | A | A | T | A | G | G | G | T | A | T | C | T | A | A | T | C | C |

| | | | | | | | | | | | | | | | | | | | | | | |
|---|---|---|---|---|---|---|---|---|---|---|---|---|---|---|---|---|---|---|---|---|---|---|
| | | *fenestrata* | | | | | | | | | | | | | | | | | | | | |
| NC034997.1 | Plecoptera | *Zelandoperla fenestrata* | A | A | T | A | A | T | A | G | G | G | T | A | T | C | T | A | A | T | C | C |
| FJ456939.1 | Heteroptera | *Aphelocheirus ellipsoideus* | A | A | T | A | A | T | A | G | G | G | T | A | T | C | T | A | A | T | C | C |
| KX505849.1 | Heteroptera | *Aphelocheirus jendeki* | A | A | T | A | A | T | A | G | G | G | T | A | T | C | T | A | A | T | C | C |
| NC037370.1 | Heteroptera | *Aphelocheirus jendeki* | A | A | T | A | A | T | A | G | G | G | T | A | T | C | T | A | A | T | C | C |
| FJ456944.1 | Heteroptera | *Aquarius paludum* | A | A | T | A | A | T | A | G | G | G | T | A | T | C | T | A | A | T | C | C |
| NC012841.1 | Heteroptera | *Aquarius paludum* | A | A | T | A | A | T | A | G | G | G | T | A | T | C | T | A | A | T | C | C |
| MK251109.1 | Heteroptera | *Diplonychus esakii* | A | A | T | A | A | T | A | G | G | G | T | A | T | C | T | A | A | T | C | C |
| FJ456940.1 | Heteroptera | *Diplonychus rusticus* | A | A | T | A | A | T | A | G | G | G | T | A | T | C | T | A | A | T | C | C |
| FJ456949.1 | Heteroptera | *Enithares tibialis* | A | A | T | A | A | T | A | G | G | G | T | A | T | C | T | A | A | T | C | C |
| NC012819.1 | Heteroptera | *Enithares tibialis* | A | A | T | A | A | T | A | G | G | G | T | A | T | C | T | A | A | T | C | C |
| KP400582.1 | Heteroptera | *Entomovelia* sp. | A | A | T | A | A | T | A | G | G | G | T | A | T | C | T | A | A | T | C | C |
| MF177288.1 | Heteroptera | *Gigantometra gigas* | A | A | T | A | A | T | A | G | G | G | T | A | T | C | T | A | A | T | C | C |

| | | | | | | | | | | | | | | | | | | | | | | |
|---|---|---|---|---|---|---|---|---|---|---|---|---|---|---|---|---|---|---|---|---|---|---|
| MF177289.1 | Heteroptera | *Gigantometra gigas* | A | A | T | A | A | T | A | G | G | G | T | A | T | C | T | A | A | T | C | C |
| MF177290.1 | Heteroptera | *Gigantometra gigas* | A | A | T | A | A | T | A | G | G | G | T | A | T | C | T | A | A | T | C | C |
| NC041084.1 | Heteroptera | *Gigantometra gigas* | A | A | T | A | A | T | A | G | G | G | T | A | T | C | T | A | A | T | C | C |
| KJ027513.1 | Heteroptera | *Helotrephes semiglobosus semiglobosus* | A | A | T | A | A | T | A | G | G | G | T | A | T | C | T | A | A | T | C | C |
| FJ456951.1 | Heteroptera | *Helotrephes* sp. | A | A | T | A | A | T | A | G | G | G | T | A | T | C | T | A | A | T | C | C |
| NC012822.1 | Heteroptera | *Helotrephes* sp. | A | A | T | A | A | T | A | G | G | G | T | A | T | C | T | A | A | T | C | C |
| FJ456945.1 | Heteroptera | *Hydrometra greeni* | A | A | T | A | A | T | A | G | G | G | T | A | T | C | T | A | A | T | C | C |
| NC012842.1 | Heteroptera | *Hydrometra greeni* | A | A | T | A | A | T | A | G | G | G | T | A | T | C | T | A | A | T | C | C |
| FJ456947.1 | Heteroptera | *Ilyocoris cimicoides* | A | A | T | A | A | T | A | G | G | G | T | A | T | C | T | A | A | T | C | C |
| NC012845.1 | Heteroptera | *Ilyocoris cimicoides* | A | A | T | A | A | T | A | G | G | G | T | A | T | C | T | A | A | T | C | C |
| FJ456948.1 | Heteroptera | *Laccotrephes robustus* | C | A | T | A | A | T | A | G | G | G | T | A | T | C | T | A | A | T | C | C |
| NC012817.1 | Heteroptera | *Laccotrephes* | C | A | T | A | A | T | A | G | G | G | T | A | T | C | T | A | A | T | C | C |

| | | | | | | | | | | | | | | | | | | | | | |
|---|---|---|---|---|---|---|---|---|---|---|---|---|---|---|---|---|---|---|---|---|---|---|
| | | *robustus* | | | | | | | | | | | | | | | | | | | | |
| FJ456946.1 | Heteroptera | *Leptopus* sp. | A | A | T | A | A | T | A | G | G | G | T | A | T | C | T | A | A | T | C | C |
| JQ910985.1 | Heteroptera | *Lethocerus deyrollei* | A | A | T | A | A | T | A | G | G | G | T | A | T | C | T | A | A | T | C | C |
| KU237288.1 | Heteroptera | *Lethocerus deyrollei* | A | A | T | A | A | T | A | G | G | G | T | A | T | C | T | A | A | T | C | C |
| KM588201.1 | Heteroptera | *Lethocerus indicus* | A | A | T | A | A | T | A | G | G | G | T | A | T | C | T | A | A | T | C | C |
| KR072671.1 | Heteroptera | *Lethocerus indicus* | A | A | T | A | A | T | A | G | G | G | T | A | T | C | T | A | A | T | C | C |
| NC027194.1 | Heteroptera | *Lethocerus indicus* | A | A | T | A | A | T | A | G | G | G | T | A | T | C | T | A | A | T | C | C |
| MN027272.1 | Heteroptera | *Limnogonus hypoleucus* | T | A | T | A | A | T | A | G | G | G | T | A | T | C | T | A | A | T | C | C |
| NC044722.1 | Heteroptera | *Limnogonus hypoleucus* | T | A | T | A | A | T | A | G | G | G | T | A | T | C | T | A | A | T | C | C |
| MN027275.1 | Heteroptera | *Limnogonus intermedius* | T | A | T | A | A | T | A | G | G | G | T | A | T | C | T | A | A | T | C | C |
| NC044724.1 | Heteroptera | *Limnogonus intermedius* | T | A | T | A | A | T | A | G | G | G | T | A | T | C | T | A | A | T | C | C |
| MN027273.1 | Heteroptera | *Limnogonus poissoni* | T | A | T | A | A | T | A | G | G | G | T | A | T | C | T | A | A | T | C | C |

| | | | | | | | | | | | | | | | | | | | | | | |
|---|---|---|---|---|---|---|---|---|---|---|---|---|---|---|---|---|---|---|---|---|---|---|
| MN027274.1 | Heteroptera | *Limnogonus poissoni* | T | A | T | A | A | T | A | G | G | G | T | A | T | C | T | A | A | T | C | C |
| MN027276.1 | Heteroptera | *Limnogonus poissoni* | T | A | T | A | A | T | A | G | G | G | T | A | T | C | T | A | A | T | C | C |
| NC044723.1 | Heteroptera | *Limnogonus poissoni* | T | A | T | A | A | T | A | G | G | G | T | A | T | C | T | A | A | T | C | C |
| FJ456942.1 | Heteroptera | *Lycorma delicatula* | A | A | T | A | A | C | A | G | G | G | T | A | T | C | T | A | A | T | C | C |
| NC012835.1 | Heteroptera | *Lycorma delicatula* | A | A | T | A | A | C | A | G | G | G | T | A | T | C | T | A | A | T | C | C |
| MN027278.1 | Heteroptera | *Metrocoris* sp. | A | A | T | A | A | T | A | G | G | G | T | A | T | C | T | A | A | T | C | C |
| MH382835.1 | Heteroptera | *Micronecta sahlbergii* | A | A | T | A | A | T | A | G | G | G | T | A | T | C | T | A | A | T | C | C |
| NC039588.1 | Heteroptera | *Micronecta sahlbergii* | A | A | T | A | A | T | A | G | G | G | T | A | T | C | T | A | A | T | C | C |
| KT031802.1 | Heteroptera | *Nepa hoffmanni* | A | A | T | A | A | T | A | G | G | G | T | A | T | C | T | A | A | T | C | C |
| NC028084.1 | Heteroptera | *Nepa hoffmanni* | A | A | T | A | A | T | A | G | G | G | T | A | T | C | T | A | A | T | C | C |
| FJ456943.1 | Heteroptera | *Nerthra indica* | A | A | T | A | A | T | A | G | G | G | T | A | T | C | T | A | A | T | C | C |
| NC012838.1 | Heteroptera | *Nerthra indica* | A | A | T | A | A | T | A | G | G | G | T | A | T | C | T | A | A | T | C | C |
| KX034086.1 | Heteroptera | *Notonecta chinensis* | A | A | T | A | A | T | A | G | G | G | T | A | T | C | T | A | A | T | C | C |
| NC036671.1 | Heteroptera | *Notonecta* | A | A | T | A | A | T | A | G | G | G | T | A | T | C | T | A | A | T | C | C |

| | | | | | | | | | | | | | | | | | | | | | | |
|---|---|---|---|---|---|---|---|---|---|---|---|---|---|---|---|---|---|---|---|---|---|---|
| | | *chinensis* | | | | | | | | | | | | | | | | | | | | |
| MH780861.1 | Heteroptera | *Notonecta montandoni* | A | A | T | A | A | T | A | G | G | G | T | A | T | C | T | A | A | T | C | C |
| NC045886.1 | Heteroptera | *Notonecta montandoni* | A | A | T | A | A | T | A | G | G | G | T | A | T | C | T | A | A | T | C | C |
| KM244707.1 | Heteroptera | *Notonectidae* sp. | A | A | T | A | A | T | A | G | G | G | T | A | T | C | T | A | A | T | C | C |
| FJ456950.1 | Heteroptera | *Ochterus marginatus* | A | A | T | A | A | T | A | G | G | G | T | A | T | C | T | A | A | T | C | C |
| JQ910988.1 | Heteroptera | *Perittopus* sp. | T | A | T | A | A | T | A | G | G | G | T | A | T | C | T | A | A | T | C | C |
| KP400583.1 | Heteroptera | *Ptilomera tigrina* | C | A | T | A | A | T | A | G | G | G | T | A | T | C | T | A | A | T | C | C |
| MN027271.1 | Heteroptera | *Rhagovelia reitteri* | A | A | T | A | A | T | A | G | G | G | T | A | T | C | T | A | A | T | C | C |
| NC044721.1 | Heteroptera | *Rhagovelia reitteri* | A | A | T | A | A | T | A | G | G | G | T | A | T | C | T | A | A | T | C | C |
| MN027279.1 | Heteroptera | *Rhagovelia* sp. | A | A | T | A | A | T | A | G | G | G | T | A | T | C | T | A | A | T | C | C |
| FJ456941.1 | Heteroptera | *Sigara septemlineata* | A | A | T | A | A | T | A | G | G | G | T | A | T | C | T | A | A | T | C | C |
| MN027277.1 | Heteroptera | *Trepobates* sp. | A | A | T | A | A | T | A | G | G | G | T | A | T | C | T | A | A | T | C | C |
| FJ456952.1 | Heteroptera | *Valentia hoffmanni* | A | A | T | A | A | T | A | G | G | G | T | A | T | C | T | A | A | T | C | C |
| NC012823.1 | Heteroptera | *Valentia hoffmanni* | A | A | T | A | A | T | A | G | G | G | T | A | T | C | T | A | A | T | C | C |

| | | | | | | | | | | | | | | | | | | | | | | | |
|---|---|---|---|---|---|---|---|---|---|---|---|---|---|---|---|---|---|---|---|---|---|---|---|
| KT425071.1 | Neuroptera | *Nallachius americanus* | | A | A | T | A | A | T | A | G | G | G | T | A | T | C | T | A | A | T | C | C |
| | | | A | 82.3 | 98.5 | 0.0 | 100.0 | 88.7 | 19.6 | 95.5 | 0.0 | 0.0 | 0.0 | 0.0 | 95.5 | 0.0 | 0.0 | 0.0 | 100.0 | 100.0 | 0.0 | 0.0 | 0.0 |
| | | | T | 14.0 | 0.8 | 100.0 | 0.0 | 0.0 | 77.0 | 0.0 | 0.0 | 0.0 | 0.0 | 100.0 | 0.0 | 100.0 | 0.0 | 100.0 | 0.0 | 0.0 | 100.0 | 0.0 | 0.0 |
| | | | G | 1.1 | 0.8 | 0.0 | 0.0 | 11.3 | 2.6 | 4.5 | 100.0 | 100.0 | 100.0 | 0.0 | 0.0 | 0.0 | 0.0 | 0.0 | 0.0 | 0.0 | 0.0 | 0.0 | 0.0 |
| | | | C | 2.6 | 0.0 | 0.0 | 0.0 | 0.0 | 0.8 | 0.0 | 0.0 | 0.0 | 0.0 | 0.0 | 4.5 | 0.0 | 100.0 | 0.0 | 0.0 | 0.0 | 0.0 | 100.0 | 100.0 |

Table S8. Nucleotide sequences of universal primer (MtInsects-12S_Hol_R) and homologous region of 209 sequences of insects used for development in this study.

| | | | | | | | | | | | | | | | | | | | | | | | |
|---|---|---|---|---|---|---|---|---|---|---|---|---|---|---|---|---|---|---|---|---|---|---|---|
| MtInsects-12S_Hol_R | | | | T | A | R | T | A | G | G | G | T | A | T | C | T | A | A | T | C | C | T | A | G |
| | | | A | 0 | 209 | 202 | 0 | 209 | 0 | 0 | 0 | 0 | 209 | 0 | 0 | 0 | 209 | 209 | 0 | 0 | 0 | 0 | 209 | 0 |
| | | | T | 209 | 0 | 0 | 209 | 0 | 0 | 0 | 0 | 209 | 0 | 209 | 0 | 209 | 0 | 0 | 209 | 0 | 0 | 209 | 0 | 0 |
| | | | G | 0 | 0 | 7 | 0 | 0 | 209 | 209 | 209 | 0 | 0 | 0 | 0 | 0 | 0 | 0 | 0 | 0 | 0 | 0 | 0 | 209 |
| | | | C | 0 | 0 | 0 | 0 | 0 | 0 | 0 | 0 | 0 | 0 | 0 | 209 | 0 | 0 | 0 | 0 | 209 | 209 | 0 | 0 | 0 |
| GenBank Acc. No. | Order | Species | | | | | | | | | | | | | | | | | | | | | | |
| KF840564.1 | Megaloptera | *Acanthacorydalis orientalis* | | T | A | A | T | A | G | G | G | T | A | T | C | T | A | A | T | C | C | T | A | G |
| NC023462.1 | Megaloptera | *Acanthacorydalis orientalis* | | T | A | A | T | A | G | G | G | T | A | T | C | T | A | A | T | C | C | T | A | G |
| KU925864.1 | Megaloptera | *Archichauliodes deceptor* | | T | A | A | T | A | G | G | G | T | A | T | C | T | A | A | T | C | C | T | A | G |
| NC035504.1 | Megaloptera | *Archichauliodes deceptor* | | T | A | A | T | A | G | G | G | T | A | T | C | T | A | A | T | C | C | T | A | G |
| KU302713.1 | Megaloptera | *Chloronia mirifica* | | T | A | A | T | A | G | G | G | T | A | T | C | T | A | A | T | C | C | T | A | G |
| NC033349.1 | Megaloptera | *Chloronia mirifica* | | T | A | A | T | A | G | G | G | T | A | T | C | T | A | A | T | C | C | T | A | G |
| FJ171323.1 | Megaloptera | *Corydalus cornutus* | | T | A | A | T | A | G | G | G | T | A | T | C | T | A | A | T | C | C | T | A | G |
| NC011276.1 | Megaloptera | *Corydalus cornutus* | | T | A | A | T | A | G | G | G | T | A | T | C | T | A | A | T | C | C | T | A | G |

| | | | | | | | | | | | | | | | | | | | | | | |
|---|---|---|---|---|---|---|---|---|---|---|---|---|---|---|---|---|---|---|---|---|---|---|
| KJ806318.1 | Megaloptera | *Dysmicohermes ingens* | T | A | A | T | A | G | G | G | T | A | T | C | T | A | A | T | C | C | T | A | G |
| NC024657.1 | Megaloptera | *Dysmicohermes ingens* | T | A | A | T | A | G | G | G | T | A | T | C | T | A | A | T | C | C | T | A | G |
| JQ351950.1 | Megaloptera | *Neochauliodes bowringi* | T | A | A | T | A | G | G | G | T | A | T | C | T | A | A | T | C | C | T | A | G |
| NC023444.1 | Megaloptera | *Neochauliodes bowringi* | T | A | A | T | A | G | G | G | T | A | T | C | T | A | A | T | C | C | T | A | G |
| KF785799.1 | Megaloptera | *Neochauliodes fraternus* | T | A | A | T | A | G | G | G | T | A | T | C | T | A | A | T | C | C | T | A | G |
| NC025282.1 | Megaloptera | *Neochauliodes fraternus* | T | A | A | T | A | G | G | G | T | A | T | C | T | A | A | T | C | C | T | A | G |
| KX821680.1 | Megaloptera | *Neochauliodes parasparsus* | T | A | A | T | A | G | G | G | T | A | T | C | T | A | A | T | C | C | T | A | G |
| JX110703.1 | Megaloptera | *Neochauliodes punctatolosus* | T | A | A | T | A | G | G | G | T | A | T | C | T | A | A | T | C | C | T | A | G |
| NC018772.1 | Megaloptera | *Neochauliodes punctatolosus* | T | A | A | T | A | G | G | G | T | A | T | C | T | A | A | T | C | C | T | A | G |
| KF771894.1 | Megaloptera | *Neochauliodes rotundatus* | T | A | A | T | A | G | G | G | T | A | T | C | T | A | A | T | C | C | T | A | G |
| NC025281.1 | Megaloptera | *Neochauliodes rotundatus* | T | A | A | T | A | G | G | G | T | A | T | C | T | A | A | T | C | C | T | A | G |

| | | | | | | | | | | | | | | | | | | | | | | |
|---|---|---|---|---|---|---|---|---|---|---|---|---|---|---|---|---|---|---|---|---|---|---|
| KP126231.1 | Megaloptera | *Neoneuromus tonkinensis* | T | A | A | T | A | G | G | G | T | A | T | C | T | A | A | T | C | C | T | A | G |
| NC027852.1 | Megaloptera | *Neoneuromus tonkinensis* | T | A | A | T | A | G | G | G | T | A | T | C | T | A | A | T | C | C | T | A | G |
| KP126232.1 | Megaloptera | *Nevromus exterior* | T | A | A | T | A | G | G | G | T | A | T | C | T | A | A | T | C | C | T | A | G |
| NC027851.1 | Megaloptera | *Nevromus exterior* | T | A | A | T | A | G | G | G | T | A | T | C | T | A | A | T | C | C | T | A | G |
| KY230493.1 | Megaloptera | *Protochauliodes biconicus* | T | A | A | T | A | G | G | G | T | A | T | C | T | A | A | T | C | C | T | A | G |
| EU526394.1 | Megaloptera | *Protohermes concolorus* | T | A | A | T | A | G | G | G | T | A | T | C | T | A | A | T | C | C | T | A | G |
| NC011524.1 | Megaloptera | *Protohermes concolorus* | T | A | A | T | A | G | G | G | T | A | T | C | T | A | A | T | C | C | T | A | G |
| FJ859905.1 | Megaloptera | *Sialis hamata* | T | A | A | T | A | G | G | G | T | A | T | C | T | A | A | T | C | C | T | A | G |
| NC013256.1 | Megaloptera | *Sialis hamata* | T | A | A | T | A | G | G | G | T | A | T | C | T | A | A | T | C | C | T | A | G |
| KT876911.1 | Megaloptera | *Sialis* sp. | T | A | A | T | A | G | G | G | T | A | T | C | T | A | A | T | C | C | T | A | G |
| KT425084.1 | Megaloptera | *Stenosialis australiensis* | T | A | A | T | A | G | G | G | T | A | T | C | T | A | A | T | C | C | T | A | G |
| KY587199.1 | Neuroptera | *Abachrysa eureka* | T | A | A | T | A | G | G | G | T | A | T | C | T | A | A | T | C | C | T | A | G |
| NC015095.1 | Neuroptera | *Apochrysa matsumurae* | T | A | A | T | A | G | G | G | T | A | T | C | T | A | A | T | C | C | T | A | G |
| KC758703.1 | Neuroptera | *Ascalohybris subjacens* | T | A | A | T | A | G | G | G | T | A | T | C | T | A | A | T | C | C | T | A | G |

| | | | | | | | | | | | | | | | | | | | | | | |
|---|---|---|---|---|---|---|---|---|---|---|---|---|---|---|---|---|---|---|---|---|---|---|
| NC021428.1 | Neuroptera | *Ascalohybris subjacens* | T | A | A | T | A | G | G | G | T | A | T | C | T | A | A | T | C | C | T | A | G |
| FJ171324.1 | Neuroptera | *Ascaloptynx appendiculatus* | T | A | A | T | A | G | G | G | T | A | T | C | T | A | A | T | C | C | T | A | G |
| NC011277.1 | Neuroptera | *Ascaloptynx appendiculatus* | T | A | A | T | A | G | G | G | T | A | T | C | T | A | A | T | C | C | T | A | G |
| KT425083.1 | Neuroptera | *Balmes birmanus* | T | A | A | T | A | G | G | G | T | A | T | C | T | A | A | T | C | C | T | A | G |
| KX369241.1 | Neuroptera | *Bullanga florida* | T | A | A | T | A | G | G | G | T | A | T | C | T | A | A | T | C | C | T | A | G |
| NC032298.1 | Neuroptera | *Bullanga florida* | T | A | A | T | A | G | G | G | T | A | T | C | T | A | A | T | C | C | T | A | G |
| KT425069.1 | Neuroptera | *Chasmoptera huttii* | T | A | A | T | A | G | G | G | T | A | T | C | T | A | A | T | C | C | T | A | G |
| JX033119.1 | Neuroptera | *Chrysopa pallens* | T | A | A | T | A | G | G | G | T | A | T | C | T | A | A | T | C | C | T | A | G |
| NC019618.1 | Neuroptera | *Chrysopa pallens* | T | A | A | T | A | G | G | G | T | A | T | C | T | A | A | T | C | C | T | A | G |
| KU877169.1 | Neuroptera | *Chrysoperla externa* | T | A | A | T | A | G | G | G | T | A | T | C | T | A | A | T | C | C | T | A | G |
| NC030341.1 | Neuroptera | *Chrysoperla externa* | T | A | A | T | A | G | G | G | T | A | T | C | T | A | A | T | C | C | T | A | G |
| KP027407.1 | Neuroptera | *Chrysoperla nipponensis* | T | A | A | T | A | G | G | G | T | A | T | C | T | A | A | T | C | C | T | A | G |
| NC015093.1 | Neuroptera | *Chrysoperla nipponensis* | T | A | A | T | A | G | G | G | T | A | T | C | T | A | A | T | C | C | T | A | G |
| KT425088.1 | Neuroptera | *Climacia areolaris* | T | A | A | T | A | G | G | G | T | A | T | C | T | A | A | T | C | C | T | A | G |
| KT425068.1 | Neuroptera | *Dendroleon* | T | A | A | T | A | G | G | G | T | A | T | C | T | A | A | T | C | C | T | A | G |

| | | | | | | | | | | | | | | | | | | | | | | |
|---|---|---|---|---|---|---|---|---|---|---|---|---|---|---|---|---|---|---|---|---|---|---|
| | | *pantherinus* | | | | | | | | | | | | | | | | | | | | |
| MK301246.1 | Neuroptera | *Dendroleon pantherinus* | T | A | A | T | A | G | G | G | T | A | T | C | T | A | A | T | C | C | T | A | G |
| KT425073.1 | Neuroptera | *Dilar* sp. | T | A | A | T | A | G | G | G | T | A | T | C | T | A | A | T | C | C | T | A | G |
| FJ859906.1 | Neuroptera | *Ditaxis biseriata* | T | A | A | T | A | G | G | G | T | A | T | C | T | A | A | T | C | C | T | A | G |
| NC013257.1 | Neuroptera | *Ditaxis biseriata* | T | A | A | T | A | G | G | G | T | A | T | C | T | A | A | T | C | C | T | A | G |
| KT425087.1 | Neuroptera | *Drepanepteryx phalaenoides* | T | A | A | T | A | G | G | G | T | A | T | C | T | A | A | T | C | C | T | A | G |
| KF701327.1 | Neuroptera | *Epacanthaclisis banksi* | T | A | A | T | A | G | G | G | T | A | T | C | T | A | A | T | C | C | T | A | G |
| NC025905.1 | Neuroptera | *Epacanthaclisis banksi* | T | A | A | T | A | G | G | G | T | A | T | C | T | A | A | T | C | C | T | A | G |
| MF972496.1 | Neuroptera | *Euclimacia badia* | T | A | A | T | A | G | G | G | T | A | T | C | T | A | A | T | C | C | T | A | G |
| NC039773.1 | Neuroptera | *Euclimacia badia* | T | A | A | T | A | G | G | G | T | A | T | C | T | A | A | T | C | C | T | A | G |
| KT425080.1 | Neuroptera | *Eumantispa harmandi* | T | A | A | T | A | G | G | G | T | A | T | C | T | A | A | T | C | C | T | A | G |
| MF972495.1 | Neuroptera | *Eumantispa harmandi* | T | A | A | T | A | G | G | G | T | A | T | C | T | A | A | T | C | C | T | A | G |
| NC039772.1 | Neuroptera | *Eumantispa harmandi* | T | A | A | T | A | G | G | G | T | A | T | C | T | A | A | T | C | C | T | A | G |
| KT425072.1 | Neuroptera | *Fontecilla graphicus* | T | A | A | T | A | G | G | G | T | A | T | C | T | A | A | T | C | C | T | A | G |

| | | | | | | | | | | | | | | | | | | | | | | |
|---|---|---|---|---|---|---|---|---|---|---|---|---|---|---|---|---|---|---|---|---|---|---|
| KY364372.1 | Neuroptera | *Gatzara jezoensis* | T | A | A | T | A | G | G | G | T | A | T | C | T | A | A | T | C | C | T | A | G |
| MK341582.1 | Neuroptera | *Gryposmylus pennyi* | T | A | A | T | A | G | G | G | T | A | T | C | T | A | A | T | C | C | T | A | G |
| MN852445.1 | Neuroptera | *Hemerobius japonicus* | T | A | A | T | A | G | G | G | T | A | T | C | T | A | A | T | C | C | T | A | G |
| NC048517.1 | Neuroptera | *Hemerobius japonicus* | T | A | A | T | A | G | G | G | T | A | T | C | T | A | A | T | C | C | T | A | G |
| KT425077.1 | Neuroptera | *Heterosmylus* sp. | T | A | A | T | A | G | G | G | T | A | T | C | T | A | A | T | C | C | T | A | G |
| KT425085.1 | Neuroptera | *Inocellia fujiana* | T | A | A | T | A | G | G | G | T | A | T | C | T | A | A | T | C | C | T | A | G |
| KY587200.1 | Neuroptera | *Italochrysa insignis* | T | A | A | T | A | G | G | G | T | A | T | C | T | A | A | T | C | C | T | A | G |
| KY587201.1 | Neuroptera | *Leucochrysa pretiosa* | T | A | A | T | A | G | G | G | T | A | T | C | T | A | A | T | C | C | T | A | G |
| NC015609.1 | Neuroptera | *Libelloides macaronius* | T | A | A | T | A | G | G | G | T | A | T | C | T | A | A | T | C | C | T | A | G |
| MF972494.1 | Neuroptera | *Mantispa japonica* | T | A | A | T | A | G | G | G | T | A | T | C | T | A | A | T | C | C | T | A | G |
| NC039771.1 | Neuroptera | *Mantispa japonica* | T | A | A | T | A | G | G | G | T | A | T | C | T | A | A | T | C | C | T | A | G |
| MK472073.1 | Neuroptera | *Micromus angulatus* | T | A | G | T | A | G | G | G | T | A | T | C | T | A | A | T | C | C | T | A | G |
| NC042677.1 | Neuroptera | *Micromus angulatus* | T | A | G | T | A | G | G | G | T | A | T | C | T | A | A | T | C | C | T | A | G |
| KT425075.1 | Neuroptera | *Micromus* sp. | T | A | G | T | A | G | G | G | T | A | T | C | T | A | A | T | C | C | T | A | G |

| | | | | | | | | | | | | | | | | | | | | | | |
|---|---|---|---|---|---|---|---|---|---|---|---|---|---|---|---|---|---|---|---|---|---|---|
| KT425092.1 | Neuroptera | *Mucroberotha vesicaria* | T | A | A | T | A | G | G | G | T | A | T | C | T | A | A | T | C | C | T | A | G |
| KT425089.1 | Neuroptera | *Myiodactylus osmyloides* | T | A | A | T | A | G | G | G | T | A | T | C | T | A | A | T | C | C | T | A | G |
| MT118664.1 | Neuroptera | *Myrmeleon formicarius* | T | A | A | T | A | G | G | G | T | A | T | C | T | A | A | T | C | C | T | A | G |
| NC047285.1 | Neuroptera | *Myrmeleon formicarius* | T | A | A | T | A | G | G | G | T | A | T | C | T | A | A | T | C | C | T | A | G |
| KJ461323.1 | Neuroptera | *Myrmeleon immanis* | T | A | A | T | A | G | G | G | T | A | T | C | T | A | A | T | C | C | T | A | G |
| NC024826.1 | Neuroptera | *Myrmeleon immanis* | T | A | A | T | A | G | G | G | T | A | T | C | T | A | A | T | C | C | T | A | G |
| KT425086.1 | Neuroptera | *Negha inflata* | T | A | A | T | A | G | G | G | T | A | T | C | T | A | A | T | C | C | T | A | G |
| KT425079.1 | Neuroptera | *Nemoptera coa* | T | A | A | T | A | G | G | G | T | A | T | C | T | A | A | T | C | C | T | A | G |
| KR078257.1 | Neuroptera | *Neuronema laminatum* | T | A | A | T | A | G | G | G | T | A | T | C | T | A | A | T | C | C | T | A | G |
| NC028153.1 | Neuroptera | *Neuronema laminatum* | T | A | A | T | A | G | G | G | T | A | T | C | T | A | A | T | C | C | T | A | G |
| KT425074.1 | Neuroptera | *Nevrorthus apatelios* | T | A | A | T | A | G | G | G | T | A | T | C | T | A | A | T | C | C | T | A | G |
| KT425076.1 | Neuroptera | *Nipponeurorthus fuscinervis* | T | A | A | T | A | G | G | G | T | A | T | C | T | A | A | T | C | C | T | A | G |

| | | | | | | | | | | | | | | | | | | | | | | |
|---|---|---|---|---|---|---|---|---|---|---|---|---|---|---|---|---|---|---|---|---|---|---|
| KP264629.1 | Neuroptera | *Nothancyla verreauxi* | T | A | A | T | A | G | G | G | T | A | T | C | T | A | A | T | C | C | T | A | G |
| KP264630.1 | Neuroptera | *Nothochrysa* sp. | T | A | A | T | A | G | G | G | T | A | T | C | T | A | A | T | C | C | T | A | G |
| KJ461322.1 | Neuroptera | *Nymphes myrmeleonoides* | T | A | A | T | A | G | G | G | T | A | T | C | T | A | A | T | C | C | T | A | G |
| NC024825.1 | Neuroptera | *Nymphes myrmeleonoides* | T | A | A | T | A | G | G | G | T | A | T | C | T | A | A | T | C | C | T | A | G |
| KT425090.1 | Neuroptera | *Oliarces clara* | T | A | A | T | A | G | G | G | T | A | T | C | T | A | A | T | C | C | T | A | G |
| MN818868.1 | Neuroptera | *Osmylus atomatus* | T | A | A | T | A | G | G | G | T | A | T | C | T | A | A | T | C | C | T | A | G |
| MN818867.1 | Neuroptera | *Osmylus fulvicephalus* | T | A | A | T | A | G | G | G | T | A | T | C | T | A | A | T | C | C | T | A | G |
| MN617021.1 | Neuroptera | *Osmylus lucalatus* | T | A | A | T | A | G | G | G | T | A | T | C | T | A | A | T | C | C | T | A | G |
| MN818870.1 | Neuroptera | *Osmylus* sp. | T | A | A | T | A | G | G | G | T | A | T | C | T | A | A | T | C | C | T | A | G |
| MN818869.1 | Neuroptera | *Osmylus zheanus* | T | A | A | T | A | G | G | G | T | A | T | C | T | A | A | T | C | C | T | A | G |
| KY587202.1 | Neuroptera | *Parankylopteryx* sp. | T | A | A | T | A | G | G | G | T | A | T | C | T | A | A | T | C | C | T | A | G |
| FJ171325.1 | Neuroptera | *Polystoechotes punctatus* | T | A | A | T | A | G | G | G | T | A | T | C | T | A | A | T | C | C | T | A | G |
| NC011278.1 | Neuroptera | *Polystoechotes punctatus* | T | A | A | T | A | G | G | G | T | A | T | C | T | A | A | T | C | C | T | A | G |
| KT425082.1 | Neuroptera | *Psychopsis coelivaga* | T | A | A | T | A | G | G | G | T | A | T | C | T | A | A | T | C | C | T | A | G |

| | | | | | | | | | | | | | | | | | | | | | | | |
|---|---|---|---|---|---|---|---|---|---|---|---|---|---|---|---|---|---|---|---|---|---|---|---|
| KF626446.1 | Neuroptera | *Rapisma xizangense* | T | A | A | T | A | G | G | G | T | A | T | C | T | A | A | T | C | C | T | A | G |
| NC023362.1 | Neuroptera | *Rapisma xizangense* | T | A | A | T | A | G | G | G | T | A | T | C | T | A | A | T | C | C | T | A | G |
| KF626447.1 | Neuroptera | *Rapisma zayuanum* | T | A | A | T | A | G | G | G | T | A | T | C | T | A | A | T | C | C | T | A | G |
| NC023363.1 | Neuroptera | *Rapisma zayuanum* | T | A | A | T | A | G | G | G | T | A | T | C | T | A | A | T | C | C | T | A | G |
| KT425070.1 | Neuroptera | *Sisyra nigra* | T | A | A | T | A | G | G | G | T | A | T | C | T | A | A | T | C | C | T | A | G |
| KT425081.1 | Neuroptera | *Stenobiella* sp. | T | A | A | T | A | G | G | G | T | A | T | C | T | A | A | T | C | C | T | A | G |
| MH361300.1 | Neuroptera | *Suhpalacsa longialata* | T | A | A | T | A | G | G | G | T | A | T | C | T | A | A | T | C | C | T | A | G |
| NC039948.1 | Neuroptera | *Suhpalacsa longialata* | T | A | A | T | A | G | G | G | T | A | T | C | T | A | A | T | C | C | T | A | G |
| MK301247.1 | Neuroptera | *Suhpalacsa* sp. | T | A | A | T | A | G | G | G | T | A | T | C | T | A | A | T | C | C | T | A | G |
| MK408756.1 | Neuroptera | *Thaumatosmylus hainanus* | T | A | A | T | A | G | G | G | T | A | T | C | T | A | A | T | C | C | T | A | G |
| NC046578.1 | Neuroptera | *Thaumatosmylus hainanus* | T | A | A | T | A | G | G | G | T | A | T | C | T | A | A | T | C | C | T | A | G |
| KC515397.1 | Neuroptera | *Thyridosmylus langii* | T | A | A | T | A | G | G | G | T | A | T | C | T | A | A | T | C | C | T | A | G |
| NC021415.1 | Neuroptera | *Thyridosmylus* | T | A | A | T | A | G | G | G | T | A | T | C | T | A | A | T | C | C | T | A | G |

| | | | | | | | | | | | | | | | | | | | | | | |
|---|---|---|---|---|---|---|---|---|---|---|---|---|---|---|---|---|---|---|---|---|---|---|
| | | *langii* | | | | | | | | | | | | | | | | | | | | |
| KT425093.1 | Neuroptera | *Xanthostigma gobicola* | T | A | A | T | A | G | G | G | T | A | T | C | T | A | A | T | C | C | T | A | G |
| MH020192.1 | Coleoptera | *Abscondita anceyi* | T | A | A | T | A | G | G | G | T | A | T | C | T | A | A | T | C | C | T | A | G |
| NC039706.1 | Coleoptera | *Abscondita anceyi* | T | A | A | T | A | G | G | G | T | A | T | C | T | A | A | T | C | C | T | A | G |
| MK122952.1 | Coleoptera | *Abscondita chinensis* | T | A | A | T | A | G | G | G | T | A | T | C | T | A | A | T | C | C | T | A | G |
| MK292092.1 | Coleoptera | *Abscondita terminalis* | T | A | A | T | A | G | G | G | T | A | T | C | T | A | A | T | C | C | T | A | G |
| NC044776.1 | Coleoptera | *Abscondita terminalis* | T | A | A | T | A | G | G | G | T | A | T | C | T | A | A | T | C | C | T | A | G |
| KT876878.1 | Coleoptera | *Acilius sulcatus* | T | A | A | T | A | G | G | G | T | A | T | C | T | A | A | T | C | C | T | A | G |
| KX758085.1 | Coleoptera | *Aquatica ficta* | T | A | A | T | A | G | G | G | T | A | T | C | T | A | A | T | C | C | T | A | G |
| NC035060.1 | Coleoptera | *Aquatica ficta* | T | A | A | T | A | G | G | G | T | A | T | C | T | A | A | T | C | C | T | A | G |
| NC035755.1 | Coleoptera | *Aquatica lateralis* | T | A | A | T | A | G | G | G | T | A | T | C | T | A | A | T | C | C | T | A | G |
| KF667531.1 | Coleoptera | *Aquatica leii* | T | A | A | T | A | G | G | G | T | A | T | C | T | A | A | T | C | C | T | A | G |
| NC025276.1 | Coleoptera | *Aquatica leii* | T | A | A | T | A | G | G | G | T | A | T | C | T | A | A | T | C | C | T | A | G |
| KX758086.1 | Coleoptera | *Aquatica wuhana* | T | A | A | T | A | G | G | G | T | A | T | C | T | A | A | T | C | C | T | A | G |
| NC035061.1 | Coleoptera | *Aquatica wuhana* | T | A | A | T | A | G | G | G | T | A | T | C | T | A | A | T | C | C | T | A | G |
| KX229747.1 | Coleoptera | *Asymmetricata circumdata* | T | A | A | T | A | G | G | G | T | A | T | C | T | A | A | T | C | C | T | A | G |
| MK292113.1 | Coleoptera | *Asymmetricata* | T | A | G | T | A | G | G | G | T | A | T | C | T | A | A | T | C | C | T | A | G |

| | | | | | | | | | | | | | | | | | | | | | | |
|---|---|---|---|---|---|---|---|---|---|---|---|---|---|---|---|---|---|---|---|---|---|---|
| | | *circumdata* | | | | | | | | | | | | | | | | | | | | |
| NC032062.1 | Coleoptera | *Asymmetricata circumdata* | T | A | A | T | A | G | G | G | T | A | T | C | T | A | A | T | C | C | T | A | G |
| KJ922151.1 | Coleoptera | *Bicellonychia lividipennis* | T | A | A | T | A | G | G | G | T | A | T | C | T | A | A | T | C | C | T | A | G |
| NC030060.1 | Coleoptera | *Bicellonychia lividipennis* | T | A | A | T | A | G | G | G | T | A | T | C | T | A | A | T | C | C | T | A | G |
| KT876885.1 | Coleoptera | *Colymbetes fuscus* | T | A | A | T | A | G | G | G | T | A | T | C | T | A | A | T | C | C | T | A | G |
| MK292114.1 | Coleoptera | *Curtos bilineatus* | T | A | A | T | A | G | G | G | T | A | T | C | T | A | A | T | C | C | T | A | G |
| NC044789.1 | Coleoptera | *Curtos bilineatus* | T | A | A | T | A | G | G | G | T | A | T | C | T | A | A | T | C | C | T | A | G |
| MK609965.1 | Coleoptera | *Curtos costipennis* | T | A | A | T | A | G | G | G | T | A | T | C | T | A | A | T | C | C | T | A | G |
| MH651351.1 | Coleoptera | *Diaphanes citrinus* | T | A | G | T | A | G | G | G | T | A | T | C | T | A | A | T | C | C | T | A | G |
| MK292116.1 | Coleoptera | *Diaphanes mendax* | T | A | A | T | A | G | G | G | T | A | T | C | T | A | A | T | C | C | T | A | G |
| NC044791.1 | Coleoptera | *Diaphanes mendax* | T | A | A | T | A | G | G | G | T | A | T | C | T | A | A | T | C | C | T | A | G |
| MK292094.1 | Coleoptera | *Diaphanes nubilus* | T | A | A | T | A | G | G | G | T | A | T | C | T | A | A | T | C | C | T | A | G |
| NC044787.1 | Coleoptera | *Diaphanes nubilus* | T | A | A | T | A | G | G | G | T | A | T | C | T | A | A | T | C | C | T | A | G |
| MK292118.1 | Coleoptera | *Diaphanes pectinealis* | T | A | A | T | A | G | G | G | T | A | T | C | T | A | A | T | C | C | T | A | G |
| NC044793.1 | Coleoptera | *Diaphanes* | T | A | A | T | A | G | G | G | T | A | T | C | T | A | A | T | C | C | T | A | G |

| | | | | | | | | | | | | | | | | | | | | | | |
|---|---|---|---|---|---|---|---|---|---|---|---|---|---|---|---|---|---|---|---|---|---|---|
| | | *pectinealis* | | | | | | | | | | | | | | | | | | | | |
| MK292095.1 | Coleoptera | *Diaphanes* sp. | T | A | A | T | A | G | G | G | T | A | T | C | T | A | A | T | C | C | T | A | G |
| MG242622.1 | Coleoptera | *Ellychnia corrusca* | T | A | A | T | A | G | G | G | T | A | T | C | T | A | A | T | C | C | T | A | G |
| MK292112.1 | Coleoptera | *Emeia pseudosauteri* | T | A | A | T | A | G | G | G | T | A | T | C | T | A | A | T | C | C | T | A | G |
| NC044778.1 | Coleoptera | *Emeia pseudosauteri* | T | A | A | T | A | G | G | G | T | A | T | C | T | A | A | T | C | C | T | A | G |
| KT876897.1 | Coleoptera | *Hydroporus erythrocephalus* | T | A | A | T | A | G | G | G | T | A | T | C | T | A | A | T | C | C | T | A | G |
| KT876896.1 | Coleoptera | *Hydroporus obscurus* | T | A | A | T | A | G | G | G | T | A | T | C | T | A | A | T | C | C | T | A | G |
| KT876901.1 | Coleoptera | *Hygrotus confluens* | T | A | A | T | A | G | G | G | T | A | T | C | T | A | A | T | C | C | T | A | G |
| KT876900.1 | Coleoptera | *Hygrotus impressopunctatus* | T | A | A | T | A | G | G | G | T | A | T | C | T | A | A | T | C | C | T | A | G |
| KT876899.1 | Coleoptera | *Hygrotus inaequalis* | T | A | A | T | A | G | G | G | T | A | T | C | T | A | A | T | C | C | T | A | G |
| KX087299.1 | Coleoptera | *Hygrotus nigrolineatus* | T | A | A | T | A | G | G | G | T | A | T | C | T | A | A | T | C | C | T | A | G |
| MH427718.1 | Coleoptera | *Inflata indica* | T | A | A | T | A | G | G | G | T | A | T | C | T | A | A | T | C | C | T | A | G |
| NC039700.1 | Coleoptera | *Inflata indica* | T | A | A | T | A | G | G | G | T | A | T | C | T | A | A | T | C | C | T | A | G |
| MK292091.1 | Coleoptera | *Lamprigera* | T | A | A | T | A | G | G | G | T | A | T | C | T | A | A | T | C | C | T | A | G |

| | | | | | | | | | | | | | | | | | | | | | | |
|---|---|---|---|---|---|---|---|---|---|---|---|---|---|---|---|---|---|---|---|---|---|---|
| | | *yunnana* | | | | | | | | | | | | | | | | | | | | |
| NC044786.1 | Coleoptera | *Lamprigera* | T | A | A | T | A | G | G | G | T | A | T | C | T | A | A | T | C | C | T | A | G |
| | | *yunnana* | | | | | | | | | | | | | | | | | | | | |
| KX087302.1 | Coleoptera | *Lampyris noctiluca* | T | A | A | T | A | G | G | G | T | A | T | C | T | A | A | T | C | C | T | A | G |
| MG912994.1 | Coleoptera | *Limbodessus* | T | A | A | T | A | G | G | G | T | A | T | C | T | A | A | T | C | C | T | A | G |
| | | *palmulaoides* | | | | | | | | | | | | | | | | | | | | |
| NC037749.1 | Coleoptera | *Limbodessus* | T | A | A | T | A | G | G | G | T | A | T | C | T | A | A | T | C | C | T | A | G |
| | | *palmulaoides* | | | | | | | | | | | | | | | | | | | | |
| KT876902.1 | Coleoptera | *Liopterus* | T | A | A | T | A | G | G | G | T | A | T | C | T | A | A | T | C | C | T | A | G |
| | | *haemorrhoidalis* | | | | | | | | | | | | | | | | | | | | |
| NC022472.1 | Coleoptera | *Luciola cruciata* | T | A | A | T | A | G | G | G | T | A | T | C | T | A | A | T | C | C | T | A | G |
| MG770613.1 | Coleoptera | *Luciola curtithorax* | T | A | A | T | A | G | G | G | T | A | T | C | T | A | A | T | C | C | T | A | G |
| NC038225.1 | Coleoptera | *Luciola curtithorax* | T | A | A | T | A | G | G | G | T | A | T | C | T | A | A | T | C | C | T | A | G |
| KP313820.1 | Coleoptera | *Luciola substriata* | T | A | A | T | A | G | G | G | T | A | T | C | T | A | A | T | C | C | T | A | G |
| NC027176.1 | Coleoptera | *Luciola substriata* | T | A | A | T | A | G | G | G | T | A | T | C | T | A | A | T | C | C | T | A | G |
| FJ859901.1 | Coleoptera | *Macrogyrus* | T | A | A | T | A | G | G | G | T | A | T | C | T | A | A | T | C | C | T | A | G |
| | | *oblongus* | | | | | | | | | | | | | | | | | | | | |
| NC013249.1 | Coleoptera | *Macrogyrus* | T | A | A | T | A | G | G | G | T | A | T | C | T | A | A | T | C | C | T | A | G |
| | | *oblongus* | | | | | | | | | | | | | | | | | | | | |
| MG912995.1 | Coleoptera | *Paroster* | T | A | A | T | A | G | G | G | T | A | T | C | T | A | A | T | C | C | T | A | G |
| | | *macrosturtensis* | | | | | | | | | | | | | | | | | | | | |
| NC037750.1 | Coleoptera | *Paroster* | T | A | A | T | A | G | G | G | T | A | T | C | T | A | A | T | C | C | T | A | G |

| | | | | | | | | | | | | | | | | | | | | | | |
|---|---|---|---|---|---|---|---|---|---|---|---|---|---|---|---|---|---|---|---|---|---|---|
| | | *macrosturtensis* | | | | | | | | | | | | | | | | | | | | |
| MG912996.1 | Coleoptera | *Paroster mesosturtensis* | T | A | A | T | A | G | G | G | T | A | T | C | T | A | A | T | C | C | T | A | G |
| NC037751.1 | Coleoptera | *Paroster mesosturtensis* | T | A | A | T | A | G | G | G | T | A | T | C | T | A | A | T | C | C | T | A | G |
| MG912997.1 | Coleoptera | *Paroster microsturtensis* | T | A | A | T | A | G | G | G | T | A | T | C | T | A | A | T | C | C | T | A | G |
| NC037752.1 | Coleoptera | *Paroster microsturtensis* | T | A | A | T | A | G | G | G | T | A | T | C | T | A | A | T | C | C | T | A | G |
| KY778696.1 | Coleoptera | *Photinus pyralis* | T | A | A | T | A | G | G | G | T | A | T | C | T | A | A | T | C | C | T | A | G |
| MF686051.1 | Coleoptera | *Pteroptyx maipo* | T | A | G | T | A | G | G | G | T | A | T | C | T | A | A | T | C | C | T | A | G |
| NC036353.1 | Coleoptera | *Pteroptyx maipo* | T | A | G | T | A | G | G | G | T | A | T | C | T | A | A | T | C | C | T | A | G |
| MK292115.1 | Coleoptera | *Pyrocoelia praetexta* | T | A | A | T | A | G | G | G | T | A | T | C | T | A | A | T | C | C | T | A | G |
| NC044790.1 | Coleoptera | *Pyrocoelia praetexta* | T | A | A | T | A | G | G | G | T | A | T | C | T | A | A | T | C | C | T | A | G |
| AF452048.1 | Coleoptera | *Pyrocoelia rufa* | T | A | A | T | A | G | G | G | T | A | T | C | T | A | A | T | C | C | T | A | G |
| NC003970.1 | Coleoptera | *Pyrocoelia rufa* | T | A | A | T | A | G | G | G | T | A | T | C | T | A | A | T | C | C | T | A | G |
| MK292117.1 | Coleoptera | *Pyrocoelia thibetana* | T | A | A | T | A | G | G | G | T | A | T | C | T | A | A | T | C | C | T | A | G |
| NC044792.1 | Coleoptera | *Pyrocoelia thibetana* | T | A | A | T | A | G | G | G | T | A | T | C | T | A | A | T | C | C | T | A | G |

| | | | | | | | | | | | | | | | | | | | | | | |
|---|---|---|---|---|---|---|---|---|---|---|---|---|---|---|---|---|---|---|---|---|---|---|
| MK292111.1 | Coleoptera | *Vesta saturnalis* | T | A | A | T | A | G | G | G | T | A | T | C | T | A | A | T | C | C | T | A | G |
| NC044788.1 | Coleoptera | *Vesta saturnalis* | T | A | A | T | A | G | G | G | T | A | T | C | T | A | A | T | C | C | T | A | G |
| NC030249.1 | Diptera | *Anopheles bellator* | T | A | A | T | A | G | G | G | T | A | T | C | T | A | A | T | C | C | T | A | G |
| NC030717.1 | Diptera | *Anopheles janconnae* | T | A | A | T | A | G | G | G | T | A | T | C | T | A | A | T | C | C | T | A | G |
| MF381592.1 | Diptera | *Anopheles marajoara* | T | A | A | T | A | G | G | G | T | A | T | C | T | A | A | T | C | C | T | A | G |
| HQ335345.1 | Diptera | *Anopheles oryzalimnetes* | T | A | A | T | A | G | G | G | T | A | T | C | T | A | A | T | C | C | T | A | G |
| MF381722.1 | Diptera | *Culex bilineatus* | T | A | A | T | A | G | G | G | T | A | T | C | T | A | A | T | C | C | T | A | G |
| MF381718.1 | Diptera | *Culex lygrus* | T | A | A | T | A | G | G | G | T | A | T | C | T | A | A | T | C | C | T | A | G |
| MN626442.1 | Diptera | *Ochlerotatus taeniorhynchus* | T | A | A | T | A | G | G | G | T | A | T | C | T | A | A | T | C | C | T | A | G |
| KP721463.1 | Diptera | *Ochlerotatus vigilax* | T | A | A | T | A | G | G | G | T | A | T | C | T | A | A | T | C | C | T | A | G |
| MF957171.1 | Diptera | *Sabethes belisarioi* | T | A | A | T | A | G | G | G | T | A | T | C | T | A | A | T | C | C | T | A | G |
| KP793690.1 | Diptera | *Simulium aureohirtum* | T | A | A | T | A | G | G | G | T | A | T | C | T | A | A | T | C | C | T | A | G |
| NC029753.1 | Diptera | *Simulium aureohirtum* | T | A | A | T | A | G | G | G | T | A | T | C | T | A | A | T | C | C | T | A | G |
| MH729190.1 | Diptera | *Simulium* | T | A | A | T | A | G | G | G | T | A | T | C | T | A | A | T | C | C | T | A | G |

| | | | | | | | | | | | | | | | | | | | | | | | |
|---|---|---|---|---|---|---|---|---|---|---|---|---|---|---|---|---|---|---|---|---|---|---|---|
| | | *maculatum* | | | | | | | | | | | | | | | | | | | | | |
| NC040120.1 | Diptera | *Simulium maculatum* | T | A | A | T | A | G | G | G | T | A | T | C | T | A | A | T | C | C | T | A | G |
| KU252587.1 | Diptera | *Simulium variegatum* | T | A | A | T | A | G | G | G | T | A | T | C | T | A | A | T | C | C | T | A | G |
| NC033348.1 | Diptera | *Simulium variegatum* | T | A | A | T | A | G | G | G | T | A | T | C | T | A | A | T | C | C | T | A | G |
| JN861743.1 | Diptera | *Tipula abdominalis* | T | A | A | T | A | G | G | G | T | A | T | C | T | A | A | T | C | C | T | A | G |
| KT970065.1 | Diptera | *Tipula cockerelliana* | T | A | A | T | A | G | G | G | T | A | T | C | T | A | A | T | C | C | T | A | G |
| NC030520.1 | Diptera | *Tipula cockerelliana* | T | A | A | T | A | G | G | G | T | A | T | C | T | A | A | T | C | C | T | A | G |
| MT410842.1 | Diptera | *Tipula fascipennis* | T | A | A | T | A | G | G | G | T | A | T | C | T | A | A | T | C | C | T | A | G |
| NC050319.1 | Diptera | *Tipula fascipennis* | T | A | A | T | A | G | G | G | T | A | T | C | T | A | A | T | C | C | T | A | G |
| NC044661.1 | Diptera | *Trichoprosopon pallidiventer* | T | A | A | T | A | G | G | G | T | A | T | C | T | A | A | T | C | C | T | A | G |
| MK575491.1 | Diptera | *Uranotaenia geometrica* | T | A | A | T | A | G | G | G | T | A | T | C | T | A | A | T | C | C | T | A | G |
| MK575492.1 | Diptera | *Wyeomyia confusa* | T | A | A | T | A | G | G | G | T | A | T | C | T | A | A | T | C | C | T | A | G |
| | | A | 0.0 | 100.0 | 96.7 | 0.0 | 100.0 | 0.0 | 0.0 | 0.0 | 0.0 | 100.0 | 0.0 | 0.0 | 0.0 | 100.0 | 100.0 | 0.0 | 0.0 | 0.0 | 0.0 | 100.0 | 0.0 |

| | | | | | | | | | | | | | | | | | | | | | |
|---|---|---|---|---|---|---|---|---|---|---|---|---|---|---|---|---|---|---|---|---|---|
| T | 100.0 | 0.0 | 0.0 | 100.0 | 0.0 | 0.0 | 0.0 | 0.0 | 100.0 | 0.0 | 100.0 | 0.0 | 100.0 | 0.0 | 0.0 | 100.0 | 0.0 | 0.0 | 100.0 | 0.0 | 0.0 |
| G | 0.0 | 0.0 | 3.3 | 0.0 | 0.0 | 100.0 | 100.0 | 100.0 | 0.0 | 0.0 | 0.0 | 0.0 | 0.0 | 0.0 | 0.0 | 0.0 | 0.0 | 0.0 | 0.0 | 0.0 | 100.0 |
| C | 0.0 | 0.0 | 0.0 | 0.0 | 0.0 | 0.0 | 0.0 | 0.0 | 0.0 | 0.0 | 0.0 | 100.0 | 0.0 | 0.0 | 0.0 | 0.0 | 100.0 | 100.0 | 0.0 | 0.0 | 0.0 |

Table S9. Nucleotide sequences of universal primer (MtInsects-12S_cad_F) and homologous region of 31 sequences of trichopteran insects used for development in this study.

| | | | | | | | | | | | | | | | | | | | | | | | | |
|---|---|---|---|---|---|---|---|---|---|---|---|---|---|---|---|---|---|---|---|---|---|---|---|---|
| MtInsects-12S_cad_F | | | | T | T | G | K | G | C | C | A | G | C | A | R | T | Y | G | C | G | G | T | W | A |
| | | | A | 0 | 0 | 0 | 0 | 0 | 0 | 0 | 31 | 0 | 0 | 29 | 24 | 0 | 0 | 0 | 0 | 0 | 2 | 0 | 0 | 31 |
| | | | T | 29 | 31 | 0 | 29 | 0 | 2 | 0 | 0 | 0 | 0 | 0 | 0 | 31 | 23 | 0 | 0 | 0 | 0 | 29 | 31 | 0 |
| | | | G | 0 | 0 | 31 | 2 | 31 | 0 | 0 | 0 | 31 | 0 | 0 | 7 | 0 | 0 | 31 | 0 | 31 | 29 | 0 | 0 | 0 |
| | | | C | 2 | 0 | 0 | 0 | 0 | 29 | 31 | 0 | 0 | 31 | 0 | 0 | 0 | 8 | 0 | 31 | 0 | 0 | 0 | 0 | 0 |
| GenBank Acc. No. | Order | Species | | | | | | | | | | | | | | | | | | | | | | |
| KF756944.1 | Trichoptera | *Apatania* sp. | | T | T | G | T | G | C | C | A | G | C | A | G | T | T | G | C | G | G | T | T | A |
| MT410856.1 | Trichoptera | *Beraea pullata* | | T | T | G | T | G | C | C | A | G | C | A | A | T | T | G | C | G | G | T | T | A |
| MG669126.1 | Trichoptera | *Cheumatopsyche analis* | | T | T | G | T | G | C | C | A | G | C | A | A | T | T | G | C | G | G | T | T | A |
| NC_036955.1 | Trichoptera | *Cheumatopsyche analis* | | T | T | G | T | G | C | C | A | G | C | A | A | T | T | G | C | G | G | T | T | A |
| KX385010.1 | Trichoptera | *Cheumatopsyche brevilineata* | | T | T | G | G | G | C | C | A | G | C | A | A | T | T | G | C | G | G | T | T | A |
| NC_043769.1 | Trichoptera | *Cheumatopsyche brevilineata* | | T | T | G | G | G | C | C | A | G | C | A | A | T | T | G | C | G | G | T | T | A |
| MG669125.1 | Trichoptera | *Cheumatopsyche campyla* | | T | T | G | T | G | C | C | A | G | C | A | A | T | T | G | C | G | G | T | T | A |
| NC_036954.1 | Trichoptera | *Cheumatopsyche campyla* | | T | T | G | T | G | C | C | A | G | C | A | A | T | T | G | C | G | G | T | T | A |

| | | | | | | | | | | | | | | | | | | | | | | | |
|---|---|---|---|---|---|---|---|---|---|---|---|---|---|---|---|---|---|---|---|---|---|---|---|
| MG669123.1 | Trichoptera | *Cheumatopsyche speciosa* | T | T | G | T | G | C | C | A | G | C | A | A | T | T | G | C | G | G | T | T | A |
| NC_036952.1 | Trichoptera | *Cheumatopsyche speciosa* | T | T | G | T | G | C | C | A | G | C | A | A | T | T | G | C | G | G | T | T | A |
| KF717094.1 | Trichoptera | *Eubasilissa regina* | T | T | G | T | G | C | C | A | G | C | A | A | T | C | G | C | G | G | T | T | A |
| KF756943.1 | Trichoptera | *Eubasilissa regina* | T | T | G | T | G | C | C | A | G | C | A | A | T | C | G | C | G | G | T | T | A |
| NC_023374.1 | Trichoptera | *Eubasilissa regina* | T | T | G | T | G | C | C | A | G | C | A | A | T | C | G | C | G | G | T | T | A |
| KX385011.1 | Trichoptera | *Hydatophylax nigrovittatus* | T | T | G | T | G | C | C | A | G | C | A | G | T | C | G | C | G | G | T | T | A |
| NC_043770.1 | Trichoptera | *Hydatophylax nigrovittatus* | T | T | G | T | G | C | C | A | G | C | A | G | T | C | G | C | G | G | T | T | A |
| KF717095.1 | Trichoptera | *Hydromanicus wulaianus* | T | T | G | T | G | C | C | A | G | C | A | A | T | T | G | C | G | G | T | T | A |
| NC_036156.1 | Trichoptera | *Hydromanicus wulaianus* | T | T | G | T | G | C | C | A | G | C | A | A | T | T | G | C | G | G | T | T | A |
| MT628580.1 | Trichoptera | *Hydropsyche angustipennis* | T | T | G | T | G | C | C | A | G | C | A | A | T | T | G | C | G | G | T | T | A |
| MG669122.1 | Trichoptera | *Hydropsyche orris* | T | T | G | T | G | C | C | A | G | C | A | A | T | T | G | C | G | G | T | T | A |

| | | | | | | | | | | | | | | | | | | | | | | | |
|---|---|---|---|---|---|---|---|---|---|---|---|---|---|---|---|---|---|---|---|---|---|---|---|
| NC_036951.1 | Trichoptera | *Hydropsyche orris* | T | T | G | T | G | C | C | A | G | C | A | A | T | T | G | C | G | G | T | T | A |
| MG669121.1 | Trichoptera | *Hydropsyche simulans* | T | T | G | T | G | C | C | A | G | C | A | A | T | T | G | C | G | G | T | T | A |
| NC_036950.1 | Trichoptera | *Hydropsyche simulans* | T | T | G | T | G | C | C | A | G | C | A | A | T | T | G | C | G | G | T | T | A |
| MK077681.1 | Trichoptera | *Limnephilus hyalinus* | T | T | G | T | G | C | C | A | G | C | A | G | T | C | G | C | G | G | T | T | A |
| NC_044710.1 | Trichoptera | *Limnephilus hyalinus* | T | T | G | T | G | C | C | A | G | C | A | G | T | C | G | C | G | G | T | T | A |
| MG980616.1 | Trichoptera | *Phryganea cinerea* | T | T | G | T | G | C | C | A | G | C | A | A | T | T | G | C | G | G | T | T | A |
| NC_039714.1 | Trichoptera | *Phryganea cinerea* | T | T | G | T | G | C | C | A | G | C | A | A | T | T | G | C | G | G | T | T | A |
| KX385012.1 | Trichoptera | *Phryganopsyche latipennis* | C | T | G | T | G | T | C | A | G | C | ? | G | T | T | G | C | G | A | ? | T | A |
| NC_43771.1 | Trichoptera | *Phryganopsyche latipennis* | C | T | G | T | G | T | C | A | G | C | ? | G | T | T | G | C | G | A | ? | T | A |
| MG669124.1 | Trichoptera | *Potamyia flava* | T | T | G | T | G | C | C | A | G | C | A | A | T | T | G | C | G | G | T | T | A |
| NC_036953.1 | Trichoptera | *Potamyia flava* | T | T | G | T | G | C | C | A | G | C | A | A | T | T | G | C | G | G | T | T | A |
| KP455291.1 | Trichoptera | *Thremma gallicum* | T | T | G | T | G | C | C | A | G | C | A | A | T | C | G | C | G | G | T | T | A |

| | | | | | | | | | | | | | | | | | | | | | |
|---|---|---|---|---|---|---|---|---|---|---|---|---|---|---|---|---|---|---|---|---|---|
| A | 0.0 | 0.0 | 0.0 | 0.0 | 0.0 | 0.0 | 0.0 | 100.0 | 0.0 | 0.0 | 100.0 | 77.4 | 0.0 | 0.0 | 0.0 | 0.0 | 0.0 | 6.5 | 0.0 | 0.0 | 100.0 |
| T | 93.5 | 100.0 | 0.0 | 93.5 | 0.0 | 6.5 | 0.0 | 0.0 | 0.0 | 0.0 | 0.0 | 0.0 | 100.0 | 74.2 | 0.0 | 0.0 | 0.0 | 0.0 | 100.0 | 100.0 | 0.0 |
| G | 0.0 | 0.0 | 100.0 | 6.5 | 100.0 | 0.0 | 0.0 | 0.0 | 100.0 | 0.0 | 0.0 | 22.6 | 0.0 | 0.0 | 100.0 | 0.0 | 100.0 | 93.5 | 0.0 | 0.0 | 0.0 |
| C | 6.5 | 0.0 | 0.0 | 0.0 | 0.0 | 93.5 | 100.0 | 0.0 | 0.0 | 100.0 | 0.0 | 0.0 | 0.0 | 25.8 | 0.0 | 100.0 | 0.0 | 0.0 | 0.0 | 0.0 | 0.0 |

Table S10. Nucleotide sequences of universal primer (MtInsects-12S_cad_R) and homologous region of 31 sequences of trichopteran insects used for development in this study.

| | | | | | | | | | | | | | | | | | | | | | | | |
|---|---|---|---|---|---|---|---|---|---|---|---|---|---|---|---|---|---|---|---|---|---|---|---|
| MtInsects-12S_cad_R | | | | W | A | T | A | R | T | R | G | R | G | T | A | T | C | T | A | A | T | Y | C |
| | | | A | 6 | 31 | 0 | 31 | 29 | 0 | 29 | 2 | 3 | 0 | 0 | 31 | 0 | 0 | 0 | 31 | 31 | 0 | 0 | 0 |
| | | | T | 25 | 0 | 31 | 0 | 0 | 31 | 0 | 0 | 0 | 0 | 31 | 0 | 29 | 0 | 31 | 0 | 0 | 31 | 1 | 0 |
| | | | G | 0 | 0 | 0 | 0 | 2 | 0 | 2 | 29 | 28 | 29 | 0 | 0 | 0 | 0 | 0 | 0 | 0 | 0 | 0 | 0 |
| | | | C | 0 | 0 | 0 | 0 | 0 | 0 | 0 | 0 | 0 | 2 | 0 | 0 | 2 | 31 | 0 | 0 | 0 | 0 | 30 | 31 |
| GenBank Acc. No. | Order | Species | | | | | | | | | | | | | | | | | | | | | |
| KF756944.1 | Trichoptera | *Apatania* sp. | | T | A | T | A | A | T | A | G | G | G | T | A | T | C | T | A | A | T | C | C |
| MT410856.1 | Trichoptera | *Beraea pullata* | | A | A | T | A | A | T | A | G | A | G | T | A | T | C | T | A | A | T | T | C |
| MG669126.1 | Trichoptera | *Cheumatopsyche analis* | | T | A | T | A | A | T | A | G | G | G | T | A | T | C | T | A | A | T | C | C |
| NC036955.1 | Trichoptera | *Cheumatopsyche analis* | | T | A | T | A | A | T | A | G | G | G | T | A | T | C | T | A | A | T | C | C |
| KX385010.1 | Trichoptera | *Cheumatopsyche brevilineata* | | A | A | T | A | A | T | A | G | G | G | T | A | T | C | T | A | A | T | C | C |
| NC043769.1 | Trichoptera | *Cheumatopsyche brevilineata* | | A | A | T | A | A | T | A | G | G | G | T | A | T | C | T | A | A | T | C | C |
| MG669125.1 | Trichoptera | *Cheumatopsyche campyla* | | T | A | T | A | A | T | A | G | G | G | T | A | T | C | T | A | A | T | C | C |
| NC036954.1 | Trichoptera | *Cheumatopsyche campyla* | | T | A | T | A | A | T | A | G | G | G | T | A | T | C | T | A | A | T | C | C |
| MG669123.1 | Trichoptera | *Cheumatopsyche speciosa* | | T | A | T | A | A | T | A | G | G | G | T | A | T | C | T | A | A | T | C | C |

| | | | | | | | | | | | | | | | | | | | | | | |
|---|---|---|---|---|---|---|---|---|---|---|---|---|---|---|---|---|---|---|---|---|---|---|
| NC036952.1 | Trichoptera | *Cheumatopsyche speciosa* | T | A | T | A | A | T | A | G | G | G | T | A | T | C | T | A | A | T | C | C |
| KF717094.1 | Trichoptera | *Eubasilissa regina* | T | A | T | A | A | T | A | G | G | G | T | A | T | C | T | A | A | T | C | C |
| KF756943.1 | Trichoptera | *Eubasilissa regina* | A | A | T | A | A | T | A | G | G | G | T | A | T | C | T | A | A | T | C | C |
| NC023374.1 | Trichoptera | *Eubasilissa regina T* | A | A | T | A | A | T | A | G | G | G | T | A | T | C | T | A | A | T | C | C |
| KX385011.1 | Trichoptera | *Hydatophylax nigrovittatus* | T | A | T | A | A | T | A | G | G | G | T | A | T | C | T | A | A | T | C | C |
| NC043770.1 | Trichoptera | *Hydatophylax nigrovittatus* | T | A | T | A | A | T | A | G | G | G | T | A | T | C | T | A | A | T | C | C |
| KF717095.1 | Trichoptera | *Hydromanicus wulaianus* | T | A | T | A | A | T | A | G | G | G | T | A | T | C | T | A | A | T | C | C |
| NC036156.1 | Trichoptera | *Hydromanicus wulaianus* | T | A | T | A | A | T | A | G | G | G | T | A | T | C | T | A | A | T | C | C |
| MT628580.1 | Trichoptera | *Hydropsyche angustipennis* | A | A | T | A | A | T | A | G | G | G | T | A | T | C | T | A | A | T | C | C |
| MG669122.1 | Trichoptera | *Hydropsyche orris* | T | A | T | A | A | T | A | G | G | G | T | A | T | C | T | A | A | T | C | C |
| NC036951.1 | Trichoptera | *Hydropsyche orris* | T | A | T | A | A | T | A | G | G | G | T | A | T | C | T | A | A | T | C | C |
| MG669121.1 | Trichoptera | *Hydropsyche simulans* | T | A | T | A | A | T | A | G | G | G | T | A | T | C | T | A | A | T | C | C |
| NC036950.1 | Trichoptera | *Hydropsyche simulans* | T | A | T | A | A | T | A | G | G | G | T | A | T | C | T | A | A | T | C | C |
| MK077681.1 | Trichoptera | *Limnephilus hyalinus* | T | A | T | A | G | T | G | G | G | G | T | A | T | C | T | A | A | T | C | C |
| NC044710.1 | Trichoptera | *Limnephilus hyalinus* | T | A | T | A | G | T | G | G | G | G | T | A | T | C | T | A | A | T | C | C |
| MG980616.1 | Trichoptera | *Phryganea cinerea* | T | A | T | A | A | T | A | G | G | G | T | A | T | C | T | A | A | T | C | C |

| | | | | | | | | | | | | | | | | | | | | | | | |
|---|---|---|---|---|---|---|---|---|---|---|---|---|---|---|---|---|---|---|---|---|---|---|---|
| NC039714.1 | Trichoptera | *Phryganea cinerea* | | T | A | T | A | A | T | A | G | G | G | T | A | T | C | T | A | A | T | C | C |
| KX385012.1 | Trichoptera | *Phryganopsyche latipennis* | | T | A | T | A | A | T | A | A | A | C | T | A | C | C | T | A | A | T | C | C |
| NC043771.1 | Trichoptera | *Phryganopsyche latipennis* | | T | A | T | A | A | T | A | A | A | C | T | A | C | C | T | A | A | T | C | C |
| MG669124.1 | Trichoptera | *Potamyia flava* | | T | A | T | A | A | T | A | G | G | G | T | A | T | C | T | A | A | T | C | C |
| NC036953.1 | Trichoptera | *Potamyia flava* | | T | A | T | A | A | T | A | G | G | G | T | A | T | C | T | A | A | T | C | C |
| KP455291.1 | Trichoptera | *Thremma gallicum* | | T | A | T | A | A | T | A | G | G | G | T | A | T | C | T | A | A | T | C | C |
| | | | A | 19.4 | 100.0 | 0.0 | 100.0 | 93.5 | 0.0 | 93.5 | 6.5 | 9.7 | 0.0 | 0.0 | 100.0 | 0.0 | 0.0 | 0.0 | 100.0 | 100.0 | 0.0 | 0.0 | 0.0 |
| | | | T | 80.6 | 0.0 | 100.0 | 0.0 | 0.0 | 100.0 | 0.0 | 0.0 | 0.0 | 0.0 | 100.0 | 0.0 | 93.5 | 0.0 | 100.0 | 0.0 | 0.0 | 100.0 | 3.2 | 0.0 |
| | | | G | 0.0 | 0.0 | 0.0 | 0.0 | 6.5 | 0.0 | 6.5 | 93.5 | 90.3 | 93.5 | 0.0 | 0.0 | 0.0 | 0.0 | 0.0 | 0.0 | 0.0 | 0.0 | 0.0 | 0.0 |
| | | | C | 0.0 | 0.0 | 0.0 | 0.0 | 0.0 | 0.0 | 0.0 | 0.0 | 0.0 | 6.5 | 0.0 | 0.0 | 6.5 | 100.0 | 0.0 | 0.0 | 0.0 | 0.0 | 96.8 | 100.0 |

Table S11. Nucleotide sequences of universal primer (AQdb-12S_Hem_R) and homologous region of 265 sequences of insects used for development in this study.

| | | | | | | | | | | | | | | | | | | | | | | |
|---|---|---|---|---|---|---|---|---|---|---|---|---|---|---|---|---|---|---|---|---|---|---|
| AQdb-12S_Hem_R | | | | C | T | A | C | T | W | T | G | T | T | A | C | G | A | C | T | T | R | T |
| | | | A | 0 | 0 | 265 | 0 | 0 | 190 | 0 | 0 | 0 | 0 | 265 | 0 | 0 | 265 | 0 | 0 | 0 | 265 | 0 |
| | | | T | 2 | 265 | 0 | 0 | 265 | 73 | 265 | 0 | 265 | 265 | 0 | 0 | 0 | 0 | 0 | 265 | 265 | 0 | 265 |
| | | | G | 0 | 0 | 0 | 0 | 0 | 2 | 0 | 265 | 0 | 0 | 0 | 0 | 265 | 0 | 0 | 0 | 0 | 0 | 0 |
| | | | C | 263 | 0 | 0 | 265 | 0 | 0 | 0 | 0 | 0 | 0 | 0 | 265 | 0 | 0 | 265 | 0 | 0 | 0 | 0 |
| GenBank Acc. No. | Order | Species | | | | | | | | | | | | | | | | | | | | |
| GQ502451.1 | Ephemeroptera | *Caenis* sp. | | C | T | A | C | T | A | T | G | T | T | A | C | G | A | C | T | T | A | T |
| MG910499.1 | Ephemeroptera | *Caenis* sp. | | C | T | A | C | T | A | T | G | T | T | A | C | G | A | C | T | T | A | T |
| MN356096.1 | Ephemeroptera | *Caenis* sp. | | C | T | A | C | T | A | T | G | T | T | A | C | G | A | C | T | T | A | T |
| MN807287.1 | Ephemeroptera | *Choroterpides apiculata* | | C | T | A | C | T | A | T | G | T | T | A | C | G | A | C | T | T | A | T |
| MG870104.1 | Ephemeroptera | *Epeorus herklotsi* | | C | T | A | C | T | A | T | G | T | T | A | C | G | A | C | T | T | A | T |
| MH752075.1 | Ephemeroptera | *Epeorus herklotsi* | | C | T | A | C | T | A | T | G | T | T | A | C | G | A | C | T | T | A | T |
| NC039612.1 | Ephemeroptera | *Epeorus herklotsi* | | C | T | A | C | T | A | T | G | T | T | A | C | G | A | C | T | T | A | T |
| KJ493406.1 | Ephemeroptera | *Epeorus* sp. | | C | T | A | C | T | A | T | G | T | T | A | C | G | A | C | T | T | A | T |
| KM244708.1 | Ephemeroptera | *Epeorus* sp. | | C | T | A | C | T | A | T | G | T | T | A | C | G | A | C | T | T | A | T |
| MK951674.1 | Ephemeroptera | *Epeorus* sp. | | C | T | A | C | T | T | T | G | T | T | A | C | G | A | C | T | T | A | T |
| EU591678.1 | Ephemeroptera | *Ephemera orientalis* | | C | T | A | C | T | A | T | G | T | T | A | C | G | A | C | T | T | A | T |
| NC012645.1 | Ephemeroptera | *Ephemera orientalis* | | C | T | A | C | T | A | T | G | T | T | A | C | G | A | C | T | T | A | T |
| MK951659.1 | Ephemeroptera | *Ephemera* sp. | | C | T | A | C | T | A | T | G | T | T | A | C | G | A | C | T | T | A | T |

| | | | | | | | | | | | | | | | | | | | | |
|---|---|---|---|---|---|---|---|---|---|---|---|---|---|---|---|---|---|---|---|---|
| MT274127.1 | Ephemeroptera | *Ephemera* sp. | C | T | A | C | T | A | T | G | T | T | A | C | G | A | C | T | T | A | T |
| NC050279.1 | Ephemeroptera | *Ephemera* sp. | C | T | A | C | T | A | T | G | T | T | A | C | G | A | C | T | T | A | T |
| GU936203.1 | Ephemeroptera | *Habrophlebiodes zijinensis* | C | T | A | C | T | T | T | G | T | T | A | C | G | A | C | T | T | A | T |
| HM143892.1 | Ephemeroptera | *Isonychia ignota* | C | T | A | C | T | A | T | G | T | T | A | C | G | A | C | T | T | A | T |
| MH119135.1 | Ephemeroptera | *Isonychia kiangsinensis* | C | T | A | C | T | A | T | G | T | T | A | C | G | A | C | T | T | A | T |
| NC042163.1 | Ephemeroptera | *Isonychia kiangsinensis* | C | T | A | C | T | A | T | G | T | T | A | C | G | A | C | T | T | A | T |
| MK951658.1 | Ephemeroptera | *Isonychia* sp. | C | T | A | C | T | A | T | G | T | T | A | C | G | A | C | T | T | A | T |
| HM004123.1 | Ephemeroptera | *Paegniodes cupulatus* | C | T | A | C | T | A | T | G | T | T | A | C | G | A | C | T | T | A | T |
| MK951673.1 | Ephemeroptera | *Parafronurus* sp. | C | T | A | C | T | T | T | G | T | T | A | C | G | A | C | T | T | A | T |
| EU349015.1 | Ephemeroptera | *Parafronurus youi* | C | T | A | C | T | T | T | G | T | T | A | C | G | A | C | T | T | A | T |
| NC011359.1 | Ephemeroptera | *Parafronurus youi* | C | T | A | C | T | T | T | G | T | T | A | C | G | A | C | T | T | A | T |
| KM244674.1 | Ephemeroptera | *Potamanthus* sp. | C | T | A | C | T | A | T | G | T | T | A | C | G | A | C | T | T | A | T |
| MK951670.1 | Ephemeroptera | *Rhithrogena* sp. | C | T | A | C | T | T | T | G | T | T | A | C | G | A | C | T | T | A | T |
| MT274128.1 | Ephemeroptera | *Serratella* sp. | C | T | A | C | T | G | T | G | T | T | A | C | G | A | C | T | T | A | T |
| MT274129.1 | Ephemeroptera | *Serratella* sp. | C | T | A | C | T | A | T | G | T | T | A | C | G | A | C | T | T | A | T |
| NC050280.1 | Ephemeroptera | *Serratella* sp. | C | T | A | C | T | G | T | G | T | T | A | C | G | A | C | T | T | A | T |
| NC050281.1 | Ephemeroptera | *Serratella* sp. | C | T | A | C | T | A | T | G | T | T | A | C | G | A | C | T | T | A | T |
| MT274130.1 | Ephemeroptera | *Serratella zapekinae* | C | T | A | C | T | A | T | G | T | T | A | C | G | A | C | T | T | A | T |

| | | | | | | | | | | | | | | | | | | | | |
|---|---|---|---|---|---|---|---|---|---|---|---|---|---|---|---|---|---|---|---|---|
| NC050282.1 | Ephemeroptera | *Serratella zapekinae* | C | T | A | C | T | A | T | G | T | T | A | C | G | A | C | T | T | A | T |
| FJ606783.1 | Ephemeroptera | *Siphlonurus immanis* | C | T | A | C | T | A | T | G | T | T | A | C | G | A | C | T | T | A | T |
| NC013822.1 | Ephemeroptera | *Siphlonurus immanis* | C | T | A | C | T | A | T | G | T | T | A | C | G | A | C | T | T | A | T |
| KM244684.1 | Ephemeroptera | *Siphlonurus* sp. | C | T | A | C | T | A | T | G | T | T | A | C | G | A | C | T | T | A | T |
| HQ875717.1 | Ephemeroptera | *Siphluriscus chinensis* | C | T | A | C | T | A | T | G | T | T | A | C | G | A | C | T | T | A | T |
| MT274131.1 | Ephemeroptera | *Torleya grandiforceps* | C | T | A | C | T | A | T | G | T | T | A | C | G | A | C | T | T | A | T |
| NC050283.1 | Ephemeroptera | *Torleya grandiforceps* | C | T | A | C | T | A | T | G | T | T | A | C | G | A | C | T | T | A | T |
| MT274132.1 | Ephemeroptera | *Torleya nepalica* | C | T | A | C | T | A | T | G | T | T | A | C | G | A | C | T | T | A | T |
| NC050284.1 | Ephemeroptera | *Torleya nepalica* | C | T | A | C | T | A | T | G | T | T | A | C | G | A | C | T | T | A | T |
| HM067837.1 | Ephemeroptera | *Vietnamella dabieshanensis* | C | T | A | C | T | A | T | G | T | T | A | C | G | A | C | T | T | A | T |
| KM244655.1 | Ephemeroptera | *Vietnamella* sp. | C | T | A | C | T | A | T | G | T | T | A | C | G | A | C | T | T | A | T |
| MH751435.1 | Odonata | *Acisoma panorpoides* | C | T | A | C | T | A | T | G | T | T | A | C | G | A | C | T | T | A | T |
| NC042689.1 | Odonata | *Acisoma panorpoides* | C | T | A | C | T | A | T | G | T | T | A | C | G | A | C | T | T | A | T |
| KX161841.1 | Odonata | *Anax imperator* | C | T | A | C | T | A | T | G | T | T | A | C | G | A | C | T | T | A | T |
| NC031821.1 | Odonata | *Anax imperator* | C | T | A | C | T | A | T | G | T | T | A | C | G | A | C | T | T | A | T |
| KP233805.1 | Odonata | *Atrocalopteryx atrata* | C | T | A | C | T | A | T | G | T | T | A | C | G | A | C | T | T | A | T |
| NC027181.1 | Odonata | *Atrocalopteryx atrata* | C | T | A | C | T | A | T | G | T | T | A | C | G | A | C | T | T | A | T |
| KM658172.1 | Odonata | *Brachythemis* | C | T | A | C | T | A | T | G | T | T | A | C | G | A | C | T | T | A | T |

| | | | | | | | | | | | | | | | | | | | | | |
|---|---|---|---|---|---|---|---|---|---|---|---|---|---|---|---|---|---|---|---|---|---|
| | | *contaminata* | | | | | | | | | | | | | | | | | | | |
| NC026305.1 | Odonata | *Brachythemis contaminata* | C | T | A | C | T | A | T | G | T | T | A | C | G | A | C | T | T | A | T |
| EU591677.1 | Odonata | *Davidius lunatus* | C | T | A | C | T | A | T | G | T | T | A | C | G | A | C | T | T | A | T |
| NC012644.1 | Odonata | *Davidius lunatus* | C | T | A | C | T | A | T | G | T | T | A | C | G | A | C | T | T | A | T |
| MH751436.1 | Odonata | *Deielia phaon* | C | T | A | C | T | A | T | G | T | T | A | C | G | A | C | T | T | A | T |
| NC042690.1 | Odonata | *Deielia phaon* | C | T | A | C | T | A | T | G | T | T | A | C | G | A | C | T | T | A | T |
| MF716899.1 | Odonata | *Enallagma cyathigerum* | C | T | A | C | T | A | T | G | T | T | A | C | G | A | C | T | T | A | T |
| MK522522.1 | Odonata | *Epophthalmia elegans* | C | T | A | C | T | A | T | G | T | T | A | C | G | A | C | T | T | A | T |
| NC046756.1 | Odonata | *Epophthalmia elegans* | C | T | A | C | T | A | T | G | T | T | A | C | G | A | C | T | T | A | T |
| HM126547.1 | Odonata | *Euphaea formosa* | C | T | A | C | T | A | T | G | T | T | A | C | G | A | C | T | T | A | T |
| NC014493.1 | Odonata | *Euphaea formosa* | C | T | A | C | T | A | T | G | T | T | A | C | G | A | C | T | T | A | T |
| KM244659.1 | Odonata | *Hydrobasileus croceus* | C | T | A | C | T | A | T | G | T | T | A | C | G | A | C | T | T | A | T |
| NC025758.1 | Odonata | *Hydrobasileus croceus* | C | T | A | C | T | A | T | G | T | T | A | C | G | A | C | T | T | A | T |
| KU958378.1 | Odonata | *Ischnura elegans* | C | T | A | C | T | A | T | G | T | T | A | C | G | A | C | T | T | A | T |
| MK951668.1 | Odonata | *Ischnura elegans* | C | T | A | C | T | A | T | G | T | T | A | C | G | A | C | T | T | A | T |
| NC031824.1 | Odonata | *Ischnura elegans* | C | T | A | C | T | A | T | G | T | T | A | C | G | A | C | T | T | A | T |

| | | | | | | | | | | | | | | | | | | | | | |
|---|---|---|---|---|---|---|---|---|---|---|---|---|---|---|---|---|---|---|---|---|---|
| KC878732.1 | Odonata | *Ischnura pumilio* | C | T | A | C | T | A | T | G | T | T | A | C | G | A | C | T | T | A | T |
| NC021617.1 | Odonata | *Ischnura pumilio* | C | T | A | C | T | A | T | G | T | T | A | C | G | A | C | T | T | A | T |
| MF990748.1 | Odonata | *Macromia daimoji* | C | T | A | C | T | A | T | G | T | T | A | C | G | A | C | T | T | A | T |
| NC041425.1 | Odonata | *Macromia daimoji* | C | T | A | C | T | A | T | G | T | T | A | C | G | A | C | T | T | A | T |
| MK722304.1 | Odonata | *Matrona basilaris* | C | T | A | C | T | A | T | G | T | T | A | C | G | A | C | T | T | A | T |
| MK722305.1 | Odonata | *Matrona basilaris* | C | T | A | C | T | A | T | G | T | T | A | C | G | A | C | T | T | A | T |
| KU958377.1 | Odonata | *Megaloprepus caerulatus* | C | T | A | C | T | A | T | G | T | T | A | C | G | A | C | T | T | A | T |
| NC031823.1 | Odonata | *Megaloprepus caerulatus* | C | T | A | C | T | A | T | G | T | T | A | C | G | A | C | T | T | A | T |
| KU871065.1 | Odonata | *Mnais costalis* | C | T | A | C | T | A | T | G | T | T | A | C | G | A | C | T | T | A | T |
| KY402222.1 | Odonata | *Nannophya pygmaea* | C | T | A | C | T | A | T | G | T | T | A | C | G | A | C | T | T | A | T |
| NC039410.1 | Odonata | *Nannophya pygmaea* | C | T | A | C | T | A | T | G | T | T | A | C | G | A | C | T | T | A | T |
| KU361233.1 | Odonata | *Orthetrum chrysis* | C | T | A | C | T | A | T | G | T | T | A | C | G | A | C | T | T | A | T |
| NC032048.1 | Odonata | *Orthetrum chrysis* | C | T | A | C | T | A | T | G | T | T | A | C | G | A | C | T | T | A | T |
| KU361232.1 | Odonata | *Orthetrum glaucum* | C | T | A | C | T | A | T | G | T | T | A | C | G | A | C | T | T | A | T |
| NC032047.1 | Odonata | *Orthetrum glaucum* | C | T | A | C | T | A | T | G | T | T | A | C | G | A | C | T | T | A | T |
| MH751437.1 | Odonata | *Orthetrum melania* | C | T | A | C | T | A | T | G | T | T | A | C | G | A | C | T | T | A | T |
| NC042732.1 | Odonata | *Orthetrum melania* | C | T | A | C | T | A | T | G | T | T | A | C | G | A | C | T | T | A | T |
| KU361234.1 | Odonata | *Orthetrum sabina* | C | T | A | C | T | A | T | G | T | T | A | C | G | A | C | T | T | A | T |
| MH751438.1 | Odonata | *Orthetrum sabina* | C | T | A | C | T | A | T | G | T | T | A | C | G | A | C | T | T | A | T |
| NC032049.1 | Odonata | *Orthetrum sabina* | C | T | A | C | T | A | T | G | T | T | A | C | G | A | C | T | T | A | T |

| | | | | | | | | | | | | | | | | | | | | | |
|---|---|---|---|---|---|---|---|---|---|---|---|---|---|---|---|---|---|---|---|---|---|
| KU361235.1 | Odonata | *Orthetrum testaceum* | C | T | A | C | T | A | T | G | T | T | A | C | G | A | C | T | T | A | T |
| MH751439.1 | Odonata | *Orthetrum testaceum* | C | T | A | C | T | A | T | G | T | T | A | C | G | A | C | T | T | A | T |
| NC032050.1 | Odonata | *Orthetrum testaceum* | C | T | A | C | T | A | T | G | T | T | A | C | G | A | C | T | T | A | T |
| AB126005.1 | Odonata | *Orthetrum triangulare melania* | C | T | A | C | T | A | T | G | T | T | A | C | G | A | C | T | T | A | T |
| MK951669.1 | Odonata | *Paracercion v-nigrum* | C | T | A | C | T | A | T | G | T | T | A | C | G | A | C | T | T | A | T |
| KP233804.1 | Odonata | *Platycnemis foliacea* | C | T | A | C | T | T | T | G | T | T | A | C | G | A | C | T | T | A | T |
| NC027180.1 | Odonata | *Platycnemis foliacea* | C | T | A | C | T | T | T | G | T | T | A | C | G | A | C | T | T | A | T |
| FJ606784.1 | Odonata | *Pseudolestes mirabilis* | C | T | A | C | T | A | T | G | T | T | A | C | G | A | C | T | T | A | T |
| NC020636.1 | Odonata | *Pseudolestes mirabilis* | C | T | A | C | T | A | T | G | T | T | A | C | G | A | C | T | T | A | T |
| MF150044.1 | Odonata | *Psolodesmus mandarinus* | C | T | A | C | T | A | T | G | T | T | A | C | G | A | C | T | T | A | T |
| NC042442.1 | Odonata | *Psolodesmus mandarinus* | C | T | A | C | T | A | T | G | T | T | A | C | G | A | C | T | T | A | T |
| MG594801.1 | Odonata | *Somatochlora hineana* | C | T | A | C | T | A | T | G | T | T | A | C | G | A | C | T | T | A | T |
| MH751440.1 | Odonata | *Tramea virginia* | C | T | A | C | T | A | T | G | T | T | A | C | G | A | C | T | T | A | T |
| NC042691.1 | Odonata | *Tramea virginia* | C | T | A | C | T | A | T | G | T | T | A | C | G | A | C | T | T | A | T |
| MH751441.1 | Odonata | *Trigomphus carus* | C | T | A | C | T | A | T | G | T | T | A | C | G | A | C | T | T | A | T |
| NC042733.1 | Odonata | *Trigomphus carus* | C | T | A | C | T | A | T | G | T | T | A | C | G | A | C | T | T | A | T |

| | | | | | | | | | | | | | | | | | | | | | |
|---|---|---|---|---|---|---|---|---|---|---|---|---|---|---|---|---|---|---|---|---|---|
| JX050224.1 | Odonata | *Vestalis melania* | C | T | A | C | T | A | T | G | T | T | A | C | G | A | C | T | T | A | T |
| NC023233.1 | Odonata | *Vestalis melania* | C | T | A | C | T | A | T | G | T | T | A | C | G | A | C | T | T | A | T |
| KM199685.1 | Plecoptera | *Acroneuria hainana* | C | T | A | C | T | A | T | G | T | T | A | C | G | A | C | T | T | A | T |
| NC026104.1 | Plecoptera | *Acroneuria hainana* | C | T | A | C | T | A | T | G | T | T | A | C | G | A | C | T | T | A | T |
| KX091846.1 | Plecoptera | *Agnetina aequalis* | C | T | A | C | T | A | T | G | T | T | A | C | G | A | C | T | T | A | T |
| MH085446.1 | Plecoptera | *Amphinemura longispina* | C | T | A | C | T | A | T | G | T | T | A | C | G | A | C | T | T | A | T |
| NC044748.1 | Plecoptera | *Amphinemura longispina* | C | T | A | C | T | A | T | G | T | T | A | C | G | A | C | T | T | A | T |
| KX091847.1 | Plecoptera | *Amphinemura sp.* | C | T | A | C | T | A | T | G | T | T | A | C | G | A | C | T | T | A | T |
| MH085447.1 | Plecoptera | *Amphinemura yao* | C | T | A | C | T | A | T | G | T | T | A | C | G | A | C | T | T | A | T |
| NC044749.1 | Plecoptera | *Amphinemura yao* | C | T | A | C | T | A | T | G | T | T | A | C | G | A | C | T | T | A | T |
| MK111413.1 | Plecoptera | *Antarctoperla michaelseni* | C | T | A | C | T | A | T | G | T | T | A | C | G | A | C | T | T | A | T |
| NC042199.1 | Plecoptera | *Antarctoperla michaelseni* | C | T | A | C | T | A | T | G | T | T | A | C | G | A | C | T | T | A | T |
| KR604721.1 | Plecoptera | *Apteroperla tikumana* | C | T | A | C | T | A | T | G | T | T | A | C | G | A | C | T | T | A | T |
| NC027698.1 | Plecoptera | *Apteroperla tikumana* | C | T | A | C | T | A | T | G | T | T | A | C | G | A | C | T | T | A | T |
| MT410838.1 | Plecoptera | *Brachyptera risi* | C | T | A | C | T | A | T | G | T | T | A | C | G | A | C | T | T | A | T |
| MG677941.1 | Plecoptera | *Calineuria stigmatica* | C | T | A | C | T | A | T | G | T | T | A | C | G | A | C | T | T | A | T |
| KX094942.1 | Plecoptera | *Capnia zijinshana* | C | T | A | C | T | A | T | G | T | T | A | C | G | A | C | T | T | A | T |
| NC034661.1 | Plecoptera | *Capnia zijinshana* | C | T | A | C | T | A | T | G | T | T | A | C | G | A | C | T | T | A | T |

| | | | | | | | | | | | | | | | | | | | | | |
|---|---|---|---|---|---|---|---|---|---|---|---|---|---|---|---|---|---|---|---|---|---|
| MG677942.1 | Plecoptera | *Caroperla siveci* | C | T | A | C | T | A | T | G | T | T | A | C | G | A | C | T | T | A | T |
| MF100783.1 | Plecoptera | *Cerconychia flectospina* | C | T | A | C | T | A | T | G | T | T | A | C | G | A | C | T | T | A | T |
| MN419914.1 | Plecoptera | *Claassenia* sp. | C | T | A | C | T | A | T | G | T | T | A | C | G | A | C | T | T | A | T |
| KC952026.1 | Plecoptera | *Cryptoperla* sp. | C | T | A | C | T | A | T | G | T | T | A | C | G | A | C | T | T | A | T |
| MK111416.1 | Plecoptera | *Diamphipnoa annulata* | C | T | A | C | T | A | T | G | T | T | A | C | G | A | C | T | T | A | T |
| NC042205.1 | Plecoptera | *Diamphipnoa annulata* | C | T | A | C | T | A | T | G | T | T | A | C | G | A | C | T | T | A | T |
| MK111417.1 | Plecoptera | *Diamphipnopsis* sp. | C | T | A | C | T | A | T | G | T | T | A | C | G | A | C | T | T | A | T |
| KF484757.1 | Plecoptera | *Dinocras cephalotes* | C | T | A | C | T | A | T | G | T | T | A | C | G | A | C | T | T | A | T |
| NC022843.1 | Plecoptera | *Dinocras cephalotes* | C | T | A | C | T | A | T | G | T | T | A | C | G | A | C | T | T | A | T |
| MG589787.1 | Plecoptera | *Doddsia occidentalis* | C | T | A | C | T | A | T | G | T | T | A | C | G | A | C | T | T | A | T |
| MK905888.1 | Plecoptera | *Etrocorema hochii* | C | T | A | C | T | A | T | G | T | T | A | C | G | A | C | T | T | A | T |
| MK905206.1 | Plecoptera | *Flavoperla* sp. | C | T | A | C | T | A | T | G | T | T | A | C | G | A | C | T | T | A | T |
| MN419916.1 | Plecoptera | *Flavoperla* sp. | C | T | A | C | T | A | T | G | T | T | A | C | G | A | C | T | T | A | T |
| MN419915.1 | Plecoptera | *Indonemoura auriformis* | C | T | A | C | T | A | T | G | T | T | A | C | G | A | C | T | T | A | T |
| MH085448.1 | Plecoptera | *Indonemoura jacobsoni* | C | T | A | C | T | A | T | G | T | T | A | C | G | A | C | T | T | A | T |
| NC044750.1 | Plecoptera | *Indonemoura jacobsoni* | C | T | A | C | T | A | T | G | T | T | A | C | G | A | C | T | T | A | T |

| | | | | | | | | | | | | | | | | | | | | |
|---|---|---|---|---|---|---|---|---|---|---|---|---|---|---|---|---|---|---|---|---|
| MH085449.1 | Plecoptera | *Indonemoura nohirae* | C | T | A | C | T | A | T | G | T | T | A | C | G | A | C | T | T | A | T |
| NC044751.1 | Plecoptera | *Indonemoura nohirae* | C | T | A | C | T | A | T | G | T | T | A | C | G | A | C | T | T | A | T |
| MF716959.1 | Plecoptera | *Isoperla bilineata* | C | T | A | C | T | A | T | G | T | T | A | C | G | A | C | T | T | A | T |
| NC038190.1 | Plecoptera | *Isoperla bilineata* | C | T | A | C | T | A | T | G | T | T | A | C | G | A | C | T | T | A | T |
| MG910457.1 | Plecoptera | *Isoperla eximia* | C | T | A | C | T | A | T | G | T | T | A | C | G | A | C | T | T | A | T |
| NC038167.1 | Plecoptera | *Isoperla eximia* | C | T | A | C | T | A | T | G | T | T | A | C | G | A | C | T | T | A | T |
| KT186102.1 | Plecoptera | *Kamimuria chungnanshana* | C | T | A | C | T | A | T | G | T | T | A | C | G | A | C | T | T | A | T |
| NC028076.1 | Plecoptera | *Kamimuria chungnanshana* | C | T | A | C | T | A | T | G | T | T | A | C | G | A | C | T | T | A | T |
| MN400755.1 | Plecoptera | *Kamimuria klapaleki* | C | T | A | C | T | A | T | G | T | T | A | C | G | A | C | T | T | A | T |
| KC894944.1 | Plecoptera | *Kamimuria wangi* | C | T | A | C | T | A | T | G | T | T | A | C | G | A | C | T | T | A | T |
| MK111414.1 | Plecoptera | *Klapopteryx armillata* | C | T | A | C | T | A | T | G | T | T | A | C | G | A | C | T | T | A | T |
| MK568475.1 | Plecoptera | *Leuctra fusca* | C | T | A | C | T | A | T | G | T | T | A | C | G | A | C | T | T | A | T |
| KP642637.1 | Plecoptera | *Mesocapnia arizonensis* | C | T | A | C | T | A | T | G | T | T | A | C | G | A | C | T | T | A | T |
| KY568983.1 | Plecoptera | *Mesocapnia daxingana* | C | T | A | C | T | A | T | G | T | T | A | C | G | A | C | T | T | A | T |
| MH085450.1 | Plecoptera | *Mesonemoura metafiligera* | C | T | A | C | T | A | T | G | T | T | A | C | G | A | C | T | T | A | T |
| NC044719.1 | Plecoptera | *Mesonemoura metafiligera* | C | T | A | C | T | A | T | G | T | T | A | C | G | A | C | T | T | A | T |

| | | | | | | | | | | | | | | | | | | | | | |
|---|---|---|---|---|---|---|---|---|---|---|---|---|---|---|---|---|---|---|---|---|---|
| MH085451.1 | Plecoptera | *Mesonemoura tritaenia* | C | T | A | C | T | A | T | G | T | T | A | C | G | A | C | T | T | A | T |
| NC044720.1 | Plecoptera | *Mesonemoura tritaenia* | C | T | A | C | T | A | T | G | T | T | A | C | G | A | C | T | T | A | T |
| MN096323.1 | Plecoptera | *Microperla geei* | C | T | A | C | T | A | T | G | T | T | A | C | G | A | C | T | T | A | T |
| MT410776.1 | Plecoptera | *Nemoura avicularis* | C | T | A | C | T | A | T | G | T | T | A | C | G | A | C | T | T | A | T |
| MT410849.1 | Plecoptera | *Nemoura cinerea* | C | T | A | C | T | A | T | G | T | T | A | C | G | A | C | T | T | A | T |
| KY940360.1 | Plecoptera | *Nemoura nankinensis* | C | T | A | C | T | A | T | G | T | T | A | C | G | A | C | T | T | A | T |
| NC034939.1 | Plecoptera | *Nemoura nankinensis* | C | T | A | C | T | A | T | G | T | T | A | C | G | A | C | T | T | A | T |
| MK290826.1 | Plecoptera | *Nemoura papilla* | C | T | A | C | T | A | T | G | T | T | A | C | G | A | C | T | T | A | T |
| MK111418.1 | Plecoptera | *Neonemura barrosi* | C | T | A | C | T | A | T | G | T | T | A | C | G | A | C | T | T | A | T |
| NC042206.1 | Plecoptera | *Neonemura barrosi* | C | T | A | C | T | A | T | G | T | T | A | C | G | A | C | T | T | A | T |
| KX091858.1 | Plecoptera | *Neoperla ignacsiveci* | C | T | A | C | T | A | T | G | T | T | A | C | G | A | C | T | T | A | T |
| KX091859.1 | Plecoptera | *Neoperla* sp. | C | T | A | C | T | A | T | G | T | T | A | C | G | A | C | T | T | A | T |
| MN400756.1 | Plecoptera | *Neoperlops gressitti* | C | T | A | C | T | A | T | G | T | T | A | C | G | A | C | T | T | A | T |
| MK111415.1 | Plecoptera | *Neuroperla schedingi* | C | T | A | C | T | A | T | G | T | T | A | C | G | A | C | T | T | A | T |
| NC042200.1 | Plecoptera | *Neuroperla schedingi* | C | T | A | C | T | A | T | G | T | T | A | C | G | A | C | T | T | A | T |
| MK686067.1 | Plecoptera | *Niponiella limbatella* | C | T | A | C | T | A | T | G | T | T | A | C | G | A | C | T | T | A | T |
| MK387068.1 | Plecoptera | *Peltoperlopsis cebuano* | C | T | A | C | T | A | T | G | T | T | A | C | G | A | C | T | T | A | T |

| | | | | | | | | | | | | | | | | | | | | |
|---|---|---|---|---|---|---|---|---|---|---|---|---|---|---|---|---|---|---|---|---|
| MF197377.1 | Plecoptera | *Perlodes* sp. | C | T | A | C | T | A | T | G | T | T | A | C | G | A | C | T | T | A | T |
| MH085452.1 | Plecoptera | *Protonemura kohnoae* | C | T | A | C | T | A | T | G | T | T | A | C | G | A | C | T | T | A | T |
| NC044752.1 | Plecoptera | *Protonemura kohnoae* | C | T | A | C | T | A | T | G | T | T | A | C | G | A | C | T | T | A | T |
| MT410850.1 | Plecoptera | *Protonemura meyeri* | C | T | A | C | T | A | T | G | T | T | A | C | G | A | C | T | T | A | T |
| NC050322.1 | Plecoptera | *Protonemura meyeri* | C | T | A | C | T | A | T | G | T | T | A | C | G | A | C | T | T | A | T |
| MH085453.1 | Plecoptera | *Protonemura orbiculata* | C | T | A | C | T | A | T | G | T | T | A | C | G | A | C | T | T | A | T |
| NC044753.1 | Plecoptera | *Protonemura orbiculata* | C | T | A | C | T | A | T | G | T | T | A | C | G | A | C | T | T | A | T |
| MG910458.1 | Plecoptera | *Pseudomegarcys japonica* | C | T | A | C | T | A | T | G | T | T | A | C | G | A | C | T | T | A | T |
| NC038168.1 | Plecoptera | *Pseudomegarcys japonica* | C | T | A | C | T | A | T | G | T | T | A | C | G | A | C | T | T | A | T |
| KU182360.1 | Plecoptera | *Pteronarcella badia* | C | T | A | C | T | A | T | G | T | T | A | C | G | A | C | T | T | A | T |
| NC029248.1 | Plecoptera | *Pteronarcella badia* | C | T | A | C | T | A | T | G | T | T | A | C | G | A | C | T | T | A | T |
| AY687866.1 | Plecoptera | *Pteronarcys princeps* | C | T | A | C | T | A | T | G | T | T | A | C | G | A | C | T | T | A | T |
| NC006133.1 | Plecoptera | *Pteronarcys princeps* | C | T | A | C | T | A | T | G | T | T | A | C | G | A | C | T | T | A | T |
| MK111419.1 | Plecoptera | *Rhopalopsole bulbifera* | C | T | A | C | T | A | T | G | T | T | A | C | G | A | C | T | T | A | T |
| NC042207.1 | Plecoptera | *Rhopalopsole* | C | T | A | C | T | A | T | G | T | T | A | C | G | A | C | T | T | A | T |

| | | | | | | | | | | | | | | | | | | | | |
|---|---|---|---|---|---|---|---|---|---|---|---|---|---|---|---|---|---|---|---|---|
| | | *bulbifera* | | | | | | | | | | | | | | | | | | |
| MH510071.1 | Plecoptera | *Scopura longa* | C | T | A | C | T | A | T | G | T | T | A | C | G | A | C | T | T | A | T |
| NC041105.1 | Plecoptera | *Scopura longa* | C | T | A | C | T | A | T | G | T | T | A | C | G | A | C | T | T | A | T |
| MK492253.1 | Plecoptera | *Sinacroneuria dabieshana* | C | T | A | C | T | A | T | G | T | T | A | C | G | A | C | T | T | A | T |
| MF716958.1 | Plecoptera | *Soliperla* sp. | C | T | A | C | T | A | T | G | T | T | A | C | G | A | C | T | T | A | T |
| NC038189.1 | Plecoptera | *Soliperla* sp. | C | T | A | C | T | A | T | G | T | T | A | C | G | A | C | T | T | A | T |
| MH085455.1 | Plecoptera | *Sphaeronemoura acutispina* | C | T | A | C | T | A | T | G | T | T | A | C | G | A | C | T | T | A | T |
| MH085454.1 | Plecoptera | *Sphaeronemoura grandicauda* | C | T | A | C | T | A | T | G | T | T | A | C | G | A | C | T | T | A | T |
| NC044754.1 | Plecoptera | *Sphaeronemoura grandicauda* | C | T | A | C | T | A | T | G | T | T | A | C | G | A | C | T | T | A | T |
| MK111420.1 | Plecoptera | *Sphaeronemoura hainana* | C | T | A | C | T | A | T | G | T | T | A | C | G | A | C | T | T | A | T |
| NC044755.1 | Plecoptera | *Sphaeronemoura hamistyla* | C | T | A | C | T | A | T | G | T | T | A | C | G | A | C | T | T | A | T |
| KR088971.1 | Plecoptera | *Styloperla* sp. | C | T | A | C | T | A | T | G | T | T | A | C | G | A | C | T | T | A | T |
| KX845569.1 | Plecoptera | *Styloperla spinicercia* | C | T | A | C | T | A | T | G | T | T | A | C | G | A | C | T | T | A | T |
| NC034809.1 | Plecoptera | *Styloperla spinicercia* | C | T | A | C | T | A | T | G | T | T | A | C | G | A | C | T | T | A | T |
| KX091865.1 | Plecoptera | *Suwallia bimaculata* | C | T | A | C | T | T | T | G | T | T | A | C | G | A | C | T | T | A | T |
| MN121757.1 | Plecoptera | *Suwallia bimaculata* | C | T | A | C | T | T | T | G | T | T | A | C | G | A | C | T | T | A | T |

| | | | | | | | | | | | | | | | | | | | | | |
|---|---|---|---|---|---|---|---|---|---|---|---|---|---|---|---|---|---|---|---|---|---|
| MF198253.1 | Plecoptera | *Suwallia teleckojensis* | C | T | A | C | T | T | T | G | T | T | A | C | G | A | C | T | T | A | T |
| NC037754.1 | Plecoptera | *Suwallia teleckojensis* | C | T | A | C | T | T | T | G | T | T | A | C | G | A | C | T | T | A | T |
| KM216826.1 | Plecoptera | *Sweltsa longistyla* | C | T | A | C | T | T | T | G | T | T | A | C | G | A | C | T | T | A | T |
| MG589786.1 | Plecoptera | *Taeniopteryx ugola* | C | T | A | C | T | A | T | G | T | T | A | C | G | A | C | T | T | A | T |
| NC037897.1 | Plecoptera | *Taeniopteryx ugola* | C | T | A | C | T | A | T | G | T | T | A | C | G | A | C | T | T | A | T |
| KM409708.1 | Plecoptera | *Togoperla* sp. | C | T | A | C | T | A | T | G | T | T | A | C | G | A | C | T | T | A | T |
| KY522907.1 | Plecoptera | *Zelandoperla fenestrata* | C | T | A | C | T | A | T | G | T | T | A | C | G | A | C | T | T | A | T |
| NC034997.1 | Plecoptera | *Zelandoperla fenestrata* | C | T | A | C | T | A | T | G | T | T | A | C | G | A | C | T | T | A | T |
| FJ456939.1 | Heteroptera | *Aphelocheirus ellipsoideus* | C | T | A | C | T | T | T | G | T | T | A | C | G | A | C | T | T | A | T |
| KX505849.1 | Heteroptera | *Aphelocheirus jendeki* | C | T | A | C | T | T | T | G | T | T | A | C | G | A | C | T | T | A | T |
| NC037370.1 | Heteroptera | *Aphelocheirus jendeki* | C | T | A | C | T | T | T | G | T | T | A | C | G | A | C | T | T | A | T |
| FJ456944.1 | Heteroptera | *Aquarius paludum* | C | T | A | C | T | T | T | G | T | T | A | C | G | A | C | T | T | A | T |
| NC012841.1 | Heteroptera | *Aquarius paludum* | C | T | A | C | T | T | T | G | T | T | A | C | G | A | C | T | T | A | T |
| MK251109.1 | Heteroptera | *Diplonychus esakii* | C | T | A | C | T | T | T | G | T | T | A | C | G | A | C | T | T | A | T |
| FJ456940.1 | Heteroptera | *Diplonychus rusticus* | C | T | A | C | T | T | T | G | T | T | A | C | G | A | C | T | T | A | T |

| | | | | | | | | | | | | | | | | | | | | | |
|---|---|---|---|---|---|---|---|---|---|---|---|---|---|---|---|---|---|---|---|---|---|
| FJ456949.1 | Heteroptera | *Enithares tibialis* | C | T | A | C | T | T | T | G | T | T | A | C | G | A | C | T | T | A | T |
| NC012819.1 | Heteroptera | *Enithares tibialis* | C | T | A | C | T | T | T | G | T | T | A | C | G | A | C | T | T | A | T |
| KP400582.1 | Heteroptera | *Entomovelia* sp. | C | T | A | C | T | T | T | G | T | T | A | C | G | A | C | T | T | A | T |
| MF177288.1 | Heteroptera | *Gigantometra gigas* | C | T | A | C | T | T | T | G | T | T | A | C | G | A | C | T | T | A | T |
| MF177289.1 | Heteroptera | *Gigantometra gigas* | C | T | A | C | T | T | T | G | T | T | A | C | G | A | C | T | T | A | T |
| NC041084.1 | Heteroptera | *Gigantometra gigas* | C | T | A | C | T | T | T | G | T | T | A | C | G | A | C | T | T | A | T |
| KJ027513.1 | Heteroptera | *Helotrephes semiglobosus semiglobosus* | C | T | A | C | T | T | T | G | T | T | A | C | G | A | C | T | T | A | T |
| FJ456951.1 | Heteroptera | *Helotrephes* sp. | C | T | A | C | T | T | T | G | T | T | A | C | G | A | C | T | T | A | T |
| NC012822.1 | Heteroptera | *Helotrephes* sp. | C | T | A | C | T | T | T | G | T | T | A | C | G | A | C | T | T | A | T |
| FJ456945.1 | Heteroptera | *Hydrometra greeni* | T | T | A | C | T | T | T | G | T | T | A | C | G | A | C | T | T | A | T |
| NC012842.1 | Heteroptera | *Hydrometra greeni* | T | T | A | C | T | T | T | G | T | T | A | C | G | A | C | T | T | A | T |
| FJ456947.1 | Heteroptera | *Ilyocoris cimicoides* | C | T | A | C | T | T | T | G | T | T | A | C | G | A | C | T | T | A | T |
| NC012845.1 | Heteroptera | *Ilyocoris cimicoides* | C | T | A | C | T | T | T | G | T | T | A | C | G | A | C | T | T | A | T |
| FJ456948.1 | Heteroptera | *Laccotrephes robustus* | C | T | A | C | T | T | T | G | T | T | A | C | G | A | C | T | T | A | T |
| NC012817.1 | Heteroptera | *Laccotrephes robustus* | C | T | A | C | T | T | T | G | T | T | A | C | G | A | C | T | T | A | T |
| FJ456946.1 | Heteroptera | *Leptopus* sp. | C | T | A | C | T | T | T | G | T | T | A | C | G | A | C | T | T | A | T |
| JQ910985.1 | Heteroptera | *Lethocerus deyrollei* | C | T | A | C | T | T | T | G | T | T | A | C | G | A | C | T | T | A | T |
| KU237288.1 | Heteroptera | *Lethocerus deyrollei* | C | T | A | C | T | T | T | G | T | T | A | C | G | A | C | T | T | A | T |

| | | | | | | | | | | | | | | | | | | | | |
|---|---|---|---|---|---|---|---|---|---|---|---|---|---|---|---|---|---|---|---|---|
| KM588201.1 | Heteroptera | *Lethocerus indicus* | C | T | A | C | T | T | T | G | T | T | A | C | G | A | C | T | T | A | T |
| KR072671.1 | Heteroptera | *Lethocerus indicus* | C | T | A | C | T | T | T | G | T | T | A | C | G | A | C | T | T | A | T |
| NC027194.1 | Heteroptera | *Lethocerus indicus* | C | T | A | C | T | T | T | G | T | T | A | C | G | A | C | T | T | A | T |
| MN027272.1 | Heteroptera | *Limnogonus hypoleucus* | C | T | A | C | T | T | T | G | T | T | A | C | G | A | C | T | T | A | T |
| NC044722.1 | Heteroptera | *Limnogonus hypoleucus* | C | T | A | C | T | T | T | G | T | T | A | C | G | A | C | T | T | A | T |
| MN027275.1 | Heteroptera | *Limnogonus intermedius* | C | T | A | C | T | T | T | G | T | T | A | C | G | A | C | T | T | A | T |
| NC044724.1 | Heteroptera | *Limnogonus intermedius* | C | T | A | C | T | T | T | G | T | T | A | C | G | A | C | T | T | A | T |
| MN027273.1 | Heteroptera | *Limnogonus poissoni* | C | T | A | C | T | T | T | G | T | T | A | C | G | A | C | T | T | A | T |
| MN027274.1 | Heteroptera | *Limnogonus poissoni* | C | T | A | C | T | T | T | G | T | T | A | C | G | A | C | T | T | A | T |
| MN027276.1 | Heteroptera | *Limnogonus poissoni* | C | T | A | C | T | T | T | G | T | T | A | C | G | A | C | T | T | A | T |
| NC044723.1 | Heteroptera | *Limnogonus poissoni* | C | T | A | C | T | T | T | G | T | T | A | C | G | A | C | T | T | A | T |
| FJ456942.1 | Heteroptera | *Lycorma delicatula* | C | T | A | C | T | T | T | G | T | T | A | C | G | A | C | T | T | A | T |
| NC012835.1 | Heteroptera | *Lycorma delicatula* | C | T | A | C | T | T | T | G | T | T | A | C | G | A | C | T | T | A | T |
| MN027278.1 | Heteroptera | *Metrocoris* sp. | C | T | A | C | T | T | T | G | T | T | A | C | G | A | C | T | T | A | T |
| MH382835.1 | Heteroptera | *Micronecta sahlbergii* | C | T | A | C | T | T | T | G | T | T | A | C | G | A | C | T | T | A | T |
| NC039588.1 | Heteroptera | *Micronecta sahlbergii* | C | T | A | C | T | T | T | G | T | T | A | C | G | A | C | T | T | A | T |
| KT031802.1 | Heteroptera | *Nepa hoffmanni* | C | T | A | C | T | T | T | G | T | T | A | C | G | A | C | T | T | A | T |
| NC028084.1 | Heteroptera | *Nepa hoffmanni* | C | T | A | C | T | T | T | G | T | T | A | C | G | A | C | T | T | A | T |

| | | | | | | | | | | | | | | | | | | | | | |
|---|---|---|---|---|---|---|---|---|---|---|---|---|---|---|---|---|---|---|---|---|---|
| FJ456943.1 | Heteroptera | *Nerthra indica* | C | T | A | C | T | T | T | G | T | T | A | C | G | A | C | T | T | A | T |
| NC012838.1 | Heteroptera | *Nerthra indica* | C | T | A | C | T | T | T | G | T | T | A | C | G | A | C | T | T | A | T |
| KX034086.1 | Heteroptera | *Notonecta chinensis* | C | T | A | C | T | T | T | G | T | T | A | C | G | A | C | T | T | A | T |
| NC036671.1 | Heteroptera | *Notonecta chinensis* | C | T | A | C | T | T | T | G | T | T | A | C | G | A | C | T | T | A | T |
| MH780861.1 | Heteroptera | *Notonecta montandoni* | C | T | A | C | T | T | T | G | T | T | A | C | G | A | C | T | T | A | T |
| NC045886.1 | Heteroptera | *Notonecta montandoni* | C | T | A | C | T | T | T | G | T | T | A | C | G | A | C | T | T | A | T |
| KM244707.1 | Heteroptera | *Notonectidae* sp. | C | T | A | C | T | T | T | G | T | T | A | C | G | A | C | T | T | A | T |
| FJ456950.1 | Heteroptera | *Ochterus marginatus* | C | T | A | C | T | T | T | G | T | T | A | C | G | A | C | T | T | A | T |
| JQ910988.1 | Heteroptera | *Perittopus* sp. | C | T | A | C | T | T | T | G | T | T | A | C | G | A | C | T | T | A | T |
| KP400583.1 | Heteroptera | *Ptilomera tigrina* | C | T | A | C | T | T | T | G | T | T | A | C | G | A | C | T | T | A | T |
| MN027271.1 | Heteroptera | *Rhagovelia reitteri* | C | T | A | C | T | T | T | G | T | T | A | C | G | A | C | T | T | A | T |
| NC044721.1 | Heteroptera | *Rhagovelia reitteri* | C | T | A | C | T | T | T | G | T | T | A | C | G | A | C | T | T | A | T |
| MN027279.1 | Heteroptera | *Rhagovelia* sp. | C | T | A | C | T | T | T | G | T | T | A | C | G | A | C | T | T | A | T |
| FJ456941.1 | Heteroptera | *Sigara septemlineata* | C | T | A | C | T | T | T | G | T | T | A | C | G | A | C | T | T | A | T |
| MN027277.1 | Heteroptera | *Trepobates* sp. | C | T | A | C | T | T | T | G | T | T | A | C | G | A | C | T | T | A | T |
| FJ456952.1 | Heteroptera | *Valentia hoffmanni* | C | T | A | C | T | T | T | G | T | T | A | C | G | A | C | T | T | A | T |
| NC012823.1 | Heteroptera | *Valentia hoffmanni* | C | T | A | C | T | T | T | G | T | T | A | C | G | A | C | T | T | A | T |
| | | A | 0.0 | 0.0 | 100.0 | 0.0 | 0.0 | 71.7 | 0.0 | 0.0 | 0.0 | 0.0 | 100.0 | 0.0 | 0.0 | 100.0 | 0.0 | 0.0 | 0.0 | 100.0 | 0.0 |
| | | T | 0.8 | 100.0 | 0.0 | 0.0 | 100.0 | 27.5 | 100.0 | 0.0 | 100.0 | 100.0 | 0.0 | 0.0 | 0.0 | 0.0 | 0.0 | 100.0 | 100.0 | 0.0 | 100.0 |

| | | | | | | | | | | | | | | | | | | | |
|---|---|---|---|---|---|---|---|---|---|---|---|---|---|---|---|---|---|---|---|
| G | 0.0 | 0.0 | 0.0 | 0.0 | 0.0 | 0.8 | 0.0 | 100.0 | 0.0 | 0.0 | 0.0 | 0.0 | 100.0 | 0.0 | 0.0 | 0.0 | 0.0 | 0.0 | 0.0 |
| C | 99.2 | 0.0 | 0.0 | 100.0 | 0.0 | 0.0 | 0.0 | 0.0 | 0.0 | 0.0 | 0.0 | 100.0 | 0.0 | 0.0 | 100.0 | 0.0 | 0.0 | 0.0 | 0.0 |

Table S12. Nucleotide sequences of universal primer (AQdb-12S_Hol_R) and homologous region of 214 sequences of insects used for development in this study.

| | | | | | | | | | | | | | | | | | | | | | | | | | |
|---|---|---|---|---|---|---|---|---|---|---|---|---|---|---|---|---|---|---|---|---|---|---|---|---|---|
| AQdb-12S_Hol_R | | | | T | A | M | W | Y | C | T | A | C | T | W | T | G | T | T | A | C | G | A | C | T | T |
| | | | A | 0 | 214 | 33 | 203 | 0 | 0 | 0 | 214 | 0 | 0 | 113 | 0 | 0 | 0 | 0 | 214 | 0 | 8 | 214 | 0 | 0 | 0 |
| | | | T | 214 | 0 | 0 | 8 | 171 | 8 | 214 | 0 | 0 | 214 | 94 | 214 | 0 | 214 | 214 | 0 | 0 | 0 | 0 | 0 | 214 | 214 |
| | | | G | 0 | 0 | 0 | 3 | 0 | 0 | 0 | 0 | 0 | 0 | 0 | 0 | 214 | 0 | 0 | 0 | 0 | 206 | 0 | 0 | 0 | 0 |
| | | | C | 0 | 0 | 181 | 0 | 43 | 206 | 0 | 0 | 214 | 0 | 7 | 0 | 0 | 0 | 0 | 0 | 214 | 0 | 0 | 214 | 0 | 0 |
| GenBank Acc. No. | Order | Species | | | | | | | | | | | | | | | | | | | | | | | |
| KF840564.1 | Megaloptera | *Acanthacorydalis orientalis* | | T | A | C | A | C | C | T | A | C | T | T | T | G | T | T | A | C | G | A | C | T | T |
| NC023462.1 | Megaloptera | *Acanthacorydalis orientalis* | | T | A | C | A | C | C | T | A | C | T | T | T | G | T | T | A | C | G | A | C | T | T |
| KU925864.1 | Megaloptera | *Archichauliodes deceptor* | | T | A | C | A | C | C | T | A | C | T | T | T | G | T | T | A | C | G | A | C | T | T |
| NC035504.1 | Megaloptera | *Archichauliodes deceptor* | | T | A | C | A | C | C | T | A | C | T | T | T | G | T | T | A | C | G | A | C | T | T |
| KU302713.1 | Megaloptera | *Chloronia mirifica* | | T | A | C | A | C | C | T | A | C | T | T | T | G | T | T | A | C | G | A | C | T | T |
| NC033349.1 | Megaloptera | *Chloronia mirifica* | | T | A | C | A | C | C | T | A | C | T | T | T | G | T | T | A | C | G | A | C | T | T |
| FJ171323.1 | Megaloptera | *Corydalus cornutus* | | T | A | C | A | C | C | T | A | C | T | T | T | G | T | T | A | C | G | A | C | T | T |
| NC011276.1 | Megaloptera | *Corydalus cornutus* | | T | A | C | A | C | C | T | A | C | T | T | T | G | T | T | A | C | G | A | C | T | T |

| | | | | | | | | | | | | | | | | | | | | | | | | |
|---|---|---|---|---|---|---|---|---|---|---|---|---|---|---|---|---|---|---|---|---|---|---|---|---|
| KJ806318.1 | Megaloptera | *Dysmicohermes ingens* | T | A | C | A | C | C | T | A | C | T | C | T | G | T | T | A | C | G | A | C | T | T |
| NC024657.1 | Megaloptera | *Dysmicohermes ingens* | T | A | C | A | C | C | T | A | C | T | C | T | G | T | T | A | C | G | A | C | T | T |
| JQ351950.1 | Megaloptera | *Neochauliodes bowringi* | T | A | C | A | C | C | T | A | C | T | T | T | G | T | T | A | C | G | A | C | T | T |
| NC023444.1 | Megaloptera | *Neochauliodes bowringi* | T | A | C | A | C | C | T | A | C | T | T | T | G | T | T | A | C | G | A | C | T | T |
| KF785799.1 | Megaloptera | *Neochauliodes fraternus* | T | A | C | A | C | C | T | A | C | T | T | T | G | T | T | A | C | G | A | C | T | T |
| NC025282.1 | Megaloptera | *Neochauliodes fraternus* | T | A | C | A | C | C | T | A | C | T | T | T | G | T | T | A | C | G | A | C | T | T |
| KX821680.1 | Megaloptera | *Neochauliodes parasparsus* | T | A | C | A | C | C | T | A | C | T | T | T | G | T | T | A | C | G | A | C | T | T |
| JX110703.1 | Megaloptera | *Neochauliodes punctatolosus* | T | A | C | A | T | C | T | A | C | T | T | T | G | T | T | A | C | G | A | C | T | T |
| NC018772.1 | Megaloptera | *Neochauliodes punctatolosus* | T | A | C | A | T | C | T | A | C | T | T | T | G | T | T | A | C | G | A | C | T | T |
| KF771894.1 | Megaloptera | *Neochauliodes rotundatus* | T | A | C | A | C | C | T | A | C | T | T | T | G | T | T | A | C | G | A | C | T | T |
| NC025281.1 | Megaloptera | *Neochauliodes rotundatus* | T | A | C | A | C | C | T | A | C | T | T | T | G | T | T | A | C | G | A | C | T | T |

| | | | | | | | | | | | | | | | | | | | | | | | | |
|---|---|---|---|---|---|---|---|---|---|---|---|---|---|---|---|---|---|---|---|---|---|---|---|---|
| KP126231.1 | Megaloptera | *Neoneuromus tonkinensis* | T | A | C | A | C | C | T | A | C | T | T | T | G | T | T | A | C | G | A | C | T | T |
| NC027852.1 | Megaloptera | *Neoneuromus tonkinensis* | T | A | C | A | C | C | T | A | C | T | T | T | G | T | T | A | C | G | A | C | T | T |
| KP126232.1 | Megaloptera | *Nevromus exterior* | T | A | C | A | C | C | T | A | C | T | T | T | G | T | T | A | C | G | A | C | T | T |
| NC027851.1 | Megaloptera | *Nevromus exterior* | T | A | C | A | C | C | T | A | C | T | T | T | G | T | T | A | C | G | A | C | T | T |
| KY230493.1 | Megaloptera | *Protochauliodes biconicus* | T | A | C | A | C | T | T | A | C | T | T | T | G | T | T | A | C | G | A | C | T | T |
| EU526394.1 | Megaloptera | *Protohermes concolorus* | T | A | C | A | T | C | T | A | C | T | T | T | G | T | T | A | C | G | A | C | T | T |
| NC011524.1 | Megaloptera | *Protohermes concolorus* | T | A | C | A | T | C | T | A | C | T | T | T | G | T | T | A | C | G | A | C | T | T |
| FJ859905.1 | Megaloptera | *Sialis hamata* | T | A | C | A | T | C | T | A | C | T | A | T | G | T | T | A | C | G | A | C | T | T |
| NC013256.1 | Megaloptera | *Sialis hamata* | T | A | C | A | T | C | T | A | C | T | A | T | G | T | T | A | C | G | A | C | T | T |
| KT876911.1 | Megaloptera | *Sialis* sp. | T | A | C | A | T | C | T | A | C | T | A | T | G | T | T | A | C | G | A | C | T | T |
| KT425084.1 | Megaloptera | *Stenosialis australiensis* | T | A | C | A | T | C | T | A | C | T | A | T | G | T | T | A | C | G | A | C | T | T |
| KY587199.1 | Neuroptera | *Abachrysa eureka* | T | A | C | A | T | C | T | A | C | T | A | T | G | T | T | A | C | G | A | C | T | T |
| NC015095.1 | Neuroptera | *Apochrysa matsumurae* | T | A | C | A | T | C | T | A | C | T | A | T | G | T | T | A | C | G | A | C | T | T |
| KC758703.1 | Neuroptera | *Ascalohybris subjacens* | T | A | C | A | T | C | T | A | C | T | A | T | G | T | T | A | C | G | A | C | T | T |

| | | | | | | | | | | | | | | | | | | | | | | | | |
|---|---|---|---|---|---|---|---|---|---|---|---|---|---|---|---|---|---|---|---|---|---|---|---|---|
| NC021428.1 | Neuroptera | *Ascalohybris subjacens* | T | A | C | A | T | C | T | A | C | T | A | T | G | T | T | A | C | G | A | C | T | T |
| FJ171324.1 | Neuroptera | *Ascaloptynx appendiculatus* | T | A | C | A | T | C | T | A | C | T | T | T | G | T | T | A | C | G | A | C | T | T |
| NC011277.1 | Neuroptera | *Ascaloptynx appendiculatus* | T | A | C | A | T | C | T | A | C | T | T | T | G | T | T | A | C | G | A | C | T | T |
| KT425083.1 | Neuroptera | *Balmes birmanus* | T | A | C | A | T | C | T | A | C | T | A | T | G | T | T | A | C | G | A | C | T | T |
| KX369241.1 | Neuroptera | *Bullanga florida* | T | A | C | A | T | C | T | A | C | T | T | T | G | T | T | A | C | G | A | C | T | T |
| NC032298.1 | Neuroptera | *Bullanga florida* | T | A | C | A | T | C | T | A | C | T | T | T | G | T | T | A | C | G | A | C | T | T |
| KT425069.1 | Neuroptera | *Chasmoptera huttii* | T | A | C | A | C | C | T | A | C | T | T | T | G | T | T | A | C | G | A | C | T | T |
| JX033119.1 | Neuroptera | *Chrysopa pallens* | T | A | C | A | T | C | T | A | C | T | A | T | G | T | T | A | C | G | A | C | T | T |
| NC019618.1 | Neuroptera | *Chrysopa pallens* | T | A | C | A | T | C | T | A | C | T | A | T | G | T | T | A | C | G | A | C | T | T |
| KU877169.1 | Neuroptera | *Chrysoperla externa* | T | A | C | A | T | C | T | A | C | T | A | T | G | T | T | A | C | G | A | C | T | T |
| NC030341.1 | Neuroptera | *Chrysoperla externa* | T | A | C | A | T | C | T | A | C | T | A | T | G | T | T | A | C | G | A | C | T | T |
| KP027407.1 | Neuroptera | *Chrysoperla nipponensis* | T | A | C | A | T | C | T | A | C | T | A | T | G | T | T | A | C | G | A | C | T | T |
| NC015093.1 | Neuroptera | *Chrysoperla nipponensis* | T | A | C | A | T | C | T | A | C | T | A | T | G | T | T | A | C | G | A | C | T | T |
| KT425088.1 | Neuroptera | *Climacia areolaris* | T | A | C | A | T | C | T | A | C | T | T | T | G | T | T | A | C | G | A | C | T | T |
| KT425078.1 | Neuroptera | *Coniopteryx* sp. | T | A | C | A | C | C | T | A | C | T | T | T | G | T | T | A | C | G | A | C | T | T |

| | | | | | | | | | | | | | | | | | | | | | | | | |
|---|---|---|---|---|---|---|---|---|---|---|---|---|---|---|---|---|---|---|---|---|---|---|---|---|
| KT425068.1 | Neuroptera | *Dendroleon pantherinus* | T | A | C | A | T | C | T | A | C | T | A | T | G | T | T | A | C | G | A | C | T | T |
| MK301246.1 | Neuroptera | *Dendroleon pantherinus* | T | A | C | A | T | C | T | A | C | T | A | T | G | T | T | A | C | G | A | C | T | T |
| KT425073.1 | Neuroptera | *Dilar* sp. | T | A | C | A | T | C | T | A | C | T | T | T | G | T | T | A | C | G | A | C | T | T |
| FJ859906.1 | Neuroptera | *Ditaxis biseriata* | T | A | C | A | T | C | T | A | C | T | A | T | G | T | T | A | C | G | A | C | T | T |
| NC013257.1 | Neuroptera | *Ditaxis biseriata* | T | A | C | A | T | C | T | A | C | T | A | T | G | T | T | A | C | G | A | C | T | T |
| KT425087.1 | Neuroptera | *Drepanepteryx phalaenoides* | T | A | A | A | C | C | T | A | C | T | T | T | G | T | T | A | C | G | A | C | T | T |
| KF701327.1 | Neuroptera | *Epacanthaclisis banksi* | T | A | C | A | T | C | T | A | C | T | A | T | G | T | T | A | C | G | A | C | T | T |
| NC025905.1 | Neuroptera | *Epacanthaclisis banksi* | T | A | C | A | T | C | T | A | C | T | A | T | G | T | T | A | C | G | A | C | T | T |
| MF972496.1 | Neuroptera | *Euclimacia badia* | T | A | C | A | T | C | T | A | C | T | A | T | G | T | T | A | C | G | A | C | T | T |
| NC039773.1 | Neuroptera | *Euclimacia badia* | T | A | C | A | T | C | T | A | C | T | A | T | G | T | T | A | C | G | A | C | T | T |
| KT425080.1 | Neuroptera | *Eumantispa harmandi* | T | A | C | A | T | C | T | A | C | T | A | T | G | T | T | A | C | G | A | C | T | T |
| MF972495.1 | Neuroptera | *Eumantispa harmandi* | T | A | C | A | T | C | T | A | C | T | A | T | G | T | T | A | C | G | A | C | T | T |
| NC039772.1 | Neuroptera | *Eumantispa harmandi* | T | A | C | A | T | C | T | A | C | T | A | T | G | T | T | A | C | G | A | C | T | T |
| KT425072.1 | Neuroptera | *Fontecilla* | T | A | C | A | T | T | T | A | C | T | A | T | G | T | T | A | C | G | A | C | T | T |

| | | | | | | | | | | | | | | | | | | | | | | | | |
|---|---|---|---|---|---|---|---|---|---|---|---|---|---|---|---|---|---|---|---|---|---|---|---|---|
| | | *graphicus* | | | | | | | | | | | | | | | | | | | | | | |
| KY364372.1 | Neuroptera | *Gatzara jezoensis* | T | A | C | A | T | C | T | A | C | T | A | T | G | T | T | A | C | G | A | C | T | T |
| MK341582.1 | Neuroptera | *Gryposmylus pennyi* | T | A | C | A | T | C | T | A | C | T | A | T | G | T | T | A | C | G | A | C | T | T |
| MN852445.1 | Neuroptera | *Hemerobius japonicus* | T | A | C | A | T | C | T | A | C | T | T | T | G | T | T | A | C | G | A | C | T | T |
| NC048517.1 | Neuroptera | *Hemerobius japonicus* | T | A | C | A | T | C | T | A | C | T | T | T | G | T | T | A | C | G | A | C | T | T |
| KT425077.1 | Neuroptera | *Heterosmylus* sp. | T | A | C | A | T | C | T | A | C | T | A | T | G | T | T | A | C | G | A | C | T | T |
| KT425085.1 | Neuroptera | *Inocellia fujiana* | T | A | C | A | T | C | T | A | C | T | T | T | G | T | T | A | C | G | A | C | T | T |
| KY587200.1 | Neuroptera | *Italochrysa insignis* | T | A | C | A | T | C | T | A | C | T | A | T | G | T | T | A | C | G | A | C | T | T |
| KY587201.1 | Neuroptera | *Leucochrysa pretiosa* | T | A | C | A | T | C | T | A | C | T | A | T | G | T | T | A | C | G | A | C | T | T |
| NC015609.1 | Neuroptera | *Libelloides macaronius* | T | A | C | A | C | C | T | A | C | T | A | T | G | T | T | A | C | G | A | C | T | T |
| MF972494.1 | Neuroptera | *Mantispa japonica* | T | A | C | A | T | C | T | A | C | T | A | T | G | T | T | A | C | G | A | C | T | T |
| NC039771.1 | Neuroptera | *Mantispa japonica* | T | A | C | A | T | C | T | A | C | T | A | T | G | T | T | A | C | G | A | C | T | T |
| MK472073.1 | Neuroptera | *Micromus angulatus* | T | A | C | A | T | C | T | A | C | T | T | T | G | T | T | A | C | G | A | C | T | T |
| NC042677.1 | Neuroptera | *Micromus angulatus* | T | A | C | A | T | C | T | A | C | T | T | T | G | T | T | A | C | G | A | C | T | T |

| | | | | | | | | | | | | | | | | | | | | | | | | |
|---|---|---|---|---|---|---|---|---|---|---|---|---|---|---|---|---|---|---|---|---|---|---|---|---|
| KT425075.1 | Neuroptera | *Micromus* sp. | T | A | C | A | T | C | T | A | C | T | T | T | G | T | T | A | C | G | A | C | T | T |
| KT425092.1 | Neuroptera | *Mucroberotha vesicaria* | T | A | C | A | T | C | T | A | C | T | A | T | G | T | T | A | C | G | A | C | T | T |
| KT425089.1 | Neuroptera | *Myiodactylus osmyloides* | T | A | C | A | T | C | T | A | C | T | A | T | G | T | T | A | C | G | A | C | T | T |
| MT118664.1 | Neuroptera | *Myrmeleon formicarius* | T | A | C | A | C | C | T | A | C | T | A | T | G | T | T | A | C | G | A | C | T | T |
| NC047285.1 | Neuroptera | *Myrmeleon formicarius* | T | A | C | A | C | C | T | A | C | T | A | T | G | T | T | A | C | G | A | C | T | T |
| KJ461323.1 | Neuroptera | *Myrmeleon immanis* | T | A | C | A | C | C | T | A | C | T | A | T | G | T | T | A | C | G | A | C | T | T |
| NC024826.1 | Neuroptera | *Myrmeleon immanis* | T | A | C | A | C | C | T | A | C | T | A | T | G | T | T | A | C | G | A | C | T | T |
| KT425071.1 | Neuroptera | *Nallachius americanus* | T | A | A | G | T | C | T | A | C | T | T | T | G | T | T | A | C | G | A | C | T | T |
| KT425086.1 | Neuroptera | *Negha inflata* | T | A | C | A | T | C | T | A | C | T | T | T | G | T | T | A | C | G | A | C | T | T |
| KT425079.1 | Neuroptera | *Nemoptera coa* | T | A | C | A | C | C | T | A | C | T | C | T | G | T | T | A | C | G | A | C | T | T |
| KR078257.1 | Neuroptera | *Neuronema laminatum* | T | A | C | A | T | C | T | A | C | T | T | T | G | T | T | A | C | G | A | C | T | T |
| NC028153.1 | Neuroptera | *Neuronema laminatum* | T | A | C | A | T | C | T | A | C | T | T | T | G | T | T | A | C | G | A | C | T | T |
| KT425074.1 | Neuroptera | *Nevrorthus* | T | A | C | A | T | C | T | A | C | T | A | T | G | T | T | A | C | G | A | C | T | T |

| | | | | | | | | | | | | | | | | | | | | | | | |
|---|---|---|---|---|---|---|---|---|---|---|---|---|---|---|---|---|---|---|---|---|---|---|---|
| | | *apatelios* | | | | | | | | | | | | | | | | | | | | | |
| KT425076.1 | Neuroptera | *Nipponeurorthus fuscinervis* | T | A | C | A | T | C | T | A | C | T | A | T | G | T | T | A | C | G | A | C | T | T |
| KP264629.1 | Neuroptera | *Nothancyla verreauxi* | T | A | C | A | T | C | T | A | C | T | A | T | G | T | T | A | C | G | A | C | T | T |
| KP264630.1 | Neuroptera | *Nothochrysa* sp. | T | A | C | A | T | C | T | A | C | T | A | T | G | T | T | A | C | G | A | C | T | T |
| KJ461322.1 | Neuroptera | *Nymphes myrmeleonoides* | T | A | C | G | T | C | T | A | C | T | A | T | G | T | T | A | C | G | A | C | T | T |
| NC024825.1 | Neuroptera | *Nymphes myrmeleonoides* | T | A | C | G | T | C | T | A | C | T | A | T | G | T | T | A | C | G | A | C | T | T |
| KT425090.1 | Neuroptera | *Oliarces clara* | T | A | C | A | T | C | T | A | C | T | A | T | G | T | T | A | C | G | A | C | T | T |
| MN818868.1 | Neuroptera | *Osmylus atomatus* | T | A | C | A | T | C | T | A | C | T | A | T | G | T | T | A | C | G | A | C | T | T |
| MN818867.1 | Neuroptera | *Osmylus fulvicephalus* | T | A | C | A | T | C | T | A | C | T | A | T | G | T | T | A | C | G | A | C | T | T |
| MN617021.1 | Neuroptera | *Osmylus lucalatus* | T | A | C | A | T | C | T | A | C | T | A | T | G | T | T | A | C | G | A | C | T | T |
| MN818870.1 | Neuroptera | *Osmylus* sp. | T | A | C | A | T | C | T | A | C | T | A | T | G | T | T | A | C | G | A | C | T | T |
| MN818869.1 | Neuroptera | *Osmylus zheanus* | T | A | C | A | T | C | T | A | C | T | A | T | G | T | T | A | C | G | A | C | T | T |
| KY587202.1 | Neuroptera | *Parankylopteryx* sp. | T | A | C | A | T | C | T | A | C | T | A | T | G | T | T | A | C | G | A | C | T | T |
| FJ171325.1 | Neuroptera | *Polystoechotes punctatus* | T | A | C | A | T | C | T | A | C | T | A | T | G | T | T | A | C | G | A | C | T | T |
| NC011278.1 | Neuroptera | *Polystoechotes* | T | A | C | A | T | C | T | A | C | T | A | T | G | T | T | A | C | G | A | C | T | T |

| | | | | | | | | | | | | | | | | | | | | | | | | |
|---|---|---|---|---|---|---|---|---|---|---|---|---|---|---|---|---|---|---|---|---|---|---|---|---|
| | | *punctatus* | | | | | | | | | | | | | | | | | | | | | | |
| KT425082.1 | Neuroptera | *Psychopsis coelivaga* | T | A | C | A | C | C | T | A | C | T | A | T | G | T | T | A | C | G | A | C | T | T |
| KF626446.1 | Neuroptera | *Rapisma xizangense* | T | A | C | A | T | C | T | A | C | T | A | T | G | T | T | A | C | G | A | C | T | T |
| NC023362.1 | Neuroptera | *Rapisma xizangense* | T | A | C | A | T | C | T | A | C | T | A | T | G | T | T | A | C | G | A | C | T | T |
| KF626447.1 | Neuroptera | *Rapisma zayuanum* | T | A | C | A | T | C | T | A | C | T | A | T | G | T | T | A | C | G | A | C | T | T |
| NC023363.1 | Neuroptera | *Rapisma zayuanum* | T | A | C | A | T | C | T | A | C | T | A | T | G | T | T | A | C | G | A | C | T | T |
| KT425067.1 | Neuroptera | *Semidalis aleyrodiformis* | T | A | C | A | C | C | T | A | C | T | T | T | G | T | T | A | C | G | A | C | T | T |
| KT425070.1 | Neuroptera | *Sisyra nigra* | T | A | C | A | T | C | T | A | C | T | T | T | G | T | T | A | C | G | A | C | T | T |
| KT425081.1 | Neuroptera | *Stenobiella* sp. | T | A | C | A | T | C | T | A | C | T | A | T | G | T | T | A | C | G | A | C | T | T |
| MH361300.1 | Neuroptera | *Suhpalacsa longialata* | T | A | C | A | T | C | T | A | C | T | A | T | G | T | T | A | C | G | A | C | T | T |
| NC039948.1 | Neuroptera | *Suhpalacsa longialata* | T | A | C | A | T | C | T | A | C | T | A | T | G | T | T | A | C | G | A | C | T | T |
| MK301247.1 | Neuroptera | *Suhpalacsa* sp. | T | A | C | A | T | C | T | A | C | T | A | T | G | T | T | A | C | G | A | C | T | T |
| MK408756.1 | Neuroptera | *Thaumatosmylus hainanus* | T | A | C | A | C | C | T | A | C | T | A | T | G | T | T | A | C | G | A | C | T | T |

| | | | | | | | | | | | | | | | | | | | | | | | | |
|---|---|---|---|---|---|---|---|---|---|---|---|---|---|---|---|---|---|---|---|---|---|---|---|---|
| NC046578.1 | Neuroptera | *Thaumatosmylus hainanus* | T | A | C | A | C | C | T | A | C | T | A | T | G | T | T | A | C | G | A | C | T | T |
| KC515397.1 | Neuroptera | *Thyridosmylus langii* | T | A | C | A | C | C | T | A | C | T | A | T | G | T | T | A | C | G | A | C | T | T |
| NC021415.1 | Neuroptera | *Thyridosmylus langii* | T | A | C | A | C | C | T | A | C | T | A | T | G | T | T | A | C | G | A | C | T | T |
| MH020192.1 | Coreoptera | *Abscondita anceyi* | T | A | A | A | T | C | T | A | C | T | T | T | G | T | T | A | C | G | A | C | T | T |
| NC039706.1 | Coreoptera | *Abscondita anceyi* | T | A | A | A | T | C | T | A | C | T | T | T | G | T | T | A | C | G | A | C | T | T |
| MK122952.1 | Coreoptera | *Abscondita chinensis* | T | A | A | A | T | C | T | A | C | T | T | T | G | T | T | A | C | G | A | C | T | T |
| MK292092.1 | Coreoptera | *Abscondita terminalis* | T | A | A | A | T | C | T | A | C | T | T | T | G | T | T | A | C | G | A | C | T | T |
| NC044776.1 | Coreoptera | *Abscondita terminalis* | T | A | A | A | T | C | T | A | C | T | T | T | G | T | T | A | C | G | A | C | T | T |
| KT876878.1 | Coreoptera | *Acilius sulcatus* | T | A | C | A | T | C | T | A | C | T | A | T | G | T | T | A | C | G | A | C | T | T |
| KX758085.1 | Coreoptera | *Aquatica ficta* | T | A | A | A | T | C | T | A | C | T | T | T | G | T | T | A | C | G | A | C | T | T |
| NC035060.1 | Coreoptera | *Aquatica ficta* | T | A | A | A | T | C | T | A | C | T | T | T | G | T | T | A | C | G | A | C | T | T |
| NC035755.1 | Coreoptera | *Aquatica lateralis* | T | A | A | A | T | C | T | A | C | T | T | T | G | T | T | A | C | G | A | C | T | T |
| KF667531.1 | Coreoptera | *Aquatica leii* | T | A | A | A | T | C | T | A | C | T | T | T | G | T | T | A | C | G | A | C | T | T |
| NC025276.1 | Coreoptera | *Aquatica leii* | T | A | A | A | T | C | T | A | C | T | T | T | G | T | T | A | C | G | A | C | T | T |
| KX758086.1 | Coreoptera | *Aquatica wuhana* | T | A | A | A | T | C | T | A | C | T | T | T | G | T | T | A | C | G | A | C | T | T |
| NC035061.1 | Coreoptera | *Aquatica wuhana* | T | A | A | A | T | C | T | A | C | T | T | T | G | T | T | A | C | G | A | C | T | T |

| | | | | | | | | | | | | | | | | | | | | | | | |
|---|---|---|---|---|---|---|---|---|---|---|---|---|---|---|---|---|---|---|---|---|---|---|---|
| KX229747.1 | Coreoptera | *Asymmetricata circumdata* | T | A | A | A | T | C | T | A | C | T | T | T | G | T | T | A | C | G | A | C | T | T |
| MK292113.1 | Coreoptera | *Asymmetricata circumdata* | T | A | A | A | T | C | T | A | C | T | T | T | G | T | T | A | C | G | A | C | T | T |
| NC032062.1 | Coreoptera | *Asymmetricata circumdata* | T | A | A | A | T | C | T | A | C | T | T | T | G | T | T | A | C | G | A | C | T | T |
| KJ922151.1 | Coreoptera | *Bicellonychia lividipennis* | T | A | C | A | C | C | T | A | C | T | T | T | G | T | T | A | C | G | A | C | T | T |
| NC030060.1 | Coreoptera | *Bicellonychia lividipennis* | T | A | C | A | C | C | T | A | C | T | T | T | G | T | T | A | C | G | A | C | T | T |
| KT876885.1 | Coreoptera | *Colymbetes fuscus* | T | A | C | A | T | C | T | A | C | T | A | T | G | T | T | A | C | G | A | C | T | T |
| MK292114.1 | Coreoptera | *Curtos bilineatus* | T | A | A | A | T | C | T | A | C | T | C | T | G | T | T | A | C | G | A | C | T | T |
| NC044789.1 | Coreoptera | *Curtos bilineatus* | T | A | A | A | T | C | T | A | C | T | C | T | G | T | T | A | C | G | A | C | T | T |
| MK609965.1 | Coreoptera | *Curtos costipennis* | T | A | A | A | T | C | T | A | C | T | A | T | G | T | T | A | C | G | A | C | T | T |
| MH651351.1 | Coreoptera | *Diaphanes citrinus* | T | A | A | A | T | C | T | A | C | T | A | T | G | T | T | A | C | G | A | C | T | T |
| MK292116.1 | Coreoptera | *Diaphanes mendax* | T | A | C | A | T | C | T | A | C | T | T | T | G | T | T | A | C | G | A | C | T | T |
| NC044791.1 | Coreoptera | *Diaphanes mendax* | T | A | C | A | T | C | T | A | C | T | T | T | G | T | T | A | C | G | A | C | T | T |
| MK292094.1 | Coreoptera | *Diaphanes nubilus* | T | A | C | A | T | C | T | A | C | T | A | T | G | T | T | A | C | G | A | C | T | T |
| NC044787.1 | Coreoptera | *Diaphanes nubilus* | T | A | C | A | T | C | T | A | C | T | A | T | G | T | T | A | C | G | A | C | T | T |

| | | | | | | | | | | | | | | | | | | | | | | | | |
|---|---|---|---|---|---|---|---|---|---|---|---|---|---|---|---|---|---|---|---|---|---|---|---|---|
| MK292118.1 | Coreoptera | *Diaphanes pectinealis* | T | A | C | A | T | C | T | A | C | T | A | T | G | T | T | A | C | G | A | C | T | T |
| NC044793.1 | Coreoptera | *Diaphanes pectinealis* | T | A | C | A | T | C | T | A | C | T | A | T | G | T | T | A | C | G | A | C | T | T |
| MK292095.1 | Coreoptera | *Diaphanes* sp. | T | A | C | A | T | C | T | A | C | T | A | T | G | T | T | A | C | G | A | C | T | T |
| MG242622.1 | Coreoptera | *Ellychnia corrusca* | T | A | C | A | T | C | T | A | C | T | T | T | G | T | T | A | C | G | A | C | T | T |
| MK292112.1 | Coreoptera | *Emeia pseudosauteri* | T | A | C | A | T | C | T | A | C | T | A | T | G | T | T | A | C | G | A | C | T | T |
| NC044778.1 | Coreoptera | *Emeia pseudosauteri* | T | A | C | A | T | C | T | A | C | T | A | T | G | T | T | A | C | G | A | C | T | T |
| KT876897.1 | Coreoptera | *Hydroporus erythrocephalus* | T | A | C | A | T | C | T | A | C | T | A | T | G | T | T | A | C | G | A | C | T | T |
| KT876896.1 | Coreoptera | *Hydroporus obscurus* | T | A | C | A | T | C | T | A | C | T | A | T | G | T | T | A | C | G | A | C | T | T |
| KT876901.1 | Coreoptera | *Hygrotus confluens* | T | A | C | A | T | C | T | A | C | T | T | T | G | T | T | A | C | G | A | C | T | T |
| KT876900.1 | Coreoptera | *Hygrotus impressopunctatus* | T | A | A | A | T | C | T | A | C | T | T | T | G | T | T | A | C | G | A | C | T | T |
| KT876899.1 | Coreoptera | *Hygrotus inaequalis* | T | A | C | A | T | C | T | A | C | T | T | T | G | T | T | A | C | G | A | C | T | T |
| KX087299.1 | Coreoptera | *Hygrotus nigrolineatus* | T | A | C | A | T | C | T | A | C | T | T | T | G | T | T | A | C | G | A | C | T | T |

| | | | | | | | | | | | | | | | | | | | | | | | | |
|---|---|---|---|---|---|---|---|---|---|---|---|---|---|---|---|---|---|---|---|---|---|---|---|---|
| MH427718.1 | Coreoptera | *Inflata indica* | T | A | A | A | T | C | T | A | C | T | T | T | G | T | T | A | C | G | A | C | T | T |
| NC039700.1 | Coreoptera | *Inflata indica* | T | A | A | A | T | C | T | A | C | T | T | T | G | T | T | A | C | G | A | C | T | T |
| MK292091.1 | Coreoptera | *Lamprigera yunnana* | T | A | C | A | C | C | T | A | C | T | C | T | G | T | T | A | C | G | A | C | T | T |
| NC044786.1 | Coreoptera | *Lamprigera yunnana* | T | A | C | A | C | C | T | A | C | T | C | T | G | T | T | A | C | G | A | C | T | T |
| KX087302.1 | Coreoptera | *Lampyris noctiluca* | T | A | C | A | T | C | T | A | C | T | T | T | G | T | T | A | C | G | A | C | T | T |
| MG912994.1 | Coreoptera | *Limbodessus palmulaoides* | T | A | C | A | C | C | T | A | C | T | A | T | G | T | T | A | C | G | A | C | T | T |
| NC037749.1 | Coreoptera | *Limbodessus palmulaoides* | T | A | C | A | C | C | T | A | C | T | A | T | G | T | T | A | C | G | A | C | T | T |
| KT876902.1 | Coreoptera | *Liopterus haemorrhoidalis* | T | A | C | A | T | C | T | A | C | T | T | T | G | T | T | A | C | G | A | C | T | T |
| NC022472.1 | Coreoptera | *Luciola cruciata* | T | A | A | A | T | C | T | A | C | T | A | T | G | T | T | A | C | G | A | C | T | T |
| MG770613.1 | Coreoptera | *Luciola curtithorax* | T | A | A | A | T | C | T | A | C | T | T | T | G | T | T | A | C | G | A | C | T | T |
| NC038225.1 | Coreoptera | *Luciola curtithorax* | T | A | A | A | T | C | T | A | C | T | T | T | G | T | T | A | C | G | A | C | T | T |
| KP313820.1 | Coreoptera | *Luciola substriata* | T | A | A | A | T | C | T | A | C | T | T | T | G | T | T | A | C | G | A | C | T | T |
| NC027176.1 | Coreoptera | *Luciola substriata* | T | A | A | A | T | C | T | A | C | T | T | T | G | T | T | A | C | G | A | C | T | T |
| FJ859901.1 | Coreoptera | *Macrogyrus oblongus* | T | A | C | A | T | C | T | A | C | T | A | T | G | T | T | A | C | G | A | C | T | T |
| NC013249.1 | Coreoptera | *Macrogyrus oblongus* | T | A | C | A | T | C | T | A | C | T | A | T | G | T | T | A | C | G | A | C | T | T |

| | | | | | | | | | | | | | | | | | | | | | | | | |
|---|---|---|---|---|---|---|---|---|---|---|---|---|---|---|---|---|---|---|---|---|---|---|---|---|
| MG912995.1 | Coreoptera | *Paroster macrosturtensis* | T | A | C | A | T | T | T | A | C | T | T | T | G | T | T | A | C | G | A | C | T | T |
| NC037750.1 | Coreoptera | *Paroster macrosturtensis* | T | A | C | A | T | T | T | A | C | T | T | T | G | T | T | A | C | G | A | C | T | T |
| MG912996.1 | Coreoptera | *Paroster mesosturtensis* | T | A | C | A | T | T | T | A | C | T | T | T | G | T | T | A | C | G | A | C | T | T |
| NC037751.1 | Coreoptera | *Paroster mesosturtensis* | T | A | C | A | T | T | T | A | C | T | T | T | G | T | T | A | C | G | A | C | T | T |
| MG912997.1 | Coreoptera | *Paroster microsturtensis* | T | A | C | A | T | T | T | A | C | T | T | T | G | T | T | A | C | G | A | C | T | T |
| NC037752.1 | Coreoptera | *Paroster microsturtensis* | T | A | C | A | T | T | T | A | C | T | T | T | G | T | T | A | C | G | A | C | T | T |
| KY778696.1 | Coreoptera | *Photinus pyralis* | T | A | C | A | T | C | T | A | C | T | T | T | G | T | T | A | C | G | A | C | T | T |
| MF686051.1 | Coreoptera | *Pteroptyx maipo* | T | A | A | A | T | C | T | A | C | T | T | T | G | T | T | A | C | G | A | C | T | T |
| NC036353.1 | Coreoptera | *Pteroptyx maipo* | T | A | A | A | T | C | T | A | C | T | T | T | G | T | T | A | C | G | A | C | T | T |
| MK292115.1 | Coreoptera | *Pyrocoelia praetexta* | T | A | C | A | T | C | T | A | C | T | T | T | G | T | T | A | C | G | A | C | T | T |
| NC044790.1 | Coreoptera | *Pyrocoelia praetexta* | T | A | C | A | T | C | T | A | C | T | T | T | G | T | T | A | C | G | A | C | T | T |
| AF452048.1 | Coreoptera | *Pyrocoelia rufa* | T | A | C | A | T | C | T | A | C | T | T | T | G | T | T | A | C | G | A | C | T | T |
| NC003970.1 | Coreoptera | *Pyrocoelia rufa* | T | A | C | A | T | C | T | A | C | T | T | T | G | T | T | A | C | G | A | C | T | T |
| MK292117.1 | Coreoptera | *Pyrocoelia* | T | A | C | A | T | C | T | A | C | T | T | T | G | T | T | A | C | G | A | C | T | T |

| | | | | | | | | | | | | | | | | | | | | | | | | |
|---|---|---|---|---|---|---|---|---|---|---|---|---|---|---|---|---|---|---|---|---|---|---|---|---|
| | | *thibetana* | | | | | | | | | | | | | | | | | | | | | | |
| NC044792.1 | Coreoptera | *Pyrocoelia thibetana* | T | A | C | A | T | C | T | A | C | T | T | T | G | T | T | A | C | G | A | C | T | T |
| MK292111.1 | Coreoptera | *Vesta saturnalis* | T | A | A | A | T | C | T | A | C | T | T | T | G | T | T | A | C | G | A | C | T | T |
| NC044788.1 | Coreoptera | *Vesta saturnalis* | T | A | A | A | T | C | T | A | C | T | T | T | G | T | T | A | C | G | A | C | T | T |
| NC030249.1 | Diptera | *Anopheles bellator* | T | A | C | A | T | C | T | A | C | T | A | T | G | T | T | A | C | G | A | C | T | T |
| NC030717.1 | Diptera | *Anopheles janconnae* | T | A | C | A | T | C | T | A | C | T | A | T | G | T | T | A | C | G | A | C | T | T |
| MF381592.1 | Diptera | *Anopheles marajoara* | T | A | C | A | T | C | T | A | C | T | A | T | G | T | T | A | C | G | A | C | T | T |
| HQ335345.1 | Diptera | *Anopheles oryzalimnetes* | T | A | C | A | T | C | T | A | C | T | A | T | G | T | T | A | C | G | A | C | T | T |
| MF381722.1 | Diptera | *Culex bilineatus* | T | A | C | A | T | C | T | A | C | T | A | T | G | T | T | A | C | A | A | C | T | T |
| MF381718.1 | Diptera | *Culex lygrus* | T | A | C | A | T | C | T | A | C | T | A | T | G | T | T | A | C | A | A | C | T | T |
| MH316119.1 | Diptera | *Lutzia halifaxii* | T | A | C | A | T | C | T | A | C | T | A | T | G | T | T | A | C | A | A | C | T | T |
| MN626442.1 | Diptera | *Ochlerotatus taeniorhynchus* | T | A | C | A | T | C | T | A | C | T | A | T | G | T | T | A | C | A | A | C | T | T |
| KP721463.1 | Diptera | *Ochlerotatus vigilax* | T | A | C | A | T | C | T | A | C | T | A | T | G | T | T | A | C | A | A | C | T | T |
| MF957171.1 | Diptera | *Sabethes belisarioi* | T | A | C | A | T | C | T | A | C | T | A | T | G | T | T | A | C | A | A | C | T | T |
| KP793690.1 | Diptera | *Simulium* | T | A | C | T | T | C | T | A | C | T | A | T | G | T | T | A | C | G | A | C | T | T |

| | | | | | | | | | | | | | | | | | | | | | | | | |
|---|---|---|---|---|---|---|---|---|---|---|---|---|---|---|---|---|---|---|---|---|---|---|---|---|
| | | *aureohirtum* | | | | | | | | | | | | | | | | | | | | | | |
| NC029753.1 | Diptera | *Simulium aureohirtum* | T | A | C | T | T | C | T | A | C | T | A | T | G | T | T | A | C | G | A | C | T | T |
| MH729190.1 | Diptera | *Simulium maculatum* | T | A | C | T | T | C | T | A | C | T | A | T | G | T | T | A | C | G | A | C | T | T |
| NC040120.1 | Diptera | *Simulium maculatum* | T | A | C | T | T | C | T | A | C | T | A | T | G | T | T | A | C | G | A | C | T | T |
| MT410847.1 | Diptera | *Simulium noelleri* | T | A | C | T | T | C | T | A | C | T | A | T | G | T | T | A | C | G | A | C | T | T |
| NC050320.1 | Diptera | *Simulium noelleri* | T | A | C | T | T | C | T | A | C | T | A | T | G | T | T | A | C | G | A | C | T | T |
| KU252587.1 | Diptera | *Simulium variegatum* | T | A | C | T | T | C | T | A | C | T | A | T | G | T | T | A | C | G | A | C | T | T |
| NC033348.1 | Diptera | *Simulium variegatum* | T | A | C | T | T | C | T | A | C | T | A | T | G | T | T | A | C | G | A | C | T | T |
| JN861743.1 | Diptera | *Tipula abdominalis* | T | A | C | A | T | C | T | A | C | T | A | T | G | T | T | A | C | G | A | C | T | T |
| KT970065.1 | Diptera | *Tipula cockerelliana* | T | A | C | A | T | C | T | A | C | T | A | T | G | T | T | A | C | G | A | C | T | T |
| NC030520.1 | Diptera | *Tipula cockerelliana* | T | A | C | A | T | C | T | A | C | T | A | T | G | T | T | A | C | G | A | C | T | T |
| MT410842.1 | Diptera | *Tipula fascipennis* | T | A | C | A | T | C | T | A | C | T | A | T | G | T | T | A | C | G | A | C | T | T |
| NC050319.1 | Diptera | *Tipula fascipennis* | T | A | C | A | T | C | T | A | C | T | A | T | G | T | T | A | C | G | A | C | T | T |
| NC044661.1 | Diptera | *Trichoprosopon pallidiventer* | T | A | C | A | T | C | T | A | C | T | A | T | G | T | T | A | C | A | A | C | T | T |

| | | | | | | | | | | | | | | | | | | | | | | | | | |
|---|---|---|---|---|---|---|---|---|---|---|---|---|---|---|---|---|---|---|---|---|---|---|---|---|---|
| MK575491.1 | Diptera | *Uranotaenia geometrica* | | T | A | C | A | T | C | T | A | C | T | A | T | G | T | T | A | C | G | A | C | T | T |
| MK575492.1 | Diptera | *Wyeomyia confusa* | | T | A | C | A | T | C | T | A | C | T | A | T | G | T | T | A | C | A | A | C | T | T |
| | | | A | 0.0 | 100.0 | 15.4 | 94.9 | 0.0 | 0.0 | 0.0 | 100.0 | 0.0 | 0.0 | 52.8 | 0.0 | 0.0 | 0.0 | 0.0 | 100.0 | 0.0 | 3.7 | 100.0 | 0.0 | 0.0 | 0.0 |
| | | | T | 100.0 | 0.0 | 0.0 | 3.7 | 79.9 | 3.7 | 100.0 | 0.0 | 0.0 | 100.0 | 43.9 | 100.0 | 0.0 | 100.0 | 100.0 | 0.0 | 0.0 | 0.0 | 0.0 | 0.0 | 100.0 | 100.0 |
| | | | G | 0.0 | 0.0 | 0.0 | 1.4 | 0.0 | 0.0 | 0.0 | 0.0 | 0.0 | 0.0 | 0.0 | 0.0 | 100.0 | 0.0 | 0.0 | 0.0 | 0.0 | 96.3 | 0.0 | 0.0 | 0.0 | 0.0 |
| | | | C | 0.0 | 0.0 | 84.6 | 0.0 | 20.1 | 96.3 | 0.0 | 0.0 | 100.0 | 0.0 | 3.3 | 0.0 | 0.0 | 0.0 | 0.0 | 0.0 | 100.0 | 0.0 | 0.0 | 100.0 | 0.0 | 0.0 |

Table S13. Nucleotide sequences of universal primer (AQdb-12S_cad_R) and homologous region of 31 sequences of trichopteran insects used for development in this study.

| | | | | | | | | | | | | | | | | | | | | | | |
|---|---|---|---|---|---|---|---|---|---|---|---|---|---|---|---|---|---|---|---|---|---|---|
| AQdb-12S_cad_R | | | | A | R | Y | G | A | C | G | G | G | C | A | A | T | A | T | G | T | R | C |
| | | | A | 31 | 6 | 0 | 0 | 31 | 0 | 0 | 0 | 0 | 0 | 31 | 31 | 0 | 31 | 0 | 0 | 0 | 23 | 0 |
| | | | T | 0 | 0 | 23 | 0 | 0 | 0 | 0 | 0 | 0 | 0 | 0 | 0 | 31 | 0 | 31 | 0 | 31 | 0 | 0 |
| | | | G | 0 | 23 | 0 | 31 | 0 | 0 | 31 | 31 | 31 | 0 | 0 | 0 | 0 | 0 | 0 | 31 | 0 | 8 | 0 |
| | | | C | 0 | 0 | 8 | 0 | 0 | 31 | 0 | 0 | 0 | 31 | 0 | 0 | 0 | 0 | 0 | 0 | 0 | 0 | 31 |
| GenBank Acc. No. | Order | Species | | | | | | | | | | | | | | | | | | | | |
| KF756944.1 | Trichoptera | *Apatania* sp. | | A | A | T | G | A | C | G | G | G | C | A | A | T | A | T | G | T | G | C |
| MT410856.1 | Trichoptera | *Beraea pullata* | | A | G | T | G | A | C | G | G | G | C | A | A | T | A | T | G | T | A | C |
| MG669126.1 | Trichoptera | *Cheumatopsyche analis* | | A | G | C | G | A | C | G | G | G | C | A | A | T | A | T | G | T | A | C |
| NC036955.1 | Trichoptera | *Cheumatopsyche analis* | | A | G | C | G | A | C | G | G | G | C | A | A | T | A | T | G | T | A | C |
| KX385010.1 | Trichoptera | *Cheumatopsyche brevilineata* | | A | G | T | G | A | C | G | G | G | C | A | A | T | A | T | G | T | A | C |
| NC043769.1 | Trichoptera | *Cheumatopsyche brevilineata* | | A | G | T | G | A | C | G | G | G | C | A | A | T | A | T | G | T | A | C |
| MG669125.1 | Trichoptera | *Cheumatopsyche campyla* | | A | G | C | G | A | C | G | G | G | C | A | A | T | A | T | G | T | A | C |
| NC036954.1 | Trichoptera | *Cheumatopsyche campyla* | | A | G | C | G | A | C | G | G | G | C | A | A | T | A | T | G | T | A | C |
| MG669123.1 | Trichoptera | *Cheumatopsyche speciosa* | | A | G | C | G | A | C | G | G | G | C | A | A | T | A | T | G | T | A | C |
| NC036952.1 | Trichoptera | *Cheumatopsyche speciosa* | | A | G | C | G | A | C | G | G | G | C | A | A | T | A | T | G | T | A | C |

| | | | | | | | | | | | | | | | | | | | | | |
|---|---|---|---|---|---|---|---|---|---|---|---|---|---|---|---|---|---|---|---|---|---|
| KF717094.1 | Trichoptera | *Eubasilissa regina* | A | G | T | G | A | C | G | G | G | C | A | A | T | A | T | G | T | A | C |
| KF756943.1 | Trichoptera | *Eubasilissa regina* | A | G | T | G | A | C | G | G | G | C | A | A | T | A | T | G | T | A | C |
| NC023374.1 | Trichoptera | *Eubasilissa regina* | A | G | T | G | A | C | G | G | G | C | A | A | T | A | T | G | T | A | C |
| KX385011.1 | Trichoptera | *Hydatophylax nigrovittatus* | A | A | T | G | A | C | G | G | G | C | A | A | T | A | T | G | T | A | C |
| NC043770.1 | Trichoptera | *Hydatophylax nigrovittatus* | A | A | T | G | A | C | G | G | G | C | A | A | T | A | T | G | T | A | C |
| KF717095.1 | Trichoptera | *Hydromanicus wulaianus* | A | G | T | G | A | C | G | G | G | C | A | A | T | A | T | G | T | A | C |
| NC036156.1 | Trichoptera | *Hydromanicus wulaianus* | A | G | T | G | A | C | G | G | G | C | A | A | T | A | T | G | T | A | C |
| MT628580.1 | Trichoptera | *Hydropsyche angustipennis* | A | G | T | G | A | C | G | G | G | C | A | A | T | A | T | G | T | A | C |
| MG669122.1 | Trichoptera | *Hydropsyche orris* | A | G | T | G | A | C | G | G | G | C | A | A | T | A | T | G | T | A | C |
| NC036951.1 | Trichoptera | *Hydropsyche orris* | A | G | T | G | A | C | G | G | G | C | A | A | T | A | T | G | T | A | C |
| MG669121.1 | Trichoptera | *Hydropsyche simulans* | A | G | T | G | A | C | G | G | G | C | A | A | T | A | T | G | T | A | C |
| NC036950.1 | Trichoptera | *Hydropsyche simulans* | A | G | T | G | A | C | G | G | G | C | A | A | T | A | T | G | T | A | C |
| MK077681.1 | Trichoptera | *Limnephilus hyalinus* | A | A | T | G | A | C | G | G | G | C | A | A | T | A | T | G | T | G | C |
| NC044710.1 | Trichoptera | *Limnephilus hyalinus* | A | A | T | G | A | C | G | G | G | C | A | A | T | A | T | G | T | G | C |
| MG980616.1 | Trichoptera | *Phryganea cinerea* | A | G | T | G | A | C | G | G | G | C | A | A | T | A | T | G | T | G | C |
| NC039714.1 | Trichoptera | *Phryganea cinerea* | A | G | T | G | A | C | G | G | G | C | A | A | T | A | T | G | T | G | C |
| KX385012.1 | Trichoptera | *Phryganopsyche latipennis* | A | ? | T | G | A | C | G | G | G | C | A | A | T | A | T | G | T | G | C |
| NC043771.1 | Trichoptera | *Phryganopsyche latipennis* | A | ? | T | G | A | C | G | G | G | C | A | A | T | A | T | G | T | G | C |
| MG669124.1 | Trichoptera | *Potamyia flava* | A | G | C | G | A | C | G | G | G | C | A | A | T | A | T | G | T | A | C |

| | | | | | | | | | | | | | | | | | | | | | | |
|---|---|---|---|---|---|---|---|---|---|---|---|---|---|---|---|---|---|---|---|---|---|---|
| NC036953.1 | Trichoptera | *Potamyia flava* | | A | G | C | G | A | C | G | G | G | C | A | A | T | A | T | G | T | A | C |
| KP455291.1 | Trichoptera | *Thremma gallicum* | | A | A | T | G | A | C | G | G | G | C | A | A | T | A | T | G | T | G | C |
| | | | A | 100.0 | 20.7 | 0.0 | 0.0 | 100.0 | 0.0 | 0.0 | 0.0 | 0.0 | 0.0 | 100.0 | 100.0 | 0.0 | 100.0 | 0.0 | 0.0 | 0.0 | 74.2 | 0.0 |
| | | | T | 0.0 | 0.0 | 74.2 | 0.0 | 0.0 | 0.0 | 0.0 | 0.0 | 0.0 | 0.0 | 0.0 | 0.0 | 100.0 | 0.0 | 100.0 | 0.0 | 100.0 | 0.0 | 0.0 |
| | | | G | 0.0 | 79.3 | 0.0 | 100.0 | 0.0 | 0.0 | 100.0 | 100.0 | 100.0 | 0.0 | 0.0 | 0.0 | 0.0 | 0.0 | 0.0 | 100.0 | 0.0 | 25.8 | 0.0 |
| | | | C | 0.0 | 0.0 | 25.8 | 0.0 | 0.0 | 100.0 | 0.0 | 0.0 | 0.0 | 100.0 | 0.0 | 0.0 | 0.0 | 0.0 | 0.0 | 0.0 | 0.0 | 0.0 | 100.0 |
